# Conserved and specialized features of thalamocortical wiring revealed by single-cell projection mapping in mouse and marmoset

**DOI:** 10.64898/2026.07.07.736957

**Authors:** Angela Y. Fan, Jack T. Scott, Noah M. Kuehn, Maryam Majeed, Sabrina Y. Cheng, Huihui Qi, Mathew T. Summers, John Hover, Tye Johnson, Ben Ouellette, Alana A. Oyama, Diana Ravens, Huiqing Zhan, Justus M. Kebschull, James A. Bourne, Xiaoyin Chen

## Abstract

Across mammals, brain regions can duplicate, expand, and diversify, requiring long-range connectivity to accommodate species-specific specializations while preserving globally ordered wiring. This challenge is especially pronounced in primate thalamocortical circuits, where select cortical fields and their thalamic partners have expanded disproportionately. Although gene expression in the thalamus follows broad and conserved gradients, how thalamocortical projections are organized at single-neuron resolution, and how this organization is reshaped by expansion, remain unknown. Here, we investigate thalamocortical projection organization by in situ sequencing and BARseq projection mapping in marmosets and mice. We profiled the gene expression of 1.5 million marmoset neurons and jointly measured gene expression and cortical projections in 708 marmoset and 1,518 mouse neurons that spanned multiple thalamic nuclei. In both species, projections of individual neurons targeted diverse areas that together spanned a large fraction of the cortex. Comparing projections at the single-neuron level and local neighborhood level revealed that marmoset thalamocortical projections were more spatially segregated, producing a point-to-point architecture. Strikingly, this local specialization coexisted with a conserved gradient that predominated over discrete anatomical borders: In the higher-order sensory thalamus of both species, gene expression and projections varied continuously across nucleus borders, and borders had only a small effect on projections. Furthermore, in both marmoset and mouse, gene expression gradients were associated with the anteroposterior locations of cortical targets. These results reconcile discrete nucleus and gradient-based models of thalamic organization and suggest that primate circuit specialization is superimposed on a conserved molecular-projection gradient.

## Introduction

Primate brains are distinguished by highly elaborated thalamocortical pathways, particularly those connecting higher-order thalamic nuclei to specialized cortical areas. Compared with those in rodents, the prefrontal cortex (PFC), the visual association cortex, and their thalamic partners are much larger and more diverse. The medial pulvinar, for example, is likely a primate-specific enlargement of the pulvinar complex that has an increased complexity of connectivity with the association and prefrontal cortex across primate evolution (Baldwin and Bourne, 2020; Córdoba-Claros et al., 2025a). The lateral geniculate nucleus (LGN), while present in all mammals, exhibits a far more complex architecture in primates, and is partitioned into laminae that physically segregate inputs by eye and cell class. These changes create a fundamental wiring problem: the connections of individual neurons must specialize to accommodate the species-specific needs that accompanied expansion, yet the thalamus preserves an ordered map of connectivity across the whole cortex. How the thalamus can specialize at the single neuron level without disrupting its orderly connectivity to the whole cortex remains unknown.

Resolving thalamocortical projections at single-neuron resolution could reveal how the thalamus is organized and how it accommodates cortical expansion. Traditionally, thalamic projections are assigned into discrete anatomical nuclei, each defined by a distinct combination of cytoarchitecture, connectivity, and function (Jones, 1998; Sherman, 2007). Recent studies, however, suggest that transcriptional patterns (Huang et al., 2026; J. W. Phillips et al., 2019), developmental lineage (Puche-Aroca et al., 2025), and activity (Petty and Bruno, 2024) are organized in gradients across thalamic nuclei. These two models predict different possibilities by which the thalamus scales. If projections are discrete across nuclei, as traditional theories dictate, serving new cortical areas would likely require adding or subdividing nuclei. If projection logic is gradient-based, new areas can be accommodated by stretching, shifting, or locally sharpening an existing topographical system. Determining which of these models dominates, and whether the dominant model differs by species, requires resolving thalamocortical projections and transcriptomic identity in the same cells at single-neuron resolution.

Spatial transcriptomics and single-neuron tracing have revealed thalamic gene-expression patterns and projection motifs separately (Dan et al., 2025; Gou et al., 2025; Huang et al., 2026; Winnubst et al., 2019; S. Yao et al., 2023), but combining transcriptomic identity, anatomical position, and long-range projections at cellular resolution and at scale remains challenging. We addressed this using BARseq (Chen et al., 2019, 2022; Muñoz-Castañeda et al., 2021; Sun et al., 2021), an in situ sequencing approach that simultaneously captures transcriptomic identity and long-range projections in thousands of individual neurons within a single animal. BARseq reads out projections by sequencing barcodes that uniquely label neurons in situ, thus allowing projections and gene expression to be determined across many neurons at the same time.

Here, we profiled gene expression in over 1.5 million common marmoset thalamic neurons, and jointly mapped projections and transcriptomic identity of 708 marmoset and 1,518 mouse thalamocortical projecting neurons. Because the data resolve projections and gene expression in the same individual neurons spanning nucleus borders in both species, it allows us to ask whether transcriptomic and projection gradients are conserved, how they relate to traditional nucleus borders, and how the larger primate brain reorganizes thalamocortical projections. Our analyses reveal that gradient organization is conserved across both species and coexists with species-specific projection differences at a more local level.

## Results

We first investigated the transcriptomic organization of the marmoset thalamus using in situ sequencing, and compared transcriptomic cell-type composition and spatial gene expression gradients with those of the mouse from the Allen Brain Cell (ABC) atlas (Z. Yao et al., 2023). We next mapped thalamocortical projections in both species using BARseq, allowing us to interpret connectivity directly in the context of transcriptomic organization. Throughout this manuscript, we use “nucleus” to refer to structures such as the lateral geniculate nucleus (LGN in marmoset and LGd in mouse), posterior thalamic nucleus group (PO), mouse lateral posterior nucleus (LP), and the divisions of the primate pulvinar (the medial pulvinar MPul, lateral pulvinar LPul, and the inferior pulvinar IPul); we use “subdivisions” to refer to a parcellation within a nucleus, such as the medial and lateral subdivisions of the LP (LPm and LPl). A complete list of abbreviations for all anatomical areas is provided (**Supp. Table 1**).

### In situ sequencing reveals transcriptomic heterogeneity of the marmoset thalamus

The transcriptomic organization of the thalamus has been extensively profiled in the mouse (Z. Yao et al., 2023; Zhang et al., 2023), but spatial transcriptomic studies in the primate thalamus have so far focused on non-single-cell resolution analysis (Huang et al., 2026; Schulmann et al., 2024) or relied on targeted techniques that utilized gene panels designed for other brain regions (Dan et al., 2025). To enable direct comparison with the transcriptomic organization of the marmoset thalamus and the mouse ABC atlas, we first set out to produce a spatial transcriptomic dataset tailored to reveal both discrete and continuous transcriptomic variation in the marmoset thalamus. We used BARseq-style in situ sequencing (Chen et al., 2025)(**Fig. 1a-b**; see **Methods**) to profile the expression of 152 genes (**Supp. Table 2**) on ten marmoset hemibrain coronal sections, spanning from the posterior end of the pulvinar to the anterior ventral thalamic groups (AP +1.80 mm to +8.00 mm)(Paxinos, 2012) (**Fig. 1c**). The gene panel combined marker genes for cell types defined in reference snRNA-seq datasets (Dan et al., 2025), genes that captured major axes of variations, classic anatomical marker genes, and additional genes of interest, yielding more discriminative power for the heterogeneity in the thalamus relative to existing panels designed for other brain regions (see **Supp. Note 1** and **ED Fig. 1** on panel design and in silico benchmarking). After removing low-quality cells (see **Methods**), the dataset contained 1,510,192 cells with 91.7 ± 53.4 (mean ± standard deviation) reads per cell and 40.7 ± 13.4 (mean ± standard deviation) unique genes per cell (**Fig. 1d**).

**Figure 1.**
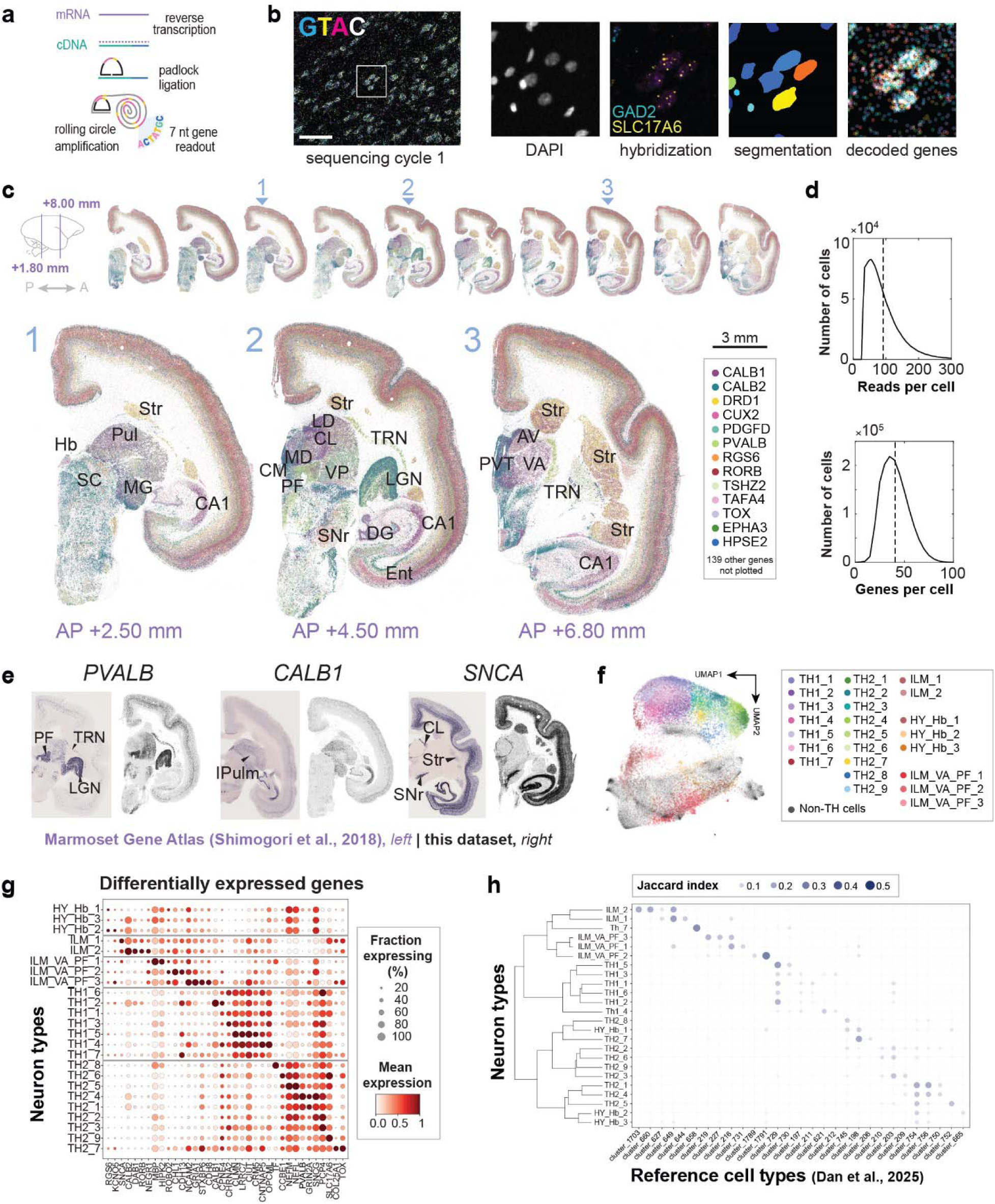
In situ sequencing reveals transcriptomic heterogeneity in the marmoset thalamus. (**a**) Schematic describing detection and amplification of target mRNAs for BARseq-style in situ sequencing. (**b**) An image of the first gene-sequencing cycle from a representative slice. The boxed area corresponds to the images on the right, showing a DAPI staining, a hybridization round that detects two highly expressed genes, the cell segmentation mask, and decoded genes from all sequencing cycles. Scale bar = 100 µm. (**c**) mRNA locations across 10 slices (*top*) and close-up images of three representative slices (*bottom*). On the brain icon, the two lines indicate the A-P span of our sampled region. For clarity, 13 out of 152 genes are plotted. AV, anteroventral thalamic nucleus; CA1, Cornu Ammonis 1 of the hippocampus; CL centrolateral thalamic nucleus; CM centromedial thalamic nucleus; DG dentate gyrus; Ent, entorhinal cortex; Hb, habenula; LD, lateral dorsal nucleus; LGN, lateral geniculate nucleus; MD, mediodorsal nucleus; MG, medial geniculate nucleus; PF, parafascicular nucleus; Pul, pulvinar; PVT, paraventricular thalamus; SC, superior colliculus; SNr, substantia nigra; Str, striatum; TRN, thalamic reticular nucleus; VA, ventral anterior thalamic nucleus; VP, ventroposterior thalamic nucleus. See **Supp. Table 1** for complete list of anatomical abbreviations used in this manuscript. Scale bar = 3 mm. (**d**) Histograms showing the distribution of read counts per cell (*top*) and gene counts per cell (*bottom*) after quality control. Dashed lines indicate medians. (**e**) Expression patterns of representative genes from the Marmoset Gene Atlas (Shimogori et al., 2018) (*left*), compared to the current dataset (*right*). (**f**) UMAP plot of gene expression of thalamic excitatory neurons, colored by thalamic neuron types. Additional hypothalamic and midbrain neurons that co-clustered with thalamic neurons at a coarser hierarchy level are shown in grey. (**g**) Expression of differentially expressed genes (x-axis) in thalamic excitatory neuron types (y-axis). Colors indicate mean expression level, and dot size indicates the fraction of cells expressing each gene. (**h**) Overlap (Jaccard index) between thalamic excitatory neuron types in this dataset (y-axis) and clusters in a reference snRNA-seq dataset (Dan et al., 2025). The dendrogram (*left*) shows the hierarchical relationship between neuron types in this dataset inferred from the mapping.

Many genes, including classical anatomical markers, were differentially expressed across major brain structures and their divisions, such as cortical layers and thalamic nuclei (**Fig. 1e**). Parvalbumin (PVALB), a well-established marker of thalamic nuclei, was highly expressed in the lateral geniculate nucleus (LGN) and the thalamic reticular nucleus, and in cells scattered throughout the cortex. Calbindin (*CALB1*), a classical negative marker of the medial portion of the inferior pulvinar (IPulm) (Baldwin and Bourne, 2020; Cusick et al., 1993), was highly expressed in the pulvinar except for the IPulm. Alpha synuclein (*SNCA*) was enriched in deep cortical layers, substantia nigra, the striatum, and the central lateral nucleus. Collectively, the spatial expression patterns of these canonical markers were consistent with those reported in the Marmoset Gene Atlas (Kita et al., 2021; Shimogori et al., 2018).

To examine differences in gene expression across thalamic nuclei, we performed hierarchical clustering to identify transcriptomically defined cell types, a common approach for exploring heterogeneity in single-cell transcriptomic data. We clustered neurons at three hierarchical levels to yield 24 excitatory thalamic neuron types (**Fig. 1f**, **ED Fig. 2**, **Supp. Note 2**). Anatomical divisions were defined by the mRNA expression of classic histological markers, including *CALB1* and *PVALB* (Jones, 1998), and by the distribution of region-specific clusters (**ED Fig. 3**, **Supp. Note 3**). As thalamocortical projection neurons are exclusively excitatory, subsequent analysis focused on these 24 neuron types.

Genes that encode classic histological markers were also differentially expressed across neuron types. Notably, *CALB1* and *PVALB*, markers that are classically associated with “core” and “matrix” thalamus (Jones, 2001), were highly enriched in the TH1 and TH2 groups, respectively (**Fig. 1g**). This differential expression was also consistent with the preferential enrichment of TH1 and TH2 neurons in first-order and higher-order thalamic nuclei, respectively (**Supp. Note 2**; **ED Fig. 2i**). To validate our neuron types against an independent dataset, we applied a k-nearest neighbor (KNN) classifier (Chen et al., 2025) to assign cell type labels from a reference snRNA-seq dataset (Dan et al., 2025) to individual neurons, and calculated the overlap with our neuron types. Most of the neuron types mapped to a small number of reference cell types (**Fig. 1h**), demonstrating that our dataset recapitulated the transcriptomic diversity observed in reference snRNA-seq datasets.

### Higher-order thalamus contains both discrete and continuous gradient organization of gene expression in mouse and marmoset

In the non-sensory thalamus, neuron types were largely confined within individual thalamic nuclei. For example, type TH1_7 was localized to the LD, and ILM_VA_PF_3 and HY_Hb_1 types were preferentially enriched in the ventroanterior magnocellular (VAmc) and anterior dorsal (AD) nuclei, respectively (**Fig. 2a**). Therefore, non-sensory thalamic nuclei can be distinguished by spatially confined neuron types.

**Figure 2.**
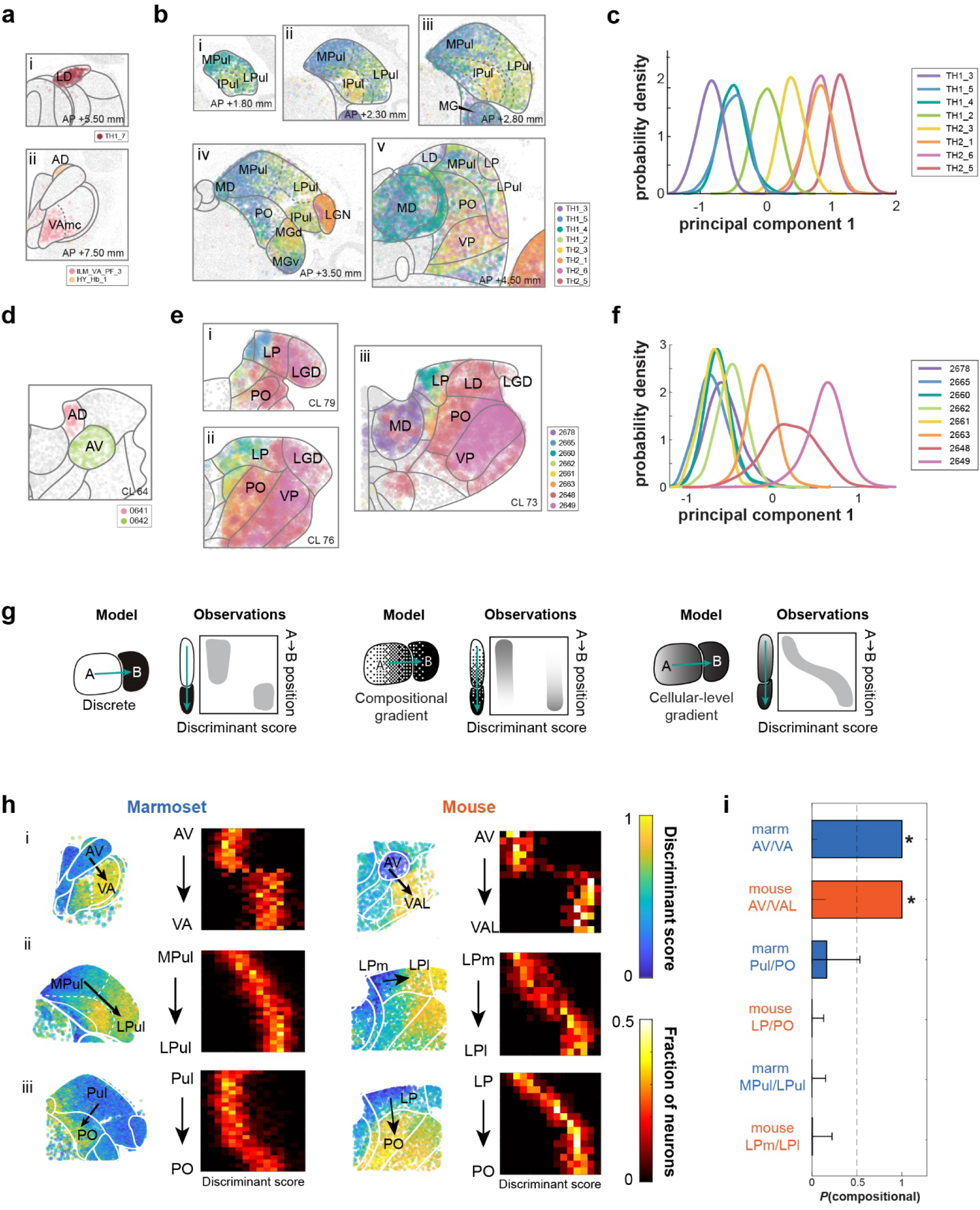
Spatial distribution of neuron types across thalamic nuclei. (**a**)-(**b**) Distribution of the indicated neuron types in the marmoset thalamus. Delineations indicate nucleus/subdivision borders (see **Supp. Note 3** for nucleus delineation criteria). (**c**) Probability density distribution of the indicated marmoset neuron types along the first principal component in the gene expression space of these neuron types. (**d**)-(**e**) Distribution of the indicated neuron types in the mouse thalamus. Delineations indicate nucleus/subdivision borders from CCF. (**f**) Probability density distribution of the indicated mouse neuron types along the first principal component in the gene expression space of these neuron types. (**g**) Three models of gene expression organization in space (*left*), with expected patterns of distributions of the gene discriminant score along the spatial axis across the two nuclei (*right*). The y-axis indicates neuron locations along the green arrows that point from the centroid of nucleus A to the centroid of nucleus B; the x-axis indicates scores along the gene discriminant vector that discriminates neurons of nuclei A and B. (**h**) Gene discriminant scores across pairs of nuclei in marmoset (*left column*) and mouse (*right column*). For each nucleus/subdivision pair, the left plot shows neurons in the thalamus colored by the discriminant scores; the right plot shows the distribution of discriminant scores (x-axis) as a function of position along the spatial vector connecting the centroids of the two nuclei/subdivisions (y-axis). Heatmap indicates the fraction of neurons at each position along the spatial vector (i.e., rows sum to 1). **(i)** Probability that the discriminant-score distribution within each transition zone is bimodal [P(compositional)]; dashed line, decision threshold. Scale bars, standard deviation from resampling. Colors indicate species. * p ≤ 0.012 after Bonferroni correction.

In contrast, in the sensory thalamus, multiple neuron types tiled nuclei in a spatially ordered sequence that extended from the higher-order thalamic nuclei into their corresponding first-order nuclei. In the pulvinar nuclei, for example, a series of neuron types spanned the anteromedial to posterolateral axis (**Fig. 2b**) in the same order as their positions along the first principal component of the gene expression of this population of neurons (**Fig. 2c**), suggesting that this spatial tiling reflects an underlying continuous gene expression gradient that dominates transcriptomic heterogeneity. Strikingly, the same sequence of neuron types appeared repeatedly across nuclei of different sensory modalities: from posteroventral to anterodorsal in the MG, and from posteromedial to anterolateral in the PO (**Fig. 2b**). In each sensory modality, these sequences of neuron types also appeared to extend into the corresponding first-order nuclei, but usually with a sharper anatomical transition (e.g., at the LGN/LP border) that could reflect modality-specific specializations in development (Frangeul et al., 2016). The recurrence of this sequence across functionally distinct nuclei indicates that the dominant axis of transcriptomic variation runs along a shared gradient rather than across nuclei.

To investigate whether the spatial patterns of gene expression are conserved across species, we reanalyzed the mouse thalamus MERSCOPE data from the Allen Brain Cell (ABC) atlas (Z. Yao et al., 2023). Similar to the patterns found in the marmoset thalamus, neuron types in nuclei that were localized anteriorly and/or medially were often confined to anatomical borders (e.g., types 0641 and 0642 in the AD and AV, respectively) (**Fig. 2d**). In contrast, those in the higher-order sensory thalamus were distributed across borders; for example, types 2678, 2665, 2660, 2662, 2661, 2663, 2648, 2649 were sequentially enriched across the LP, the mouse homolog of the marmoset Pul, from the medial end to lateral end. Like the marmoset neuron types, these neurons extended beyond the borders of LP into neighboring nuclei, such as the PO (**Fig. 2e**), and they were arranged in a similar order from VP to PO to MD. In gene expression space, these neuron clusters were similarly arranged in the same order as they were spatially (**Fig. 2f**). These observations indicate that the transcriptomic gradient organization in the sensory thalamus is conserved across mouse and marmoset.

We then investigated the nature of the gradient organization in the higher-order sensory (HOs) thalamus. Sequential tiling by neuron types in the HOs thalamus could reflect one of two possibilities: a gradient in the relative proportion of discrete neuron populations (a compositional gradient), or a gradient in gene expression across individual neurons (a cellular level gradient) (**Fig. 2g, middle and right**). This contrasts with the non-sensory thalamus, where nucleus-confined cell types likely reflect discrete organization (**Fig. 2g, left**). Distinguishing these possibilities requires examining gene expression continuously rather than through discrete clusters, since clustering forces cells into categories. We therefore used a cluster-free approach: for each pair of neighboring thalamic nuclei or subnuclei, we identified a gene discriminant vector that best separated neurons from the two regions, then examined how discriminant scores varied with neurons’ physical locations (**Methods**). In regions with distinct neuron types, such as the anteroventral nucleus (AV) and the ventral anterior (VA)/ventral anterior-lateral complex (VAL), scores shifted abruptly from one peak to another across the border (**Fig. 2h**; see **ED Fig. 4a** for robustness to read counts). In contrast, within the HOs thalamus, the score distribution shifted gradually both in nuclei of the same sensory modality (e.g., mouse LPm to LPl and marmoset MPul to LPul) and across nuclei of different sensory modalities (e.g., mouse LP to PO and marmoset Pul to PO) (**Fig. 2h**); these gradients were distinguishable from synthetic compositional gradients (**ED Fig. 4b**). To examine whether this gradient component masked a less dominant gene axis that was discrete between the two pairs of nuclei, we also examined the distribution of top principal components (**ED Fig. 4c, d**). No principal component showed the two-peaked distribution expected of a discrete component.

To further confirm that the HOs transitions were cellular-level gradients, we tested the discriminant score distributions in each transition zone for bimodality, using both a Gaussian mixture model comparison and Silverman’s test for multimodality (**Fig. 2i**; see **Methods**). Consistent with the distribution heatmaps, the AV/VA (marmoset) and AV/VAL (mouse) borders were strongly bimodal in both species [*P*(compositional) = 1.0; Silverman’s *p =* 0.012 after Bonferroni correction], indicating genuinely discrete boundaries. In contrast, the marmoset MPul/LPul and Pul/PO transitions and the mouse LP/PO and LPm/LPl transitions all favored the single-peak bridge model [*P*(compositional) ≤ 0.16 for all four pairs] with no evidence of multimodality (Silverman’s *p =* 0.285 – 0.683, raw values). These results extend previously described mesoscale gradients (Huang et al., 2026; J. W. Phillips et al., 2019) to the single-cell level and establish continuous cellular-level gradients, rather than nucleus identity, as the conserved and dominant organizing principle of transcriptomic heterogeneity across the HOs thalamus.

### BARseq reveals single-cell projection patterns of mouse thalamocortical neurons

The cellular-level transcriptomic gradients we observed across borders of nuclei stand in contrast to the traditional view of the thalamus as discrete nuclei (Jones, 2007; Sherman et al., 2006), which was, in part, defined by distinct projection patterns. Whether neurons projecting to different cortical areas are segregated by anatomical borders or are intermingled across borders remains unresolved (**Fig. 3a**). Distinguishing discrete transitions from gradients across borders requires sufficiently dense sampling of single neurons near borders. BARseq fulfills these requirements by mapping single-cell projections densely within a single brain, and relating projections to gene expression. Here, we apply BARseq to the mouse and marmoset HOs thalamus to identify conserved and species-specific features in thalamocortical circuits.

**Figure 3.**
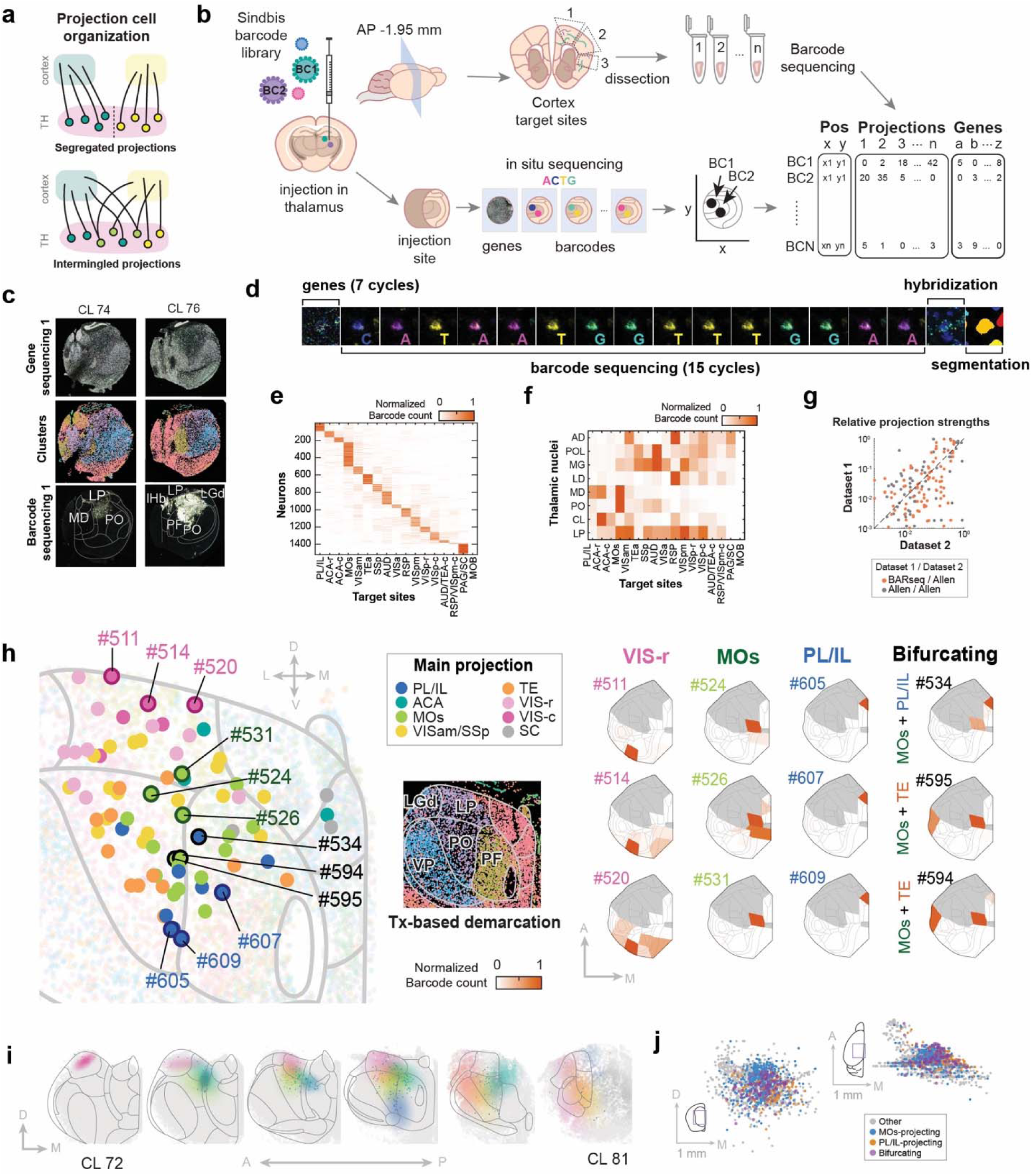
BARseq reveals single-cell projection patterns from the mouse thalamus. (**a**) Models of projection organization. Thalamic neurons either form discrete projection populations separated by a sharp border (*top*) or transition gradually between neighboring projection populations (*bottom*). (**b**) Schematic of mouse BARseq experimental workflow. (**c**) Two representative thalamic sections showing Sindbis virus injection sites. *Top*, merged image of gene rolonies from gene sequencing cycle 1. *Middle*, cells colored by coarse-level transcriptomic clusters, revealing gene expression differences across major thalamic nuclei. *Bottom*, merged image of barcoded neurons from barcode sequencing cycle 1. Major thalamic nuclei are labeled. (**d**) Representative images of in situ sequencing at a barcoded soma. Shown are the first gene sequencing cycle, 15 barcode sequencing cycles, one hybridization cycle, and the cell segmentation masks. (**e**) Projection matrix of all barcoded thalamic neurons from two mouse brains. Rows indicate individual neurons, columns indicate dissection cubelets. Colors represent row- and spike-in-normalized barcode counts. (**f**) Mean normalized barcode counts from each thalamic nucleus (y-axis) to each target domain (x-axis). (**g**) Fraction of total projections in the BARseq dataset (y-axis, *orange dots*) compared to the Allen Mouse Brain Connectivity Atlas (x-axis). Grey dots show multiple replicate Allen Atlas experiments targeting the same thalamic nuclei, included to measure within-dataset variability. (**h**) Example projection neurons on a representative section. *Left*, barcoded neurons are colored by their dominant target domain, overlaid on non-barcoded neurons colored by coarse-level transcriptomic cluster (background). The inset shows the same non-barcoded neurons colored by coarse-level clusters to indicate correspondence with nucleus boundaries. *Right*, flatmaps showing cortical projection patterns of representative single neurons projecting to each domain, and of three bifurcating neurons. The intensity of color represents the normalized barcode counts in each area. (**i**) Estimated projection density across six thalamic sections from anterior to posterior. Colors correspond to target domains in (h). (**j**) Soma locations of neurons projecting to MOs, PL/IL, or bifurcating to both, shown in two views. On the brain icons, the boxes indicate the location of the extent of the plotted region.

We first injected two mice with a barcoded Sindbis virus library at two sites per brain (**ED Fig. 5a**), one at the posterolateral portion of LP near the LP/PO border, and one at the medial portion of LP near the LP/CL/MD border. We then sequenced barcodes alongside mRNAs of 188 cell type marker genes in situ (**Fig. 3b-d**). We hierarchically clustered the neurons based on gene expression and validated clusters by mapping to reference cell types (Z. Yao et al., 2023) (**Supp. Note 4** and **ED Fig. 5b**). We then determined nucleus borders using both the expression of individual genes and coarse-level clusters (**Fig. 3c**, **ED Fig. 5c**), enabling assignment of neurons to nuclei with high accuracy. To sequence barcodes at cortical projection sites, we dissected out 17 target areas, or “cubelets,” comprising 15 cortical areas, the superior colliculus, and the olfactory bulb (negative control). All dissected target areas were registered to the Allen Common Coordinate Framework v3 (Wang et al., 2020) (CCFv3; see **Methods** and **ED Fig. 6a**) to enable comparison with external datasets and to improve future utility. Matching soma barcodes to those recovered from cortical projection sites yielded 1,518 uniquely barcoded projection neurons (**Fig. 3e**). To assess whether barcode diversity was sufficient to distinguish projections at cellular resolution, we compared the number of matching barcodes to a shuffled control. By chance, 0.2% of shuffled barcodes matched (see **Supp. Note 4**, **ED Fig. 6b, c** for details), indicating that the number of barcode-matched projections in our dataset was unlikely due to chance. Neurons from the two brains were similar in projection patterns (**ED Fig. 6d**) and innervated all dissected cortical areas except the olfactory bulb, which served as a negative control. LP contained the highest number of mapped neurons (443 of 1,518), and the remainder was distributed principally across MG, MD, LD, PO, POL, and AD (**ED Fig. 6e**).

To test whether our BARseq data were consistent with known connectivity patterns in the thalamus, we first estimated projection strengths from each thalamic nucleus to each target area by pseudobulking the total barcode counts (**Fig. 3f**) and compared them with the Allen Mouse Brain Connectivity Atlas (Harris et al., 2019) **(Fig. 3g, *orange dots*).** To establish a baseline for how well projections from two independent injections in different brains are expected to match, we also compared multiple experiments in the Allen dataset targeting the same thalamic nuclei (**Fig. 3g, *grey dots***). The BARseq-derived pseudobulk projection strengths correlated with the Allen dataset (Pearson *r =* 0.62, *p =* 3×10^-14^) to a similar degree as did injections targeting the same thalamic nuclei within the Allen dataset (Pearson *r =* 0.64, *p =* 5×10^-14^), indicating that our dataset captured differences in projection patterns across nuclei as consistently as conventional tracing.

We next asked whether BARseq data captured projection diversity at single-cell resolution by comparing it with single-cell tracing (scTracing) data from the MouseLight (Winnubst et al., 2019) and SEU-Allen (Jiang et al., 2025; Peng et al., 2021) datasets. We pooled all available scTracing neurons with somas in CL, LP, MD, MG, PO, and POL and with reconstructed cortical axons, yielding 99 neurons across the two datasets. scTracing neurons from the same nuclei as BARseq neurons resembled them in cortical terminal fields (**ED Fig. 7a**). For example, in both BARseq and scTracing datasets, we found LP neurons that projected focally to VISp, innervated RSP broadly, or bifurcated to distinct frontal areas (**ED Fig. 7a**, neurons #511, #113 and #117). To compare BARseq and scTracing data systematically, we randomly subsampled BARseq neurons to match the sample size of the scTracing dataset (n = 99 per dataset), combined the datasets at BARseq’s projection resolution, and co-clustered neurons by their projections. Although the two tracing datasets sampled neurons with distinct cortical projections (**ED Fig. 7b**), co-clustering nonetheless yielded eight clusters, each containing both BARseq and scTracing neurons (**ED Fig. 7c**). Notably, clusters 2 and 8 contained only scTracing neurons from MouseLight or SEU-Allen datasets, respectively, yet both contained BARseq neurons (**ED Fig. 7d**), indicating that the BARseq data were consistent with the union of the two scTracing datasets. Consistent with the co-clustering results, t-SNE of the combined data placed BARseq neurons throughout the full projection space, whereas the two tracing datasets occupied partially distinct regions (**ED Fig. 7e**). Together, these results demonstrate that BARseq captured projection patterns consistent with published scTracing datasets at single-cell resolution.

### Mouse thalamocortical projections are intermingled across borders

The relatively dense labeling in our dataset allowed us to examine the local organization of projection neurons at single-cell resolution. Because each neuron projected to multiple cortical targets (**ED Fig. 7f, g**) and specific combinations of targets were commonly co-innervated (**ED Fig. 7h**), a neuron’s projections could often be approximated as the sum of a few recurring components. We therefore decomposed each neuron’s projection pattern into a sum of eight “target domains” using non-negative matrix factorization (NMF) (Lee and Seung, 1999) (**ED Fig. 7i**; see **Supp. Note 5** for details and the number of target domains), a common processing approach for BARseq data that results in a compact and lower-noise representation of projection patterns (Chen et al., 2019; Sun et al., 2021).

Using this representation, we first examined how projections to different target domains were spatially distributed in individual slices. For example, in a single 20 µm section, we mapped the projections of 79 neurons that spanned LP, PO, and PF (**Fig. 3h**). Neurons were arranged along the dorsal-ventral axis by cortical projection target (**Fig. 3h**): visual domains (VIS-r, VIS-c) were represented at the dorsal pole with ACA at the medial tip; TE and MOs occupied an intermediate band laterally and medially, respectively; and PL/IL was found most ventrally. To examine how this organization varies along the anterior-posterior axis, we collapsed five-section stacks spanning 100 µm into single planes, then estimated the spatial probability of projection to each projection domain using Gaussian-kernel-smoothed Bernoulli models (**Fig. 3i**). Across neighboring stacks, the positions of projection neurons shifted gradually, but their relative arrangement remained largely stable: visual cortex-projecting neurons were consistently dorsal, ACA and PL/IL-projecting neurons were medial and/or ventral, and neurons projecting to motor, somatosensory, and sensory association areas largely occupied an intermediate zone.

Although nucleus borders were thought to separate distinct projection patterns, we found that neurons with different projections were intermingled regardless of classical thalamic boundaries. For example, in the coronal section shown, neurons projecting to VIS-r, VISam/SSp, TE, and MOs were present on both sides of the LP/PO border, and MOs- and PL/IL-projecting neurons were found in both PO and PF. The PO/PF case is particularly striking: PF is transcriptomically distinct from the HOs thalamus and has a clear transcriptomic border with PO (**Fig. 3h, *inset***), yet neurons projecting to the same cortical targets were found on both sides. In addition, some neurons (e.g., neurons #534, #594, and #595) projected to two non-adjacent cortical targets, suggesting that their projections bifurcate. These bifurcating neurons were located between single-target-projecting populations (**Fig. 3j**). The presence of bifurcating neurons and the spatial intermingling of neurons with different projections suggest that no border can discretely segregate these projection populations. Thus, these results argue against a strictly discrete nucleus partition of projection classes.

### BARseq reveals single-cell projection patterns of marmoset thalamocortical neurons

Primates possess substantially expanded higher-order visual, temporal, and prefrontal areas relative to mice. Accommodating this expanded cortical territory is likely to require corresponding changes in thalamocortical wiring, yet it is unknown whether and how single-neuron projection patterns differ across species. We therefore applied BARseq to map the projections of 708 neurons from two animals, in the marmoset pulvinar (Pul), mediodorsal nucleus (MD), posterior nucleus (PO), and adjacent thalamic nuclei, which innervate these expanded areas (Córdoba-Claros et al., 2025b; Roberts et al., 2007; Romanski et al., 1997)

To assess how well BARseq can distinguish single-cell projections in the marmoset, we first performed a pilot experiment to map MD projections. Retrograde tracers and bulk tract tracing have established the broad topography of MD-to-PFC projections in macaque (Goldman-Rakic and Porrino, 1985; J. M. Phillips et al., 2019) and marmoset (Roberts et al., 2007), but how individual MD neurons distribute their axons across PFC remains unclear. We injected a barcoded Sindbis virus library bilaterally into the MD in one animal (**Fig. 4a**; **ED Fig. 8a, b**), sequenced the barcodes in the somas in situ (**Fig. 4b**), and matched them to barcodes recovered from dissected cubelets spanning the prefrontal cortex (**ED Fig. 8c**; **Supp. Table 4**), yielding 115 (right hemisphere) and 203 (left hemisphere) MD-to-PFC projection neurons (**ED Fig. 8d**; **Supp. Note 6**). We pooled cubelets to identify vlPFC, dlPFC, mPFC, and oPFC projections. Neurons that projected to the different PFC regions were enriched at different locations both along the AP axis and within a coronal plane. Neurons in the posterior MD (AP +4.3 mm) projected predominantly to the dlPFC (**Fig. 4c**). More anteriorly (AP +5.0 mm), vlPFC-, dlPFC-, mPFC-and oPFC-projecting neurons occupied ventral, lateral, dorsal and medial MD, respectively (**Fig. 4c**). At this resolution, most neurons targeted one of the PFC regions (**ED Fig. 8d**), and even those projecting to multiple PFC regions predominantly targeted neighboring ones. These results were consistent with previous tracing and imaging studies in marmoset (Roberts et al., 2007) and macaque (Goldman-Rakic and Porrino, 1985; J. M. Phillips et al., 2019), and further show that this topography arises from neurons that focally target specific regions of the PFC.

**Figure 4.**
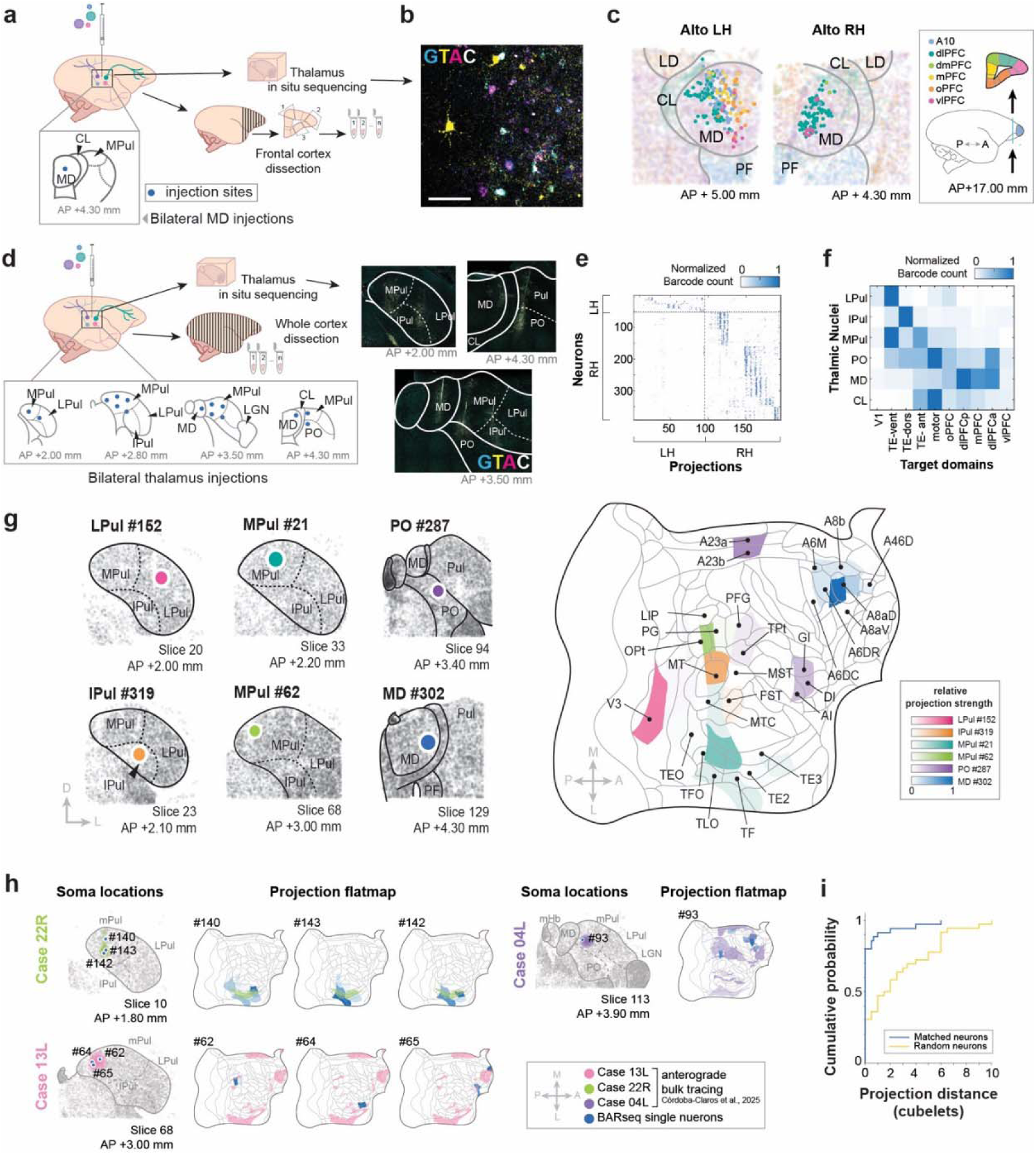
BARseq reveals single-cell projection patterns from the marmoset thalamus. (**a**) Schematic of the BARseq workflow for the MD pilot experiment (animal Alto). A barcoded virus library was injected into the MD bilaterally. The frontal cortex was dissected for sequencing, and the thalamus was sectioned for in situ sequencing. (**b**) Representative image from the second barcode sequencing cycle at the injection site. Scale bar = 100 µm. (**c**) Barcoded cells in the MD, colored by their dominant projection to each PFC subdivision. Non-barcoded cells in the background are colored by coarse-level transcriptomic cluster. The target areas that each color corresponds to are drawn on the right. LH, left hemisphere; RH, right hemisphere. (**d**) Schematic of the BARseq workflow for the full marmoset experiment (animal Hazelnut). A barcoded virus library was injected into the MD and Pul bilaterally, the whole cortex was dissected for sequencing, and the thalamus was sectioned for in situ sequencing. (**e**) Projection matrix showing the normalized projection strengths to dissected cubelets (x-axis) for each neuron (y-axis). Both axes are sorted by hemisphere. (**f**) Mean barcode counts of neurons from each thalamic nuclei (y-axis) to each target domain (x-axis), normalized by rows. (**g**) Projection patterns of representative neurons. *Top*, soma locations of each representative neuron, with section numbers and corresponding AP coordinates (Paxinos, 2012) indicated. *Bottom,* cortical projection patterns of each neuron visualized on a flatmap. Colors correspond to soma colors shown at the top. (**h**) Select single-neuron BARseq projection patterns overlaid on bulk anterograde tracing data (Córdoba-Claros et al., 2025b). *Left*, soma locations of each neuron, with section numbers and corresponding AP coordinates indicated. Colored shading indicates injection sites from bulk anterograde tracing cases; blue dots indicate BARseq neuron somas overlapping with injection sites. *Right*, cortical projection patterns visualized on a flatmap. Blue shading indicates BARseq single-neuron projection patterns; colored shading indicates bulk anterograde tracing projection zones. (**i**) Cumulative probability distribution of the distance (in dissection cubelets) between tracer-labelled projection zones and projections of matching (*blue*) or random (*yellow*) BARseq neurons. The x-axis indicates the number of dissection cubelets that are between the centroid of a BARseq neuron projection and the closest tracer-labelled region.

We next applied BARseq to two additional hemispheres to map projections and gene expression across the pulvinar, MD, and adjacent nuclei. We injected at 16 sites per hemisphere to achieve broad coverage of these regions (**Fig. 4d**) and dissected each cortical hemisphere into 94 cubelets (**Supp. Table 4**). Dissection cubelets were annotated on a marmoset magnetic resonance imaging (MRI) template (Liu et al., 2021) (**ED Fig. 9a**), and projected onto a cortical flatmap (**ED Fig. 9b, c**) that had been registered to a stereotaxic atlas (Paxinos, 2012) (**ED Fig. 9d**). In situ sequencing of barcodes and marker genes across 191 thalamic sections yielded 390 projection neurons after quality control (**Fig. 4e**). As we have done for the mouse BARseq data, we estimated the probability of assigning a target site barcode to the wrong soma to be under 0.01% by matching shuffled barcodes (**ED Fig. 9e**). We hierarchically clustered neurons into transcriptomically defined neuron types (see **ED Fig. 9f** for cluster mapping) and demarcated anatomical divisions using patterns of select genes and neuron types that were confined to specific nuclei, as described for the gene-only dataset (see **Supp. Note 3**). For some analyses, we grouped the cubelets into eight target domains using NMF, enabling examination of projections at both the original resolution and a lower resolution that better matched the mouse data for comparison (**ED Fig. 10a, b**; **Supp. Note 5**).

Individual neurons projected to diverse cortical targets in patterns that were stereotypical for their thalamic origin (**Fig. 4e-g**) (Córdoba-Claros et al., 2025b; Mundinano et al., 2019; Roberts et al., 2007; Romanski et al., 1997; Shipp, 2001). To assess agreement between BARseq single-neuron projections and existing anterograde bulk tracing data, we spatially matched BARseq somas and projections to published MPul anterograde tracing cases (Córdoba-Claros et al., 2025b) (see **Supp. Note 7**; **Fig. 4h**). Many single neurons whose somas fell within a given bulk tracing injection site projected to cortical locations overlapping with the corresponding bulk tracing-labeled projection zones (**Fig. 4h**, case 22R). In other cases, the projection patterns of individual neurons overlapped with only a subset of the bulk projection zones (**Fig. 4h**, case 13L). Across all spatially matched neurons, BARseq projections were significantly closer to the matched bulk tracer-labeled territories than the projections of neurons outside the bulk tracer injection sites (**Fig. 4i**; **Supp. Table 5**; *p =* 1×10^-5^, rank sum test). Most matched neurons (34/37, 92%) had projection patterns that overlapped with or abutted the matched bulk tracer-labeled territories, compared with 17/37 (46%, *p =* 3×10^-5^, Fisher’s exact test) randomly selected neurons with somas outside the matched injection sites. Thus, BARseq projection patterns recapitulated thalamic projection patterns seen in traditional anterograde tracing data.

### Increased selectivity and local homogeneity distinguish marmoset thalamocortical projections

While the primate thalamus is traditionally viewed as more discretely organized than the rodent thalamus, with sharply bounded nuclei and largely segregated projections to cortex (Baleydier and Morel, 1992; Jones, 2007), this has never been demonstrated at a single-cell level. Individual marmoset neurons often projected to a small region of cortex, appearing more selective than their mouse counterpart in single-neuron projections (**Fig. 5a**). To quantify this difference, we calculated the selectivity index, defined as the fraction of barcode molecules found in the strongest target domain per neuron; a high selectivity index thus indicates that a neuron projects to a more focal location in the cortex. Marmoset neurons had higher selectivity indices than mouse neurons (**Fig. 5b**; *p =* 5 ×10^-21^, Kolmogorov-Smirnov test; see **ED Fig. 10c** for robustness to the number of target domains). As a complementary measure, we calculated each neuron’s effective cortical coverage, defined as the exponential of the entropy of its barcode distribution, normalized by the number of cubelets per hemisphere (see **Methods**). Individual marmoset neurons covered 3.7% ± 2.1% (mean ± s.d.) of a cortical hemisphere, whereas mouse neurons covered a 4.4-fold larger fraction of the cortex (16% ± 10%, *p =* 2 × 10□^1^□□, rank sum test; **ED Fig. 10d**). The species difference persisted across a range of barcode count thresholds, indicating that it was not driven by differences in sampling depth (**ED Fig. 10e**). These results indicate that individual marmoset thalamic neurons project more selectively than mouse neurons, and are consistent with characterization of macaque cortical neurons (Gou et al., 2025) and amygdala neurons (Zeisler et al., 2024).

**Figure 5.**
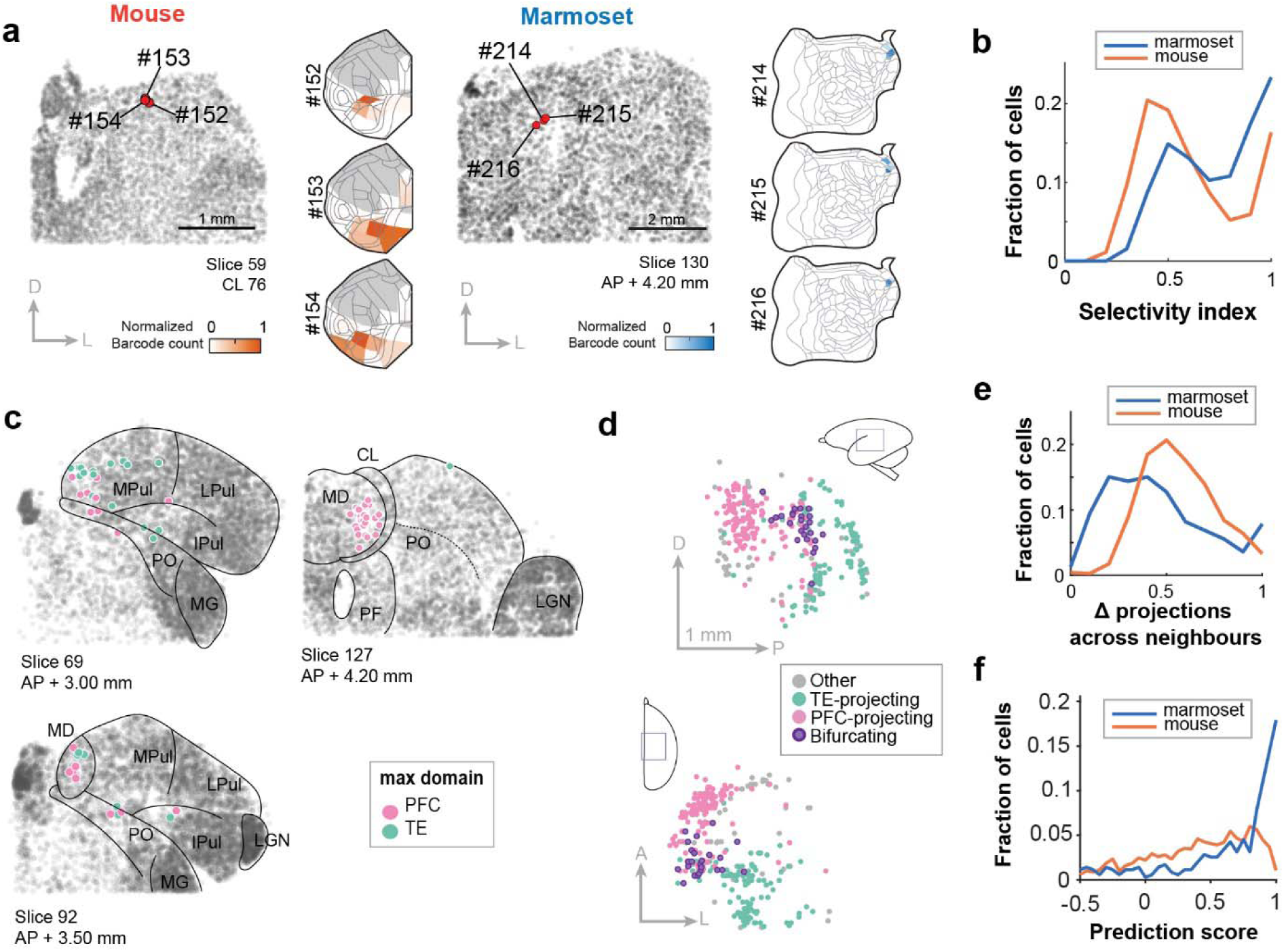
Marmoset thalamocortical projections are more spatially organized. (**a**) Projection patterns of representative neurons in mouse (*left group*) and marmoset (*right group*). Within each group, *Left*, soma locations of each representative neuron, with section numbers and corresponding AP coordinates indicated. *Right,* cortical projection patterns of each neuron visualized on a flatmap. Colors correspond to normalized barcode counts. (**b**) Distribution of the selectivity index for marmoset (*blue*) and mouse (*orange*) neurons. (**c**) Soma locations of neurons that project to the PFC or the TE in three example slices that span the transition between MPul and MD. Each section is labelled with its slice number and corresponding AP coordinate from the stereotaxic atlas (Paxinos, 2012). (**d**) Soma locations of neurons projecting to TE, PFC, or both, shown in two views. On the brain icons, the boxes indicate the location of the extent of the labelled region. (**e**) Distribution of the mean projection dissimilarity across spatial neighbors for marmoset (*blue*) and mouse (*orange*) neurons. (**f**) Distribution of prediction scores, reflecting the improvement of a local KNN model over a baseline in predicting thalamocortical projections, for mouse (*orange*) and marmoset (*blue*) neurons.

Beyond single neurons, nearby marmoset neurons also appeared to project more similarly, whereas neighboring mouse neurons diverged in their projections (**Fig. 5a**). This apparent local homogeneity raises the question of whether marmoset projections are more spatially segregated, as opposed to the strong intermingling observed in the mouse thalamus **(Fig 3h, j)**. We first examined neurons that projected to TE and/or PFC, the two areas that received most projections in this dataset. PFC-projecting neurons were predominantly found in MD but extended into the medial part of MPul (**Fig. 5c, *Slice 69***), whereas TE-projecting neurons were mostly in MPul, and extended into the posterior part (**Fig. 5c, *Slice 92***) but not the anterior part of MD (**Fig. 5c, *Slice 127***). Only a small number of neurons bifurcated to both TE and PFC (**Fig. 5d**), and the two populations remained largely segregated even within nuclei where both were present (**Fig. 5c**), suggesting that neurons with different projections were less intermingled than in mouse. To quantify the extent of intermingling, we computed the mean projection dissimilarity between each neuron and its 10 nearest spatial neighbors within a 500 µm radius. In both species, the dissimilarity distribution was broad, reflecting a substantial fraction of neurons whose neighbors projected differently (**Fig. 5e**). However, the distribution was shifted toward lower dissimilarity in marmoset, indicating that neighboring neurons projected more homogeneously than in mouse (*p =* 5 × 10^-22^, Kolmogorov-Smirnov test). This difference was robust to the number of neighbors used (**ED Fig. 10f**). Thus, thalamocortical projections are less spatially intermingled in marmoset than in mouse.

This greater local homogeneity, combined with the higher selectivity of single-neuron projections, should make a marmoset neuron’s spatial position more informative about its projection target. To test this hypothesis, we built a KNN classifier that predicted each neuron’s projection pattern from the mean projection of up to 10 nearest spatial neighbors within 500 µm. We then defined a prediction score as the reduction in the fraction of the prediction error in cosine distance over a shuffled baseline (see **Methods**); a higher score indicates that spatial location is more informative of projections. In both species, local prediction outperformed the random baseline (median 0.83 [IQR 0.47 - 0.96] in marmoset; 0.45 [IQR 0.06 - 0.73] in mouse), but the improvement was substantially larger in marmoset (**Fig. 5f**; *p =* 3 × 10^-34^, Kolmogorov-Smirnov test). These analyses show that the marmoset thalamocortical projections are more spatially ordered than those in mouse, suggesting that they form a circuit that is closer to a point-to-point organization.

### Gradient organization of thalamocortical connectivity is conserved across mouse and marmoset

The presence of the same projections across nucleus borders in both mouse and marmoset suggests that projections change across thalamic borders either as a cellular-level gradient or as a compositional gradient of two discrete projection types. To distinguish these models, we focused on two sets of borders between HOs nuclei, mouse PO/LP and marmoset MD/MPul, and one border between an HOs nucleus and a non-sensory nucleus, mouse PO/PF. For each, we scored every cell by the projection feature that best predicted its position along the axis joining the two nucleus centroids and examined how this score was distributed along that spatial axis (**Fig. 6a**). In all three pairs of nuclei, the scores varied along a diagonal from one nucleus to the other (albeit along a broad range for PO/PF). We then compared how well a single-Gaussian model (cellular gradient) or a two-Gaussian mixture with a spatially varying mixing weight (compositional gradient) could fit the data, and summarized the difference as a compositional index (see **Methods**). For all three pairs of nuclei, the compositional indices strongly favored a one-Gaussian model (**Fig. 6b**; p = 0.1 to 0.5 compared to the parametric bootstrapping null, raw values). In contrast, performing the same analysis on LP/midbrain and a synthetic compositional dataset derived from mouse PO/LP strongly favored the two-peak models (**Fig. 6b**; p < 0.025 for both after Bonferroni correction). These results indicate that projections vary as cellular-level gradients across these nuclei.

**Figure 6.**
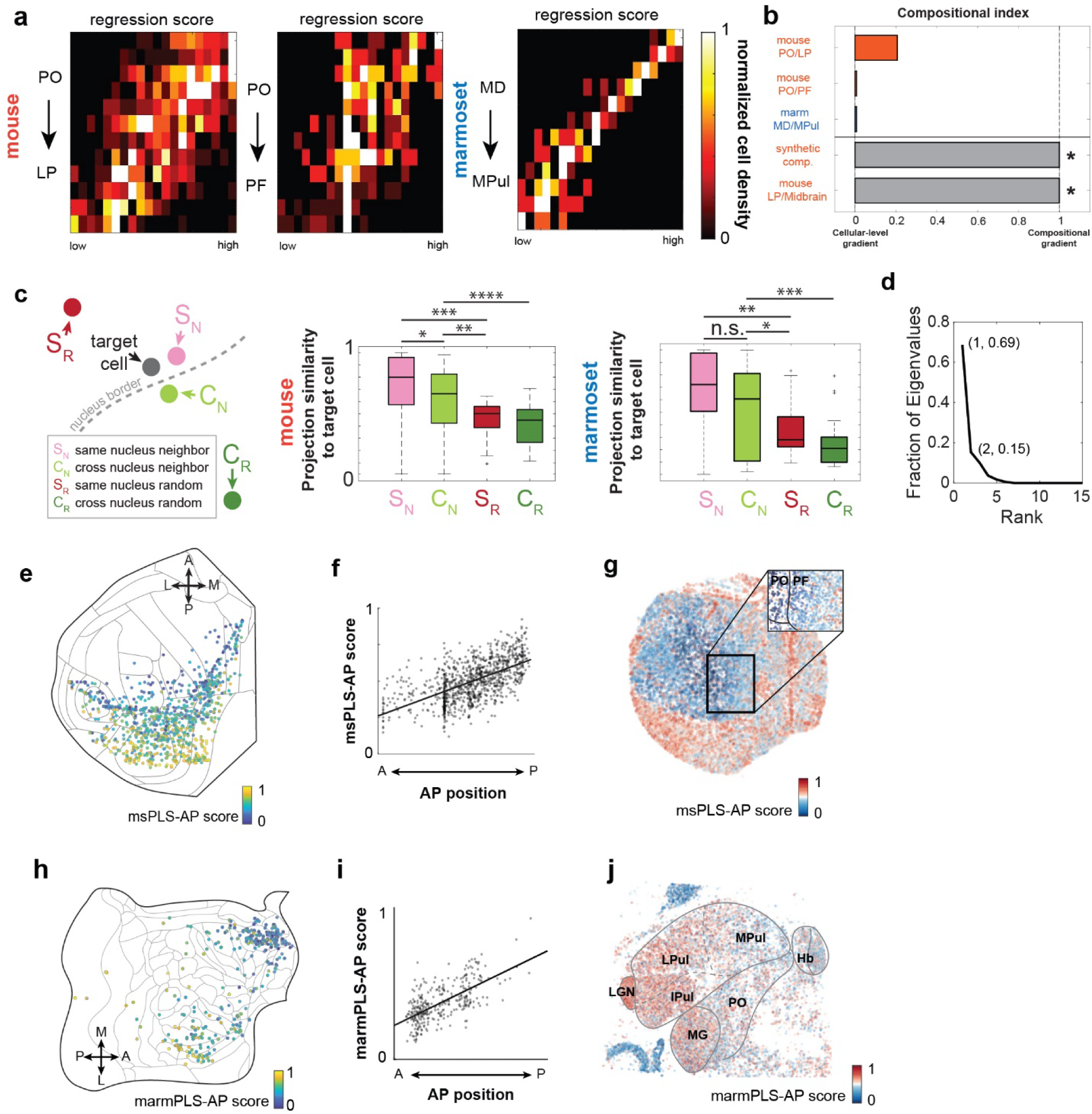
Gene expression predicts the anterior-posterior topography of thalamocortical projections in mouse and marmoset. (**a**) Heatmaps showing the distribution of projection scores on the regression axis that distinguishes the two nuclei. Y-axis indicates spatial locations between the two nuclei, and X-axis indicates projection scores. Colors indicate number of neurons, normalized by row max. (**b**) compositional index for each pair of nuclei. *p < 0.025 after Bonferroni correction. (**c**) *Left*, Schematic illustrating four categories of neuron pairs: neighboring neurons within the same nucleus (S_N_), cross-nucleus neighbors (C_N_), same-nucleus random neurons (S_R_), and cross-nucleus random neurons (C_R_). *Middle*, projection similarities between mouse neurons and the four categories of neurons shown on the left. Lines indicate medians, boxes indicate quartiles, and whiskers indicate ranges. * *p =* 0.002, ** *p =* 3×10^-11^, *** *p =* 9×10^-13^, **** *p =* 8×10^-15^, using Kolmogorov-Smirnov tests. *Right*, projection similarities between same-nucleus neighbors (S_N_), cross-nuclei neighbors (C_N_), same-nucleus random (S_R_), and cross-nucleus random (C_R_) neurons in the marmoset. Lines indicate median, boxes represent quartiles, and whiskers indicate range. * *p =* 0.001, ** *p =* 7×10^-7^, *** *p =* 9×10^-6^, all using Kolmogorov-Smirnov test. (**d**) Fraction of eigenvalues for each LDA axis. (**e**) Projection centroids of individual mouse neurons plotted on a cortical flatmap, colored by msPLS-AP score. (**f**) Relationship between the AP position of each mouse neuron’s projection centroid (x-axis) and its score along the gene axis (y-axis). Each dot represents a single neuron. Black line indicates a linear fit. (**g**) A representative mouse thalamic section with neurons colored by their msPLS-AP score. Inset contains a magnified view of the boxed area overlaid with borders, showing gradual change in msPLS-AP score across the PO/PF border. (**h**) Projection centroids of individual marmoset neurons plotted on a cortical flatmap, colored by marmPLS-AP score. (**i**) Relationship between the AP position of each marmoset neuron’s projection centroid (x-axis) and its score along the gene axis (y-axis). Each dot represents a single neuron. Black line indicates a linear fit. (**j**) A representative marmoset thalamic section with neurons colored by their marmPLS-AP score.

If projections are not confined by borders, how do thalamic borders and nucleus identity contribute to differences in projections? To test the effect of nucleus on projections, for each neuron, we identified its nearest spatial neighbors and compared projection similarity among same-nucleus neighbors, cross-border neighbors, and non-neighboring neurons from the same pair of nuclei (**Fig. 6c**). In both species, spatial proximity mattered: neighbors from the same nuclei were more similar in projections than non-neighbors from the same nuclei (**Fig. 6c**, S_N_ versus S_R_; p = 9×10^-13^ and 7×10^-7^, for mouse and marmoset, respectively, using Kolmogorov-Smirnov test). Nucleus identity, however, had a smaller effect: same-nucleus neighbors were slightly more similar than cross-border neighbors in mouse (**Fig. 6c**, S_N_ versus C_N_; *p =* 0.002), with a consistent but non-significant trend in marmoset. However, this border effect was outweighed by spatial proximity: cross-border neighbors remained more similar in projections than same-nucleus non-neighbors (**Fig. 6c**, C_N_ versus S_R_; *p =* 3×10^-11^ for mouse and *p =* 0.001 for marmoset, Kolmogorov-Smirnov test). Defining neighbors as all neurons within a set radius gave similar results across a wide range of radii (see **Supp. Table 3**), indicating that these results were robust to the choice of neighbor definition. Spatial proximity, therefore, dominates the organization of projections, whereas nucleus borders contribute only weakly, further confirming that this gradient organization across borders is conserved between mouse and marmoset.

We next examined how gene expression gradients relate to projections. To clearly distinguish between species throughout this section, we prefixed transcriptomic latent factors with “marm” and “ms” to denote marmoset and mouse, respectively. Since thalamocortical projections have traditionally been described as discrete, parallel channels (Jones, 2007; Sherman and Guillery, 2002), we began by treating projection identity as categorical: we grouped the mouse neurons into projection-defined populations based on their dominant projection, and used linear discriminant analysis (LDA) to find the gene axes that best separated them. The first two (msLDA1 and msLDA2) together accounted for 84% of the between-class variance (**Fig. 6d**), suggesting that only a small number of gene-expression dimensions were informative for projections. The low dimensionality raised the possibility that gene expression reflected the spatial arrangement of projection targets, which is similarly low dimensional. Consistent with this hypothesis, msLDA1 correlated with the anterior-posterior (AP) locations of cortical projection targets (**ED Fig. 11a**; Pearson *r =* 0.45, *p =* 2×10^-52^), whereas msLDA2 separated medial domains (PL/IL and ACA) from the dorsal and lateral domains (**ED Fig. 11b**; *p =* 3×10^-16^, rank sum test). To confirm that the correlation between msLDA1 and AP locations reflected the spatial arrangement of projections rather than chance, we shuffled the coordinates of each projection cubelet and recalculated the projection centroids. Shuffling abolished the correlation between msLDA1 and AP position (shuffled Pearson correlation *r =* 0.00 ± 0.24, *p =* 0.04 comparing real to shuffled data). This correlation between msLDA1 and the AP locations was notable because LDA was given only categorical projection labels, with no information about the spatial positions of projection targets. Thus, projection-defined populations were arranged along a gene-expression axis that is associated with the AP positions of their cortical targets.

To examine the correspondence between gene expression and AP locations of projections directly, rather than as a by-product of separating projection labels, we used partial least squares (PLS) regression to predict the flatmap coordinates of each neuron’s projection centroid from its gene expression. In the mouse dataset, we resampled neurons to equalize projection centroid density across the flatmap, and defined msPLS-AP as the linear combination of leading PLS components that best predicted AP position (msPLS-AP; **Fig. 6e, f**, Pearson *r =* 0.60, *p =* 4×10^-98^). msPLS-AP captured similar structure to msLDA1: both predicted AP target position (*r* = 0.60 and 0.45, respectively), had correlated gene loadings (**ED Fig. 11c**; Pearson *r =* 0.51, *p =* 6×10^-14^), and shared a similar dorsolateral-high, ventromedial-low expression pattern in the thalamus (**Fig. 6g**; **ED Fig. 11d, e**). This projection-associated axis is also distinct from the first principal component of thalamic gene expression described previously (Huang et al., 2026; J. W. Phillips et al., 2019), which showed a dorsomedial-high, ventrolateral-low pattern and was uncorrelated with the msPLS-AP loadings (**ED Fig. 11f, g**; Pearson *r =* 0.08, *p =* 0.3). Notably, the msPLS-AP gradient extended across the PO/PF border into PF (**Fig. 6g**), even though the same border separates distinct subclass-level neuron types (**ED Fig. 11e**). Because the mouse PO and PF project to different layers of the cortex (Harris et al., 2019), this gene axis is not a general predictor of projection “types,” but may be a specific predictor of projection AP positions.

To test whether this relationship between gene axis and projection topography is conserved across species, we then applied the same approach to identify the equivalent axis in the marmoset dataset (marmPLS-AP). marmPLS-AP was also associated with the AP positions of projection targets (*r* = 0.66, *p =* 2×10□□^2^) (**Fig. 6h, i**). Cells in the LPul scored high, and cells in the MPul, MD and PO scored low on this axis, mirroring the results from the mouse (**Fig. 6j**). An analogous axis in each species, however, does not establish that the two axes are equivalent. We therefore restricted to the 45 genes that were shared across the two gene panels and scored each cell on both the mouse and marmoset axes. The scores were correlated in both populations (**ED Fig. 11h**; Pearson *r =* 0.54, *p =* 8×10^-76^ for mouse neurons and Pearson *r =* 0.41, *p =* 4×10^-12^ for marmoset neurons), suggesting that the two axes are partly conserved across species. Shared genes contributing to these axes exhibited spatial expression patterns that crossed classical thalamic nuclei boundaries. Genes at the anterior-projecting end of the axis, including *Cdh13* and *Ncam2* (**ED Fig. 11i**), were enriched in the mouse PF and marmoset MPul, with weaker expression extending laterally in both species (**ED Fig. 11j**). Genes at the posterior-projecting end showed two types of patterns that were consistent across the two species: some genes, such as *Pvalb* (**ED Fig. 11j**), were expressed along a lateral-high, medial-low pattern, whereas other genes, such as *Calb2* (**ED Fig. 11j**) were expressed along a dorsal-high, ventral-low gradient in both species.

To summarize, the conserved projection gradients across borders (**Fig. 6a, b**), the projection-associated gene axis across the two species (**Fig. 6e, h**), and the shared transcriptomic gradients across nucleus borders (**Fig. 2**) together indicate that gradient-based organization is a conserved principle of the thalamus evident in both connectivity and gene expression.

## Discussion

By pairing spatial transcriptomic profiling with cortical projection mapping in the same neurons, BARseq provides a comprehensive view of thalamocortical organization across multiple thalamic nuclei in mouse and marmoset at single-cell resolution. Comparison of single-cell gene expression and projections across the two species revealed that marmoset thalamocortical projections were more spatially segregated than mouse neurons. Both species, however, share a conserved organization that reconciles the traditional models of discrete thalamic nuclei with the broad gradients across the thalamus revealed by recent transcriptomic studies. Projection patterns vary continuously across and within nuclei, and anatomical borders only had a small effect on projections. This organization is accompanied by a transcriptomic axis related to the anterior-posterior (AP) position of cortical targets, forming a coordinate-like landscape that bridges circuit architecture and molecular identity. This graded organization coexists with other gene expression variations that distinguish anatomical structures and/or transcriptomically defined cell types, revealing a multidimensional organization that was only partially captured by the discrete and gradient views. By integrating single-cell projection profiling with transcriptomics, we establish a gene-based reference that enables cross-species connectivity analyses at single-cell resolution, even in the presence of non-isometric expansion of brain regions.

### Projection topography is reflected in a transcriptomic coordinate axis

Our data identify a transcriptomic axis that is associated with the AP position of cortical projection targets in HOs thalamus. Earlier models of the primate pulvinar similarly proposed that cortico-pulvinar connections form a map of the cortical sheet that can cross traditional nuclei and subdivisions (Shipp, 2003). Our data broaden the idea beyond the pulvinar to other HOs nuclei. This result extends prior population-level reports linking thalamic gene-expression gradients to thalamocortical connectivity (Huang et al., 2026; J. W. Phillips et al., 2019; Puche-Aroca et al., 2025) to the single-neuron level. This gene axis could be a remnant of developmental patterning along the AP molecular gradients (Dufour et al., 2003; Fukuchi-Shimogori and Grove, 2003; Kudo et al., 2007). However, this adult gene axis is unlikely to directly reflect developmental instructions, because multiple mechanisms refine the thalamocortical projection patterns after the initial targeting (Grant et al., 2012; Sydnor et al., 2024). Alternatively, this gene axis could reflect physiological specialization tuned to the properties of cortical targets. For example, cortical areas differ in intrinsic time scales, with primary sensory regions generally operating on faster timescales, including in marmoset cortex (Canton-Josh et al., 2025; Li et al., 2025; Murray et al., 2014). Thalamic neurons projecting to different positions along the cortical AP axis may therefore require different intrinsic membrane or synaptic properties. Whether developmental in origin, functionally tuned, or both, the conservation of a gene-projection axis in both mouse and marmoset suggests that projection topography-aligned transcriptomic organization is a fundamental principle of thalamocortical wiring.

A continuous molecular coordinate system could allow projection topography to be preserved as anatomy changes. Expanded or newly differentiated cortical territories can occupy different positions along an existing thalamic coordinate axis, rather than requiring new cell types. This idea is consistent with models proposing that an AP topographical organization is maintained across the cortex, striatum, pallidum, and thalamus during forebrain evolution (Bailey and Von Bonin, 1957; Giarrocco and Averbeck, 2023), and suggests a possible model by which thalamocortical circuits can evolve alongside the cortex without sacrificing coherent organization.

### Single-cell gene expression and projection variations reconcile the discrete and gradient views of the thalamus

Our data reveal that neuronal populations with different projections do not segregate cleanly in spatial organization. This intermingled organization is also consistent with functional studies. For example, in primates, thalamocortical projections from several territories overlap across sensory and motor systems, providing anatomical routes for multisensory and sensorimotor interactions (Cappe et al., 2009). In the macaque, the medial pulvinar possesses visual, auditory, and multisensory neurons and shows cross-modality integration (Vittek et al., 2023). In mice, activity in HOs thalamus can be reshaped by behavioral training so that neurons respond to behaviorally relevant, reward-predicting stimuli regardless of stimulus modality (Petty and Bruno, 2024). Thus, our anatomical data, together with these functional studies, suggest that the organization of the thalamus could allow information to be routed flexibly according to the behavioral context rather than sensory modality alone.

The organization of the thalamus has long been described as a collection of nuclei with further subdivisions and laminations, defined by cytoarchitecture, chemoarchitecture, connectivity, and function. Recent molecular and imaging studies have instead emphasized gradients of thalamic gene expression and connectivity across classical anatomical boundaries (Huang et al., 2026; Müller et al., 2020; Oldham and Ball, 2023; J. W. Phillips et al., 2019; Puche-Aroca et al., 2025). Although the discrete and gradient views appear to conflict with each other, our data revealed that both organizations exist in gene expression and projections in the mouse and marmoset thalamus. Cortical projection patterns varied continuously across the thalamus but changed more rapidly near borders, indicating that gradient and discrete features coexist in projections. This coexistence holds even though our mapping covered only a subset of nuclei and captured only area-level differences, not other features such as laminar patterns of axonal termination (Harris et al., 2019) that could reveal further gradient and/or discrete distinctions across nuclei. Transcriptomic identity was similarly two-sided: whereas transcriptomic clusters separated broad thalamic divisions, such as the intralaminar thalamus, the projection-associated gene axis varied continuously across anatomical borders. This observation explains why discrete and gradient views have, to date, been both compelling: classical neuroanatomy seeks to describe the thalamus in discrete terms, and captures divergence in projections and/or discontinuities in gene expression as borders, whereas molecular and imaging studies seek to describe organization across the whole brain, and are better suited to capturing variations as gradients. The discrete and gradient views therefore reveal different aspects of thalamic organization that are conserved across mouse and marmoset.

### Sharper spatial organization may be a functional adaptation to brain expansion

Individual marmoset thalamic neurons projected to more selective cortical territories than those of mice, and neighboring marmoset neurons had more similar projection patterns. This species difference resembles those recently reported for the macaque prefrontal corticocortical connections, where individual neurons showed greater target specificity, fewer collaterals, and smaller brain-size-normalized arbors than mouse neurons (Gou et al., 2025). Such findings are consistent with the observation that the expansion of cortical surface area outpaces white matter volume in larger brains and that larger brains possess relatively sparser area-to-area connections (Ardesch et al., 2022). Nevertheless, white matter scaling alone is unlikely to explain our results and other reports of sparsification in primate subcortical projections (Zeisler et al., 2024). The dominant scaling theories argue that corticocortical connections are constrained by computational timing imposed by a larger brain (Ringo, 1991; Ringo et al., 1994), wiring economy (Bullmore and Sporns, 2012; Chklovskii et al., 2002; Zhang and Sejnowski, 2000), and/or geometric constraints imposed by gyrification, in which mechanical folding of the expanding cortex reshapes the trajectories available to corticocortical fibers (Mota et al., 2019; Mota and Herculano-Houzel, 2015; Tallinen et al., 2014; Van Essen, 1997). However, these factors likely act on thalamocortical projections differently: thalamocortical projections are shorter than callosal connections, occupy a small fraction of the white matter, and are radial. In fact, it may be functionally important to supplement corticocortical connections with cortico-thalamo-cortical loops to strengthen communication between key cortical areas (Jaramillo et al., 2019; Shepherd and Yamawaki, 2021; Sherman, 2016).

Instead, we propose that increased thalamocortical selectivity reflects a functional advantage. As the cortex expands and differentiates, cortical areas become more specialized in architecture, connectivity, and function (Chaplin et al., 2013; Magrou et al., 2024). Because this cortical differentiation creates distinct targets that did not exist in a smaller, less parcellated cortex, thalamocortical inputs that terminate in more confined locations can address these specialized areas separately, increasing the precision of thalamic control over cortical computation. Spatial segregation of thalamocortical projections may therefore follow cortical specialization rather than arise independently: as the cortex differentiates, more selective thalamic input becomes both possible and useful. Such local specialization, superimposed on a conserved gradient, may allow thalamocortical systems to preserve a common global plan while tuning local circuitry to the demands of a larger, more parcellated cortex.

## Supplementary notes

### Supplementary Note 1: Gene panel selection and validation

Because BARseq-style in situ sequencing, like other imaging-based spatial transcriptomic approaches, can interrogate a targeted panel of genes, we first selected genes to best capture gene expression variation in the marmoset thalamus based on two single-nucleus RNA sequencing (snRNA-seq) datasets (Dan et al., 2025; Krienen et al., 2023). We aimed to capture variation across the whole thalamus, with particular focus on resolving the transcriptomic heterogeneity of two nuclei: the pulvinar (Pul), a major higher-order sensory thalamic nucleus, and the mediodorsal nucleus (MD), a hub connecting sensory streams to the PFC. Because previous reports have shown that thalamic neurons contain both distinct transcriptomic clusters (Dan et al., 2025; Krienen et al., 2023; Z. Yao et al., 2023) and transcriptomic gradients (Huang et al., 2026; J. M. Phillips et al., 2019), we applied three complementary approaches **(ED Fig. 1a)** to capture both discrete and continuous variation.

We first used MetaMarkers (Fischer and Gillis, 2021) to identify cell type marker genes that are highly expressed and differentially expression either across coarse-level excitatory neuron types in the whole thalamus or across fine-grained cell types in MD and Pul. We subset the snRNA-seq dataset to the thalamic excitatory neurons by identifying clusters that highly express genes characteristic of thalamic excitatory neurons, such as *SLC17A6* (thalamic excitatory neuron marker), *PTPN3*, *KITLG*, *NTNG1*, *OTX2*, *MYO1B*, and *LEF1* (Kita et al., 2021; Shimogori et al., 2018). We identified clusters that are specifically enriched in the MD and Pul using a published Xenium spatial transcriptomic dataset that had been mapped to the aforementioned snRNA-seq dataset. This dataset used a gene panel that was optimized for the basal ganglia, not for the thalamus, but nonetheless distinguished cells across broad thalamic divisions. Because thalamic gene expression also vary continuously (Huang et al., 2026; J. W. Phillips et al., 2019), we next used PERSIST (Covert et al., 2023) to identify genes that best capture gene expression variation across the thalamus, irrespective of cell-type labels. We also included additional classic anatomical markers and select genes of interest in the thalamus, including CALB1, PVALB, *CACNA1C*, *DRD1*, *DRD2*, *DRD3*, *GRIN3A*, *NEGR1*, *OPMCL*, and *PTPRD*. These three sets of genes were then combined with a gene panel focused on distinguishing marmoset cortical neurons, which were designed using MetaMarkers (Fischer and Gillis, 2021).

To estimate how well the gene panel captured cellular diversity in the thalamus, we built a k-nearest-neighbor-based classifier to predict the cluster label of a neuron in the snRNA-seq dataset using various gene panels, and compared them to the performance that can be achieved by the top 1,000 highly variable genes (see **Methods**). Both specificity and sensitivity improved with more genes in the panel and approached the level achievable using highly variable genes, but plateaued between 143 and 179 genes (**ED Fig. 1b**)

The mouse gene panel is a combination of a 104-gene cortical panel (Chen et al., 2025), combined with a 84-gene thalamic panel, both of which were previously designed and validated.

### Supplementary Note 2: Hierarchical clustering of in situ sequencing data

We clustered the neurons at three levels, which we denote as hierarchy 1, 2, and 3 (H1, H2, H3; **ED Fig. 2a**). At a coarse level (H1), clustering identified cell populations that were specific to brain regions, such as the thalamus, the thalamic reticular nucleus, the striatum, the hippocampus, and cortical layers (**ED Fig. 2a-c**). We then combined two H1 types that contained excitatory neurons in the thalamus, midbrain, and hypothalamus, and reclustered them into H2 types.

Clustering at the H2 type level initially resulted in seven clusters (**ED Fig. 2d**), but two clusters were of low-quality cells and were removed from downstream analysis. Neurons in the cluster TH_broad were found in the same locations as TH1 and TH2 type neurons with no obvious enrichment at specific locations (**ED Fig. 2e**), had lower read counts per cell compared to TH1 and TH2 (**ED Fig. 2f**), and highly expressed MBP (**ED Fig. 2g**), an oligodendrocyte marker. These characteristics suggest that TH_broad included segmentation errors in which portions of neurons and oligodendrocytes were segmented as single cells, which was confirmed by manual examination of a subset of TH_broad cells. We thus excluded this cluster from further analysis. Lowqual_HY contained a small group of cells at the ventral edge of the hypothalamus and no cells in the thalamus, and was also excluded from further analyses (**ED Fig. 2h**). Two of the remaining H2 types (TH1, TH2) were confined to the thalamus and enriched in higher-order and first-order nuclei, respectively, whilst the remaining three (ILM, HY_Hb, ILM_VA) included neurons in specific nuclei either close to the midline or in the anterior portion of the thalamus, along with neurons in the midbrain, and the hypothalamus.

We reclustered each H2 type again, resulting in 24 H3 thalamic excitatory types (**Fig. 1f**). At this level, neurons in the three mixed H2 types separated into eight H3 types that included neurons in the intralaminar nuclei (centrolateral, CL and centromedial, CM), the habenula (Hb), anterior dorsal nucleus (AD), parafasicular nuclei (PF), and portions of the medial side of the ventroanterior nucleus (VAm), and 21 H3 types that were mostly in the hypothalamus and the midbrain; these non-thalamic H3 types were excluded from further analyses.

### Supplementary Note 3: Transcriptomic feature-based area demarcation

Generally, we used the mRNA expression patterns of classic anatomical markers, such as *CALB1* and *PVALB*, combined with neuron type distribution, to draw anatomical borders. Here we present detailed examples on the area demarcation in the marmoset in situ sequencing dataset (**Fig. 1** and **Fig. 2**). We first illustrate the rationale for demarcation on three representative slices in detail (**ED Fig. 3a-c**), then show the demarcation on all slices (**ED Fig. 3d**). The same approach was also applied to demarcate the marmoset BARseq dataset (**Fig. 4**)

We first used major landmarks, such as the emergence and disappearance of major thalamic nuclei, to match slices to coronal planes. We matched BARseq slice 19 to AP coordinates +2.3 in the marmoset atlas (Paxinos et al., 2026) (**ED Fig. 3a**), based on the shape of the pulvinar, which was *CALB1* positive, the emergence of the MG, and its size relative to the pulvinar. The borders of the medial division of the inferior pulvinar (IPulm) was denoted by a lack of *CALB1* expression within the pulvinar (Paxinos et al., 2026). CA1 of the hippocampus showed a thick band of *CALB1* signals in its pyramidal cell layer (Py), and the dentate gyrus was characterized by a thin ribbon of *CALB1* signal in the granular cell layer (GrDG).

We matched BARseq slices 43 to AP coordinates +3.5 (**ED Fig. 3b**). *PVALB* revealed the LGN. In addition, a thin ribbon of *PVALB*-negative cells marked the LGN’s koniocellular layer 3 (K3). K3 is the thickest koniocellular layer at this coronal section and separates the LGN magnocellular and parvocellular layers. At this coronal section, the brachium, a white matter bundle, passed through the pulvinar and connected to the dorsal LGN. Both the posterior limitans (PLi) and the thalamic reticular nucleus (TRN) were positive for *PVALB*, marking the medial-ventral and dorsal-lateral borders of the thalamic nuclei at this AP location. The medial portion of this slice was demarcated with the distinct localization of H3 types. For example, HY_Hb_1, HY_Hb_2, HY_Hb_3 were found in the medial habenula (mHb), and clusters ILM_1 and ILM_2 demarcated the centrolateral nucleus (CL) and intralaminar nuclei, which encased the MD and indicated the posterior end of the MD nucleus.

BARseq slice 91 was matched to an AP location of +5.5 (**ED Fig. 3c**). We used *CALB1* to identify the location of the lateral dorsal nucleus (LD) and the substantia nigra (SNr), both of which were *CALB1* positive. The remainder of the thalamus showed a relatively homogenous expression of *CALB1*, so we used neuron types to demarcate notable anatomical nuclei. Again, ILM_1 and ILM_2 revealed the location of the CL, but also the centromedial (CMn) and posterior parvoventricular nucleus (PVP) (**ED Fig. 3c-i**). The CMn could be demarcated by its own cell type (**ED Fig. 3c-ii**). The MD was encased by the CL/CMn nuclei, and the 3 subdivisions (MDm, MDc, MDl) could be differentiated by differential enrichment of TH1 subtypes (**ED Fig. 3c-iii**). Because the remaining nuclei and subdivisions were less distinct, we did not demarcate their borders.

This same approach was used to demarcate all hemi-coronal sections from the marmoset in-situ sequencing dataset (**ED Fig. 3d**) and the marmoset BARseq dataset. The mouse ABC atlas was pre-registered to the Allen Common Coordinate Framework v3 (CCFv3)(Wang et al., 2020), which allowed us to directly use the CCF labels.

### Supplementary Note 4: BARseq in mouse thalamus

Although we performed most of the gene expression analysis using cluster-agnostic approaches, we initially clustered the neurons hierarchically to examine the quality of the data. We mapped the clusters to cell types in reference snRNA-seq datasets (Z. Yao et al., 2023) using a kNN-based approach, and found that each mouse BARseq neuron type corresponded to one or a small number of reference clusters (**ED Fig. 5a**), indicating that this dataset recapitulated the diversity in gene expression in the reference snRNA-seq dataset. These clusters were then used to facilitate delineation of thalamic nuclei (**ED Fig. 5b**).

Because the projection sites were dissected from 300 µm thick sections, whereas in situ sequencing requires 20 µm sections, we developed an approach to allow cryosectioning the thalamus and the cortex on the same coronal planes at different thickness (**ED Fig. 6a**). We first cut 200 µm cryo-sections from both anterior and posterior ends of the brain until we saw the thalamus (Coronal level 67 for the anterior end of the thalamus, 78 for the posterior end). We punched out the thalamus with a 3 mm biopsy punch while it is still frozen, and continued to section the remainder of the brain to 200 µm sections. We sectioned the punched-out thalamus to 20 µm for in situ sequencing. We dissected all areas that receive projections from areas around the LP, based on the Allen Connectivity Atlas, into 17 cubelets (Harris et al., 2019). These include the Superior colliculus/periaqueductal gray (SC/PAG), 15 cortical regions, and the main olfactory bulb (MOB) as a negative control. All images of slices with the dissection areas marked out are provided in **Supplementary File 1**. The block-face image of each section was first matched to coronal planes in the Allen Reference Atlas based on major landmarks. We drew the dissection areas onto each matched CCF coronal plane, then transformed them into cortical flatmaps (**ED Fig. 6a**)

We first applied a linguistic complexity threshold to filter out soma barcodes with repetitive sequences (**ED Fig. 6b**). These “barcodes” were usually background fluorescence instead of true barcodes. To match the filtered soma barcodes to those in the projection sites, we first examined the distribution of minimum hamming distance between a soma barcode and projection barcodes, and compared the distribution to randomly shuffled soma barcodes (**ED Fig. 6c**). We found that a sizeable fraction of soma barcodes were perfect matches to projection barcodes (i.e., had minimal hamming distance of 0). In contrast, few shuffled barcodes were perfect matches (0.18% ± 0.04%, mean ± standard deviation across 200 shuffled barcode sets), indicating that the soma barcodes did not match by chance.

To exclude the possibility that barcodes in passing axons, dendrites, and/or nearby somas were assigned to a soma by mistake, we manually examined the sequencing images for each barcoded cell to identify 1,578 barcodes in 1,809 high confidence barcoded somas. Of these somas, 60 barcodes were found in multiple cells, 13 barcodes were found in the same cells but were imaged multiple times, because they were in the overlapping areas between adjacent imaging tiles. We filtered out barcodes that were found in multiple cells, and removed the “extra” cells that were imaged multiple times, leaving 1,518 cells with unique barcodes for downstream analysis. The observed barcode collision rate (i.e., the same barcode labelling multiple cells) was 60 / 1578 = 4%, which was consistent with previous studies (Chen et al., 2019; Sun et al., 2021). Because all of the detected double labelled barcodes have been removed from the filtered data, we consider the observed rate as an upper bound in the filtered data, and the effect on the downstream analyses was minimal due to the filtering. No barcode was found in the negative control site (MOB), or found in the brain other than the one that the soma was found in, indicating that the barcodes detected in the projection sites were specific.

### Supplementary Note 5: Non-negative matrix factorization of projection data in both species

Individual mouse neurons projected to 4.1 ± 2.5 target areas (mean ± standard deviation) (**ED Fig. 7f**), and the dominant target accounted for 61% ± 23% (mean ± standard deviation) of barcode molecules per neuron (**ED Fig. 7g**). Furthermore, certain areas were often co-innervated by the same neurons (**ED Fig. 7h**), suggesting that projection data may lie in a lower-dimensional space than the number of cubelets. This is typical of biological measurements in the nervous system, such as gene expression and neuronal activity, where the measured variables are far more numerous than the latent biological processes that generate them. Dimensionality reduction is a natural step for analyzing such data, as it can recover this underlying structure while suppressing noise. Recovering a shared low-dimensional structure also enables more accurate cross-species comparison, since the dissection resolutions of mouse and marmoset differed substantially (17 vs. 94 cubelets) and were not directly comparable in the original cubelet space. We thus applied NMF to group cubelets that were frequently co-innervated into target domains. In both mouse and marmoset datasets, we selected the number of domains based on how well they reflect prior biological knowledge. In mouse, we chose a value that resolved the major functional modalities of the mouse cortex (**ED Fig. 7i**). These included motor cortex (MOs), the auditory and temporal association areas (TE), the frontal cortex (PL/IL), and anterior cingulate cortex (ACA). Given the spatial extent of visual-associated cortex, visual areas were subdivided into three target domains: VIS-caudal (VIS-c), which encompasses retrosplenial cortex and constitutes the most posterior cortical domain; VIS-rostral (VIS-r), which contained primary visual cortex; and a third domain encompassing VIS-anteromedial and adjacent SSp cubelets (VIS-am/SSp). The superior colliculus and periaqueductal grey, dissected as a unit, formed an additional target domain.

Because the marmoset cortex was dissected at sub-area resolution, we selected the lowest number of target domains that resolved the full complement of cortical targets innervated by barcoded neurons across the pulvinar subdivisions and neighboring nuclei (**ED Fig. 10b**). Because we could not match the left and right hemisphere cubelet-to-cubelet, we performed NMF separately for the two hemispheres, and combined the data afterwards. In the right hemisphere, we identified three target domains in the temporal cortex, four in the prefrontal cortex, and one in the motor/premotor area. Of the three domains in the temporal cortex, two of them corresponded to the dorsal visual stream (TE-dorsal, including MT, FST, MST) and the ventral visual stream (TE-vent, including V4, TEO, TE), and a third domain that was more anterior to them (TE-ant, including STR and anterior TE areas). The dlPFC was further subdivided into two domains: a posterior domain (dlPFCp) encompassing the frontal eye fields (8Ad and 8Av), and an anterior domain (dlPFCa) encompassing areas 46d and 46v. Two medial prefrontal domains were also identified: a dmPFCa domain containing area 9, and a broader mPFC domain encompassing anterior cingulate (area 32) and medial orbital prefrontal cortex (areas 13/14). Motor and premotor cortex, consistent with their known connectivity with the posterior nucleus (PO), formed a separate domain. In the left hemisphere, we obtained eight domains: dlPFCa, TE-vent, TE-dorsal, TE-ant1, TE-ant2, IP-motor, V1, and vlPFC. The first three overlapped with those in the right hemisphere; TE-ant1 and TE-ant2 corresponded to TE-ant in the right hemisphere; IP-motor corresponded to Motor in the right hemisphere, but included additional intraparietal areas. The differences in the domains between the two hemispheres likely reflected differences in labelling, which captured neurons that projected to different cortical areas. For analysis purposes, we typically pooled the two hemispheres together by combining corresponding target domains. For analysis involving gene expression, we used the right hemisphere only, because the majority of the left hemisphere were sequenced with a smaller gene panel that did not fully resolve the gene-projection relationship.

### Supplementary Note 6: BARseq reveals MD – PFC projection topography in the marmoset

To validate BARseq in the marmoset, we mapped the MD projections to the PFC because of its well-documented topography in primates. We used an MRI-guided surgery robot to inject a barcoded pseudotyped Sindbis bilaterally into the MD nucleus of a pilot animal (**ED Fig. 8a**, **Methods**). We injected a variant of the Sindbis virus that was pseudotyped with the structural protein derived from the Eastern Equine Encephalitis virus (EEEV). This pseudotyped Sindbis virus has been previously optimized for barcoded projection mapping in primate brains (J Kebschull, unpublished data).

After tissue collection 48 hours post-injection, we used a vibrating microtome to section the frontal cortex of the animals into 400 µm slabs. These were frozen on microscope slides, and images were matched to the Stereotaxic atlas of the marmoset brain to demarcate dissection locations.

We sequenced the injection site for both barcodes and a small gene panel (**Supp. Table 2**). This panel was not sufficient to resolve finer differences in thalamic neurons, but was useful to parcellate thalamic nuclei. To consolidate the frontal cortical dissections with published literature, we aggregated cubelets into six regions of the PFC; Area 10 (most anterior portion of the frontal cortex), dlPFC (Area 46D, 46V, 8aV, 8aD), dmPFC (Area 9), mPFC (Area 32), oPFC (Area 13, 14, 11), and vlPFC (Area 45, 47). Projections to the frontal cortex can cover multiple contiguous regions, but usually one region had the most dominant projection (**ED Fig. 8d**). We thus categorized neurons based on their strongest projections.

Our data revealed projection differences across all three spatial axes across the MD. In the right hemisphere, where the injection was slightly more posterior, most neurons projected to the dlPFC. This dominance of dlPFC projections from the posterior end of the MD was consistent with previous imaging studies in the macaque (Saalmann et al., 2012). In the left hemisphere, where the labelled neurons were more anterior, neurons projecting to vlPFC, dlPFC, mPFC, and oPFC occupied ventral, lateral, dorsal, and medial quadrants, respectively (**Fig. 4c**). This circular topography was consistent with that described in a previous marmoset retrograde tracing study (Roberts et al., 2007). Thus, BARseq data accurately recapitulated the previously described topography of marmoset MD-PFC connectivity.

### Supplementary Note 7: Marmoset BARseq validation

To compare the marmoset projection data to existing bulk-tracing data quantitatively, we identified neurons in our data with somas that overlapped with injection sites in bulk-tracing data, then assessed overlap in the projections. To match somas in our data to bulk-tracing injections, we identified the AP locations of the centers of published injection sites, found the sections that corresponded to the same AP locations in our data, then re-traced the injection site boundary on our sections. We then identified BARseq neurons within that boundary, extended 100 µm in both the anterior and posterior direction. Because each injection site spanned about 500 µm – 800 µm within the coronal section, we expected that the real injection site would span a similar amount on the AP axis. Thus, our estimate of the injection site was conservative.

Because the published flatmap representation of the bulk tracing (Córdoba-Claros et al., 2025b) differed from the MRI-based surface projection used in our data, we needed to transform the two datasets into a common space for comparison. We manually retraced all bulk projection zones in the published dataset onto our flatmap template based on annotations in the Paxinos atlas. For each BARseq neuron, we then identified how many “targets” it projected to; each projection target was defined as a continuous zone of barcodes in dissection cubelets with a single local maximum. For each projection target in each neuron, we counted how many cubelets away it was to the closest area labelled by the matching bulk tracing. For neurons that projected to multiple targets, we used the mean of all targets (**Supp. Table 5**; **Supp. File 2**).

## Methods

### Animals and surgery

#### Marmosets

All experimental procedures were fully compliant with the Guidelines for the Care and Use of Laboratory Animals by the National Institutes of Health and approved by the Animal Care and Use Committee of the National Institute of Mental Health. Animals were housed at ambient conditions (26 - 28 °C, 40 – 60 % humidity) on a circadian cycle of 12-h light and 12-h dark, ad-libitum access to food and water. Experiments were performed on adult marmosets 1.5-6 years of age.

For all marmoset surgeries, Computed Tomography (CT) scans and Magnetic Resonance Imaging (MRI) scans were acquired up to 3 months before the procedure. DICOM volumes were converted to MINC (.mnc) format for co-registration: corresponding landmark tags placed in both volumes were used to compute a transformation matrix, and a linear transformation was applied to the CT to bring it into register with the MRI. Both volumes were imported into BrainSight (Rogue Research, Montreal, Canada) to plan injection targets and trajectories.

Anesthesia was induced with alfaxalone (8 mg kg□1, i.m.) and diazepam (1-3mg/kg i.m.) maintained with inspired isoflurane (0.5–2%) through a custom 3D printed face mask. Bupivicane (0.1 – 0.3ml, Aspen) was used as local anesthetic, injected subcutaneously. Eyes were protected with topical eye ointment for the duration of all procedures. The animal was kept on a homeothermic heat pad set at 38°C. The head was shaved and swabbed with a topical betadine (7.5 povidone-iodine) antiseptic cleanser, followed by 70% ethanol. Adults received a broad-spectrum antibiotic (Norocilin; Norbrook, Lenexa, KS, USA; 27 mg kg□^1^, 0.1 mL, i.m.), atropine (20um/kg, i.m.) and dexamethasone (0.3 mg kg□^1^, i.m.) to limit cerebral oedema, control heart rate, and were then secured in a stereotaxic frame (BrainSight). Physiological status (SpO□ and core temperature) was monitored continuously and held within defined limits.

A skin incision was made along the midline of the cranium and the skull exposed. The cranium was then registered to the co-registered CT/MRI by surface-based registration, as briefly described in (Scott et al., 2025). In brief, a robot-mounted laser (BrainSight Vet Robot, Rogue Research) projected a grid of points across the exposed bone; two calibrated cameras captured each point stereoscopically, allowing the skull surface to be reconstructed by laser triangulation. The resulting point cloud was aligned to the skull surface extracted from the CT, establishing the spatial mapping between the surgical field and the navigation coordinate system.

A craniotomy was then made over injection targets using a burr drill fitted with a 1mm ophthalmic bit. Virus was injected through a 5 μL Hamilton Neuros syringe fitted with a custom 15° bevel needle, driven by a Stoelting microinjector mounted on the BrainSight robotic arm. The needle was pierced through the dura directly to minimize cortical damage. All virus was injected at a rate of 180nl/min, and settled for 10 minutes before withdrawing the needle at 0.1mm/second. The cranial window was sealed with Kwik-Sil and the scalp incision closed with intradermal sutures (coated Vicryl 5-0). The animal was recovered in a temperature and humidity-controlled cage.

#### Mice

Animal handling was conducted according to protocols approved by the Institutional Animal Care and Use Committee (IACUC) of the Allen Institute. Animals were housed 3–5 per cage and were on a 12/12 light/dark cycle in an environmentally controlled room (humidity 40%, temperature 21 °C).

Barcoded Sindbis virus libraries were stereotaxically injected into target brain areas of C57BL/6J male mice using coordinates obtained from the Allen Reference Atlas (site 1: AP −1.96 mm, ML 1.2 mm, DV 2.7mm from Bregma; site 2: AP −2.26 mm, ML 1.5 mm, DV, 2.6mm from Bregma, 140 nL per site).

### Tissue processing for BARseq

Following Sindbis virus injection, marmosets recovered for 24 h in a temperature- and humidity-controlled ICU. To collect tissue, we euthanized animals with an overdose of pentobarbital (>100 mg kg□^1^, i.p.) and transcardially perfused them with ice-cold artificial cerebrospinal fluid (aCSF). We removed the brain and blocked out the thalamus with a razor blade. All blocks were flash-frozen in dry-ice cooled isopentane stored at −80 °C. The fresh-frozen injection site (whole thalamus) was cryosectioned at 20 μm with the chamber held at −14 °C. Ten to twelve sections were mounted per aminosilane-coated slide (Schott) and stored at −80 °C.

The remaining cortex was cryosectioned coronally at 300 μm on the same cryostat (chamber at −10 °C) to generate thick slabs for RNA extraction. Each slab was thaw-mounted onto a clean microscope slide and stored at −80 °C until microdissection. We acquired block-face images of all slabs and registered them to the *Marmoset Brain in Stereotaxic Coordinates* atlas (Paxinos, 2012) to delineate cortical areas. Microdissection was performed on a temperature-controlled metal block at −20 °C as previously described (Kebschull et al., 2016); we excised cortical regions by hand with a pre-chilled scalpel, keeping the section frozen throughout. Dissected tissue was collected into TRIzol, stored at −80 °C and shipped to the MAPseq/BARseq core facility (Cold Spring Harbour Laboratory) for sequencing and processing. For pilot animal (Alto), the cortex was instead sectioned at 300 μm on a vibratome in ice-cold aCSF. Slab images were registered to the atlas as above, and the tissue frozen at −80 °C and microdissected identically. Dissected tissue was collected into TRIzol, stored at −80 °C and shipped to the MAPseq/BARseq core facility (Cold Spring Harbour Laboratory) for sequencing and processing.

For non-surgical animals used for transcriptomic profiling, euthanasia and perfusion were performed as above. The intact hemisphere was frozen in isopentane at −60 °C and stored at −80 °C. Whole coronal sections were cut at 20 μm on the same cryostat and mounted two per aminosilane-coated slide (Schott).

For mice, we euthanized the animals 24 hours after surgery. We then removed the brain, flash-froze it in isopentane/ethanol bath and stored it at −80 °C. The cortex was cryosectioned coronally at 300 μm, and the thalamus isolated with a biopsy punch and sectioned at 20 μm for BARseq. Block-face images of the cortical slabs were registered to the Allen Mouse Brain Common Coordinate Framework (Wang et al., 2020). We then microdissected target regions, stored dissected tissue in TRIzol at −80 °C, then shipped them to the Cold Spring Harbour Laboratory MAPseq/BARseq core facility for RNA sequencing.

### Gene panel design

Candidate marker genes for marmoset thalamic neuron types were selected from marmoset single-nucleus RNA-seq (snRNA-seq) reference data (Dan et al., 2025; Krienen et al., 2023). Counts were normalized to counts per million, and differential markers were computed at both cluster and nested-subcluster levels using MetaMarkers, which ranks genes per group by area under the ROC curve (AUROC), detection rate, and fold change. Genes were retained as cluster-level markers when AUROC > 0.8, detection rate > 0.9, and fold change > 2, and as subcluster-level markers under relaxed criteria (AUROC > 0.7, detection rate > 0.9, fold change > 2). To prioritize genes for compact panels, we additionally selected the top 20 differentially expressed genes per cell type by fold change, and the top three genes per subcluster ranked by detection rate and by a composite score (log[fold change] × AUROC × detection rate); genes marking two or more subclusters were flagged as recurrent markers.

As a complementary, data-driven approach, we used PERSIST (Covert et al., 2023) to identify a compact set of genes that best explains transcriptome-wide variation across thalamic excitatory neurons. Starting from the snRNA-seq reference (44,261 neurons restricted to 3,400 highly variable genes), we constructed a binarized expression layer (expression > 0) and a log1p counts-per-million layer (normalized to 1e6), and split cells 80/20 into training and validation sets. Genes were first coarsely eliminated to a candidate set (target ≈ 500 genes) by increasing the L1 sparsity penalty (λ from 0.01 to ∼0.26, reducing 3,015 → 538 genes), after which a final set of 32 genes was selected (up to 250 training epochs per round). The selected genes were saved (persist_gradients_32.csv) and their expression visualized as dot plots across cell types at multiple taxonomy levels to confirm they captured graded and type-specific variation, and the set was cross-referenced against the marker-derived candidate panels.

Padlock probes were designed as previously described (Chen et al., 2025). Briefly, for each transcript, padlock arms (21–23 nt per arm, GC content 40–60%, melting temperature ≥ 58 °C, no homopolymers) were generated, with each padlock comprising a target-complementary 3′ arm, a fixed backbone, a gene-identifying barcode, a short linker, and a 5′ arm. Low-complexity designs were rejected using a linguistic sequence-complexity threshold (> 0.01). Specificity was verified by BLAST of each padlock’s arms against the marmoset RefSeq RNA database; an off-target was counted when it had a perfect three-nucleotide match flanking the ligation junction, no insertion or deletion within ∼6–7 nt of the junction, and a matched-arm melting temperature above threshold, and probes able to detect any non-target gene were discarded. Up to 12 probes per gene were retained, evenly spaced along the transcript. Each gene was assigned a unique 7-nt RPI barcode (gene-identification index) from the pool of unused barcodes, and the codebook was checked for unique gene–barcode assignment and minimum pairwise Hamming distance, with 10 spare barcodes included as decoding controls.

Each candidate panel was evaluated by k-nearest-neighbor (KNN) label transfer on the snRNA-seq thalamic neuron reference. For each panel, the log-normalized expression matrix was restricted to its genes, and approximate KNN was computed with k = 30 in a leave-one-out scheme. Each cell was assigned the majority label among its neighbors at both cluster and cluster subcluster resolution. Mapping accuracy was quantified as the fraction of cells whose predicted label matched the ground-truth label, and per-type sensitivity (true positives / true cells) and precision (true positives / predicted cells) were computed and plotted. To benchmark performance, the same procedure was applied to highly variable gene baselines as an upper bound, and to Xenium 300-gene panel (Dan et al., 2025) targeting the basal ganglia and 1500-gene whole-brain panels for comparison.

### In situ sequencing

We performed BARseq library preparation as previously described (Sun et al., 2021) with slight modifications. Briefly, the slides with 20 µm tissue sections were fixed in 4% PFA in PBS for 1 hour, washed in PBS and PBST (PBS with 0.5% Tween-20), dehydrated in ethanol, then rehydrated and reverse transcribed overnight at 37 °C. On the second day, the cDNAs were crosslinked, then gene padlocks were hybridized and ligated. For projection mapping experiments (**Fig. 3** and **Fig. 4**), this was followed by hybridization, gap-filling, and ligation of barcode padlocks, whereas for gene-only experiments (**Fig. 1**), we proceeded directly to the next step. We then hybridized an RCA primer and performed rolling circle amplification with phi-29XT (New England Biolabs) for 2 hours at 40 °C, followed by a second round of rolling circle amplification with phi-29 DNA polymerase (Thermo Fisher) at room temperature overnight. On the third day, we then crosslinked the samples and proceeded to in situ sequencing. A detailed step-by-step protocol is provided at https://dx.doi.org/10.17504/protocols.io.kqdg3ke9qv25/v1 (Wang and Isogai, 2025).

In situ sequencing was performed on a custom in situ sequencer, built on top of a Nikon Ti-2E microscope with Photometrics Kinetix cameras, a NanoScan OP400 objective piezo (Prior Scientific), a Celesta 7-line laser (Lumencor), an Xlight v3 (Crest Optics), and custom heating chambers from Bioptechs. Sequencing was performed using Miseq nano v2 kits (Illumina), following previously established protocols (Sun et al., 2021).

### In situ sequencing data processing and quality control

The BARseq image-processing pipeline performs a sequence of denoising and registration operations to align multichannel fluorescence images across sequencing and in-situ hybridization cycles. Image denoising begins with Noise2Void (N2V), a self-supervised deep learning approach that denoise each 2D sequencing channel independently using pre-trained convolutional neural networks trained on the specific experimental modality, while preserving non-fluorescence reference channels (e.g., transmitted-light DIC) unprocessed. Sequencing cycle registration operates at 40× magnification and combines channel profile correction, background subtraction, and translation-based image alignment. Channel shift corrections account for dichroic-induced aberrations by applying per-channel affine transformations. Background subtraction uses rolling-ball morphological opening with a radius of approximately 20 pixels to remove uneven illumination and tissue autofluorescence while preserving cellular signal. Channel bleedthrough is corrected by dividing each channel by empirically measured bleedthrough profiles (stored as calibration matrices), which quantify the fractional signal leakage from one channel into others. Image alignment is achieved via normalized cross-correlation on high-frequency features: images are divided into 256×256 blocks, blocks are ranked by total intensity and the brightest fraction (controlled by subsampling rate) are used for registration to avoid bias from bright soma regions, correlation peaks are refined at 5× upsampled resolution using FFT-based convolution, and translation offsets are estimated from the peak of the mean correlation across blocks, with optional intensity thresholding to exclude pixels above a brightness threshold (soma masking). Barcode-to-gene registration aligns barcode sequencing cycles to registered gene expression reference cycles by computing translation-only alignment between a single reference channel of each modality using normalized cross-correlation on edge-cropped regions, then applying this 2D translation uniformly to all barcode channels. In-situ hybridization (hyb) cycle registration to the first sequencing cycle incorporates median filtering for denoising, rolling-ball background subtraction at approximately 30 pixel radius, optional processing of nuclear counterstain channels, bleedthrough correction using hyb-specific channel profiles, and an additional high-pass filtering step (rolling-ball subtraction at 30 pixel radius) to isolate rolony features before translation-based registration via multimodal optimizer. Tile stitching reconstructs whole-slice images from the grid of 40× fields of view using phase-correlation-based image registration (Microscopy Image Stitching Tool / MIST algorithm in Fiji), which estimates tile positions by cross-correlating overlapping regions, refines positions via exhaustive search over 32 candidate starting offsets, and outputs the stitched image along with transformation matrices. Visual validation of registration quality is performed by warping the aligned RGB composite images from each 20× field using the stitching transformation matrices, compositing them onto a canvas via maximum-intensity projection with tile boundary markings, downsampling by 5× for rapid visualization, and saving the result for manual inspection to confirm alignment fidelity before proceeding to downstream analysis.

### Iterative clustering and cell-type annotation

For transcriptomic analysis, the files are then exported to h5ad and rds format for further processing in python. Cells with fewer than 5 detected genes or fewer than 30 total counts, or lacking a section assignment, were excluded. We normalized counts by logcp10 and computed principal components without highly-variable-gene selection or scaling. Principal components were batch-corrected across brains with Harmony (theta = 2), and a shared nearest-neighbor graph (15 neighbors, 30 components) was clustered with the Leiden algorithm (resolution 1.0), yielding 18 clusters. We then combined two H1 clusters that contained glutamatergic thalamic neurons, recomputed PCA, and subclustered at resolution 0.5, giving seven H2 groups. Each H2 group (excluding one low-quality group) was independently subclustered at resolution 1.0, yielding 57 H3 clusters. Marmoset and mouse BARseq data (with projection mapping) were clustered using a similar approach.

To map clusters to cell types in reference datasets, we labelled each BARseq cell by k-nearest-neighbor mapping (k = 15) onto a reference taxonomy using ‘BiocNeighbors::queryKNN’. We restricted both datasets to the genes shared between the panel and the reference, excluding the broad neurotransmitter markers Slc17a7, Slc17a6, Gad1 and Gad2. To correct for platform differences in capture, we rescaled each reference gene by the ratio of its mean count in the query to its mean count in the reference (f_gene = rowMeans(query)/rowMeans(reference)) and then column-normalized both matrices to zero mean and unit length (cosine normalization, without rank transformation). Each cell received the majority label among its 15 neighbors, with a confidence equal to the fraction of neighbors supporting that label. We summarized the correspondence between BARseq and reference types with the Jaccard index, J = n_AB / (Σ_row + Σ_col − n_AB). We mapped the marmoset projection neurons to a marmoset thalamic-excitatory reference and the mouse neurons to the Allen ABC atlas thalamic glutamatergic neurons at both the supertype and cluster levels.

### Demarcation of thalamic nuclei

We drew nucleus and subdivision borders manually on registered sections using marker-gene landmarks (principally PVALB and CALB1) and the stereotaxic atlas, and applied them across all sections to assign each neuron to a nucleus. We reconstructed sections in a common space by assigning each an anterior–posterior coordinate from landmarks (medial habenula, lateral geniculate, MD, lateral dorsal nucleus), verifying that physical inter-section distances (20 µm per section) matched the atlas progression. For further details see **Supp. Note 3, ED Fig. 3**.

### Gene-expression gradient test across nucleus borders

To distinguish a discrete border, a compositional gradient, and a continuous single-cell gradient, we analyzed representative sections containing pairs of neighboring nuclei (marmoset: anteroventral/ventroanterior and medial/lateral pulvinar; mouse: lateral-posterior/posterior). We normalized gene expression to log2(1 + 100 × counts/total) for the marmoset in situ sequencing data and log2(1 + 1000 × counts/total) for the mouse ABC atlas data and reduced both to 30 principal components. We fit a regularized linear discriminant (‘fitcdiscr’, Delta = 1×10□^3^, Gamma = 0.5) to the two areas in this space and projected all cells onto the discriminant. We binned cells into 15 discriminant-score bins and 25 spatial bins (15 for the mouse) over the 2nd–98th percentile of the physical axis (10th–90th for the mouse) within a band of 500 µm (200 µm for the mouse), and asked whether the observed per-bin distribution was reproduced by a purely compositional model in which each bin draws a fraction f_b = (m_b − m_pure2)/(m_pure1 − m_pure2) (clamped to [0,1]) of its cells from the first pure pool and the remainder from the second. We assessed robustness to detection efficiency by binomially subsampling counts at rates of 1.0, 0.5, 0.25, 0.10 and 0.05. To build a distribution that reflects a compositional gradient, we defined "pure" reference pools as the 20% neurons at the extreme end along the spatial axis between the two areas, then reconstructed each bin using these reference cells such that they preserve the mean discriminant score. Separately, we computed the first principal component of gene expression and plotted the probability density along this axis.

Two complementary statistics were computed. First, one- and two-component Gaussian mixtures were fitted to a tight interior window (± 2 bins), and the BIC difference dBIC = BIC(1) − BIC(2) was mapped to a two-peak preference P(compositional) = 1/(1 + exp(−dBIC/2)), with a smoothed nonparametric bootstrap (1,000 resamples, Gaussian jitter h = 0.5·s.d.·n^(−1/5)) giving the ± 1 s.d. interval. Second, a Silverman critical-bandwidth test for unimodality was run on a wider window (± 5 bins marmoset, ± 3 bins mouse), estimating the smallest Gaussian-kernel bandwidth yielding ≤ 1 mode and testing H0 (unimodality) by variance-corrected smoothed bootstrap (500 resamples). Because the Gaussian mixture comparison is effectively a test of Gaussianity that broad unimodal bridges can spuriously fail, the two-peak versus bridge call was carried by the Silverman test, with P(compositional) reporting direction and magnitude.

### Registration of dissected tissue to reference templates

We dissected the entire marmoset cortical hemisphere that is ipsilateral to the injections into 94 cubelets (**Supp. Table 4**). Frontal, temporal, and parietal regions were dissected at a higher resolution, because the nuclei we injected in have strong projections to these areas based on previous tracing studies(Córdoba-Claros et al., 2025b). For each cubelet, a two-dimensional region of interest (ROI) was drawn on the matched cortical slice from the Marmoset Brain Mapping Template space v3 (Liu et al., 2021) and extrapolated in three dimensions to match the section thickness. The MRI volume and ROIs were then and projected onto the cortical surface with 3dVol2Surf (AFNI)(Cox, 1996). To provide a canonical parcellation reference for each cubelet, and to account for inter-animal variability and dissection imprecision, the Paxinos cortical atlas was overlaid on the resulting flatmap.

For the mouse, each dissection slab was registered to the corresponding plane of the Allen Mouse Brain CCF manually, and the dissected region was delineated on that plane. The dissections were then flattened to a flatmap using the ccf_streamlines package (https://github.com/AllenInstitute/ccf_streamlines.git).

### Projection data processing

Projection barcodes were error-corrected as previously described (Kebschull et al., 2016), and only barcodes with at least 5 UMI counts in one cubelet were kept. Barcodes seen in the in situ sequencing data were then matched to those in the dissected cubelets, allowing no mismatch, to construct projection matrices. To estimate the rate of wrong matches due to barcode collision, we shuffled each digit across the barcodes and performed the same matching to the projection data. Most downstream analysis were then performed on the projection matrix after normalizing barcode counts in each cubelet by the recovered spike-in molecule numbers, followed by row normalization. For projection domains, we performed non-negative matrix factorization on log2(1 + 1000 × normalized counts). For some analyses, we binarized the projection matrices using a fixed threshold (projection domains ≥ 3, cubelet-level projections ≥ 0.1).

To visualize the spatial organization of projections to target domain, we estimated a smooth probability map over the flatmap of registered soma positions for each target domain. Sections were binned into consecutive groups of five along the anteroposterior axis, and maps were estimated independently per target domain and bin. For each domain–bin with at least five labelled neurons, we discretized the projections and modelled the probability of the module label as an anisotropic 2D Gaussian, *p*(*x*) = *A* exp(−½ (*x*−μ)□ Σ□^1^ (*x*−μ)), fitted by maximum likelihood under a Bernoulli model (Nelder–Mead), with penalties discouraging extreme ellipticity and regularizing Σ and μ towards the empirical distribution of labelled somata.

### Validation of projections against bulk mouse anterograde tracing

To compare the mouse BARseq data to the Allen Connectivity Atlas, we summed row-normalized barcode counts from neurons per source nucleus, generating a source nucleus by cortical target pseudobulked projection matrix. From the Allen Connectivity Atlas, structure-unionized data were retrieved from the brain-map RMA API, using ipsilateral projection signal excluding the injection site, summed over the acronyms mapped to each of the 11 targets. For each experiment, we used the projection_volume, normalized by sum within each experiment. When multiple experiments existed for the same source nucleus, we averaged the normalized projection_volume vectors across the experiments. We then calculated Pearson correlation between the pseudobulked BARseq data and the normalized projection_volume of the Allen Connectivity Atlas. To estimate a ceiling for the correlation from multiple experiments on the same source nuclei in the Allen Connectivity Atlas, we held out each experiment, calculated the average projection vector from the remaining experiments, and calculated the correlation in the same way as the BARseq/Allen comparison. The Allen Connectivity Atlas experiments used were:

LD: 272969333, 113784293, 272967913
MD: 272970747, 114291646, 156931568
PO: 174781014, 100147785, 180708524
LP: 146658879, 266585624
MG: 180520257 (excluded from the Allen/Allen pairwise correlation, which required same-source replicates)

### Validation of projections with single-neuron axonal reconstructions

We compared BARseq projections with 99 single-neuron reconstructions from the MouseLight and SEU-Allen datasets. We transformed each single-neuron morphology, as defined in the SWC files onto a two-dimensional flatmap with normalized cortical depth. To co-cluster BARseq data with scTracing data, we randomly subsampled BARseq data to 99 neurons, reduced the resolution of the scTracing data to match BARseq, and the two were pooled and clustered jointly by Ward’s hierarchical clustering on Euclidean distances, across a range of cluster numbers (2–12). Both log-transformed and row-normalized projection matrices produced qualitatively similar co-clustering.

For visualization, the pooled neurons were embedded in two dimensions by t-SNE and colored by dataset of origin. To identify directly comparable individual neurons, we row-normalized projection profiles in both datasets and, for each BARseq neuron, found its nearest traced neuron by Euclidean distance (*k*-nearest-neighbor search).

### Validating marmoset BARseq neuron projection patterns against published bulk anterograde tracing

We compared the projection patterns of BARseq neurons with published bulk anterograde tracing by matching BARseq soma locations to bulk injection sites and assessing projection correspondence. For each published injection site, we retraced its boundary onto the AP-matched sections in our data and assigned all enclosed BARseq neurons, extending the window 100 µm anteriorly and posteriorly. Given injection extents of ∼500–800 µm in plane, this provides a conservative estimate of injection volume. To register the two datasets, we manually retraced all bulk-traced projection zones onto our MRI-based flatmap template using Paxinos atlas annotations. For each neuron, we defined projection targets as continuous zones of barcode signal across cubelets with a single local maximum, and measured the cubelet distance from each target to the nearest matching bulk-traced area, averaging across targets for neurons with multiple projections (**Supp. Table 5**; **Supp. File 2**).

### Spatial organization comparison between mouse and marmoset

We computed each neuron’s selectivity index as the max of its row-normalized projection falling in a single target domain. Separately, we computed each neuron’s projection coverage as its effective number of targets (two raised to the Shannon entropy of its projection) divided by the number of targets.

To test whether neighboring neurons share projections, we compared the similarity of each neuron’s projection to those of its nearest spatial neighbors. For each neuron we computed the mean normalized Euclidean distance between its projection patterns to the target domains and those of its 8 nearest neighbors within 200 µm.

To test how spatial locations can inform of projections, we computed a spatial-informativeness score defined as 1 − (E_spatial / E_random), where E_spatial is the cosine error between a neuron’s projection pattern and the pattern predicted from the mean of its up to 10 nearest neighbors within 500 µm, and E_random is the corresponding cosine error when the prediction is instead formed from an equal number of randomly chosen neurons, averaged over 50 random draws.

### Spatial organization of projections across nucleus borders

To test how nucleus identity contributed to projection differences, for each neuron we computed the cosine similarity of its projections to target domains to those of its nearest spatial neighbors—using the 2 nearest neighbors in the mouse and the 5 nearest in the marmoset. We grouped source-neighbor pairs based on their nucleus identity into the same-nucleus neighbor (S_N_) and cross-nucleus neighbor (C_N_) groups. For each pair, we then shuffled projection patterns among cells of the same area (200 shuffles) to obtain measures for the same-nucleus random (S_R_) and cross-nucleus random (C_R_) groups.

### Cellular versus compositional projection transitions

To distinguish whether projections vary as cellular level gradients or compositional gradients across borders, we compared two generative models of a one-dimensional projection score laid out along the spatial axis connecting the centroids of a pair of nuclei. In the marmoset data, axial position was rank-transformed to account for the much larger span of the pulvinar compared to the MD, but raw positions were used for the mouse data. A projection score was computed as the least-squares (Moore–Penrose) projection of the z-scored projections onto axial position, and lightly jittered to break ties from near-single-target vectors. Two models were fitted: a shifting single-Gaussian model with a per-bin mean and a single shared variance (B + 1 parameters), and a compositional model of two globally fixed Gaussians with a per-bin mixing weight, fitted by expectation-maximization (B + 4 parameters). Model preference was defined as the difference in Bayesian Information Content between the two models. Because ΔBIC is not interpretable in isolation, it was calibrated against two parametric bootstraps (200 each): a single-Gaussian null and a compositional-expected reference simulated from the respective fitted models. We defined a compositional index as the ΔBIC, normalized to the range of the differences across the two parametric bootstraps. Aa one-sided p value against the cellular null was calculated by comparing to the bootstrap. As positive controls, we constructed a synthetic compositional mixture based on the mouse PO/LP data, and also applied the same analysis to the LP/midbrain neurons.

### Identifying gene expression axes associated with projections

We first computed linear-discriminant axes in gene expression space that separate neurons grouped by their dominant projection domains. We extracted the first 15 principal components from logcp100 gene expression data, assigned each neuron within the thalamic nuclei to its strongest projection target domain, and performed linear discriminant analysis to identify the first two discriminant axes. We correlated the first axis with the anterior–posterior position of each neuron’s projection centroid, assessing significance against a null that shuffled the cortical cubelet positions (10,000 permutations, two-tailed), and tested whether the second axis separated medially projecting (PL/IL, ACA) from dorsolaterally projecting neurons with a Wilcoxon rank-sum test.

To relate gene expression directly to projection topography, we calculated each neuron’s projection target position w as the projection-weighted mean of the flatmap cubelet coordinates; for mouse the AP coordinate was the flatmap y-axis, whereas for marmoset each flatmap centroid was mapped through a per-pixel flatmap-to-MRI registration to real MRI AP coordinates. Because AP and ML are spatially correlated across projection neurons, we reweighted the sample by inverse-propensity weighting, binning AP and ML each into five quantile bins and assigning each cell a weight equal to the product of the marginal bin probabilities divided by the joint, clipped at the 0.98 quantile. On a weighted resample of 20,000 draws, a single-target partial-least-squares regression of gene expression onto AP was fitted with four latent components directly on the full gene panel, yielding one gene-weight vector per species (mouse, 188 genes; marmoset, 152 genes) aligned posterior-positive. To compare across species, we restricted to the 45 genes present in both BARseq panels, and scored the same cells on both axes.

### Statistics

Group comparisons used two-sample Kolmogorov–Smirnov tests, Wilcoxon rank-sum tests or paired Wilcoxon signed-rank tests as indicated; associations used Pearson or Spearman correlation; and classifier performance used the area under the ROC curve with cross-validation. P values are reported with Holm-Bonferroni or Bonferroni correction, as indicated in the text, or without correction if no correction was mentioned.

### Data and Code Availability

Bulk projection RNA sequencing data are in the process of being deposited to SRA. In situ sequencing data are deposited and being processed at Brain Image Library. Processed data and all scripts used for data processing and analysis are available upon request.

## Acknowledgements

We thank Kevin Marche and Xiaofeng (Lisa) Zhang, and the staff of the Central Animal Facility (CAF) and the Veterinary Medicine and Resources Branch (VMRB, NIMH) for providing care and support for the research animals; Aixin Zhang for technical support; C. Gerfen, J. Gillis, B. Averbeck, T. Shimogori, and H. Zeng for their helpful comments on the manuscript. This work was supported by the National Institutes of Health grants ZIAMH002984 (J.A.B.), 5U01NS132161 (X.C., J.M.K), 5R01MH133181 (X.C.), DP2MH132940 (X.C.), a Klingenstein-Simons Fellowship (J.M.K), a Packard Fellowship (J.M.K), and by a Human Frontier Science Program (HFSP) Program Grant (J.A.B.). A.Y.F, N.M.K, J.T.S and J.A.B are supported by the National Institutes of Mental Health (NIMH) Intramural Research Program. A.Y.F is in part supported by the Commonwealth of Australia through an Australian Government Research Training Program Scholarship. The authors thank the Allen Institute founder, Paul G. Allen, for his vision, encouragement, and support. The content is solely the responsibility of the authors and does not necessarily represent the official views of the National Institutes of Health.

## Author contributions

Conceptualization: A.Y.F., J.A.B., X.C.; Investigation: A.Y.F., J.T.S., N.M.K., S.C., T.J., B.O., A.A.O., D.R., H.Z., J.A.B., X.C.; Methodology: A.Y.F., J.S., H.Z., J.A.B., X.C.; Resources: A.Y.F., M.M., H.Q., M.T.S., H.Z., J.A.B., X.C.; Validation: A.Y.F., X.C.; Data curation: A.Y.F., H.Z., X.C.; Software: A.Y.F., J.T.S., J.H., X.C.; Formal analysis: A.Y.F., J.T.S., X.C.; Visualization: A.Y.F., J.T.S., J.A.B., X.C.; Writing – original draft: A.Y.F., J.A.B., X.C.; Writing – review and editing: A.Y.F., J.A.B., X.C.; Supervision: M.M., H.Z., J.M.K., J.A.B., X.C.; Project administration: J.A.B., X.C.; Funding acquisition: J.M.K., J.A.B., X.C

## Extended Data Figures

**Extended Data Figure 1.**
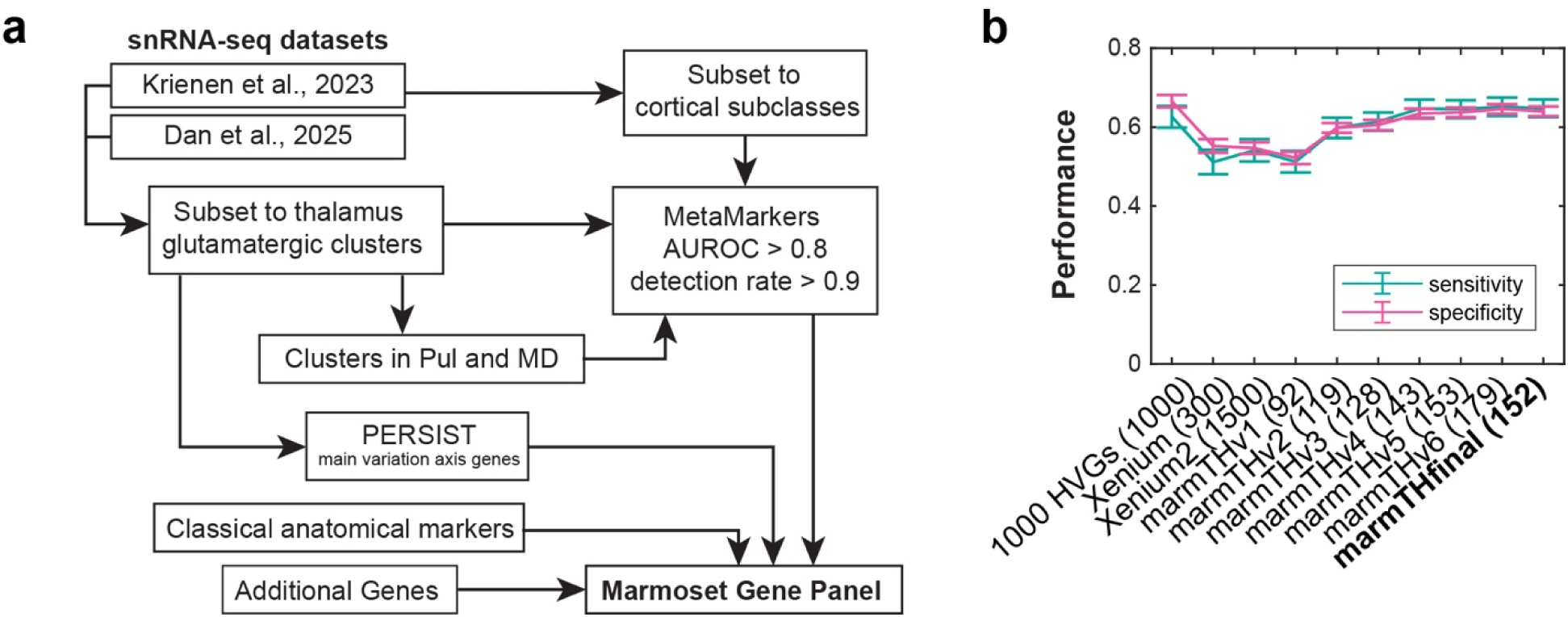
Gene panel performance. (**a**) Schematic of the workflow for designing the marmoset gene panel. We first identified thalamic glutamatergic neuron clusters, cortical neurons, and clusters that are enriched in MD and the pulvinar in two single nucleus RNA-sequencing data (Dan et al., 2025; Krienen et al., 2023). We then applied three complementary approaches, based on MetaMarkers, PERSIST, and manual selection to find cell type marker genes in the cortex and the thalamus, and genes that reflect major axes of variations in the thalamus. (**b**) In silico assessment of sensitivity (*blue*) and specificity (*pink*) of cluster assignment, using various gene panels against a snRNA-seq thalamus dataset (Dan et al., 2025). The gene panels included 1000 highly variable genes (HVGs, chosen as a benchmark), a 300-gene Xenium panel targeted to basal ganglia (Dan et al., 2025), a 1500-gene Xenium panel expanded from the 300-gene panel, and various BARseq panels with the indicated number of genes, which were designed to resolve thalamic excitatory neurons.

**Extended Data Figure 2.**
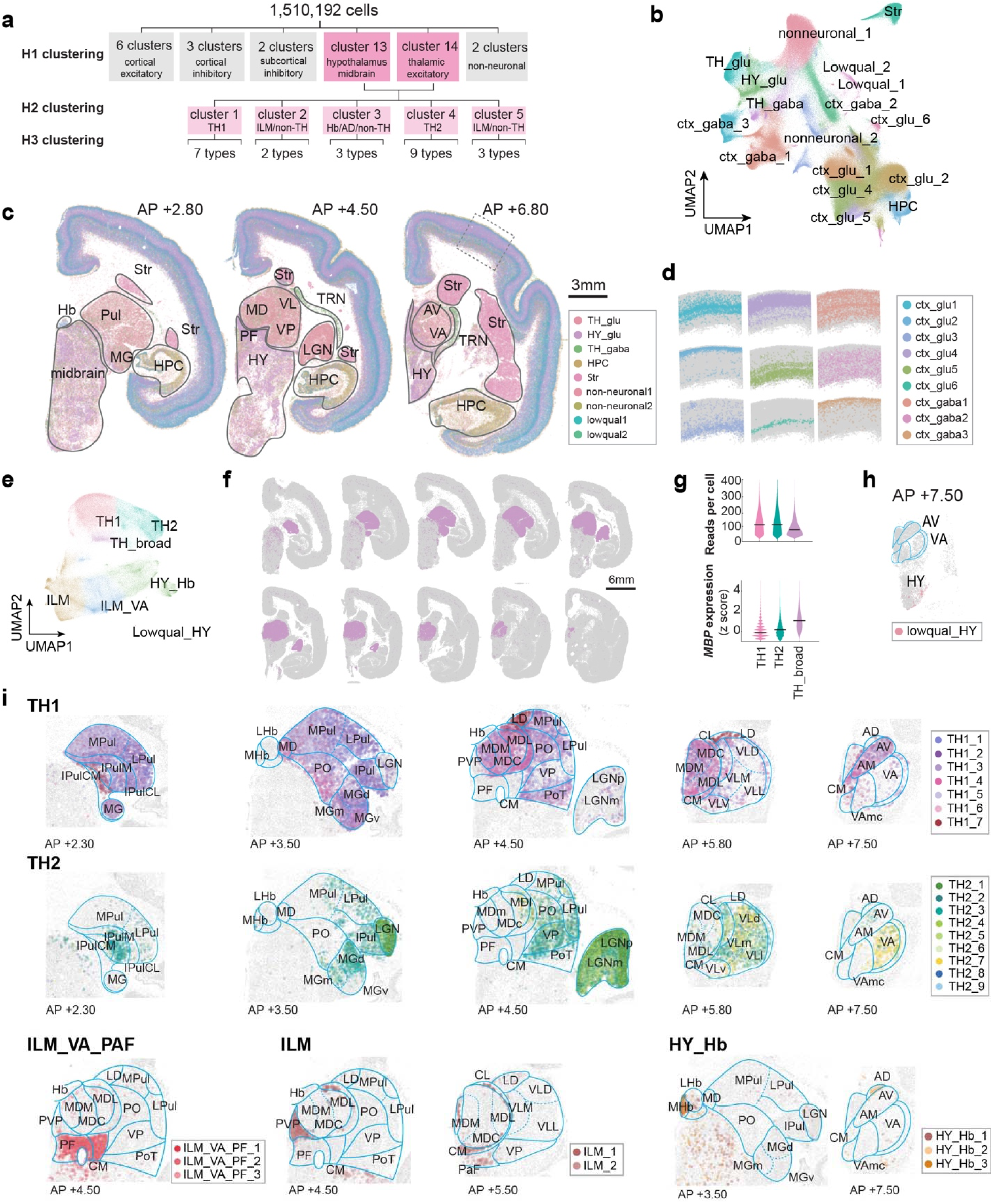
Hierarchical clustering and spatial distribution of neuron types. (**a**) Workflow for hierarchical clustering. (**b**) UMAP plot of gene expression of all cells after QC. Cells are colored by H1 clusters. (**c**) Distribution of H1 cell types on three representative slices. Scale bar = 3 mm. AV, anteroventral nucleus of the thalamus; Hb, habenula; HPC, hippocampal cortex; HY, hypothalamus; LGN, lateral geniculate nucleus; MD, mediodorsal nucleus of the thalamus; MG, medial geniculate nucleus; PF, parafascicular nucleus; Pul, pulvinar; Str, striatum; TRN, thalamic reticular nucleus; VA, ventroanterior nucleus of the thalamus; VL, ventrolateral nucleus of the thalamus; VP, ventroposterior nucleus of the thalamus. (**d**) The laminar distribution of H1 cortical cell types in the lateral cortex. The area shown corresponds to the dashed box in (c). (**e**) UMAP plot of gene expression of cells after subsetting to H2 clusters that contain thalamic excitatory neurons. Cells are colored by H2 clusters from the second level of clustering. Labels indicate H2 clusters. (**f**) Distribution of TH_broad clusters across all slices. (**g**) Violin plots showing the distribution of reads per cell (*top*) and z-scored expression of *MBP* (*bottom*) in TH1, TH2, and TH_Broad clusters. (**h**) Spatial distribution of Lowqual_HY cluster in slice at AP +7.5 mm, the slice where most of Lowqual_HY cells are found. (**i**) Distribution of thalamic excitatory neuron types from each indicated H2 cluster in representative slices. Lines indicate nucleus borders. Colors indicate neuron types (H3 clusters). H2 cluster names are indicated on the upper left corner. AD, anterodorsal thalamic nucleus; AM, anteromedial thalamic nucleus; CL, centrolateral thalamic nucleus; CMn, central medial thalamic nucleus; lHb, lateral habenula; mHB, medial habenula; LD, laterodorsal thalamic nucleus; LGNm, lateral geniculate nucleus, magnocellular layer; LGNp, lateral geniculate nucleus, parvocellular layer; MDc, mediodorsal thalamic nucleus, central part; MDl, mediodorsal thalamic nucleus, lateral part; MDm, mediodorsal thalamic nucleus, medial part; MGd, medial geniculate nucleus, dorsal part; MGm, medial geniculate nucleus, medial part; MGv, medial geniculate nucleus, ventral part; PO, posterior thalamic nucleus group; PoT, posterior thalamic nucleus group, triangular part; IPulCL, inferior pulvinar, caudolateral part; IPulCM, inferior pulvinar, caudomedial part; IPulM, inferior pulvinar, medial part; LPul, lateral pulvinar; MPul, medial pulvinar; PVP, paraventricular thalamic nucleus, posterior part; VA, ventral anterior thalamic nucleus; VAmc, ventral anterior thalamic nucleus, magnocellular part; VLd, ventral lateral thalamic nucleus, dorsal part; VLl, ventral lateral thalamic nucleus, lateral part; VLm, ventral lateral thalamic nucleus, medial part; VLv, ventral lateral thalamic nucleus, ventral part.

**Extended Data Figure 3.**
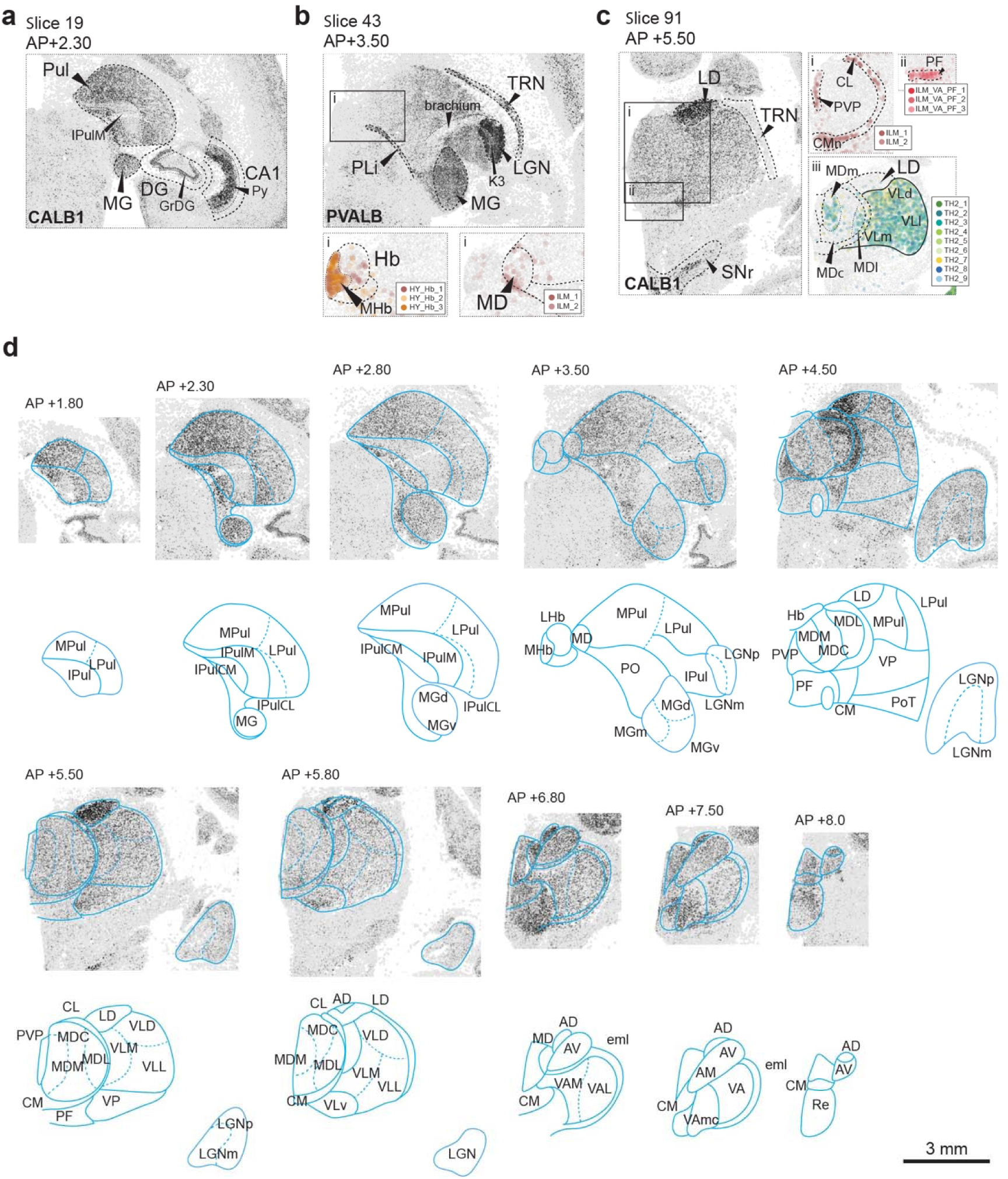
Demarcation of nucleus borders in marmoset. (**a**)-(**c**) Three representative slices in the dataset showing expression of *PVALB* or *CALB1* (black dots) that delineates nucleus borders. In (b) and (c), colored plots show the indicated neuron types in the boxed areas in the *PVALB* and *CALB1* plots. (**d**) Demarcations drawn on top all slices showing the expression of *CALB1* (*top*), with anatomical labels over the nucleus boundaries (*bottom*). Dashed lines separate nuclei or subdivision that were defined by relatively location and were not based on clear distinction in gene expression in this dataset. eml, external medullary lamina; GrDG, granular cell layer of the dentate gyrus; K3, koniocellular layer 3; PLi, posterior limitans; Re, reunions nucleus of the thalamus; TRN, thalamic reticular nucleus. For other abbreviations, see **ED Fig. 2**.

**Extended Data Figure 4.**
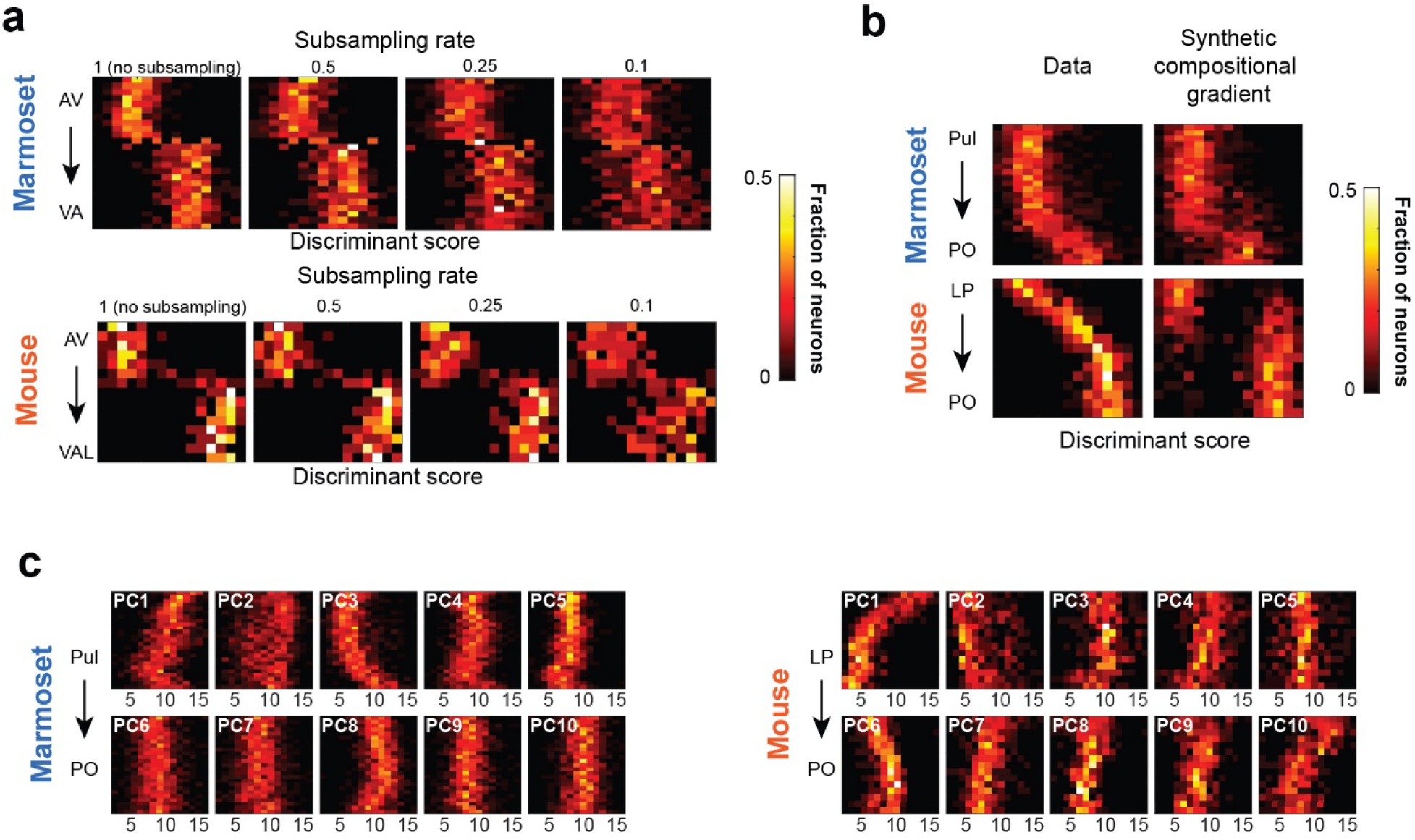
Gene expression gradient in the mouse HOs thalamus. (**a**) Heatmaps showing the distribution of discriminant scores (x-axis) as a function of position along the spatial vector connecting the centroids of the two nuclei (y-axis), with varying degree of mRNA subsampling, in marmoset (*top*) and mouse (*bottom).* Numbers at the top indicate mRNA subsampling rates. (**b**) Heatmaps showing the distribution of discriminant scores (x-axis) in marmoset (*top*) and mouse (*bottom),* as a function of position along the spatial vector connecting the centroids of the two nuclei (y-axis) in real data (*left*) and in a synthetic compositional gradient (*right*). (**c**)(**d**) Heatmaps showing the distribution of principal component scores (x-axis) as a function of position along the spatial vector connecting the centroids of the two nuclei (y-axis) in marmoset (c) and mouse (d)

**Extended Data Figure 5.**
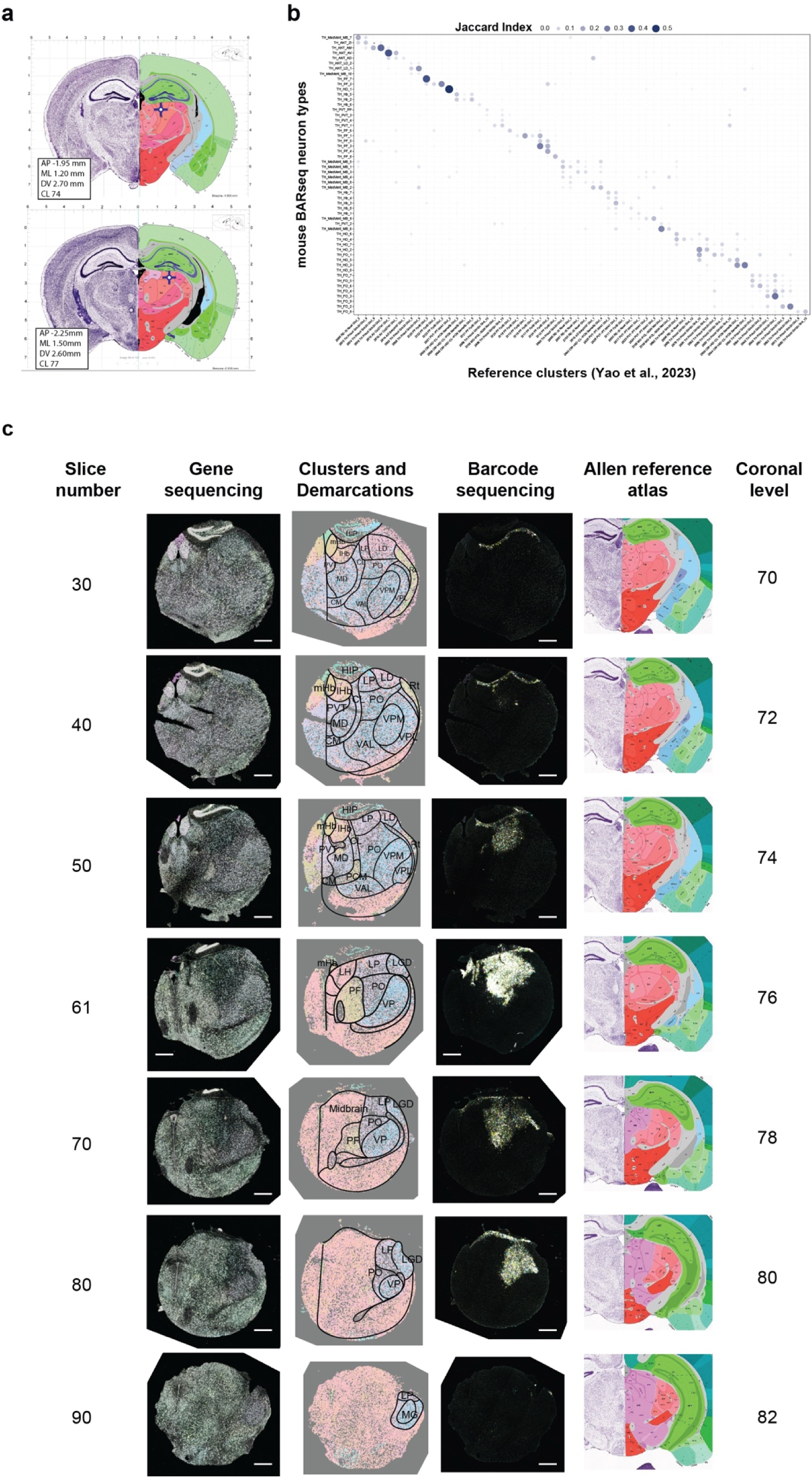
Mouse BARseq slice demarcation. (**a**) Injection coordinates for mouse BARseq, visualized on CCF coronal planes. **(b)** Overlap (Jaccard index) between thalamic excitatory neuron types in the mouse BARseq dataset (y-axis) and clusters in a reference snRNA-seq dataset (Z. Yao et al., 2023).(**c**) Representative slices from mouse 818353 are shown to illustrate area demarcation. The images shown included, from left to right, the first sequencing cycle for genes, coarse cluster labels with demarcation, first sequencing cycle for barcodes, and reference atlas template. Slice numbers and the corresponding coronal levels are shown. Slices from mouse 839168 were demarcated using the same approach. Scale bars = 500 µm.

**Extended Data Figure 6.**
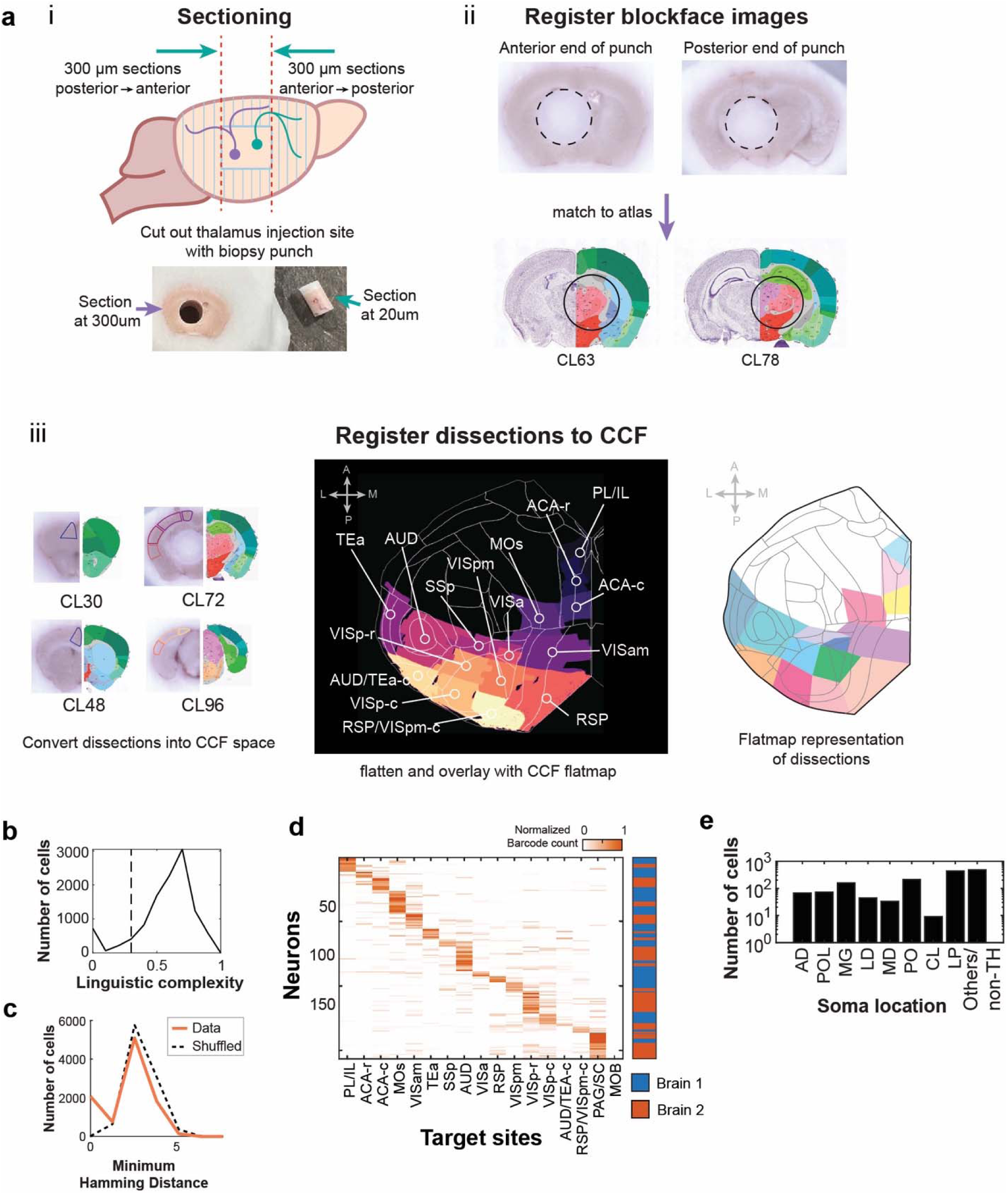
Injection, dissection and quality control for the mouse BARseq experiment. **(a)** Schematic for tissue dissection and sectioning. (*i)* dissection procedure to obtain thin (20 µm) slices from the injection site, and thick (200 µm) sections for cortical dissection. (*ii*) the first and last coronal sections that contained the thalamus (after the thalamus was punched out), matched to corresponding atlas coronal planes. (*iii*) Dissection areas are drawn on matched CCF coronal planes and transformed to flatmap space. Labels on flatmap indicate cubelet names. (**b**) Distribution of linguistic complexity of soma barcodes. (**c**) Distribution of minimum Hamming distances from each soma barcode (orange - Data) or a shuffled barcode set (black dashed line) to barcodes recovered from target sites. (**d**) Projection (row normalized) of neurons from two brains. The first brain (818353) has been resampled to match the number of cells from the second brain (839168), and the combined dataset is sorted by the area that received the strongest projections. (**e**) The number of projection neurons labelled in each thalamic nuclei.

**Extended Data Figure 7.**
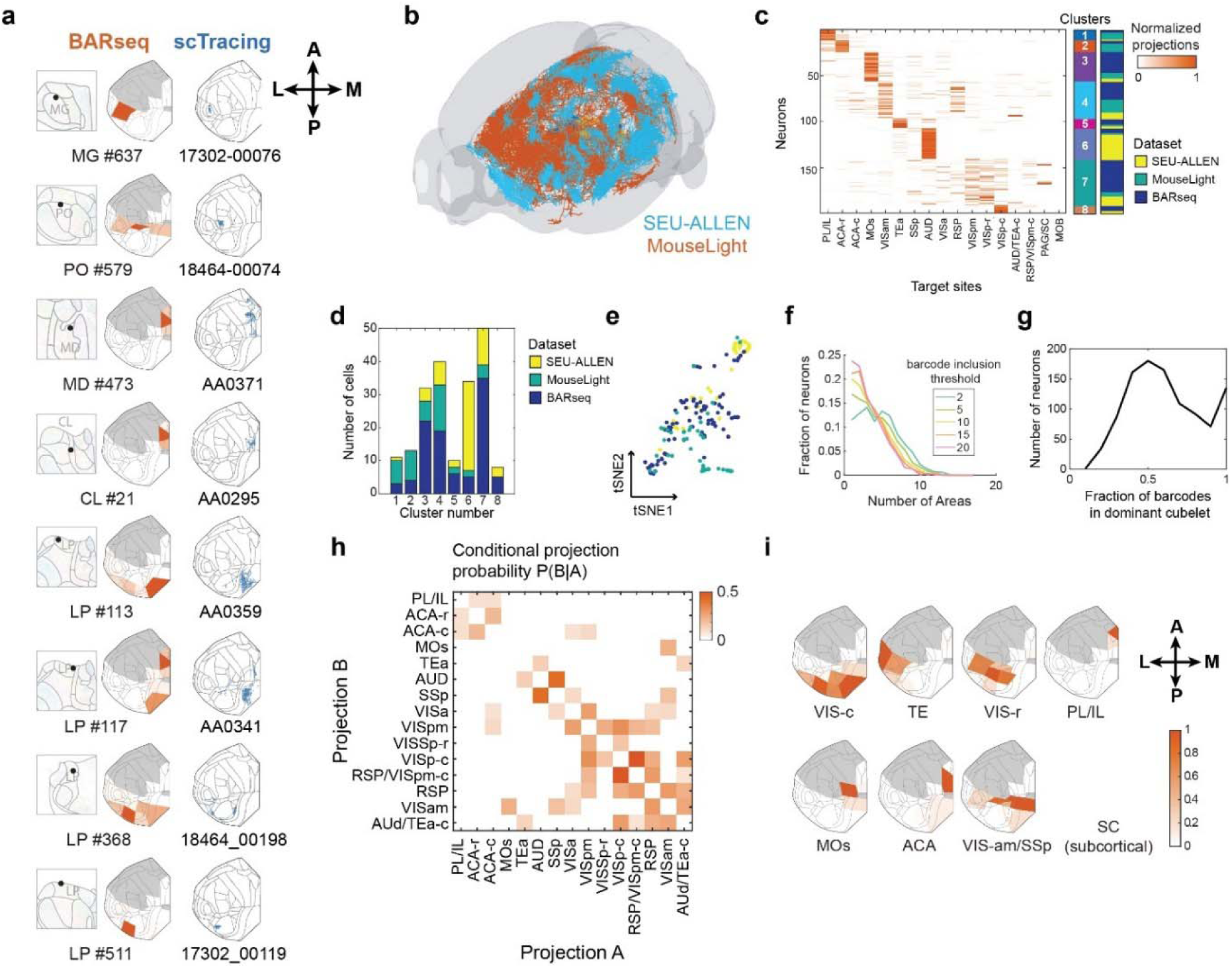
Comparing BARseq data to single-cell tracing data. (**a**) Single neuron examples from BARseq and matching single-cell tracing data. (*left*) BARseq neuron soma locations; (*center*), projection pattern on cortex; (*right*), single-cell tracing axons. (**b**) All 99 traced neurons from SEU-ALLEN (*blue*) and MouseLight (*orange*). (**c**) Row-normalized projections of combined BARseq and single-cell tracing datasets. First colored sidebar indicates eight co-clusters from hierarchal clustering. Second side bar depicts the source of the neurons. (**d**) Distribution of neuron sources across co-clusters. Colors indicate the source of the neurons. (**e**) tSNE plot showing projections of neurons. Colors indicate the source of the neurons. (**f**) Fraction of neurons (y-axis) with projections to the indicated number of areas (x-axis). Line colors indicate thresholds at which a projection is considered present. (**g**) Distribution of the fraction of barcodes in the dominant cubelet. (**h**) Pairwise correlation of projection barcode counts across targets. Areas are ordered by optimal leaf ordering (Bar-Joseph et al., 2001). (**i**) Cortical flatmaps showing the eight NMF-based target domains. The SC target domain is subcortical and could not be shown on a flatmap. Colors indicate normalized barcode counts.

**Extended Data Figure 8.**
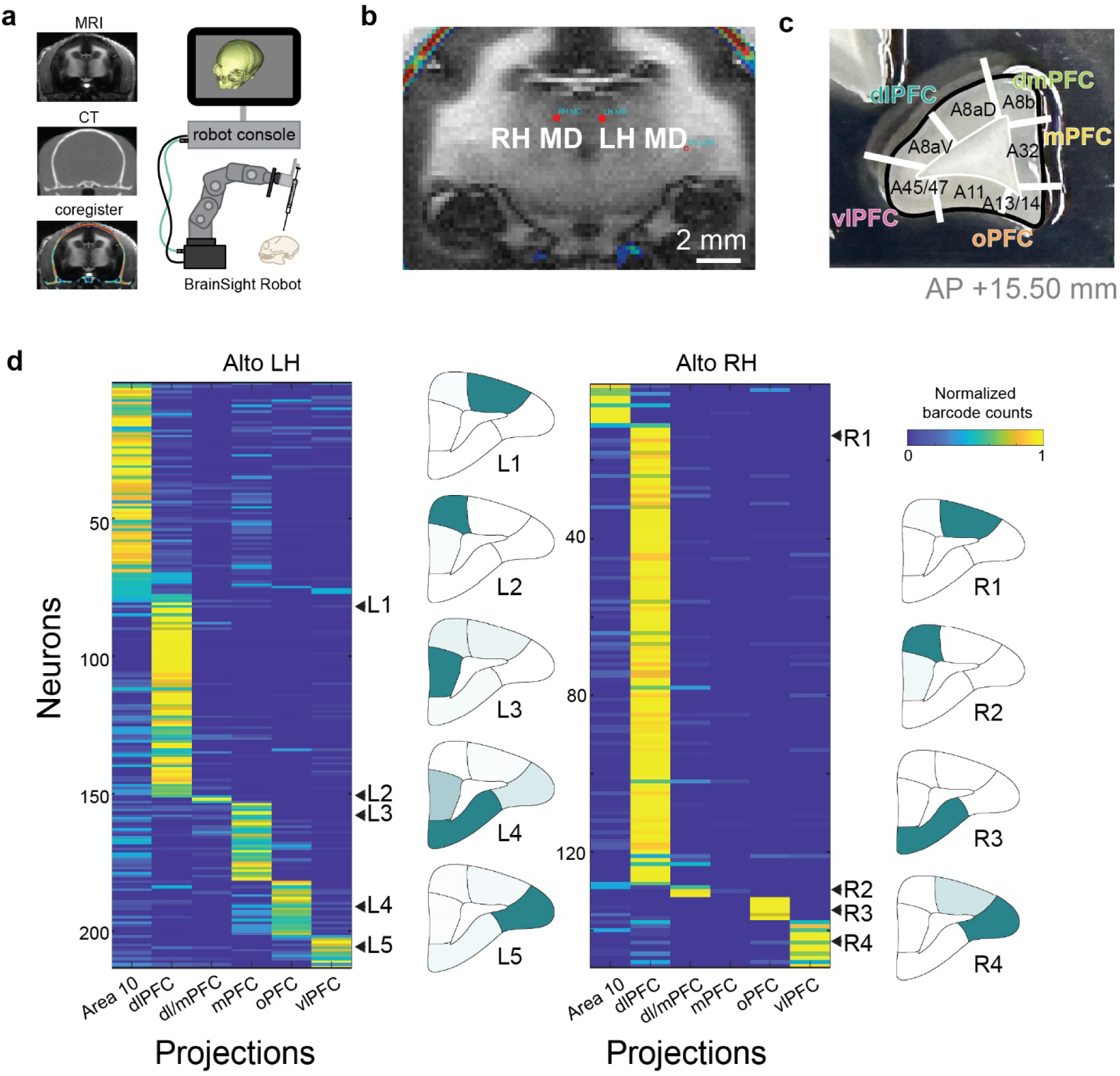
BARseq reveals marmoset MD – PFC projections. (**a**) In surgeries, we co-registered MRI with CT, defined projection sites based on MRI landmarks, then performed surgery using a BrainSight robot. (**b**) A coronal plane of the MRI showing the two MD injection sites. (**c**) An image of a frontal cortex slice, with dissection cubelets marked. Area names for each cubelet, and the pooled region names are indicated. (**d**) Normalized projection strengths of MD neurons that project to the PFC in the two brains. Colors indicate row normalized barcode counts. Columns indicate the pooled regions, and rows indicate single neurons. Example neurons are indicated by arrowheads on the right side of each projection matrix. The projections of these neurons are shown on flatmaps.

**Extended Data Figure 9.**
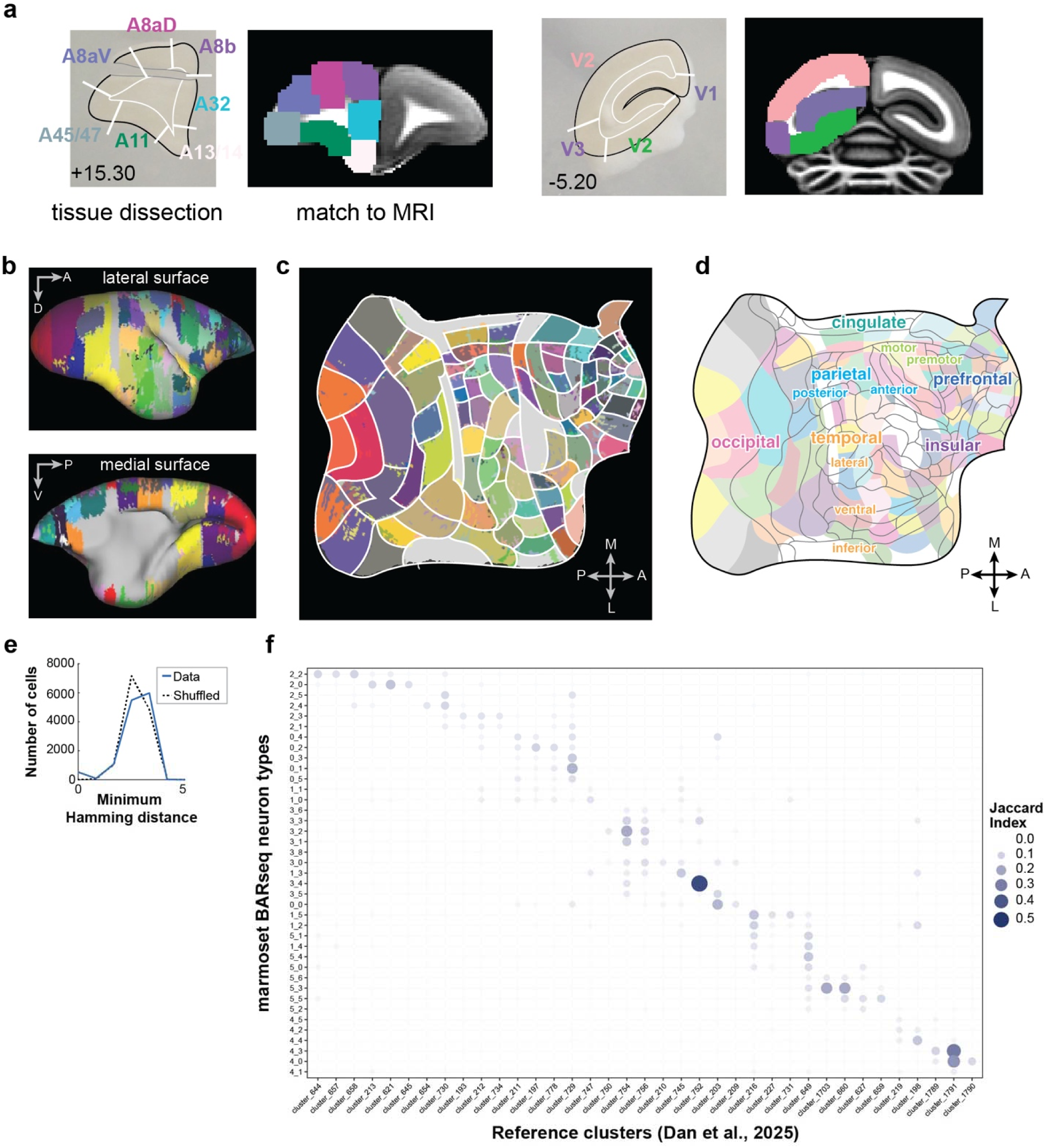
Spatial registration and transcriptomic mapping of marmoset BARseq data. (**a**) Two representative slices that were registered to MRI. In each pair of images, the image on the left shows the slice, with dissection areas overlaid on top. The same areas are registered to the MRI on the image to the right. AP locations are indicated. (**b**)(**c**) All cubelets of a hemisphere, shown on MRI after registration (b) and after mapping onto a flatmap (c). Gray areas indicate areas that did not map onto dissected cubelets. All of these locations correspond to blocking surfaces, and may reflect registration errors across blocks and/or tissue loss during blocking. (**d**) Smoothed dissection cubelets, overlaid with area borders. (**e**) Distribution of minimum Hamming distances from soma barcodes (*blue line*) or a shuffled barcode set (*black dashed line*) to barcodes recovered from dissected cubelets. (**f**) Overlap (Jaccard index) between thalamic excitatory neuron types in the marmoset BARseq dataset (y-axis) and clusters in a reference snRNA-seq dataset (Dan et al., 2025) (x-axis).

**Extended Data Figure 10.**
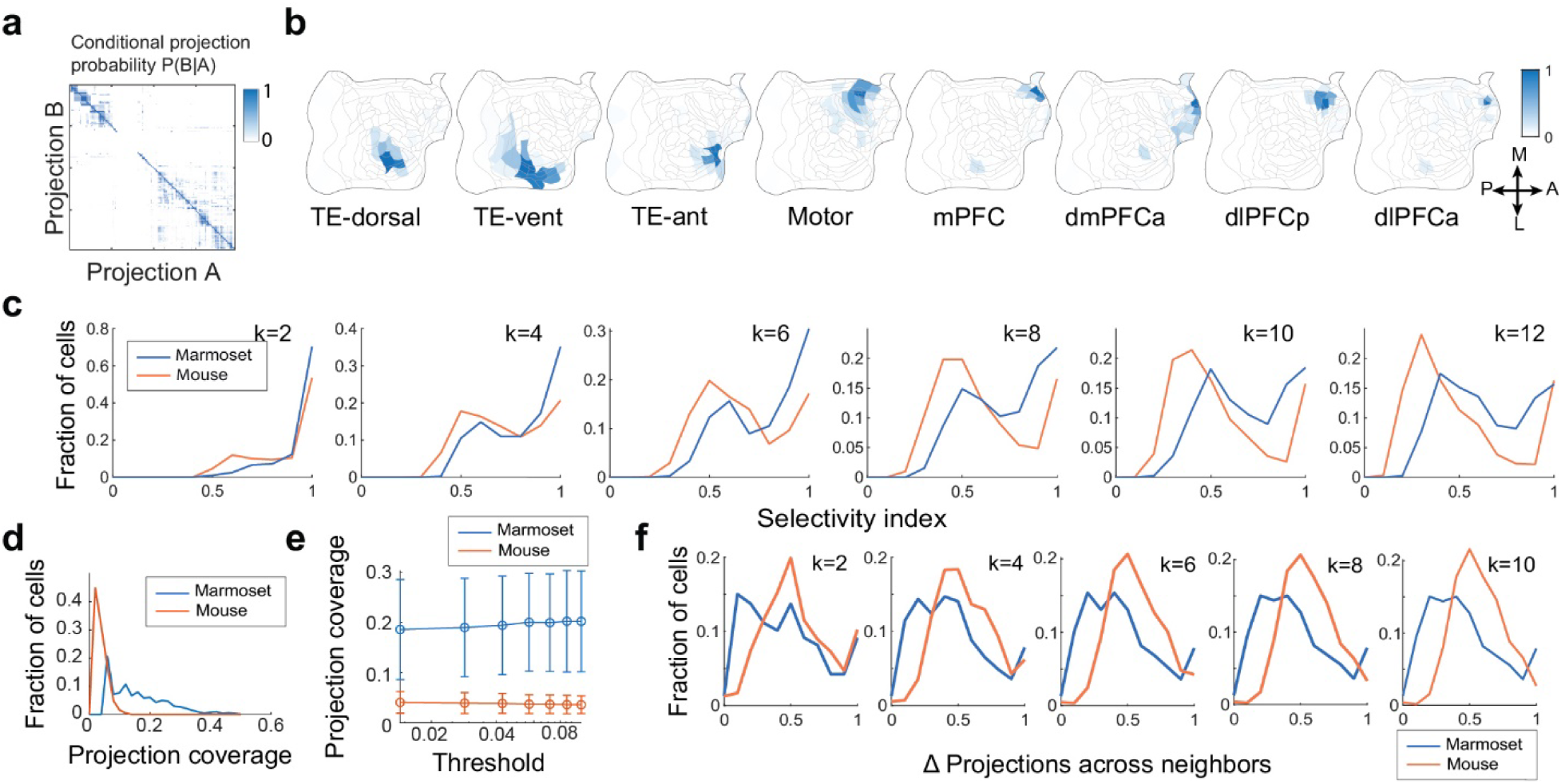
Spatial organization of marmoset thalamocortical projections. (**a**) Probability of a neuron projecting to area B conditioned on it projecting to area A. Areas are sorted using optimal leaf ordering (Bar-Joseph et al., 2001). (**b**) Flatmaps showing the patterns of each projection domain. (**c**) Distribution of selectivity index at mouse (*orange*) and marmoset (*blue*) neurons at different target domain numbers (indicated on top). (**d**) Distribution of projection coverage of mouse (*orange*) and marmoset (*blue*) neurons. (**e**) Means ± standard deviations of projection coverage of mouse (*orange*) and marmoset (*blue*) neurons when filtered by the indicated total projection threshold. (**f**) Distribution of projection differences to spatial neighbors for mouse (*orange*) and marmoset (*blue*) neurons at different neighbor numbers (indicated on top).

**Extended Data Figure 11.**
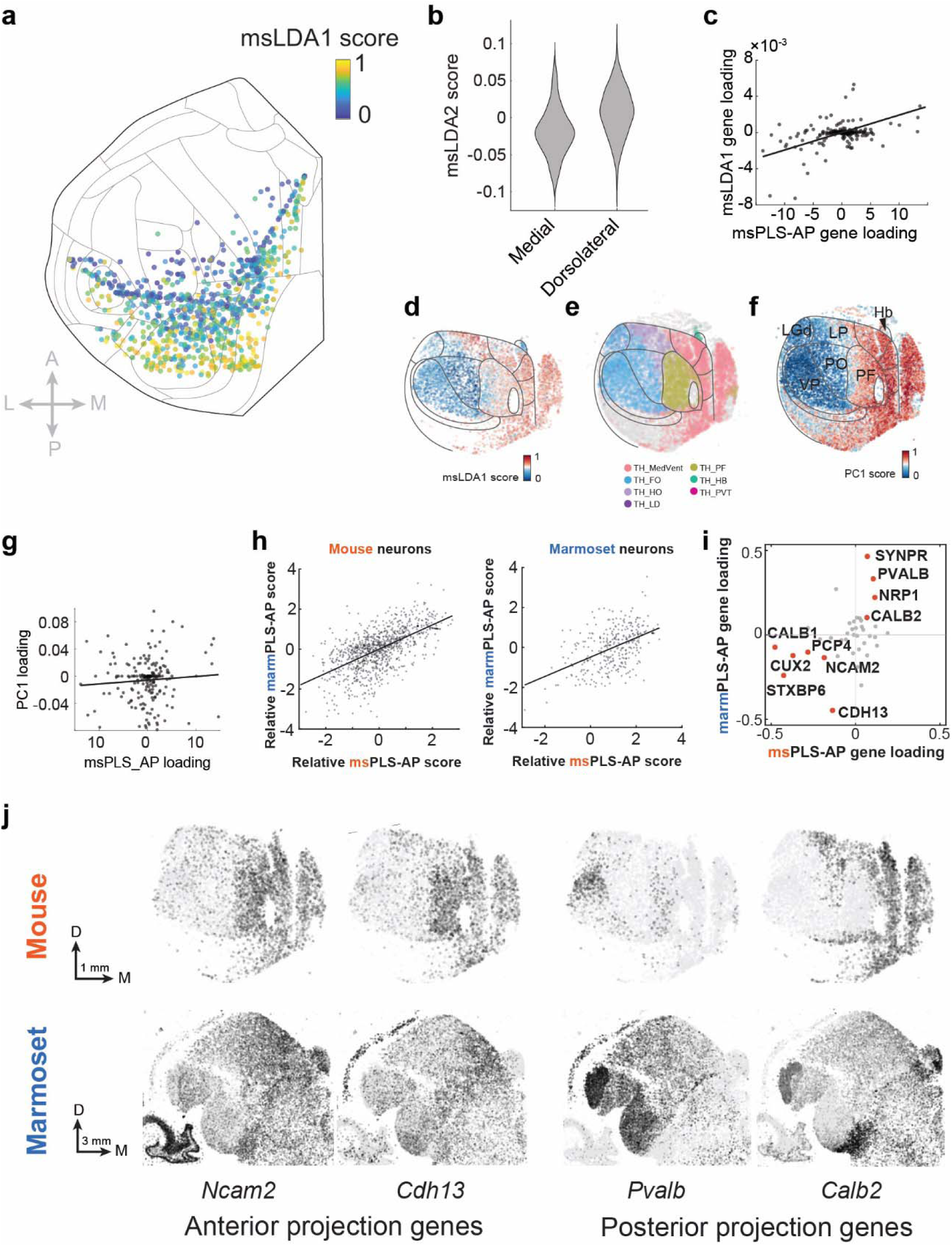
Gene expression of mouse thalamic neurons predicts projections. (**a**) Projection centroids of individual mouse neurons plotted on a cortical flatmap, colored by msLDA1 score. (**b**) Violin plots showing the distribution of msLDA2 scores in medial projections (ACA and PL/IL) and in dorsolateral projections (all other cortical projections). (**c**) Correspondence between mouse gene loadings on msLDA1 (y-axis) and msPLS-AP (x-axis). Each dot represents one gene. Black line indicates best linear fit, Pearson *r =* 0.50, *p =* 6×10^-14^. (**d**)(**e**)(**f)** A representative mouse thalamic section showing neurons colored by their msLDA1 score (d), coarse-level neuron types (e), and by PC1 score (f). (**g**) Gene loadings on PC1 (x-axis) and msPLS-AP (y-axis). Each dot represents one gene. Black line indicates best linear fit, Pearson *r =* 0.08, *p =* 0.3. (**h**) Z-scores of mouse thalamic excitatory neurons (*left*) and marmoset thalamic excitatory neurons (*right*) on both the shared-gene msPLS-AP and marmPLS-AP axes. (**i**) Loadings of genes that are shared between the msPLS-AP and marmPLS-AP axes. Genes with high absolute values are marked. (**j**) The expression of the indicated genes in the mouse (*top*) and the marmoset thalamus (*bottom*).

## Supplementary Files

***Supplementary Table 1: Anatomical area abbreviations***

***Supplementary Table 2: Gene panels and oligos***

***Supplementary Table 3: Examination of border effects on projection similarity in mouse***

***Supplementary Table 4: List of all dissected cubelets in mouse and marmoset***

***Supplementary Table 5: List of matched neurons to (Córdoba-Claros et al., 2025b)***

***Supplementary File 1: zip file containing dissection slice images for the BARseq experiments***

***Supplementary File 2: zip file containing flatmap representations of all matched neurons to (Córdoba-Claros et al., 2025b)***

## References

Ardesch, D.J., Scholtens, L.H., de Lange, S.C., Roumazeilles, L., Khrapitchev, A.A., Preuss, T.M., Rilling, J.K., Mars, R.B., van den Heuvel, M.P., 2022. Scaling Principles of White Matter Connectivity in the Human and Nonhuman Primate Brain. Cereb. Cortex 32, 2831–2842. 10.1093/cercor/bhab384

Bailey, P., Von Bonin, G., 1957. Evolution of the cerebral cortex: organ of the mind. Whats New 13–19.

Baldwin, M.K.L., Bourne, J.A., 2020. The Evolution of Subcortical Pathways to the Extrastriate Cortex, in: Evolutionary Neuroscience. Elsevier, pp. 565–587. 10.1016/B978-0-12-820584-6.00024-6

Baleydier, C., Morel, A., 1992. Segregated thalamocortical pathways to inferior parietal and inferotemporal cortex in macaque monkey. Vis. Neurosci. 8, 391–405. 10.1017/s0952523800004922

Bar-Joseph, Z., Gifford, D.K., Jaakkola, T.S., 2001. Fast optimal leaf ordering for hierarchical clustering. Bioinformatics 17 Suppl 1, S22–29. 10.1093/bioinformatics/17.suppl_1.s22

Bullmore, E., Sporns, O., 2012. The economy of brain network organization. Nat. Rev. Neurosci. 13, 336–349. 10.1038/nrn3214

Canton-Josh, J.E., Ackert-Smith, L.A., Costa, R.M., Pinto, L., 2025. Network dynamics underlying activity-timescale differences between cortical regions. 10.1101/2025.09.19.677442

Cappe, C., Morel, A., Barone, P., Rouiller, E.M., 2009. The thalamocortical projection systems in primate: an anatomical support for multisensory and sensorimotor interplay. Cereb. Cortex 19, 2025–2037. 10.1093/cercor/bhn228

Chaplin, T.A., Yu, H.-H., Soares, J.G.M., Gattass, R., Rosa, M.G.P., 2013. A Conserved Pattern of Differential Expansion of Cortical Areas in Simian Primates. J. Neurosci. 33, 15120–15125. 10.1523/JNEUROSCI.2909-13.2013

Chen, X., Fischer, S., Rue, M.C.P., Zhang, A., Mukherjee, D., Kanold, P.O., Gillis, J., Zador, A.M., 2025. Whole-cortex in situ sequencing reveals input-dependent area identity. Nature 647, 203–212. 10.1038/s41586-024-07221-6

Chen, X., Sun, Y.-C., Zhan, H., Kebschull, J.M., Fischer, S., Matho, K., Huang, Z.J., Gillis, J., Zador, A.M., 2019. High-Throughput Mapping of Long-Range Neuronal Projection Using In Situ Sequencing. Cell 179, 772–786.e19. 10.1016/j.cell.2019.09.023

Chen, Y., Chen, X., Baserdem, B., Zhan, H., Li, Y., Davis, M.B., Kebschull, J.M., Zador, A.M., Koulakov, A.A., Albeanu, D.F., 2022. High-throughput sequencing of single neuron projections reveals spatial organization in the olfactory cortex. Cell 185, 4117–4134.e28. 10.1016/j.cell.2022.09.038

Chklovskii, D.B., Schikorski, T., Stevens, C.F., 2002. Wiring Optimization in Cortical Circuits. Neuron 34, 341–347. 10.1016/S0896-6273(02)00679-7

Córdoba-Claros, M.A., Rubio-Garrido, P., de Lima, R.R.M., Morais, P.L.A.G., do Nascimento, E.S., Cavalcante, J.S., Clascá, F., 2025a. Projection Motifs and Wiring Logic of Medial Pulvinar Thalamocortical Axons in the Marmoset Monkey. J. Neurosci. 45, e1837242025. 10.1523/JNEUROSCI.1837-24.2025

Córdoba-Claros, M.A., Rubio-Garrido, P., De Lima, R.R.M., Morais, P.L.A.G., Do Nascimento, E.S., Cavalcante, J.S., Clascá, F., 2025b. Projection Motifs and Wiring Logic of Medial Pulvinar Thalamocortical Axons in the Marmoset Monkey. J. Neurosci. 45, e1837242025. 10.1523/JNEUROSCI.1837-24.2025

Covert, I., Gala, R., Wang, T., Svoboda, K., Sümbül, U., Lee, S.-I., 2023. Predictive and robust gene selection for spatial transcriptomics. Nat. Commun. 14, 2091. 10.1038/s41467-023-37392-1

Cox, R.W., 1996. AFNI: Software for Analysis and Visualization of Functional Magnetic Resonance Neuroimages. Comput. Biomed. Res. 29, 162–173. 10.1006/cbmr.1996.0014

Cusick, C.G., Scripter, J.L., Darensbourg, J.G., Weber, J.T., 1993. Chemoarchitectonic subdivisions of the visual pulvinar in monkeys and their connectional relations with the middle temporal and rostral dorsolateral visual areas, MT and DLr. J. Comp. Neurol. 336, 1–30. 10.1002/cne.903360102

Dan, S., Turner, M.A., DeBerardine, M., Caceres, L., Nieto-Caballero, V.E., Schembri, J., Levandowski, K., Cardenas, C., Wang, N., Li, C., Basu, S., Zheng, H., Liu, X., Höllt, T., Feliciano, J., Zhang, Q., Nakamoto, R., McMillen, D.A., Martin, N., Cuevas, N.V., Olsen, P., Nagra, J., Campos, J., VanNess, M.M., Waters, J., Ransford, S., Juneau, Z., Hastings, S., Barta, S., Ruiz, A., Ariza, J., Raphael, B.J., Zeng, H., Lein, E.S., Lelieveldt, B., Feng, G., Long, B., Krienen, F.M., 2025. Spatial patterning of transcriptional and regulatory programs in the primate subcortex. 10.1101/2025.11.22.689869

Dufour, A., Seibt, J., Passante, L., Depaepe, V., Ciossek, T., Frisén, J., Kullander, K., Flanagan, J.G., Polleux, F., Vanderhaeghen, P., 2003. Area specificity and topography of thalamocortical projections are controlled by ephrin/Eph genes. Neuron 39, 453–465. 10.1016/s0896-6273(03)00440-9

Fischer, S., Gillis, J., 2021. How many markers are needed to robustly determine a cell’s type? iScience 24, 103292. 10.1016/j.isci.2021.103292

Frangeul, L., Pouchelon, G., Telley, L., Lefort, S., Luscher, C., Jabaudon, D., 2016. A cross-modal genetic framework for the development and plasticity of sensory pathways. Nature 538, 96–98. 10.1038/nature19770

Fukuchi-Shimogori, T., Grove, E.A., 2003. Emx2 patterns the neocortex by regulating FGF positional signaling. Nat. Neurosci. 6, 825–831. 10.1038/nn1093

Giarrocco, F., Averbeck, B.B., 2023. Anatomical organization of forebrain circuits in the primate. Brain Struct. Funct. 228, 393–411. 10.1007/s00429-022-02586-8

Goldman-Rakic, P.S., Porrino, L.J., 1985. The primate mediodorsal (MD) nucleus and its projection to the frontal lobe. J. Comp. Neurol. 242, 535–560. 10.1002/cne.902420406

Gou, L., Wang, Y., Gao, L., Liu, S., Wang, M., Chai, Q., Fang, J., Zhan, L., Shen, X., Jiang, T., Ren, W., Ren, M., Jia, X., Xiao, C., Li, A., Li, X., Luo, Q., Okazawa, G., Yang, T., Liu, Z., Poo, M., Yang, X., Shen, Z., Xu, C., Yan, J., 2025. Single-neuron projectomes of macaque prefrontal cortex reveal refined axon targeting and arborization. Cell 188, 3806–3822.e24. 10.1016/j.cell.2025.06.005

Grant, E., Hoerder-Suabedissen, A., Molnár, Z., 2012. Development of the Corticothalamic Projections. Front. Neurosci. 6, 53. 10.3389/fnins.2012.00053

Harris, J.A., Mihalas, S., Hirokawa, K.E., Whitesell, J.D., Choi, H., Bernard, A., Bohn, P., Caldejon, S., Casal, L., Cho, A., Feiner, A., Feng, D., Gaudreault, N., Gerfen, C.R., Graddis, N., Groblewski, P.A., Henry, A.M., Ho, A., Howard, R., Knox, J.E., Kuan, L., Kuang, X., Lecoq, J., Lesnar, P., Li, Y., Luviano, J., McConoughey, S., Mortrud, M.T., Naeemi, M., Ng, L., Oh, S.W., Ouellette, B., Shen, E., Sorensen, S.A., Wakeman, W., Wang, Q., Wang, Y., Williford, A., Phillips, J.W., Jones, A.R., Koch, C., Zeng, H., 2019. Hierarchical organization of cortical and thalamic connectivity. Nature 575, 195–202. 10.1038/s41586-019-1716-z

Huang, Z., Yang, Q., Li, S., Zhu, X., Wang, H., Lin, J., Zhan, Y., Wu, Y., Wang, Z., Majka, P., Qu, H., Atapour, N., Yang, T., Lin, Y., Cui, L., Yao, Y.-G., Liang, Z., Liu, Z., Li, Chao, Wei, W., Zhou, Y., Ma, S., Shen, Z., Wei, X., Xu, X., Liu, S., Li, Chengyu, Poo, M., Liu, L., Rosa, M.G.P., Sun, Y., Hao, S., Liu, C., 2026. An opposing molecular gradient axis underlies primate cortical organization. Science 392, eaea2673. 10.1126/science.aea2673

Jaramillo, J., Mejias, J.F., Wang, X.-J., 2019. Engagement of Pulvino-cortical Feedforward and Feedback Pathways in Cognitive Computations. Neuron 101, 321–336.e9. 10.1016/j.neuron.2018.11.023

Jiang, S., Wang, L., Yun, Z., Chen, H., Liu, L., Yao, J., Peng, H., 2025. NeuroXiv: AI-powered open databasing and dynamic mining of brain-wide neuron morphometry. Nat. Methods 22, 1195–1198. 10.1038/s41592-025-02687-2

Jones, E.G., 2007. Thalamus, Second edition. ed. Cambridge University Press, Cambridge (GB).

Jones, E.G., 2001. The thalamic matrix and thalamocortical synchrony. Trends Neurosci. 24, 595–601. 10.1016/s0166-2236(00)01922-6

Jones, E.G., 1998. Viewpoint: the core and matrix of thalamic organization. Neuroscience 85, 331–345. 10.1016/s0306-4522(97)00581-2

Kebschull, J.M., Garcia da Silva, P., Reid, A.P., Peikon, I.D., Albeanu, D.F., Zador, A.M., 2016. High-Throughput Mapping of Single-Neuron Projections by Sequencing of Barcoded RNA. Neuron 91, 975–987. 10.1016/j.neuron.2016.07.036

Kita, Y., Nishibe, H., Wang, Y., Hashikawa, T., Kikuchi, S.S., U, M., Yoshida, A.C., Yoshida, C., Kawase, T., Ishii, S., Skibbe, H., Shimogori, T., 2021. Cellular-resolution gene expression profiling in the neonatal marmoset brain reveals dynamic species- and region-specific differences. Proc. Natl. Acad. Sci. 118, e2020125118. 10.1073/pnas.2020125118

Krienen, F.M., Levandowski, K.M., Zaniewski, H., Del Rosario, R.C.H., Schroeder, M.E., Goldman, M., Wienisch, M., Lutservitz, A., Beja-Glasser, V.F., Chen, C., Zhang, Q., Chan, K.Y., Li, K.X., Sharma, J., McCormack, D., Shin, T.W., Harrahill, A., Nyase, E., Mudhar, G., Mauermann, A., Wysoker, A., Nemesh, J., Kashin, S., Vergara, J., Chelini, G., Dimidschstein, J., Berretta, S., Deverman, B.E., Boyden, E., McCarroll, S.A., Feng, G., 2023. A marmoset brain cell census reveals regional specialization of cellular identities. Sci. Adv. 9, eadk3986. 10.1126/sciadv.adk3986

Kudo, L.C., Karsten, S.L., Chen, J., Levitt, P., Geschwind, D.H., 2007. Genetic Analysis of Anterior-Posterior Expression Gradients in the Developing Mammalian Forebrain. Cereb. Cortex 17, 2108–2122. 10.1093/cercor/bhl118

Lee, D.D., Seung, H.S., 1999. Learning the parts of objects by non-negative matrix factorization. Nature 401, 788–791. 10.1038/44565

Li, G., Li, S., Wang, X.-J., 2025. A hierarchy of time constants and reliable signal propagation in the marmoset cerebral cortex. Nat. Commun. 16, 11640. 10.1038/s41467-025-66699-4

Liu, C., Yen, C.C.-C., Szczupak, D., Tian, X., Glen, D., Silva, A.C., 2021. Marmoset Brain Mapping V3: Population multi-modal standard volumetric and surface-based templates. NeuroImage 226, 117620. 10.1016/j.neuroimage.2020.117620

Magrou, L., Joyce, M.K.P., Froudist-Walsh, S., Datta, D., Wang, X.-J., Martinez-Trujillo, J., Arnsten, A.F.T., 2024. The meso-connectomes of mouse, marmoset, and macaque: network organization and the emergence of higher cognition. Cereb. Cortex 34, bhae174. 10.1093/cercor/bhae174

Mota, B., Dos Santos, S.E., Ventura-Antunes, L., Jardim-Messeder, D., Neves, K., Kazu, R.S., Noctor, S., Lambert, K., Bertelsen, M.F., Manger, P.R., Sherwood, C.C., Kaas, J.H., Herculano-Houzel, S., 2019. White matter volume and white/gray matter ratio in mammalian species as a consequence of the universal scaling of cortical folding. Proc. Natl. Acad. Sci. U. S. A. 116, 15253–15261. 10.1073/pnas.1716956116

Mota, B., Herculano-Houzel, S., 2015. BRAIN STRUCTURE. Cortical folding scales universally with surface area and thickness, not number of neurons. Science 349, 74–77. 10.1126/science.aaa9101

Müller, E.J., Munn, B., Hearne, L.J., Smith, J.B., Fulcher, B., Arnatkevičiūtė, A., Lurie, D.J., Cocchi, L., Shine, J.M., 2020. Core and matrix thalamic sub-populations relate to spatio-temporal cortical connectivity gradients. NeuroImage 222, 117224. 10.1016/j.neuroimage.2020.117224

Mundinano, I.-C., Kwan, W.C., Bourne, J.A., 2019. Retinotopic specializations of cortical and thalamic inputs to area MT. Proc. Natl. Acad. Sci. U. S. A. 116, 23326–23331. 10.1073/pnas.1909799116

Muñoz-Castañeda, R., Zingg, B., Matho, K.S., Chen, X., Wang, Q., Foster, N.N., Li, A., Narasimhan, A., Hirokawa, K.E., Huo, B., Bannerjee, S., Korobkova, L., Park, C.S., Park, Y.-G., Bienkowski, M.S., Chon, U., Wheeler, D.W., Li, Xiangning, Wang, Yun, Naeemi, M., Xie, P., Liu, L., Kelly, K., An, X., Attili, S.M., Bowman, I., Bludova, A., Cetin, A., Ding, L., Drewes, R., D’Orazi, F., Elowsky, C., Fischer, S., Galbavy, W., Gao, L., Gillis, J., Groblewski, P.A., Gou, L., Hahn, J.D., Hatfield, J.T., Hintiryan, H., Huang, J.J., Kondo, H., Kuang, X., Lesnar, P., Li, Xu, Li, Y., Lin, M., Lo, D., Mizrachi, J., Mok, S., Nicovich, P.R., Palaniswamy, R., Palmer, J., Qi, X., Shen, E., Sun, Y.-C., Tao, H.W., Wakemen, W., Wang, Yimin, Yao, S., Yuan, J., Zhan, H., Zhu, M., Ng, L., Zhang, L.I., Lim, B.K., Hawrylycz, M., Gong, H., Gee, J.C., Kim, Y., Chung, K., Yang, X.W., Peng, H., Luo, Q., Mitra, P.P., Zador, A.M., Zeng, H., Ascoli, G.A., Josh Huang, Z., Osten, P., Harris, J.A., Dong, H.-W., 2021. Cellular anatomy of the mouse primary motor cortex. Nature 598, 159–166. 10.1038/s41586-021-03970-w

Murray, J.D., Bernacchia, A., Freedman, D.J., Romo, R., Wallis, J.D., Cai, X., Padoa-Schioppa, C., Pasternak, T., Seo, H., Lee, D., Wang, X.-J., 2014. A hierarchy of intrinsic timescales across primate cortex. Nat. Neurosci. 17, 1661–1663. 10.1038/nn.3862

Oldham, S., Ball, G., 2023. A phylogenetically-conserved axis of thalamocortical connectivity in the human brain. Nat. Commun. 14, 6032. 10.1038/s41467-023-41722-8

Paxinos, G., 2012. The marmoset brain in stereotaxic coordinates, 1st ed. ed. Academic Press, London.

Paxinos, G., Rosa, M., Merlin, S., Majka, P., 2026. The marmoset brain in stereotaxic coordinates, Second edition. ed. Academic Press, Place of publication not identified.

Peng, H., Xie, P., Liu, L., Kuang, X., Wang, Yimin, Qu, L., Gong, H., Jiang, S., Li, A., Ruan, Z., Ding, L., Yao, Z., Chen, C., Chen, M., Daigle, T.L., Dalley, R., Ding, Z., Duan, Y., Feiner, A., He, P., Hill, C., Hirokawa, K.E., Hong, G., Huang, L., Kebede, S., Kuo, H.-C., Larsen, R., Lesnar, P., Li, L., Li, Q., Li, X., Li, Yaoyao, Li, Yuanyuan, Liu, A., Lu, D., Mok, S., Ng, L., Nguyen, T.N., Ouyang, Q., Pan, J., Shen, E., Song, Y., Sunkin, S.M., Tasic, B., Veldman, M.B., Wakeman, W., Wan, W., Wang, P., Wang, Q., Wang, T., Wang, Yaping, Xiong, F., Xiong, W., Xu, W., Ye, M., Yin, L., Yu, Y., Yuan, Jia, Yuan, Jing, Yun, Z., Zeng, S., Zhang, S., Zhao, S., Zhao, Z., Zhou, Z., Huang, Z.J., Esposito, L., Hawrylycz, M.J., Sorensen, S.A., Yang, X.W., Zheng, Y., Gu, Z., Xie, W., Koch, C., Luo, Q., Harris, J.A., Wang, Yun, Zeng, H., 2021. Morphological diversity of single neurons in molecularly defined cell types. Nature 598, 174–181. 10.1038/s41586-021-03941-1

Petty, G.H., Bruno, R.M., 2024. Attentional modulation of secondary somatosensory and visual thalamus of mice. eLife 13, RP97188. 10.7554/eLife.97188

Phillips, J.M., Fish, L.R., Kambi, N.A., Redinbaugh, M.J., Mohanta, S., Kecskemeti, S.R., Saalmann, Y.B., 2019. Topographic organization of connections between prefrontal cortex and mediodorsal thalamus: Evidence for a general principle of indirect thalamic pathways between directly connected cortical areas. NeuroImage 189, 832–846. 10.1016/j.neuroimage.2019.01.078

Phillips, J.W., Schulmann, A., Hara, E., Winnubst, J., Liu, C., Valakh, V., Wang, L., Shields, B.C., Korff, W., Chandrashekar, J., Lemire, A.L., Mensh, B., Dudman, J.T., Nelson, S.B., Hantman, A.W., 2019. A repeated molecular architecture across thalamic pathways. Nat. Neurosci. 22, 1925–1935. 10.1038/s41593-019-0483-3

Puche-Aroca, L., Andrés-Bayón, B., Wilson, E.S., Javed, A., Torregrosa-Mira, A., López-Atalaya, J.P., López-Bendito, G., 2025. Early lineage divergence segregates sensory and non-sensory thalamic circuits. 10.1101/2025.07.17.665342

Ringo, J.L., 1991. Neuronal interconnection as a function of brain size. Brain. Behav. Evol. 38, 1–6. 10.1159/000114375

Ringo, J.L., Doty, R.W., Demeter, S., Simard, P.Y., 1994. Time is of the essence: a conjecture that hemispheric specialization arises from interhemispheric conduction delay. Cereb. Cortex 4, 331–343. 10.1093/cercor/4.4.331

Roberts, A.C., Tomic, D.L., Parkinson, C.H., Roeling, T.A., Cutter, D.J., Robbins, T.W., Everitt, B.J., 2007. Forebrain connectivity of the prefrontal cortex in the marmoset monkey (Callithrix jacchus): an anterograde and retrograde tract-tracing study. J. Comp. Neurol. 502, 86–112. 10.1002/cne.21300

Romanski, L.M., Giguere, M., Bates, J.F., Goldman-Rakic, P.S., 1997. Topographic organization of medial pulvinar connections with the prefrontal cortex in the rhesus monkey. J. Comp. Neurol. 379, 313–332.

Saalmann, Y.B., Pinsk, M.A., Wang, L., Li, X., Kastner, S., 2012. The pulvinar regulates information transmission between cortical areas based on attention demands. Science 337, 753–756. 10.1126/science.1223082

Schulmann, A., Feng, N., Auluck, P.K., Mukherjee, A., Komal, R., Leng, Y., Gao, C., Williams Avram, S.K., Roy, S., Usdin, T.B., Xu, Q., Imamovic, V., Patel, Y., Akula, N., Raznahan, A., Menon, V., Roussos, P., Duncan, L., Elkahloun, A., Singh, J., Kelly, M.C., Halassa, M.M., Hattar, S., Penzo, M.A., Marenco, S., McMahon, F.J., 2024. A conserved cell-type gradient across the human mediodorsal and paraventricular thalamus. 10.1101/2024.09.03.611112

Scott, J.T., Mendivez Vasquez, B.L., Stewart, B.J., Panacheril, D.D., Rajit, D.K.J., Fan, A.Y., Bourne, J.A., 2025. CalliCog is an open-source cognitive neuroscience toolkit for freely behaving nonhuman primates. Cell Rep. Methods 5, 101034. 10.1016/j.crmeth.2025.101034

Shepherd, G.M.G., Yamawaki, N., 2021. Untangling the cortico-thalamo-cortical loop: cellular pieces of a knotty circuit puzzle. Nat. Rev. Neurosci. 22, 389–406. 10.1038/s41583-021-00459-3

Sherman, S.M., 2016. Thalamus plays a central role in ongoing cortical functioning. Nat. Neurosci. 19, 533–541. 10.1038/nn.4269

Sherman, S.M., 2007. The thalamus is more than just a relay. Curr. Opin. Neurobiol. 17, 417–422. 10.1016/j.conb.2007.07.003

Sherman, S.M., Guillery, R.W., 2002. The role of the thalamus in the flow of information to the cortex. Philos. Trans. R. Soc. Lond. B. Biol. Sci. 357, 1695–1708. 10.1098/rstb.2002.1161

Sherman, S.M., Guillery, R.W., Sherman, S.M., 2006. Exploring the thalamus and its role in cortical function, 2nd ed. ed. MIT Press, Cambridge, Mass.

Shimogori, T., Abe, A., Go, Y., Hashikawa, T., Kishi, N., Kikuchi, S.S., Kita, Y., Niimi, K., Nishibe, H., Okuno, M., Saga, K., Sakurai, M., Sato, M., Serizawa, T., Suzuki, S., Takahashi, E., Tanaka, M., Tatsumoto, S., Toki, M., U, M., Wang, Y., Windak, K.J., Yamagishi, H., Yamashita, K., Yoda, T., Yoshida, A.C., Yoshida, C., Yoshimoto, T., Okano, H., 2018. Digital gene atlas of neonate common marmoset brain. Neurosci. Res. 128, 1–13. 10.1016/j.neures.2017.10.009

Shipp, S., 2003. The functional logic of cortico–pulvinar connections. Philos. Trans. R. Soc. Lond. B. Biol. Sci. 358, 1605–1624. 10.1098/rstb.2002.1213

Shipp, S., 2001. Corticopulvinar connections of areas V5, V4, and V3 in the macaque monkey: a dual model of retinal and cortical topographies. J. Comp. Neurol. 439, 469–490. 10.1002/cne.1363

Sun, Y.-C., Chen, X., Fischer, S., Lu, S., Zhan, H., Gillis, J., Zador, A.M., 2021. Integrating barcoded neuroanatomy with spatial transcriptional profiling enables identification of gene correlates of projections. Nat. Neurosci. 24, 873–885. 10.1038/s41593-021-00842-4

Sydnor, V.J., Bagautdinova, J., Larsen, B., Arcaro, M.J., Barch, D.M., Bassett, D.S., Alexander-Bloch, A.F., Cook, P.A., Covitz, S., Franco, A.R., Gur, R.E., Gur, R.C., Mackey, A.P., Mehta, K., Meisler, S.L., Milham, M.P., Moore, T.M., Müller, E.J., Roalf, D.R., Salo, T., Schubiner, G., Seidlitz, J., Shinohara, R.T., Shine, J.M., Yeh, F.-C., Cieslak, M., Satterthwaite, T.D., 2024. A sensorimotor-association axis of thalamocortical connection development. 10.1101/2024.06.13.598749

Tallinen, T., Chung, J.Y., Biggins, J.S., Mahadevan, L., 2014. Gyrification from constrained cortical expansion. Proc. Natl. Acad. Sci. U. S. A. 111, 12667–12672. 10.1073/pnas.1406015111

Van Essen, D.C., 1997. A tension-based theory of morphogenesis and compact wiring in the central nervous system. Nature 385, 313–318. 10.1038/385313a0

Vittek, A.-L., Juan, C., Nowak, L.G., Girard, P., Cappe, C., 2023. Multisensory integration in neurons of the medial pulvinar of macaque monkey. Cereb. Cortex 33, 4202–4215. 10.1093/cercor/bhac337

Wang, J., Isogai, Y., 2025. BARseq 2.5 v1. 10.17504/protocols.io.kqdg3ke9qv25/v1

Wang, Q., Ding, S.-L., Li, Y., Royall, J., Feng, D., Lesnar, P., Graddis, N., Naeemi, M., Facer, B., Ho, A., Dolbeare, T., Blanchard, B., Dee, N., Wakeman, W., Hirokawa, K.E., Szafer, A., Sunkin, S.M., Oh, S.W., Bernard, A., Phillips, J.W., Hawrylycz, M., Koch, C., Zeng, H., Harris, J.A., Ng, L., 2020. The Allen Mouse Brain Common Coordinate Framework: A 3D Reference Atlas. Cell 181, 936–953.e20. 10.1016/j.cell.2020.04.007

Winnubst, J., Bas, E., Ferreira, T.A., Wu, Z., Economo, M.N., Edson, P., Arthur, B.J., Bruns, C., Rokicki, K., Schauder, D., Olbris, D.J., Murphy, S.D., Ackerman, D.G., Arshadi, C., Baldwin, P., Blake, R., Elsayed, A., Hasan, M., Ramirez, D., Dos Santos, B., Weldon, M., Zafar, A., Dudman, J.T., Gerfen, C.R., Hantman, A.W., Korff, W., Sternson, S.M., Spruston, N., Svoboda, K., Chandrashekar, J., 2019. Reconstruction of 1,000 Projection Neurons Reveals New Cell Types and Organization of Long-Range Connectivity in the Mouse Brain. Cell 179, 268–281.e13. 10.1016/j.cell.2019.07.042

Yao, S., Wang, Q., Hirokawa, K.E., Ouellette, B., Ahmed, R., Bomben, J., Brouner, K., Casal, L., Caldejon, S., Cho, A., Dotson, N.I., Daigle, T.L., Egdorf, T., Enstrom, R., Gary, A., Gelfand, E., Gorham, M., Griffin, F., Gu, H., Hancock, N., Howard, R., Kuan, L., Lambert, S., Lee, E.K., Luviano, J., Mace, K., Maxwell, M., Mortrud, M.T., Naeemi, M., Nayan, C., Ngo, N.-K., Nguyen, T., North, K., Ransford, S., Ruiz, A., Seid, S., Swapp, J., Taormina, M.J., Wakeman, W., Zhou, T., Nicovich, P.R., Williford, A., Potekhina, L., McGraw, M., Ng, L., Groblewski, P.A., Tasic, B., Mihalas, S., Harris, J.A., Cetin, A., Zeng, H., 2023. A whole-brain monosynaptic input connectome to neuron classes in mouse visual cortex. Nat. Neurosci. 26, 350–364. 10.1038/s41593-022-01219-x

Yao, Z., Van Velthoven, C.T.J., Kunst, M., Zhang, M., McMillen, D., Lee, C., Jung, W., Goldy, J., Abdelhak, A., Aitken, M., Baker, K., Baker, P., Barkan, E., Bertagnolli, D., Bhandiwad, A., Bielstein, C., Bishwakarma, P., Campos, J., Carey, D., Casper, T., Chakka, A.B., Chakrabarty, R., Chavan, S., Chen, M., Clark, M., Close, J., Crichton, K., Daniel, S., DiValentin, P., Dolbeare, T., Ellingwood, L., Fiabane, E., Fliss, T., Gee, J., Gerstenberger, J., Glandon, A., Gloe, J., Gould, J., Gray, J., Guilford, N., Guzman, J., Hirschstein, D., Ho, W., Hooper, M., Huang, M., Hupp, M., Jin, K., Kroll, M., Lathia, K., Leon, A., Li, S., Long, B., Madigan, Z., Malloy, J., Malone, J., Maltzer, Z., Martin, N., McCue, R., McGinty, R., Mei, N., Melchor, J., Meyerdierks, E., Mollenkopf, T., Moonsman, S., Nguyen, T.N., Otto, S., Pham, T., Rimorin, C., Ruiz, A., Sanchez, R., Sawyer, L., Shapovalova, N., Shepard, N., Slaughterbeck, C., Sulc, J., Tieu, M., Torkelson, A., Tung, H., Valera Cuevas, N., Vance, S., Wadhwani, K., Ward, K., Levi, B., Farrell, C., Young, R., Staats, B., Wang, M.-Q.M., Thompson, C.L., Mufti, S., Pagan, C.M., Kruse, L., Dee, N., Sunkin, S.M., Esposito, L., Hawrylycz, M.J., Waters, J., Ng, L., Smith, K., Tasic, B., Zhuang, X., Zeng, H., 2023. A high-resolution transcriptomic and spatial atlas of cell types in the whole mouse brain. Nature 624, 317–332. 10.1038/s41586-023-06812-z

Zeisler, Z.R., Heslin, K.A., Stoll, F.M., Hof, P.R., Clem, R.L., Rudebeck, P.H., 2024. Comparative basolateral amygdala connectomics reveals dissociable single-neuron projection patterns to frontal cortex in macaques and mice. Curr. Biol. CB 34, 3249–3257.e3. 10.1016/j.cub.2024.06.012

Zhang, K., Sejnowski, T.J., 2000. A universal scaling law between gray matter and white matter of cerebral cortex. Proc. Natl. Acad. Sci. U. S. A. 97, 5621–5626. 10.1073/pnas.090504197

Zhang, M., Pan, X., Jung, W., Halpern, A.R., Eichhorn, S.W., Lei, Z., Cohen, L., Smith, K.A., Tasic, B., Yao, Z., Zeng, H., Zhuang, X., 2023. Molecularly defined and spatially resolved cell atlas of the whole mouse brain. Nature 624, 343–354. 10.1038/s41586-023-06808-9

