## Supplementary material for "Conserved and specialized features of thalamocortical wiring revealed by single-cell projection mapping in mouse and marmoset": Supp. File 1: Alto_mapseq_dissection_annotations.pdf

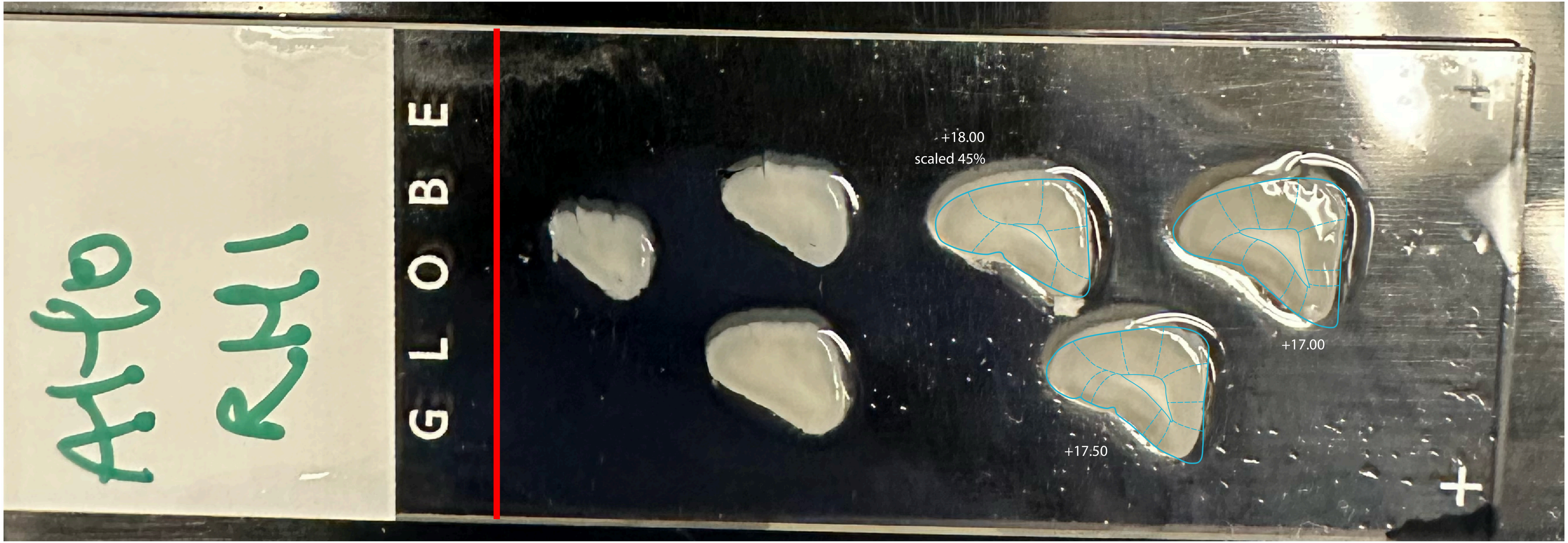


Alto_RH_1.jpg

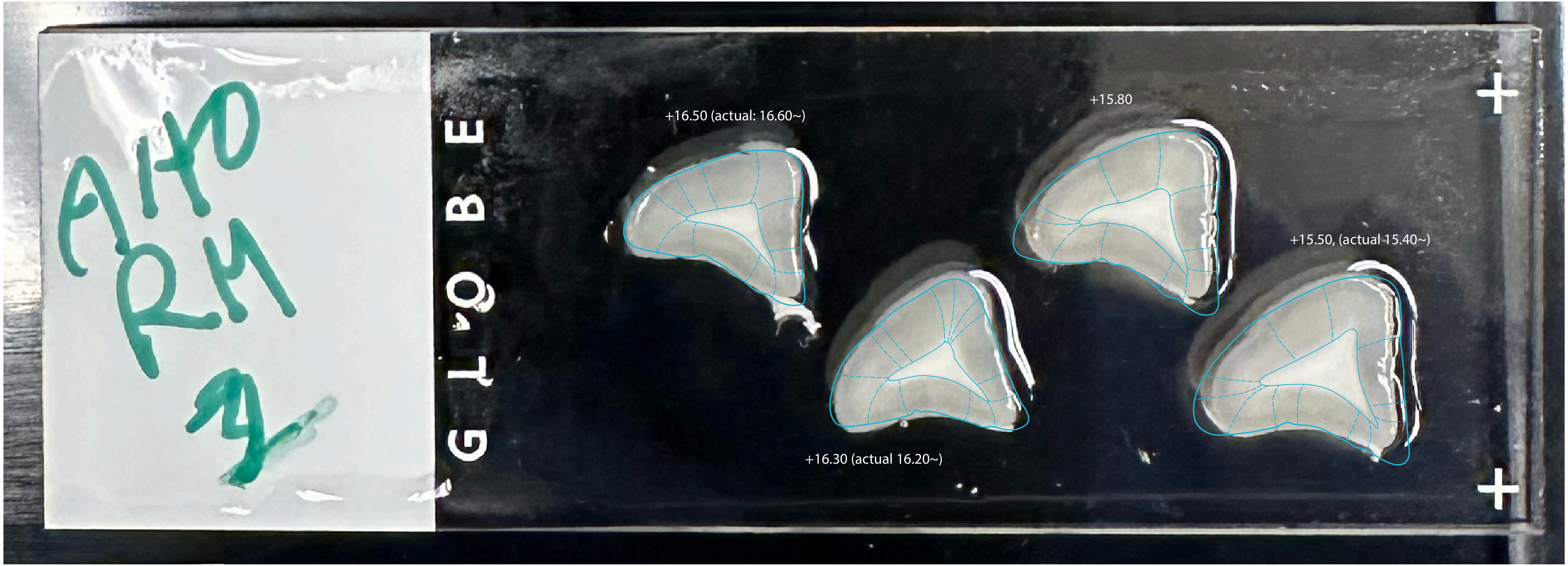
Alto
RH
GLOBE
+16.50 (actual: 16.60~)
+15.80
+15.50, (actual 15.40~)
+16.30 (actual 16.20~)

Alto_RH_2.jpg

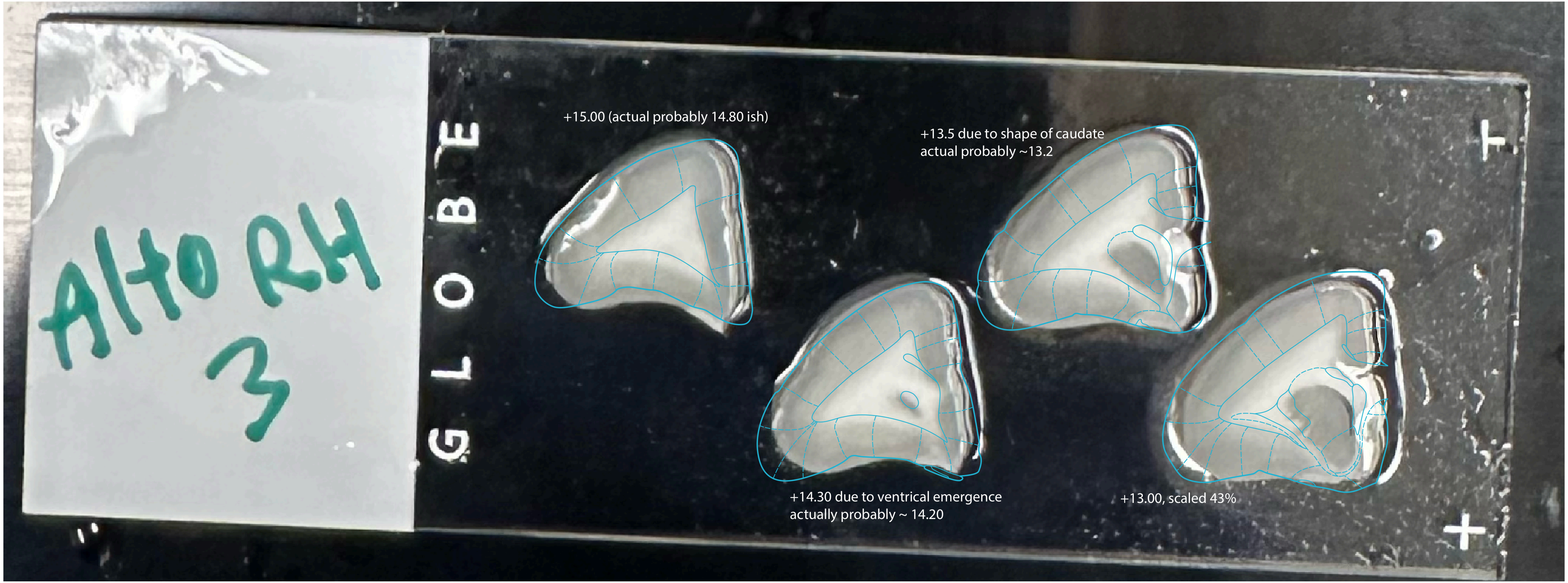
Alto RH
3
GLOBE
+15.00 (actual probably 14.80 ish)
+13.5 due to shape of caudate
actual probably ~13.2
+14.30 due to ventrical emergence
actually probably ~ 14.20
+13.00, scaled 43%

Alto_RH_3.jpg

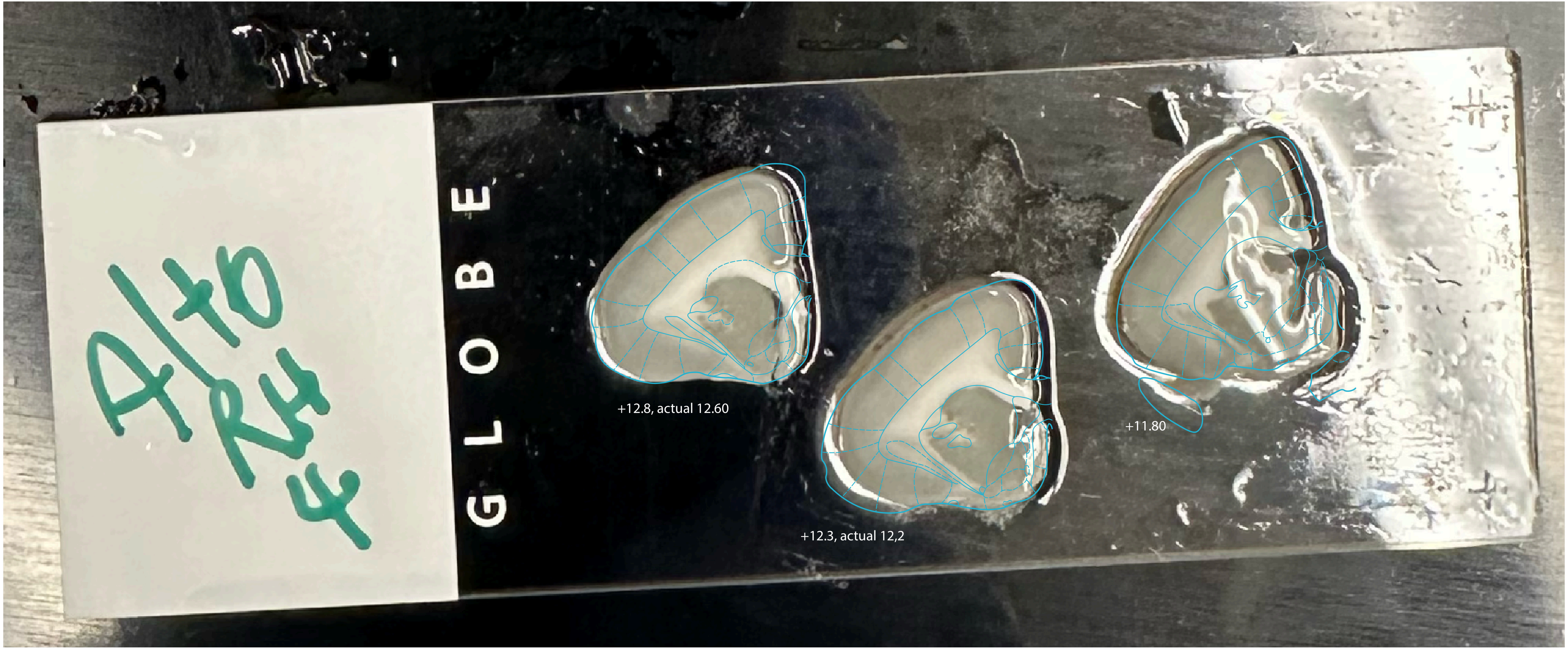


Alto_RH_4.jpg

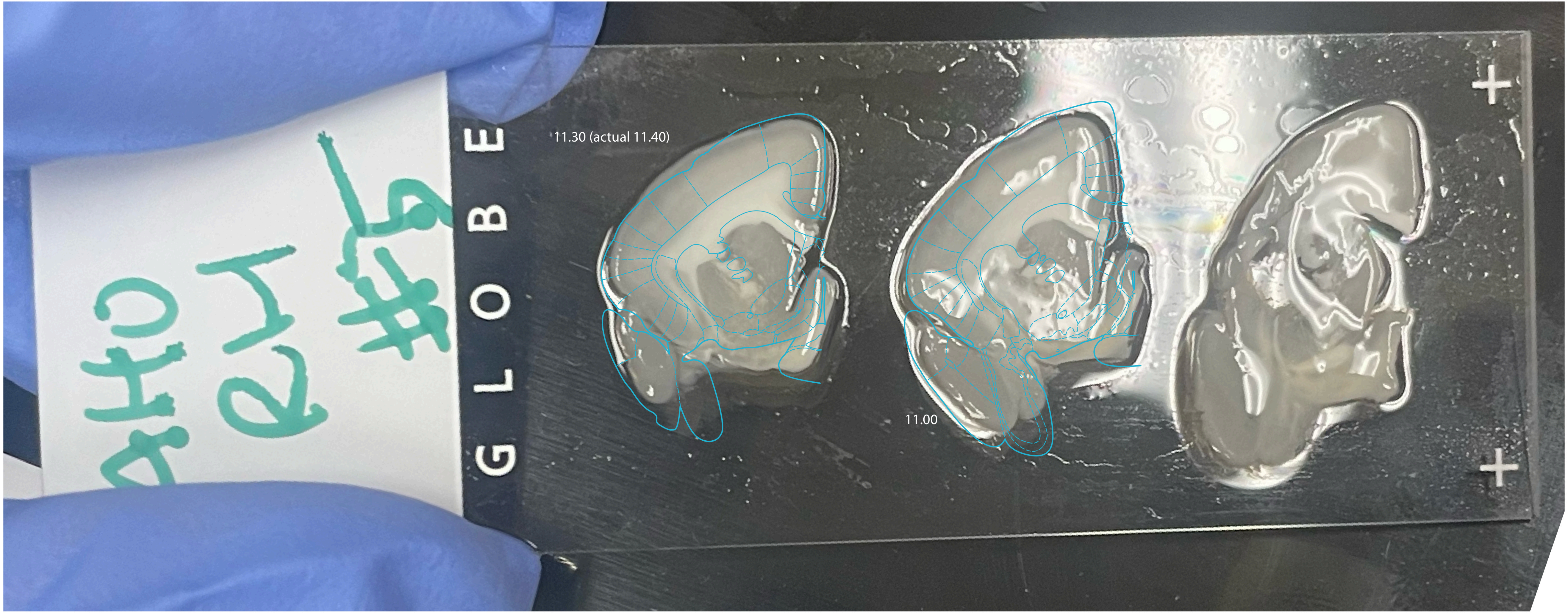


Alto_RH_5.jpg

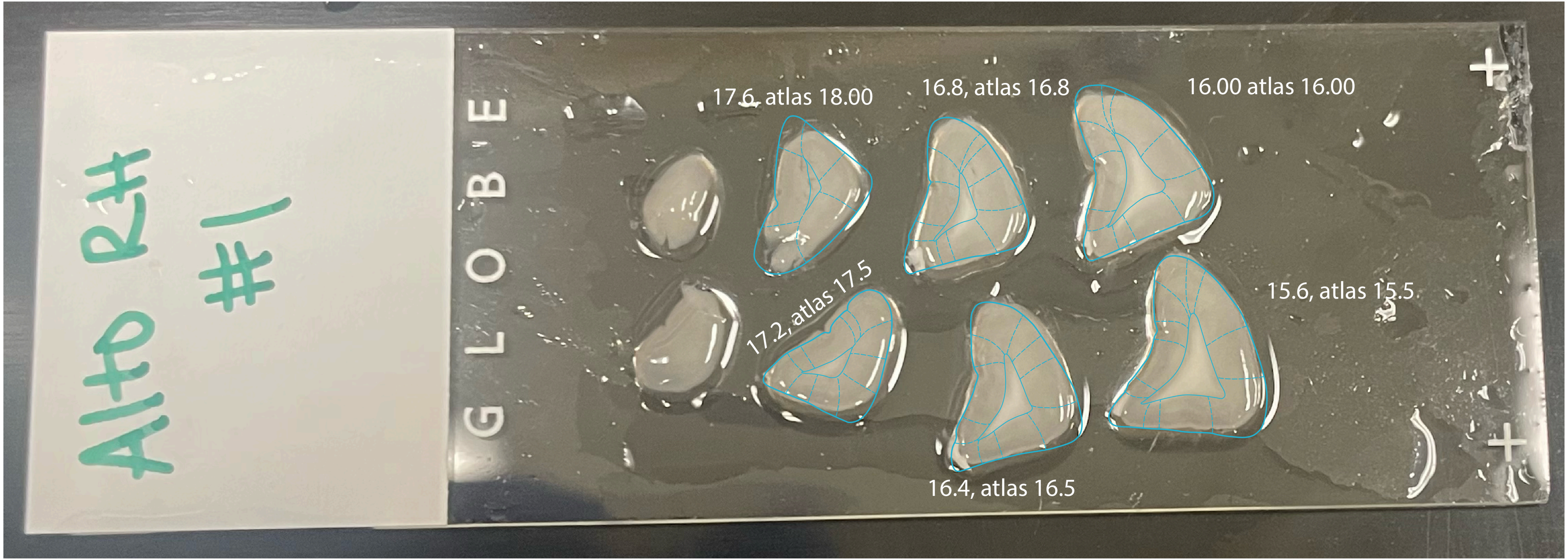
Alto RH #1
GLOBE
17.6, atlas 18.00
16.8, atlas 16.8
16.00 atlas 16.00
17.2, atlas 17.5
15.6, atlas 15.5
16.4, atlas 16.5

Alto_LH_1.jpg

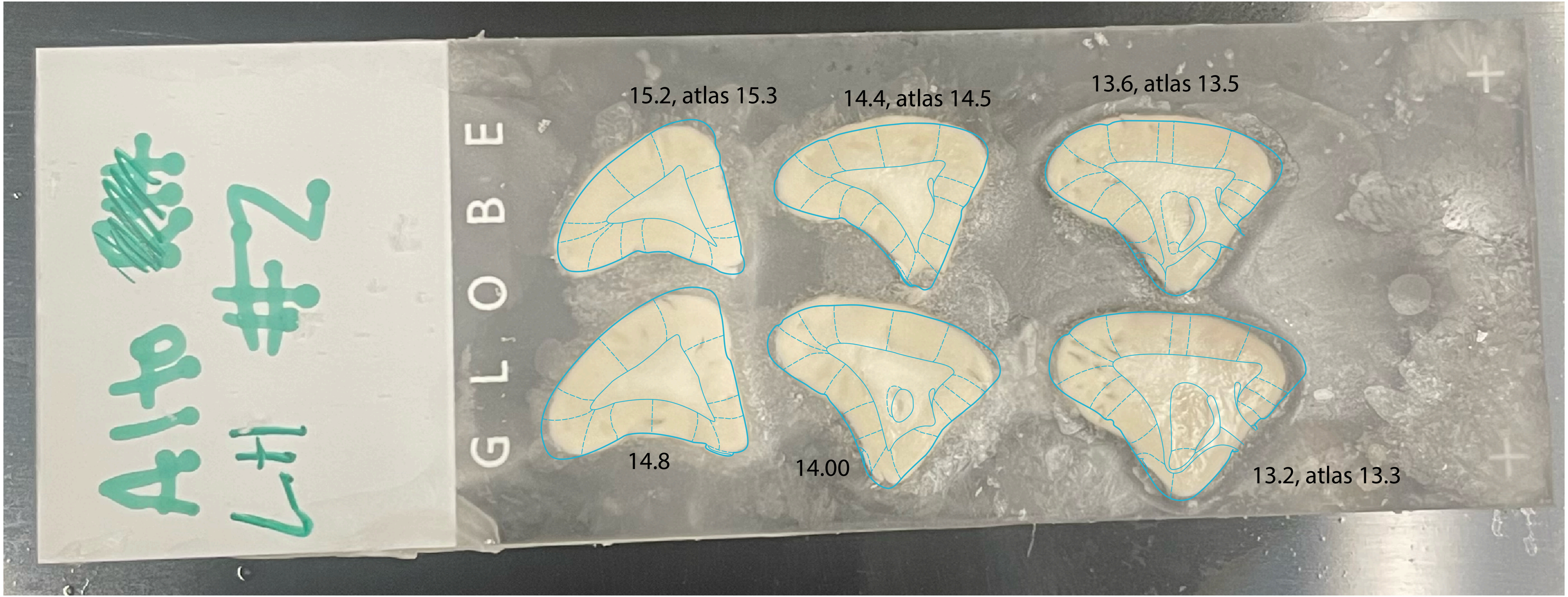
Alto #2
LH
GLOBE
15.2, atlas 15.3
14.4, atlas 14.5
13.6, atlas 13.5
14.8
14.00
13.2, atlas 13.3

Alto_LH_2.jpg

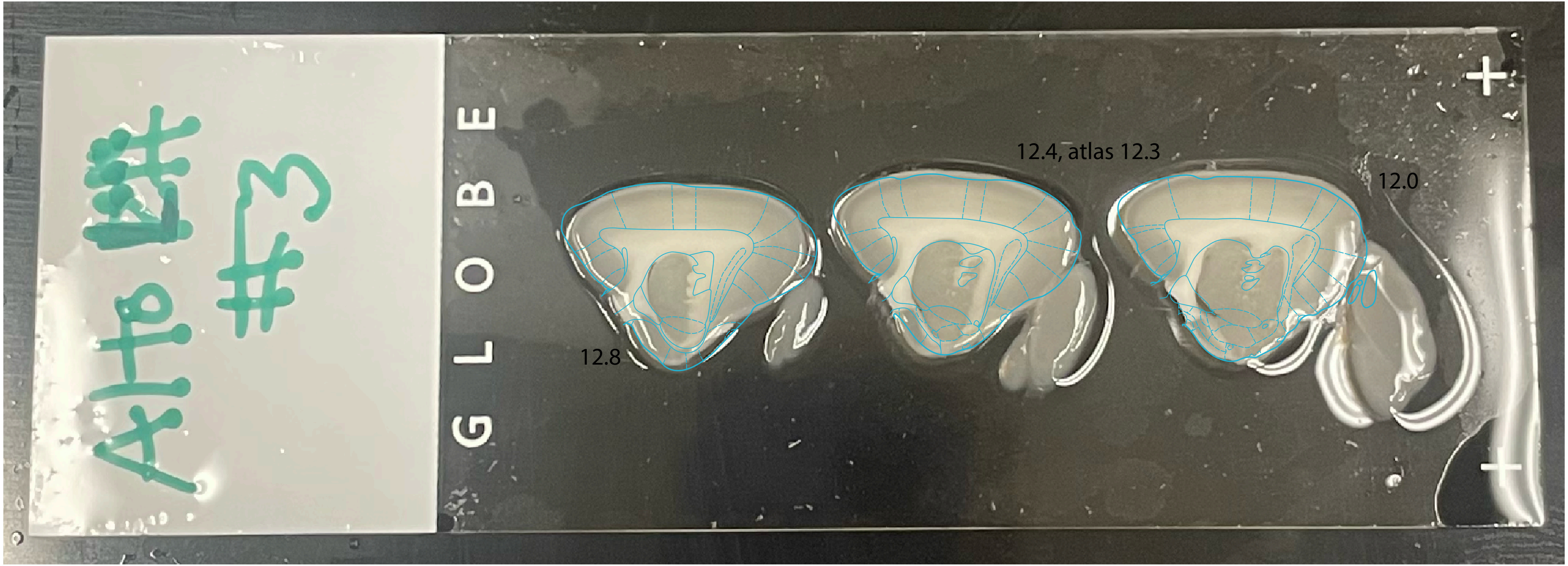


Alto_LH_3.jpg

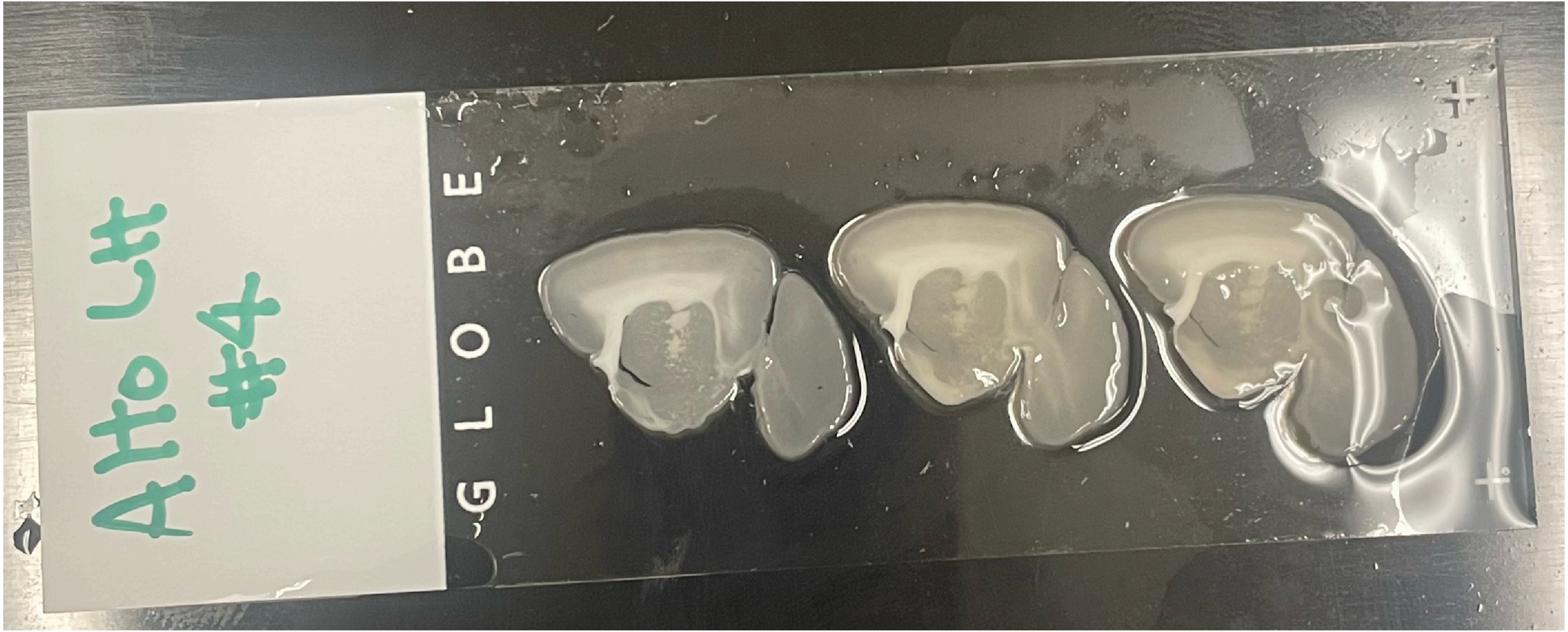


Alto_LH_4.jpg

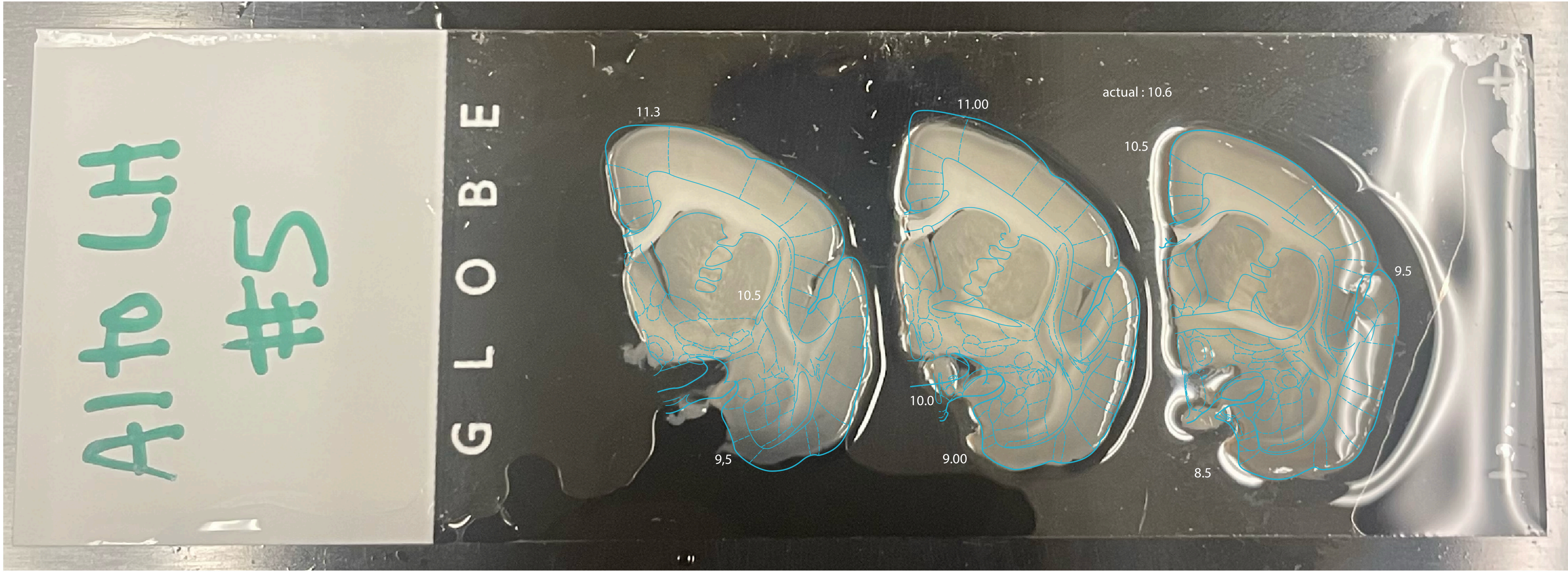


Alto_LH_5.jpg

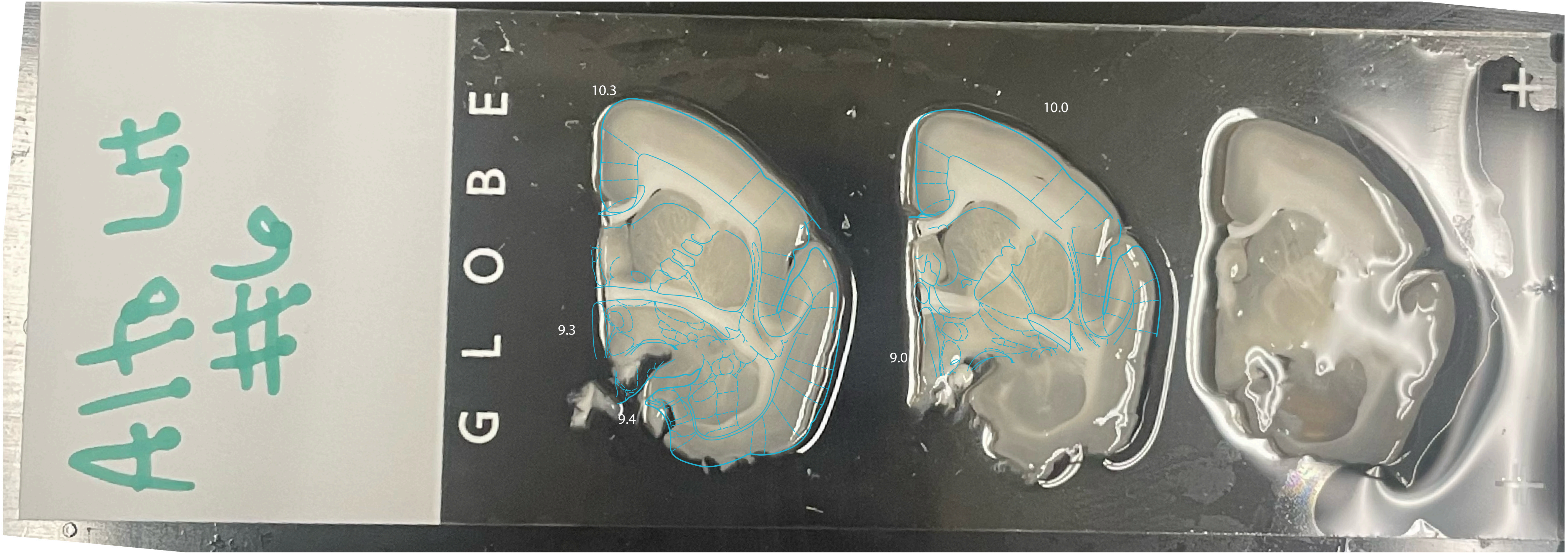


Alto_LH_6.jpg
