## Supplementary material for "Conserved and specialized features of thalamocortical wiring revealed by single-cell projection mapping in mouse and marmoset": Supp. File 1: Hazelnut_dissection_plan_annotated.pdf

### HEMISPHERE

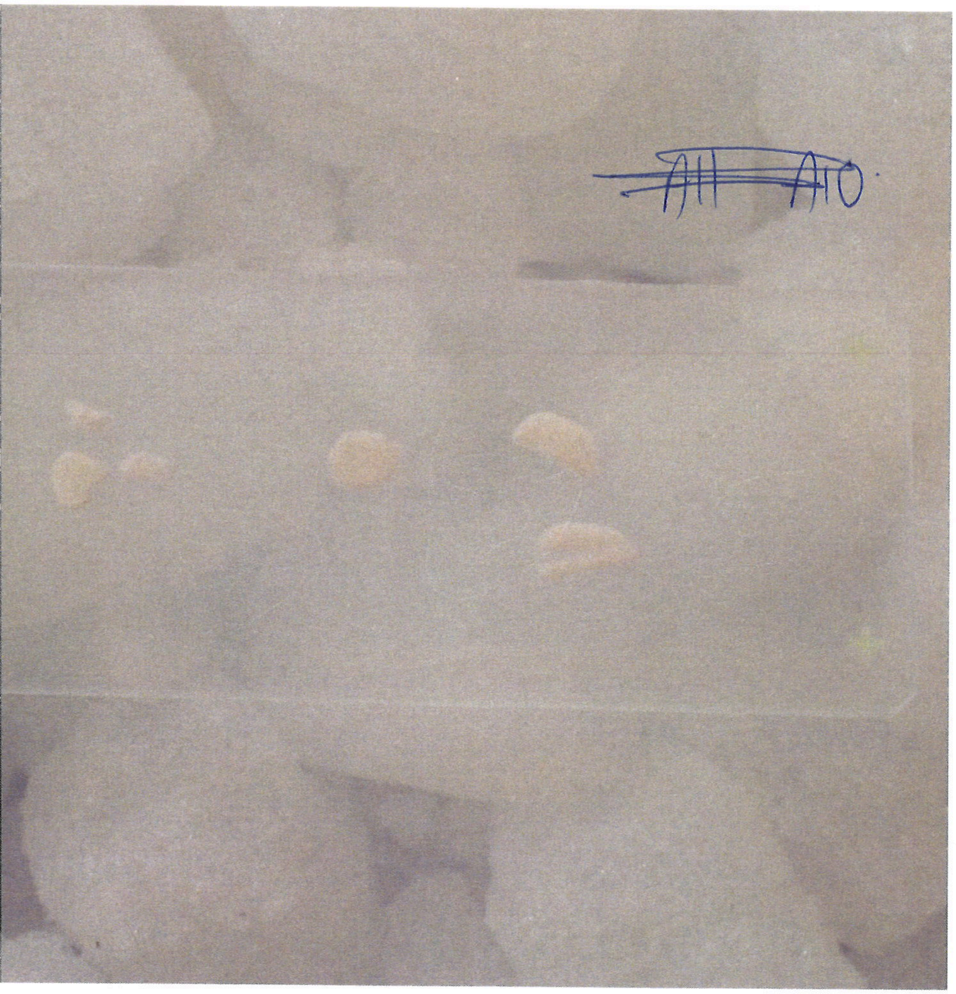

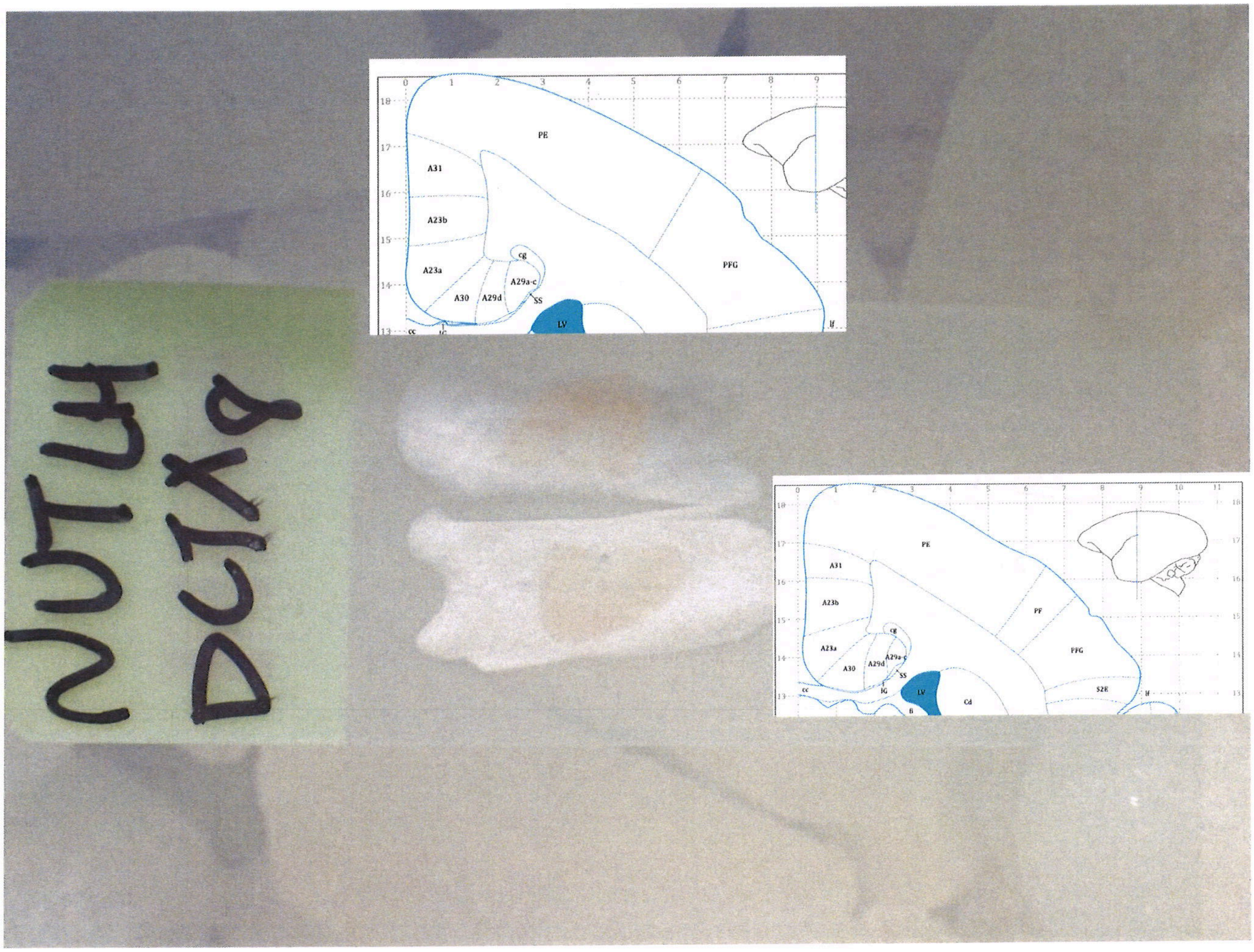


RIGHT HEMISPHERE

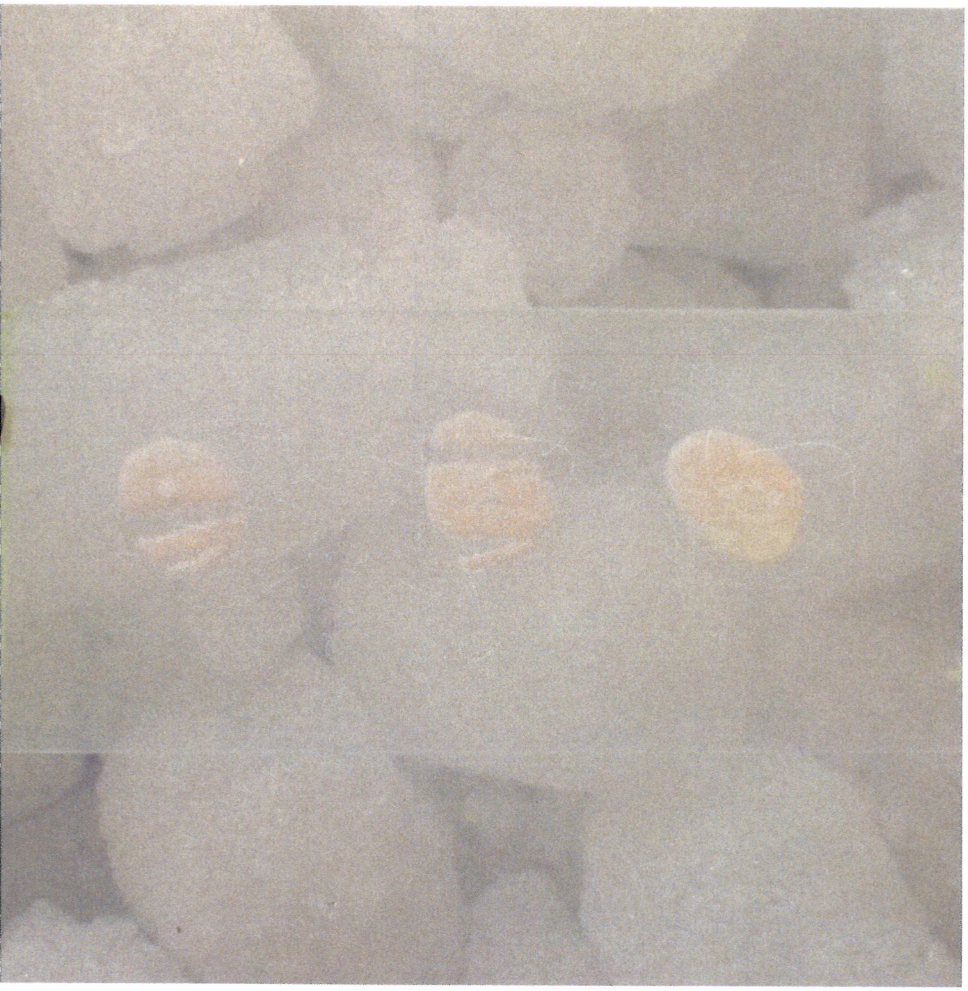

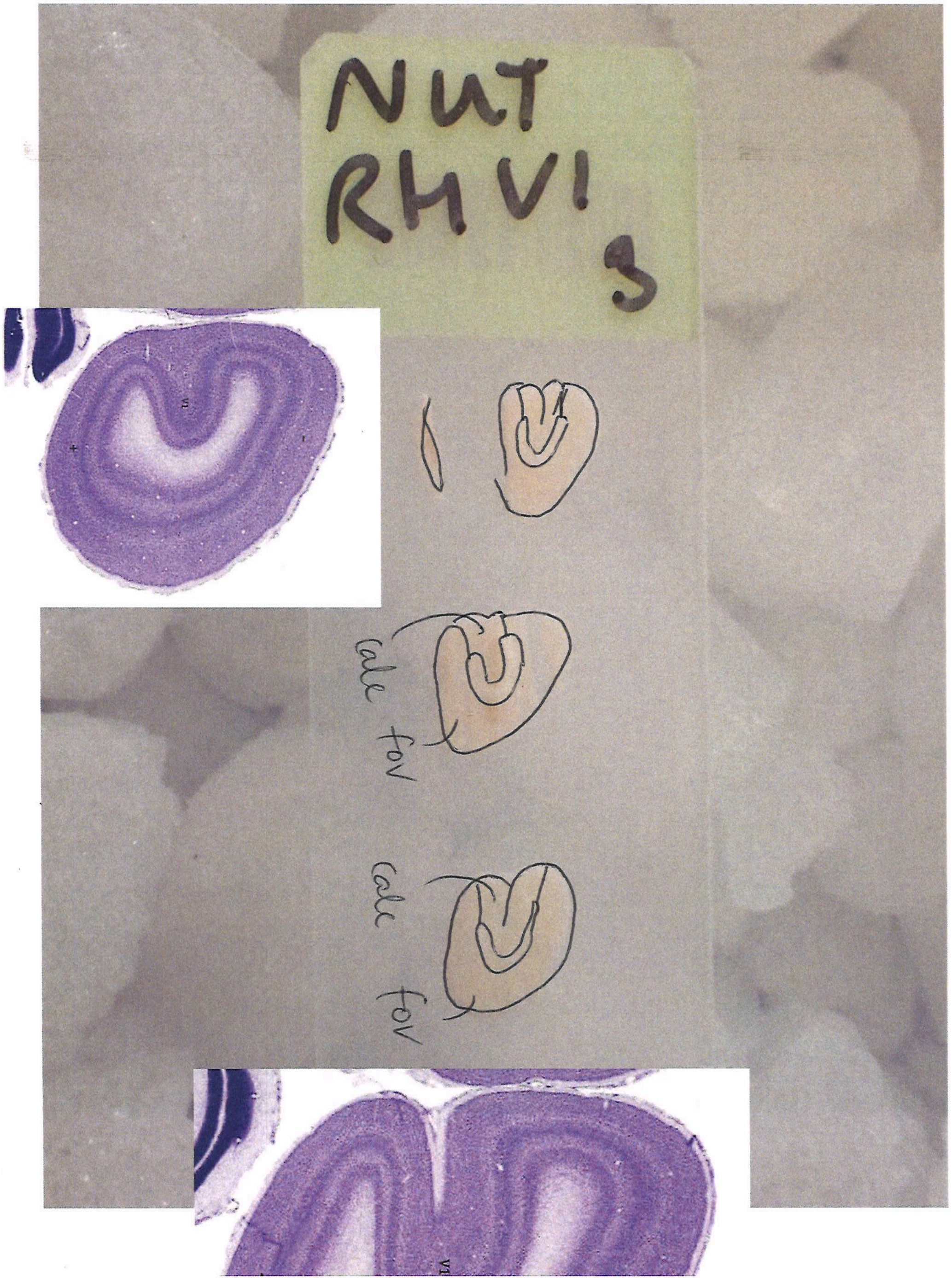
NUT
RHV1
3
calc
fov
calc
fov

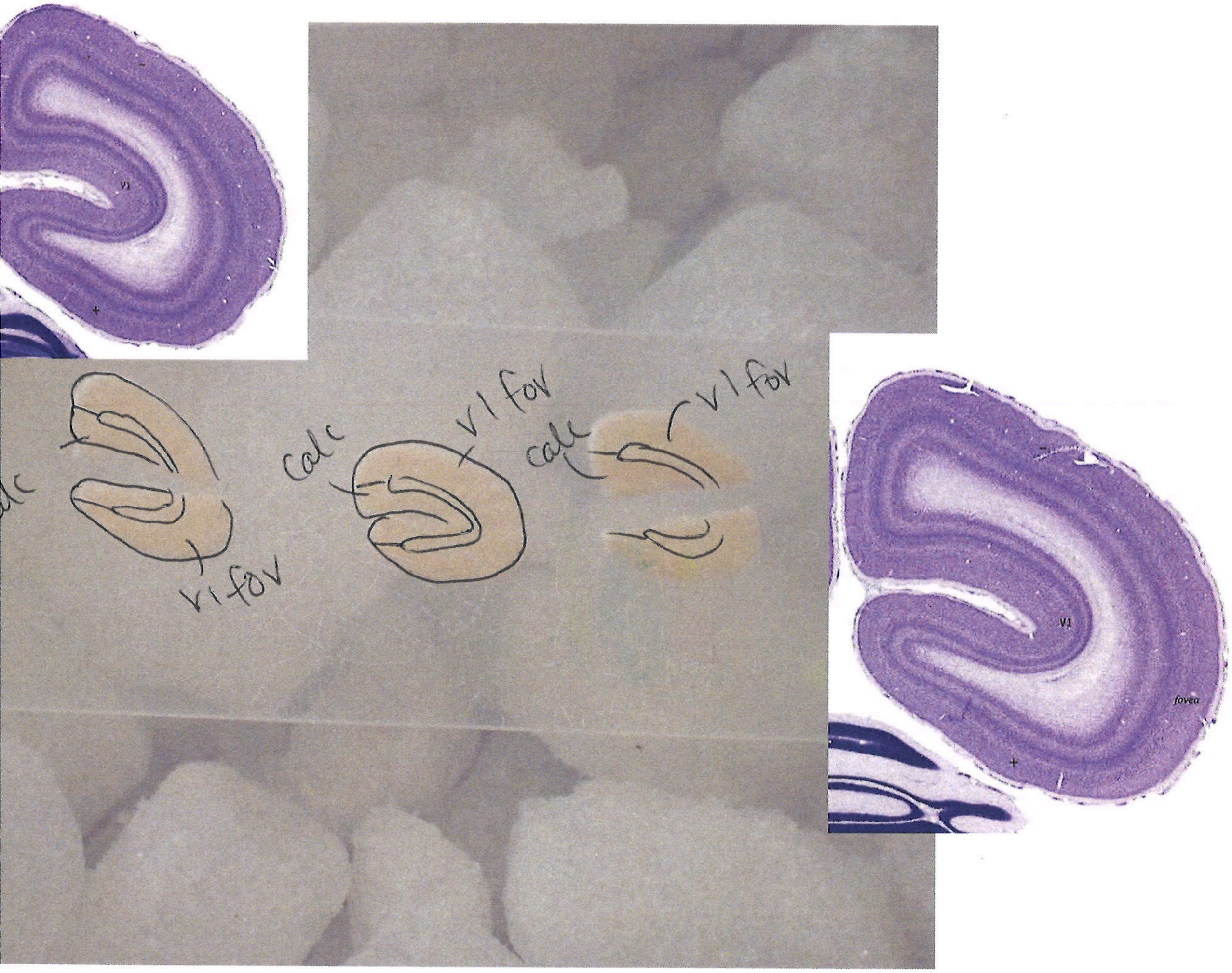
V1
v1 fov
calc
v1 fov
calc
v1 fov
V1
fovea

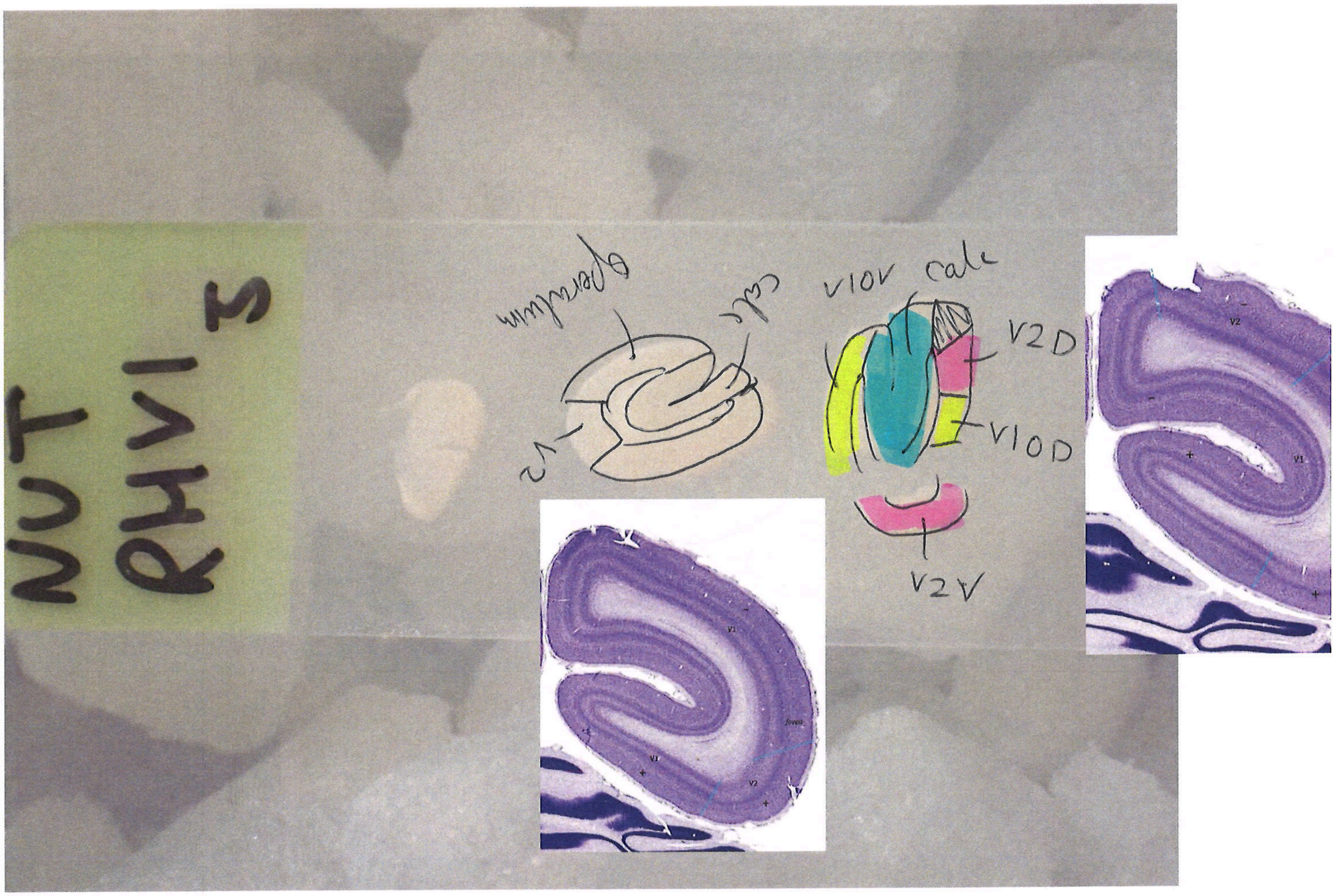
NUT
RHV1
3
operculum
calc
V2
V1OV
calc
V2D
V1OD
V2V

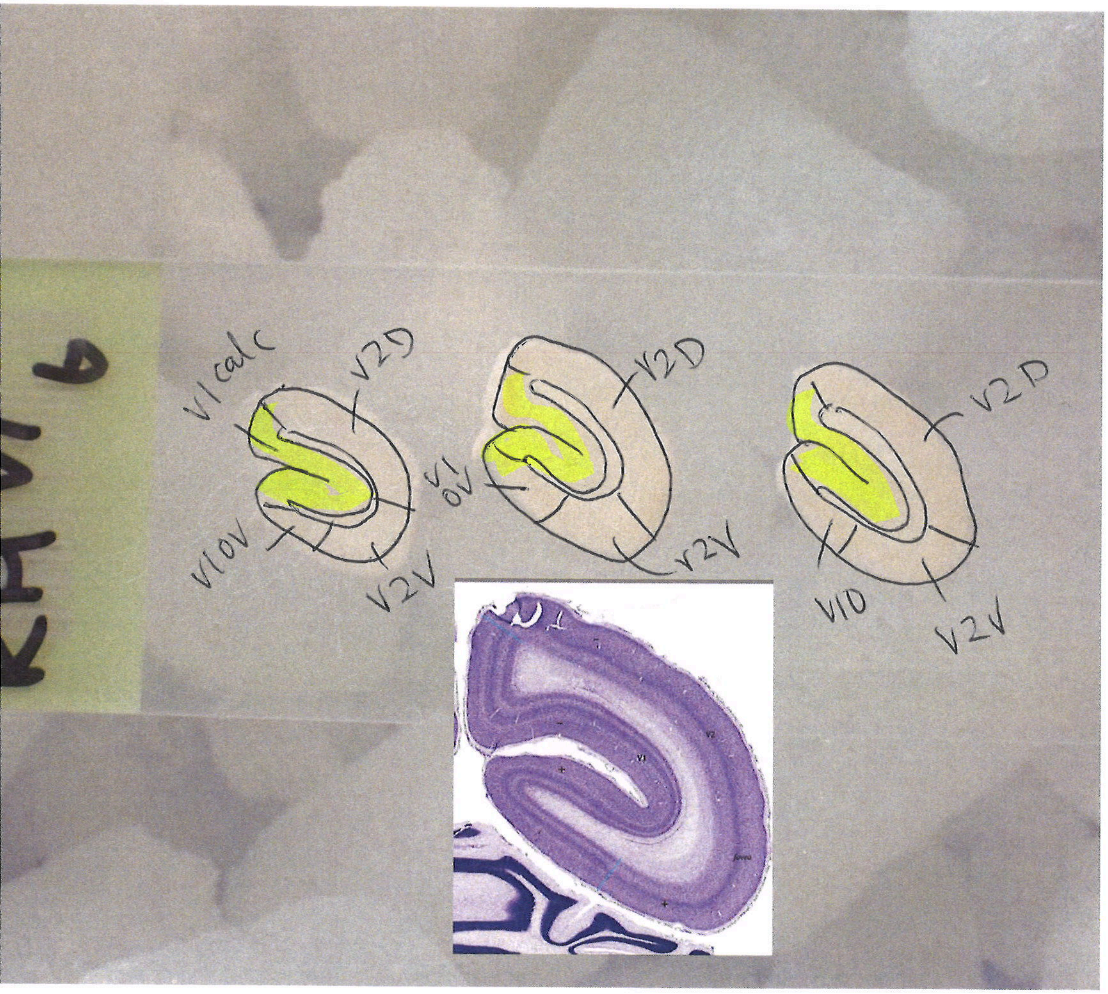
V1 calc
V2D
V1OV
V2V
V1 OV
V2D
V2V
V1D
V2D
V2V

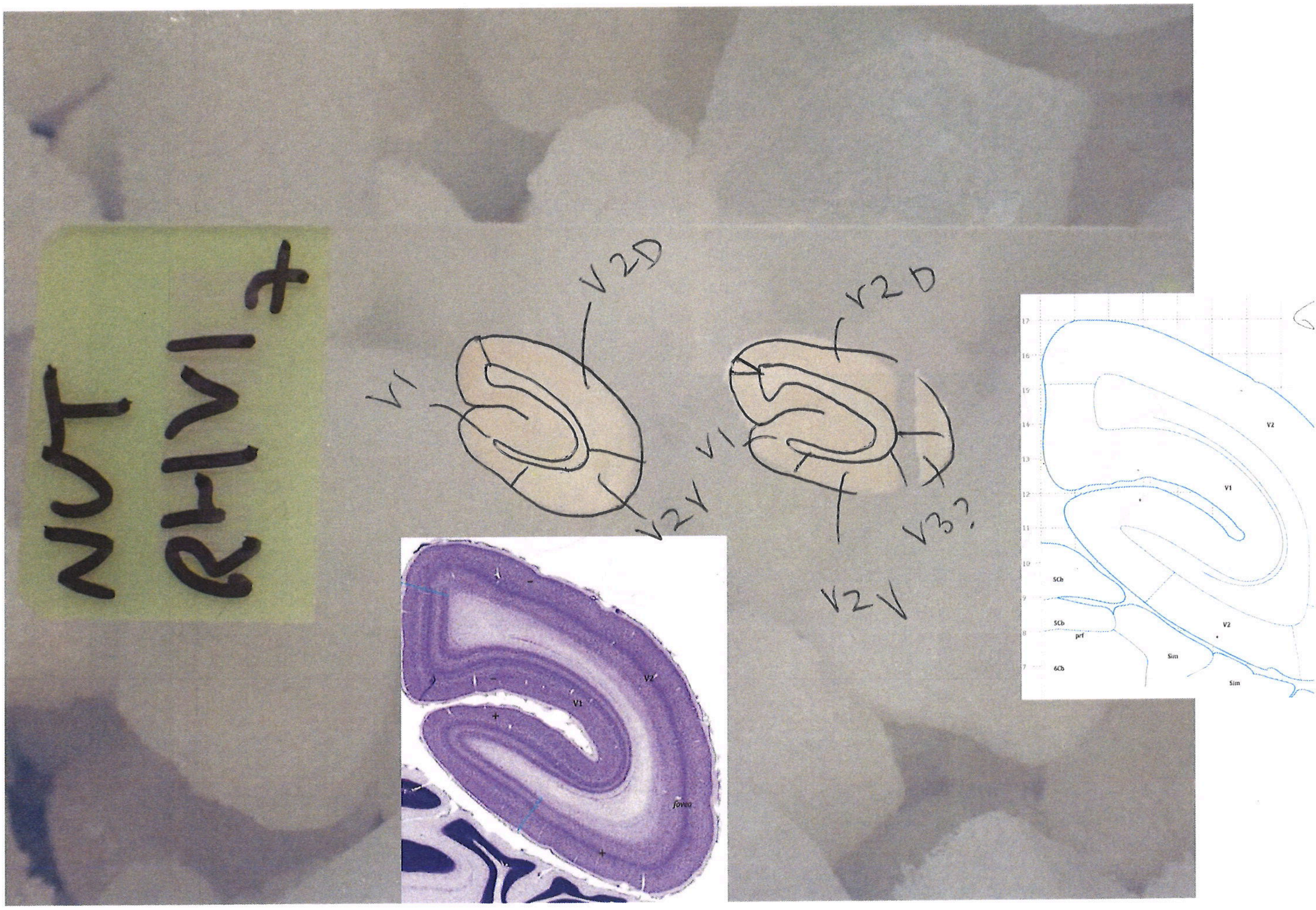
NUT
RH V1 +
V1
V2D
V2V
V1
V2D
V3?
V2V
V1
V2
fovea
V2
V1
V2
SCb
SCb
prf
6Cb
Sim
Sim

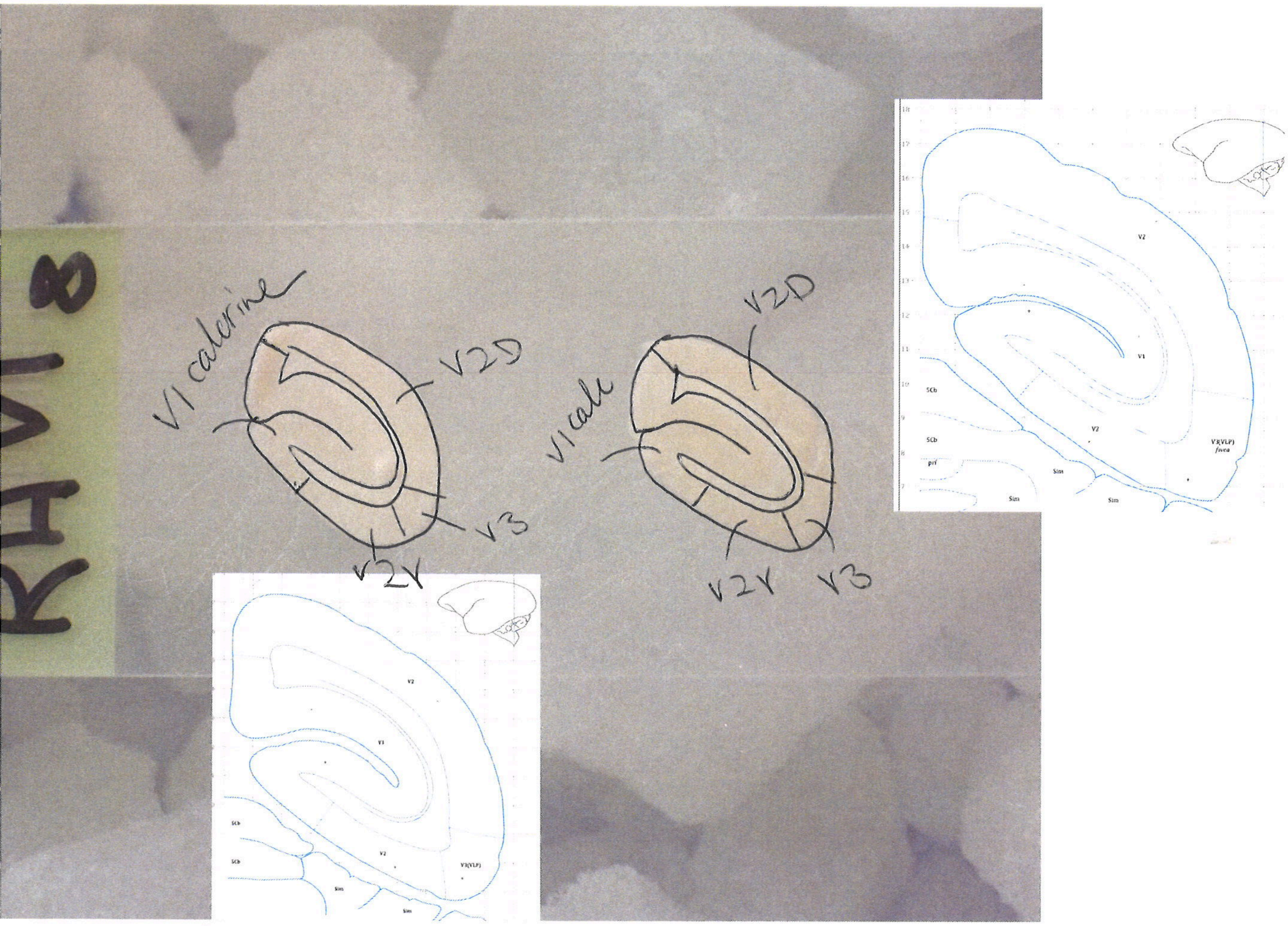
R4V1 8
V1 calcarine
V2D
V3
V2V
V1 calc
V2D
V2V
V3
V2
V1
SCb
Sim
V3(VLP)
fovea

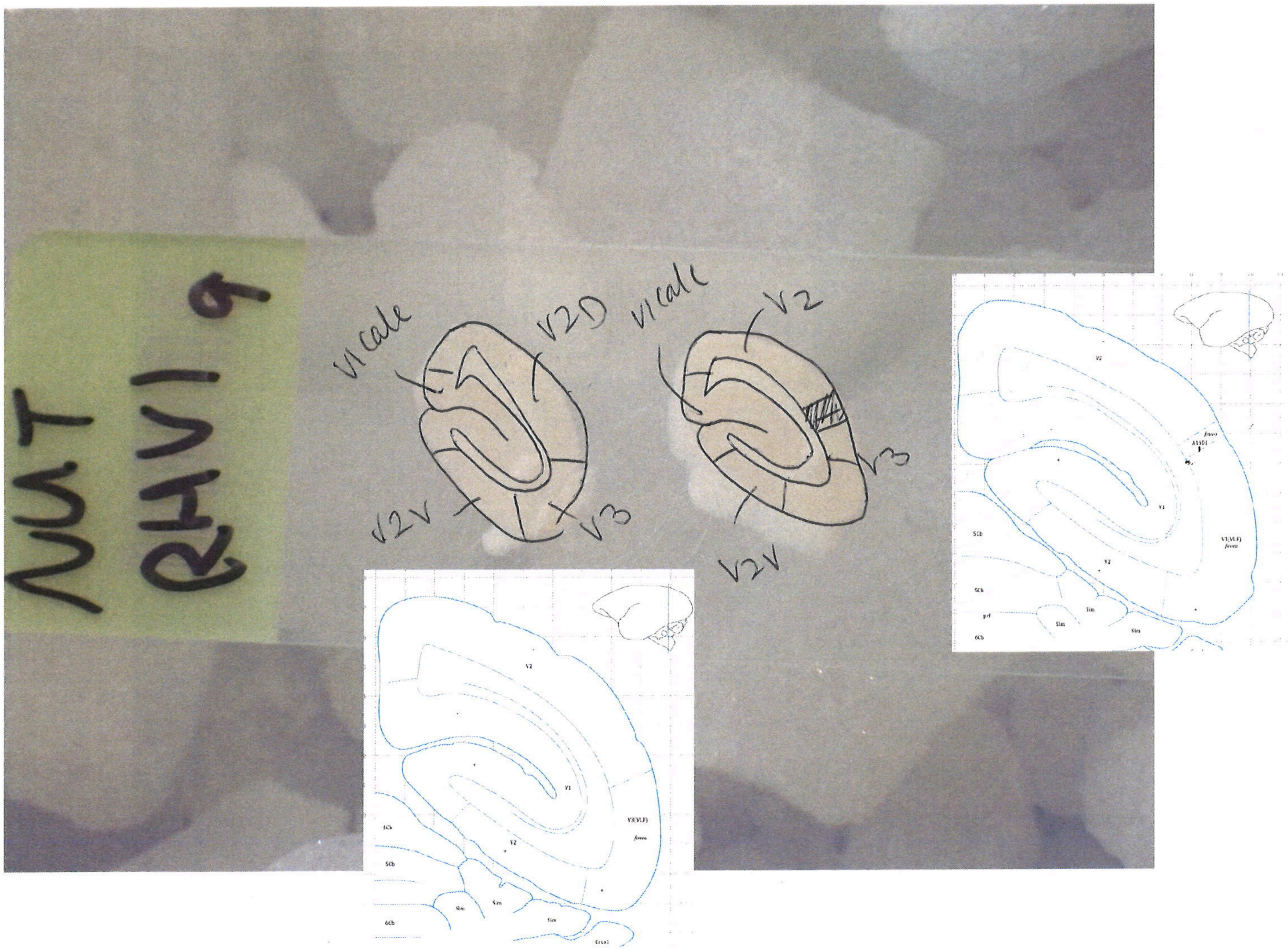
NUT
RHV1 9
V1 calc
V2D
V2V
V3
V1 calc
V2
V3
V2V

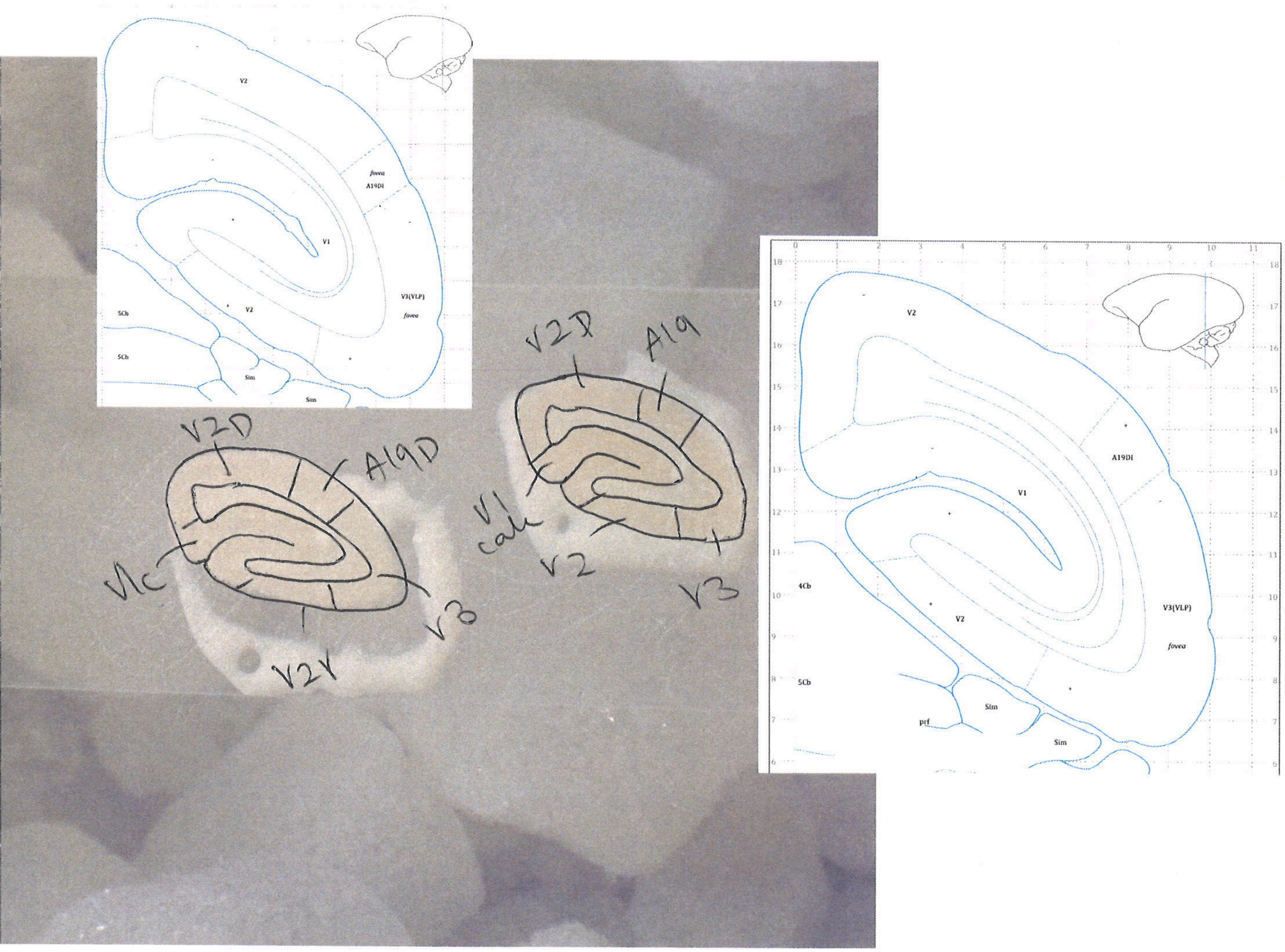
V2
fovea
A19DI
V1
V3(VLP)
fovea
5Cb
V2
5Cb
Sim
Sim
V2D
A19D
V1c
V3
V2V
V2D
A19
V1
calc
V2
V3
V2
A19DI
V1
4Cb
V2
V3(VLP)
fovea
5Cb
Sim
prf
Sim

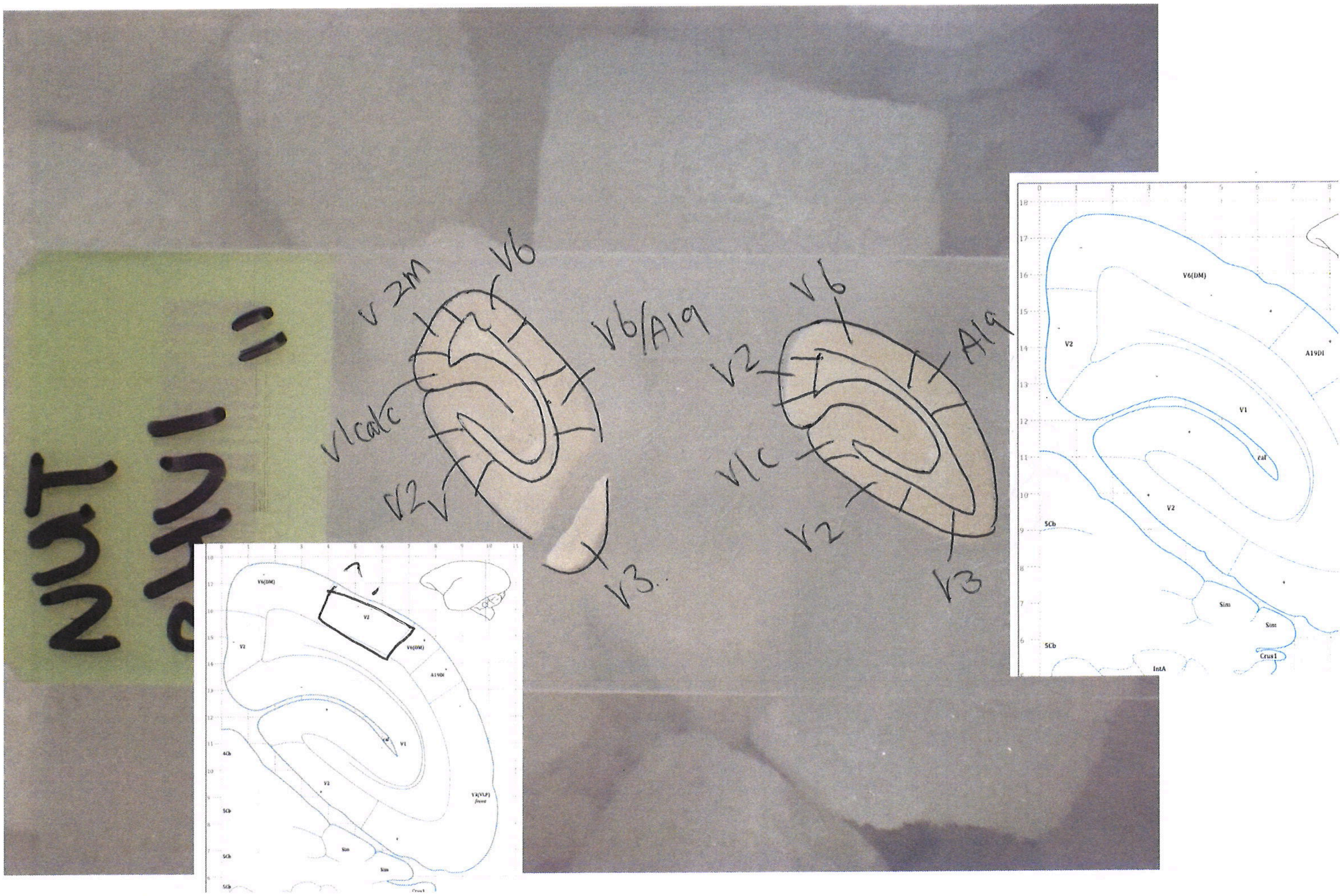
V2M
V6
V6/A19
V1calc
V2
V3.
V6
A19
V2
V1C
V2
V3
V6(DM)
V2
A19Dl
V1
cal
5Cb
Sim
Crus1
IntA

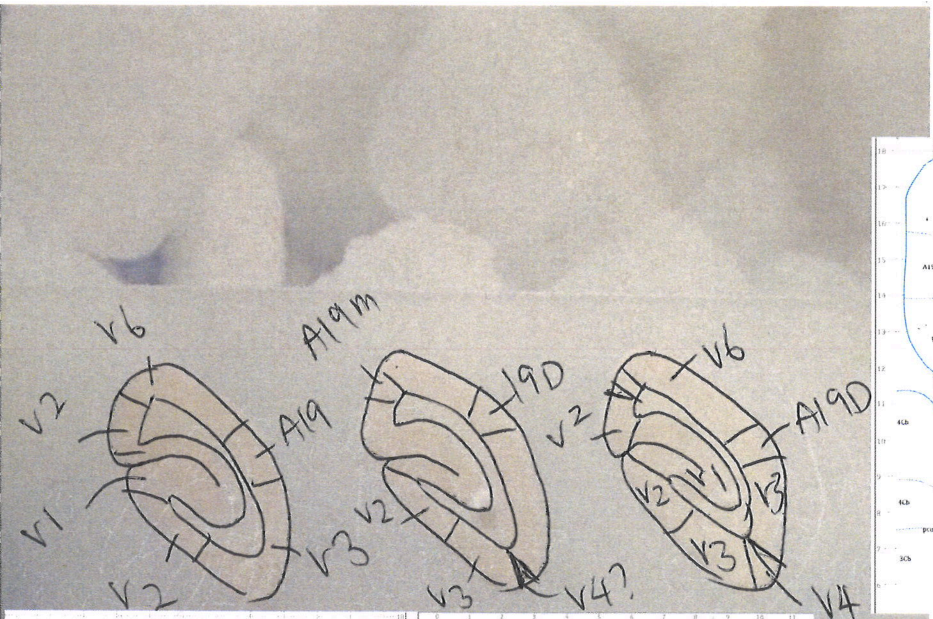
V6
V2
A19
V1
V2
V3
A19m
19D
V2
V3
V4?
V6
V2
A19D
V2
V1
V3
V3
V4

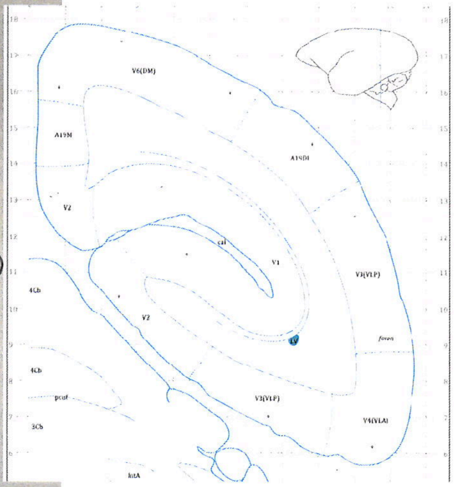
V6(DM)
A19M
A19DI
V2
cal
V1
V3(VLP)
4Cb
V2
fovea
LV
V3(VLP)
pcuf
3Cb
V4(VLA)
IntA

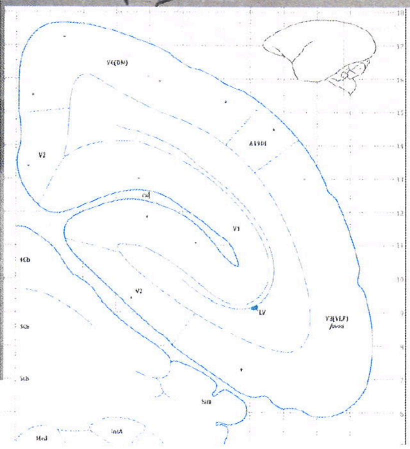
V6(DM)
A19DI
V2
cal
V1
4Cb
V2
LV
V3(VLP)
fovea
IntA

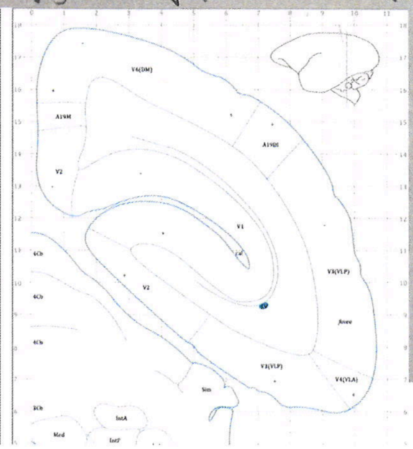
V6(DM)
A19M
A19DI
V2
V1
cal
4Cb
V2
V3(VLP)
fovea
V3(VLP)
V4(VLA)
Sim
3Cb
IntA
Med

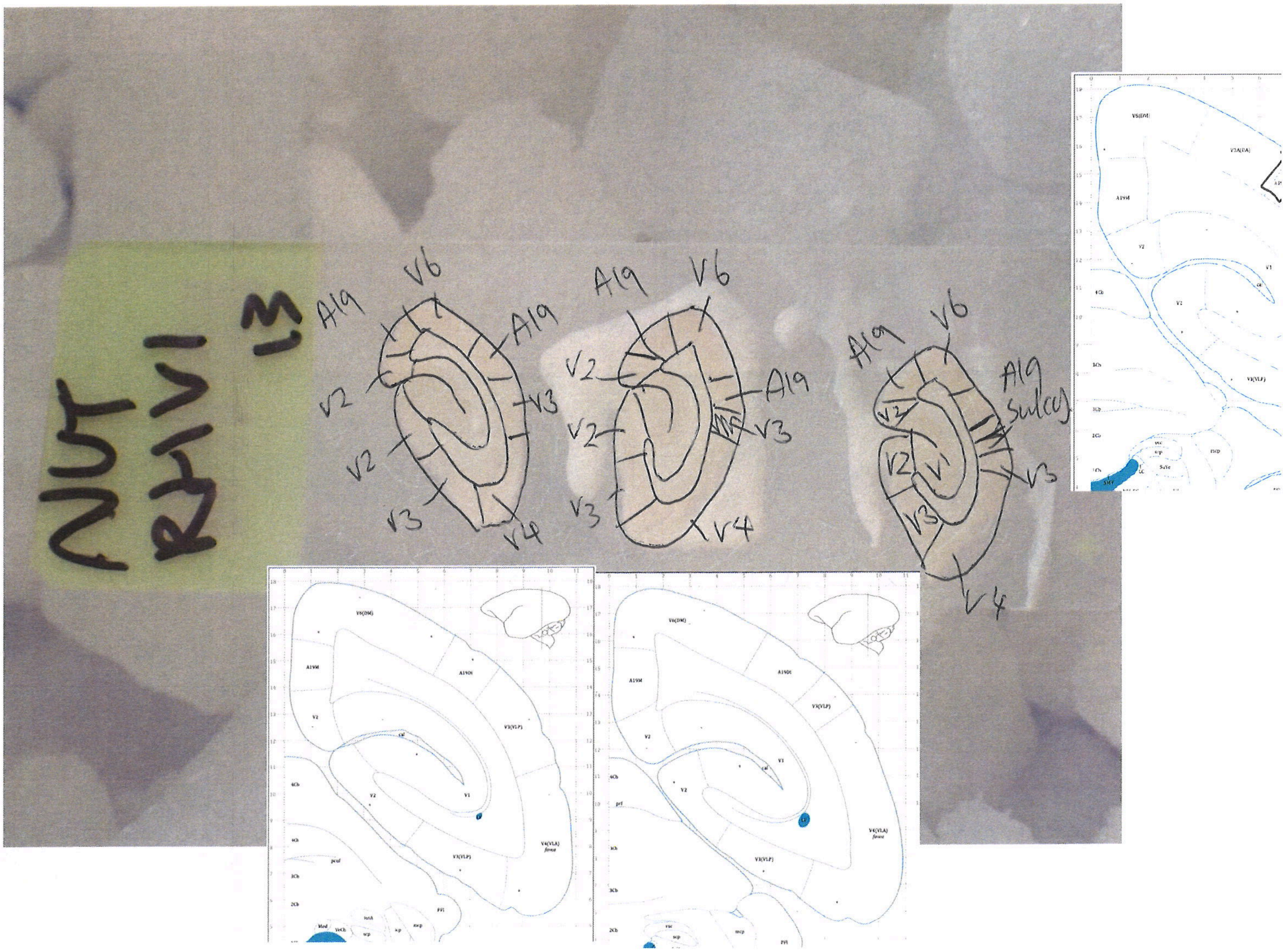
NUT
RH V1
13
A19
V6
A19
V2
V3
V2
V3
V4
A19
V6
V2
A19
V2
V3
V3
V4
A19
V6
A19
Sulcus
V2
V2
V1
V3
V3
V4
V6(DM)
A19M
A19DI
V3(VLP)
V2
V1
V4(VLA)
fovea
cal

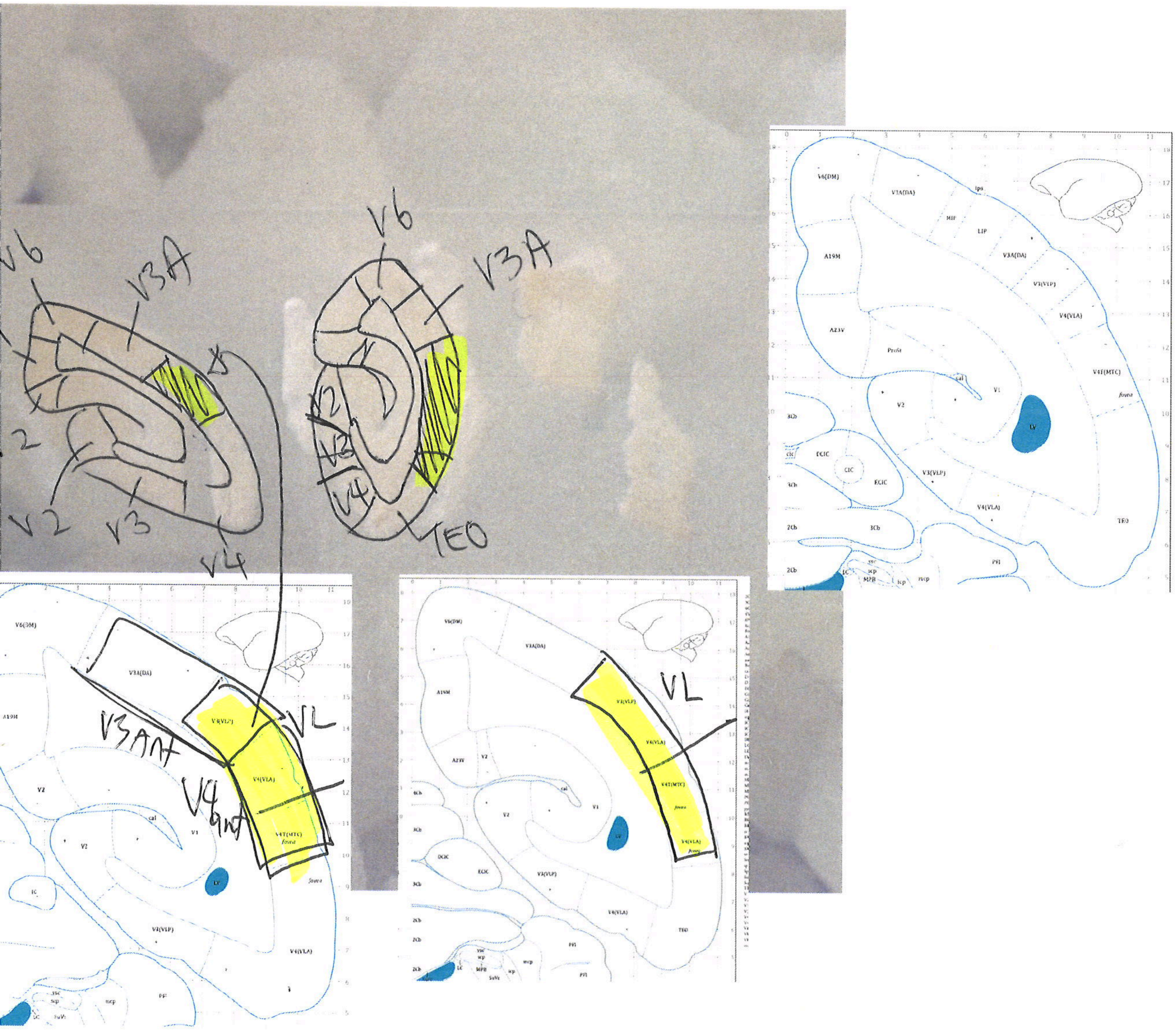

V6
V3A
V2
V3
V4
TEO
VL
V3ant
V4ant
V6(DM)
V3A(DA)
A19M
V3(VLP)
V4(VLA)
V4T(MTC)
fovea
V1
V2
LV
TEO

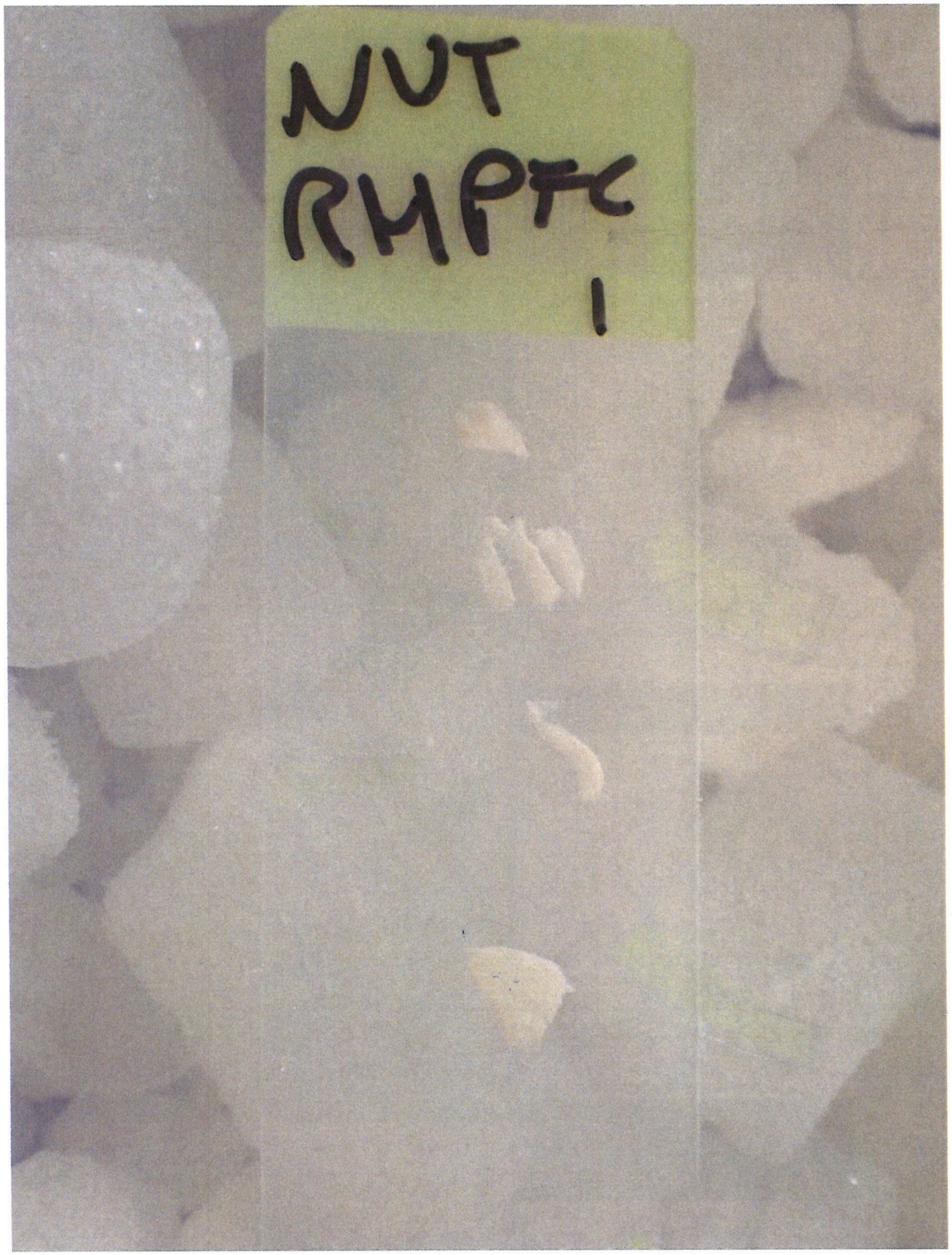
NUT
RMPFC
1

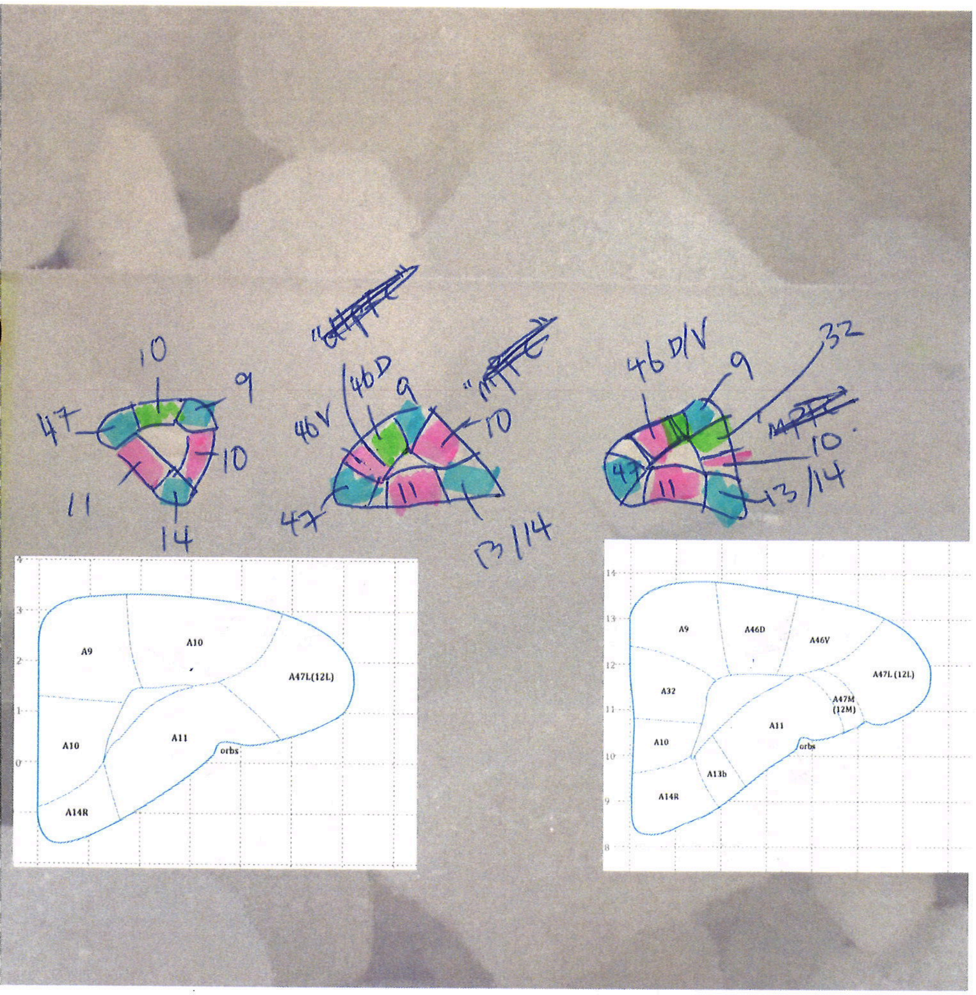
10
9
47
10
11
14
46V
46D
9
10
47
11
13/14
46D/V
9
32
10
11
13/14
A9
A10
A47L(12L)
A10
A11
orbs
A14R
A9
A46D
A46V
A47L (12L)
A32
A47M
(12M)
A11
A10
orbs
A13b
A14R

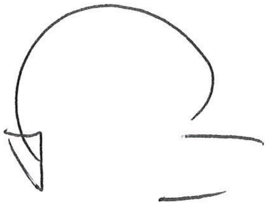

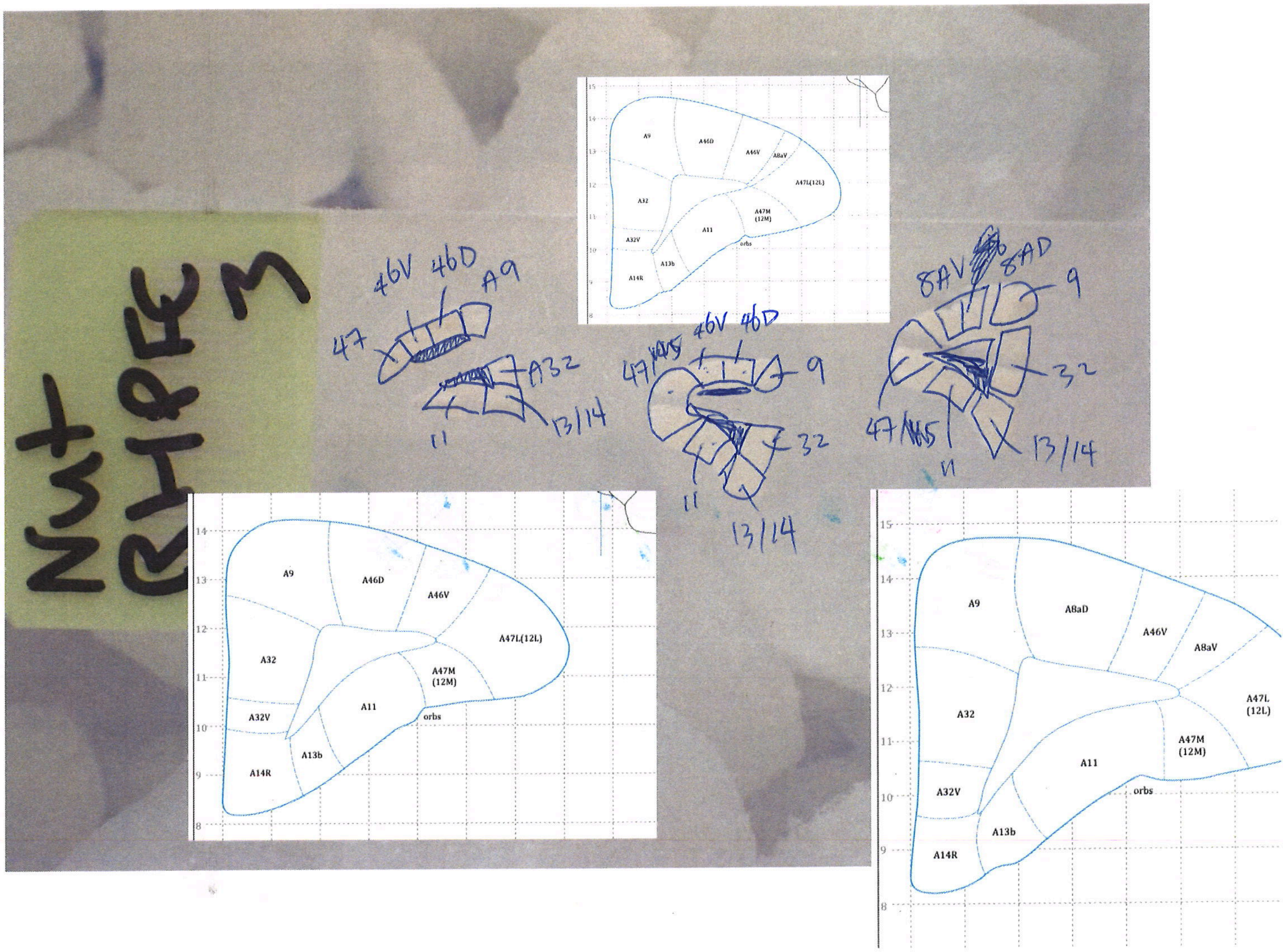
Nut
RHPFC 3
46V 46D A9
47
A32
11
13/14
46V 46D
47
9
32
8AV 8AD
47
A9
A46D
A46V
A8aV
A47L(12L)
A32
A47M
(12M)
A11
A32V
A13b
A14R
orbs
A8aD
A47L
(12L)

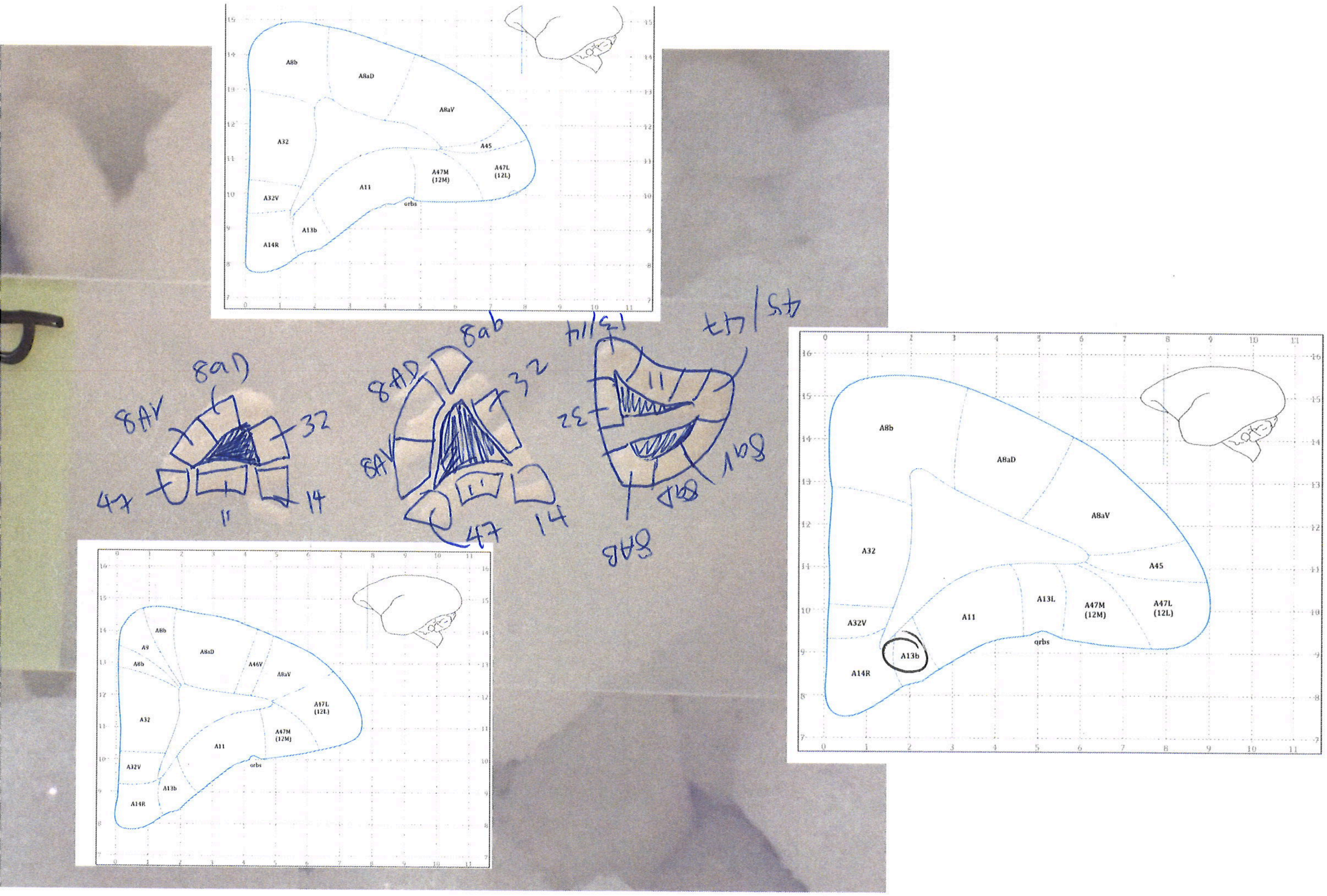
A8b
A8aD
A8aV
A45
A32
A47M
(12M)
A47L
(12L)
A11
A32V
orbs
A13b
A14R
8aD
8AV
32
47
11
14
8ab
8AD
8AV
47
14
13/14
45/47
32
A8b
A8aD
A8aV
A46V
A47L
(12L)
A47M
(12M)
A11
A32
A32V
A13b
A14R
orbs
A8b
A8aD
A8aV
A45
A32
A13L
A47M
(12M)
A47L
(12L)
A11
A32V
A13b
orbs
A14R

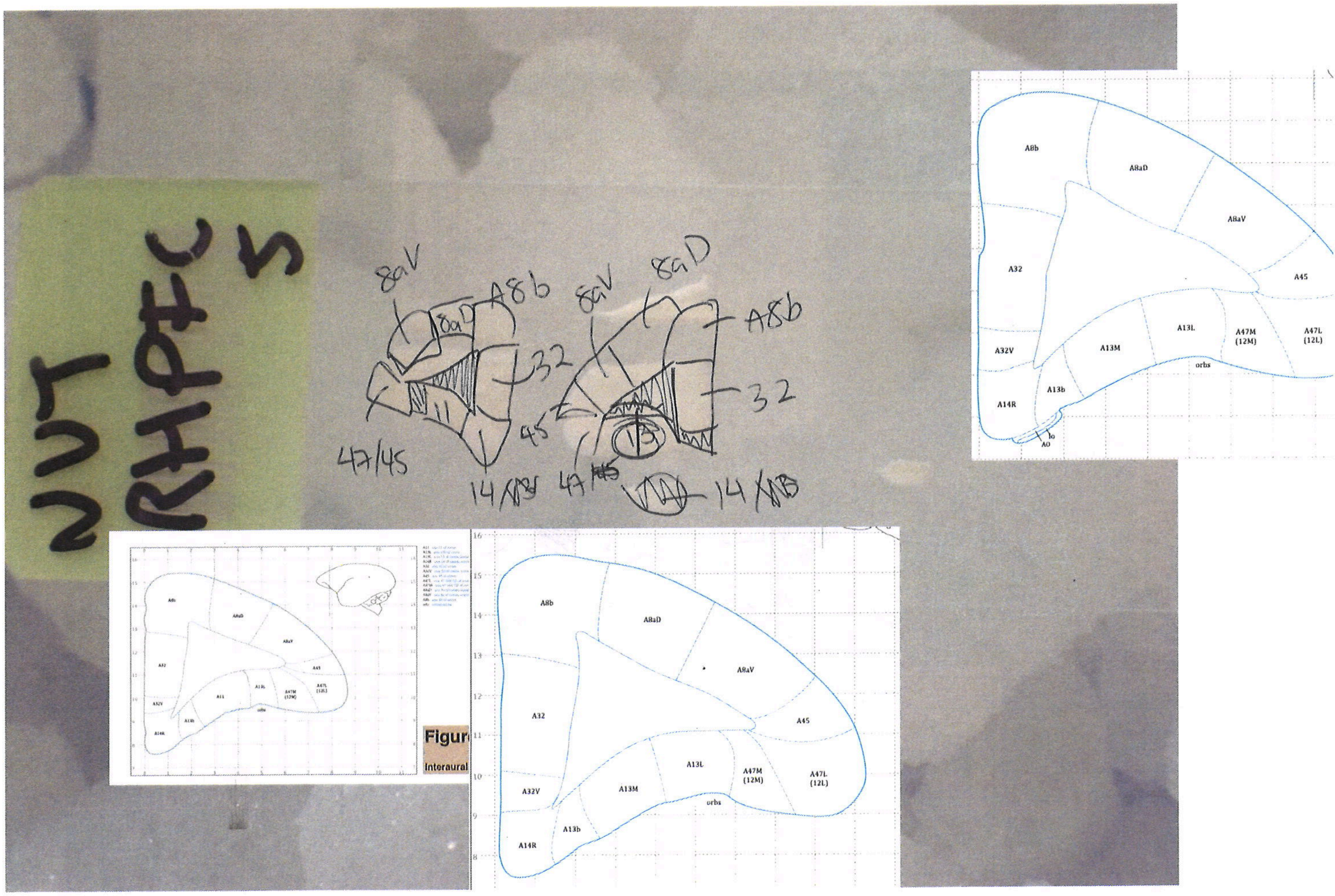

NUT
RHPFC
5
8aV
8aD
A8b
32
47/45
45
14
A8b
A8aD
A8aV
A32
A45
A32V
A13M
A13L
A47M (12M)
A47L (12L)
orbs
A13b
A14R
Figur
Interaural

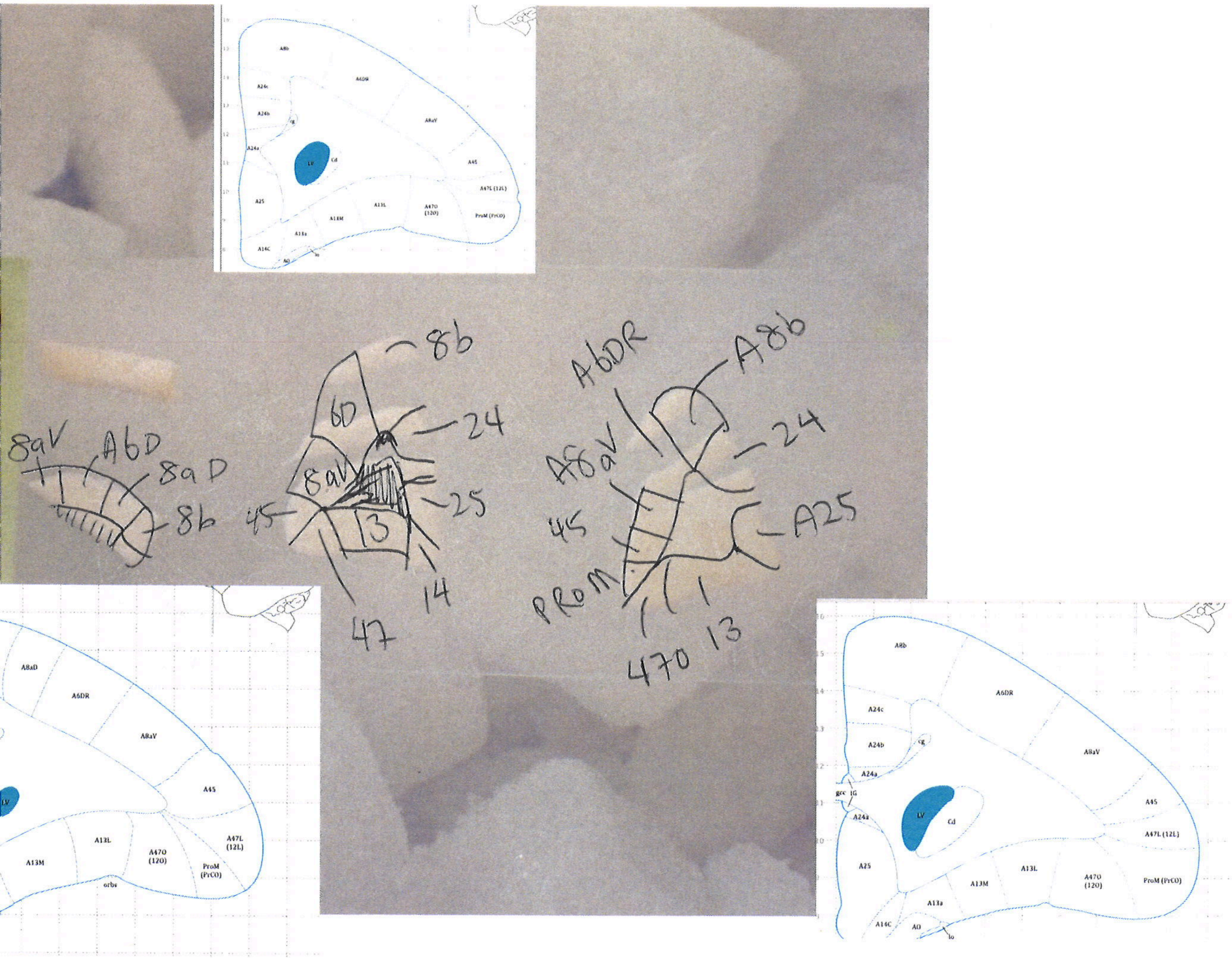
8aV
A6D
8aD
8b
6D
8b
24
8aV
25
45
13
47
14
A6DR
A8b
24
A8aV
45
A25
PROM
470
13
A8b
A6DR
A24c
A24b
A8aV
A24a
LV
Cd
A45
A47L (12L)
A25
A13L
A47O (12O)
A13M
ProM (PrCO)
A13a
A14C
AO
A8aD

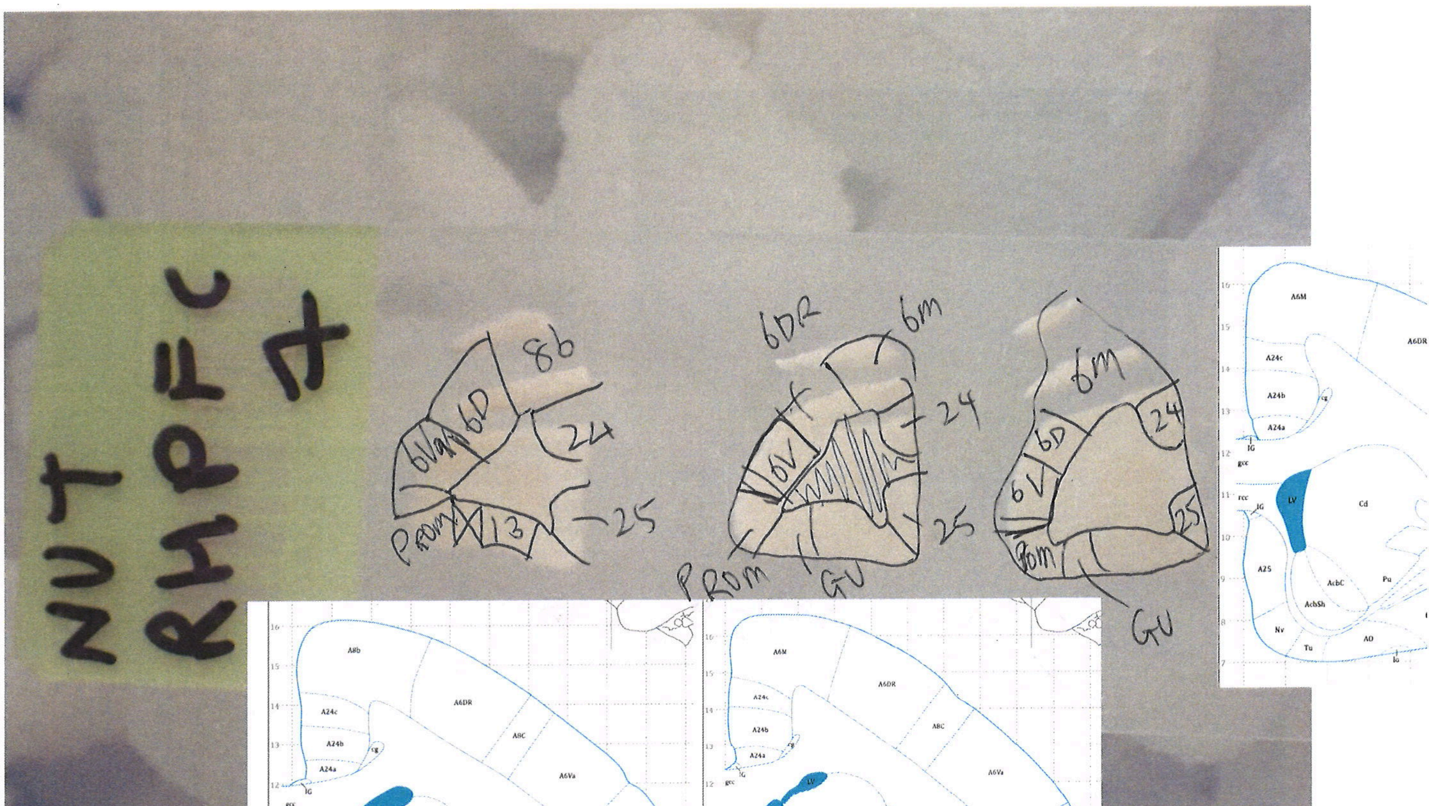
NUT
RHPFC
4
8b
6Va
6D
24
13
PROM
25
6DR
6m
6V
GU
6m
6D
6V
24
25
Prom
GU

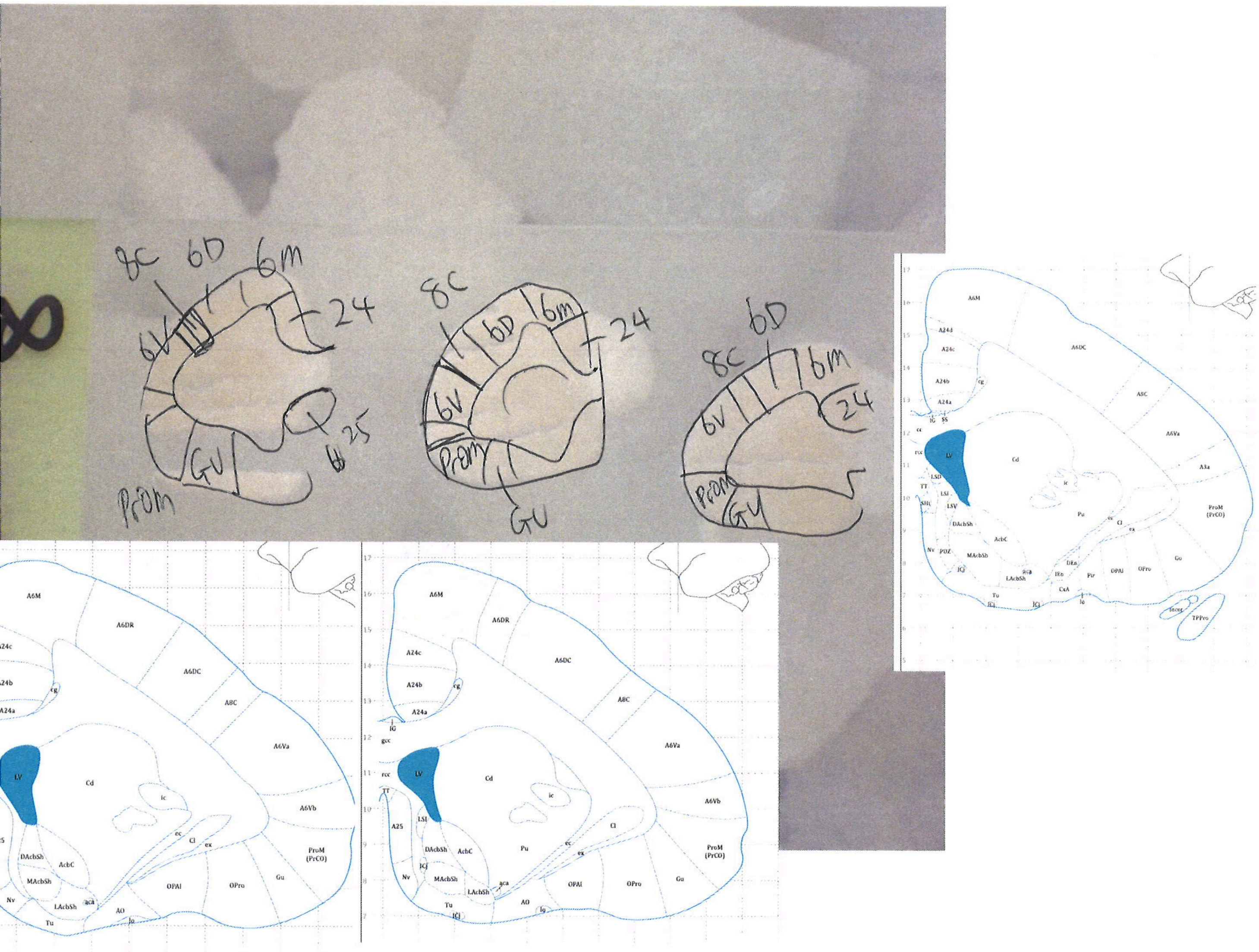

8C
6D
6m
24
6V
GU
25
ProM
A6M
A6DR
A6DC
A8C
A6Va
A6Vb
A24d
A24c
A24b
A24a
A25
A3a
cg
IG
gcc
rcc
TT
LV
Cd
ic
Pu
ec
Cl
ex
LSI
LSD
LSV
SHi
DAcbSh
AcbC
MAcbSh
LAcbSh
aca
Nv
ICj
Tu
AO
lo
OPAl
OPro
Gu
ProM (PrCO)
Pir
IEn
DEn
CxA
Incez
TPPro

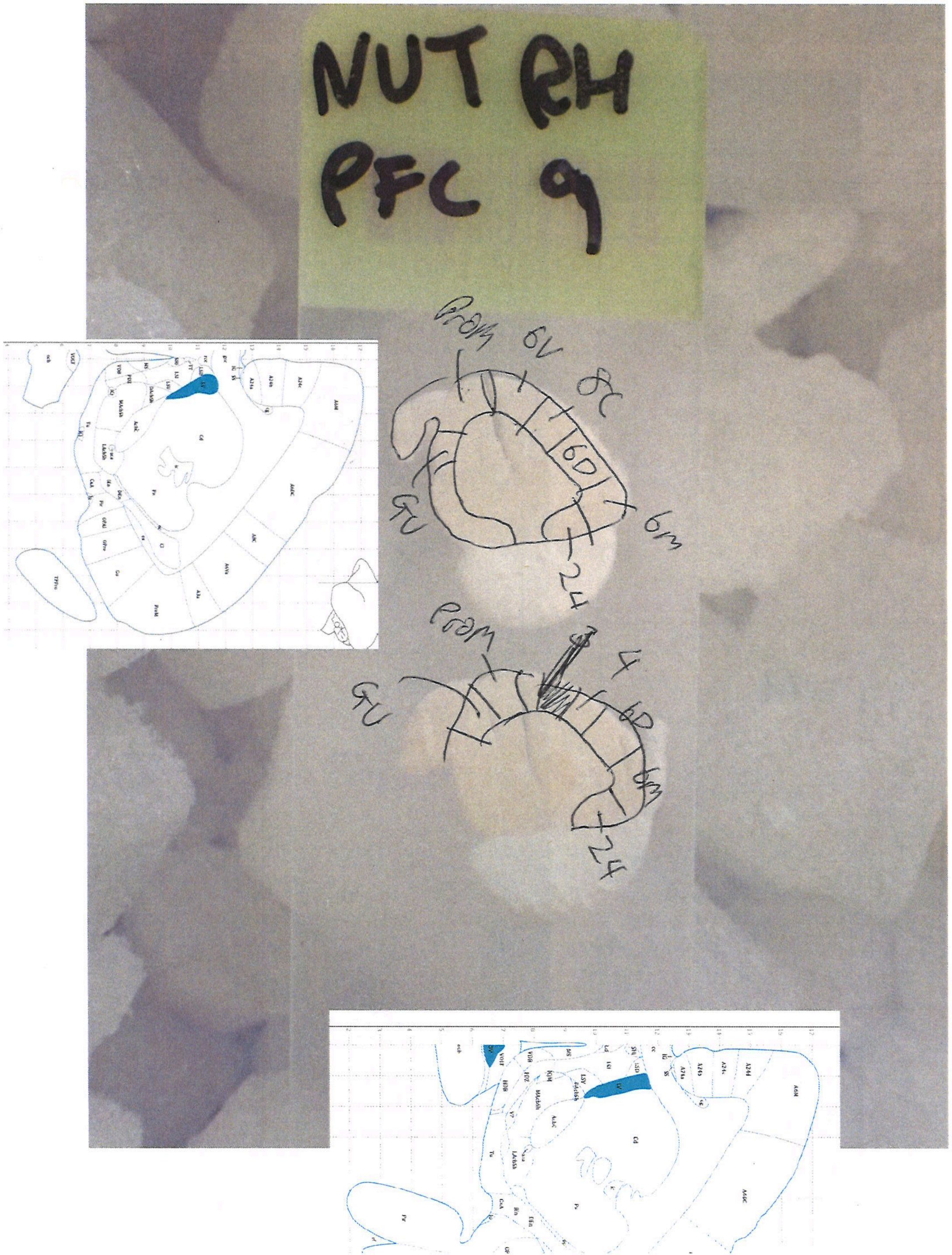
NUT RH
PFC 9
ProM
6V
8C
6D
6m
GU
24
4
6D
6m
24

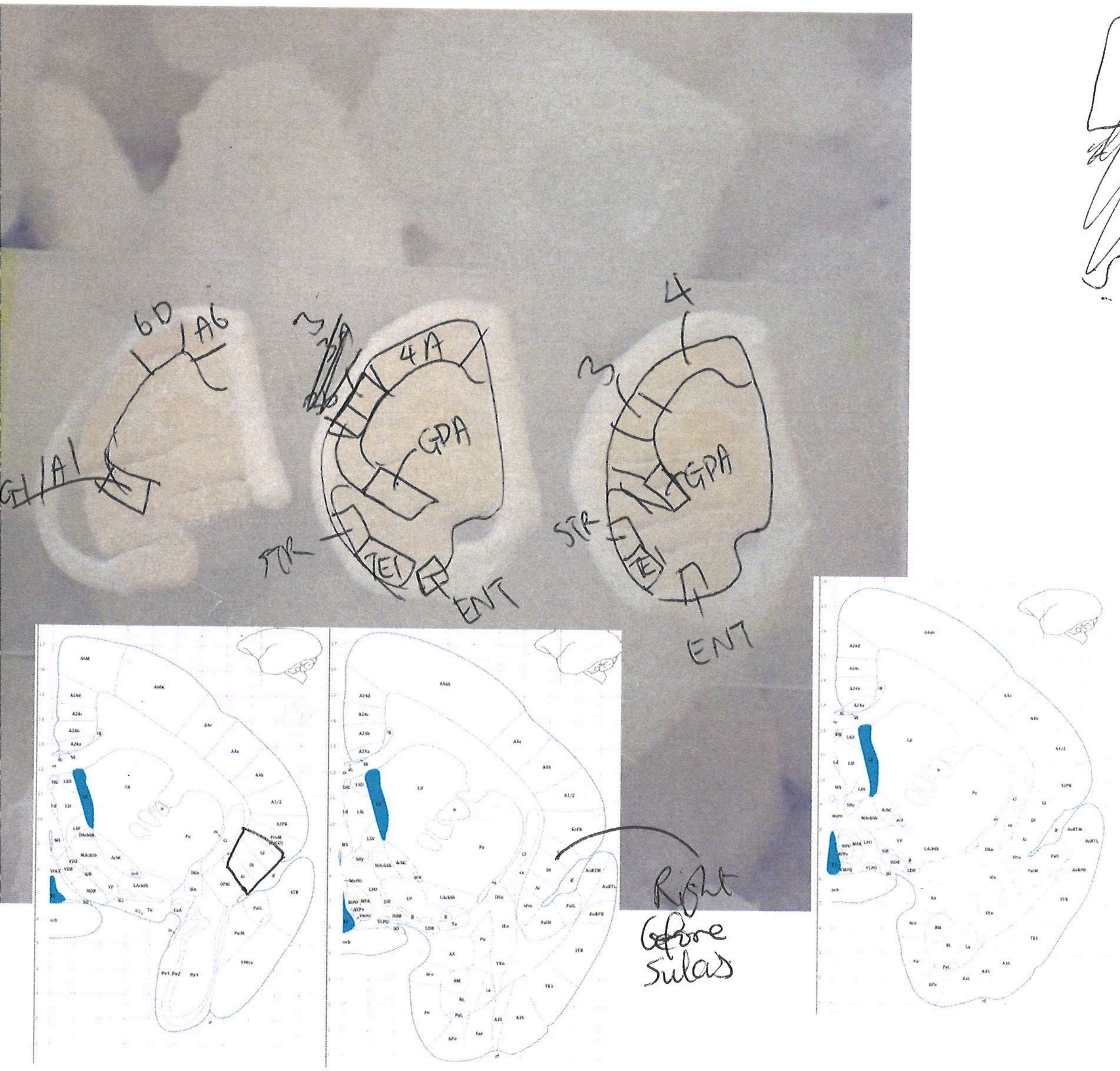
6D
A6
GI/AI
4A
GPA
STR
TE1
ENT
3
4
GPA
STR
TE1
ENT
Right before Sulcus

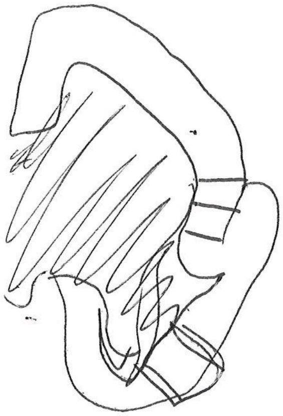

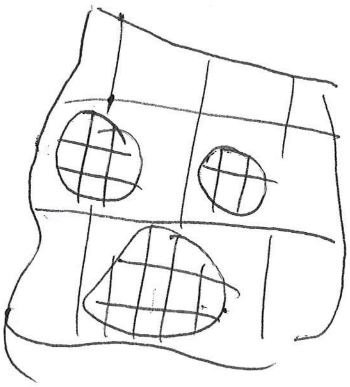

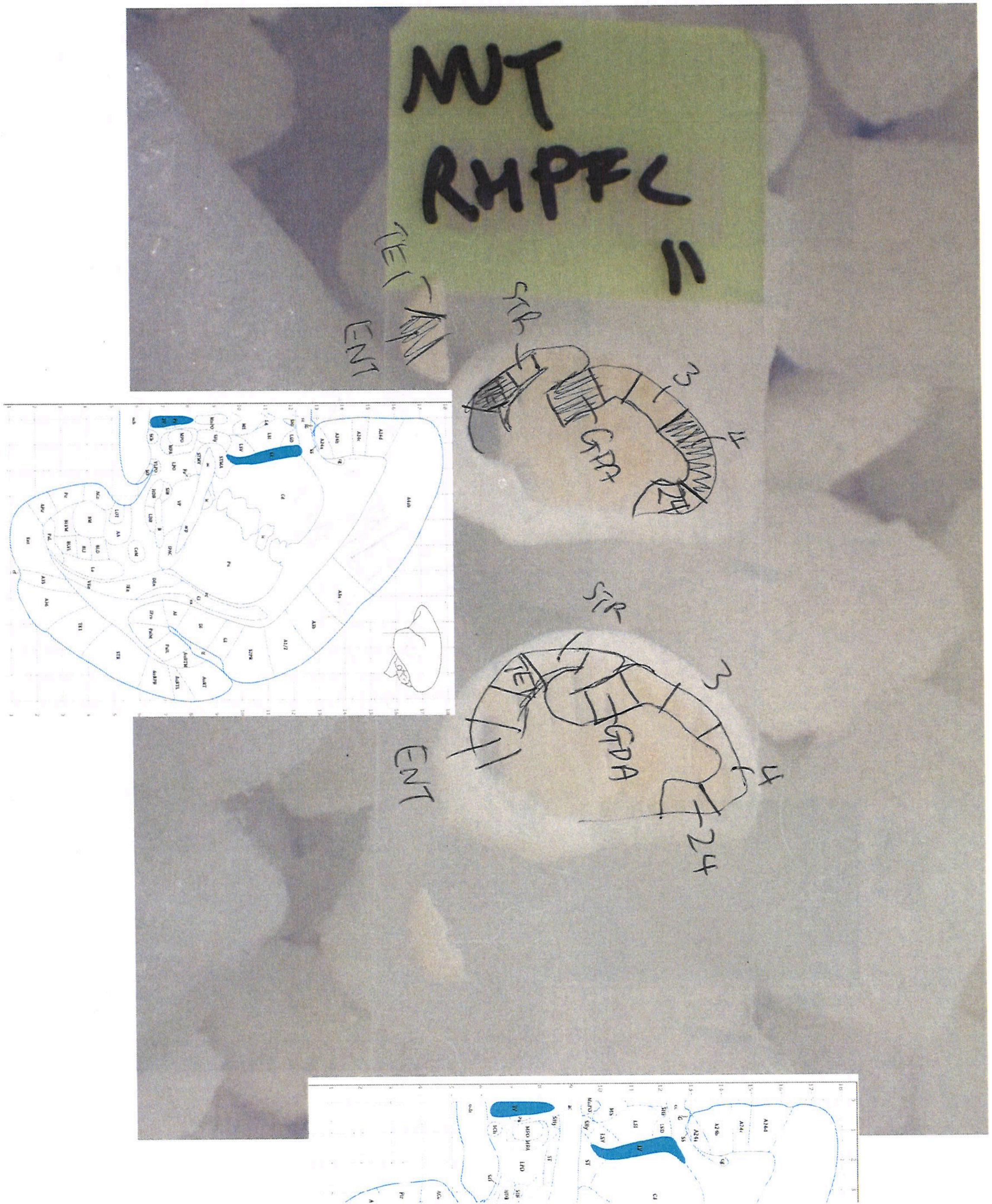
MUT
RHPFC
11
TET
ENT
STR
3
4
GPA
24
STR
TET
3
GPDA
ENT
4
24

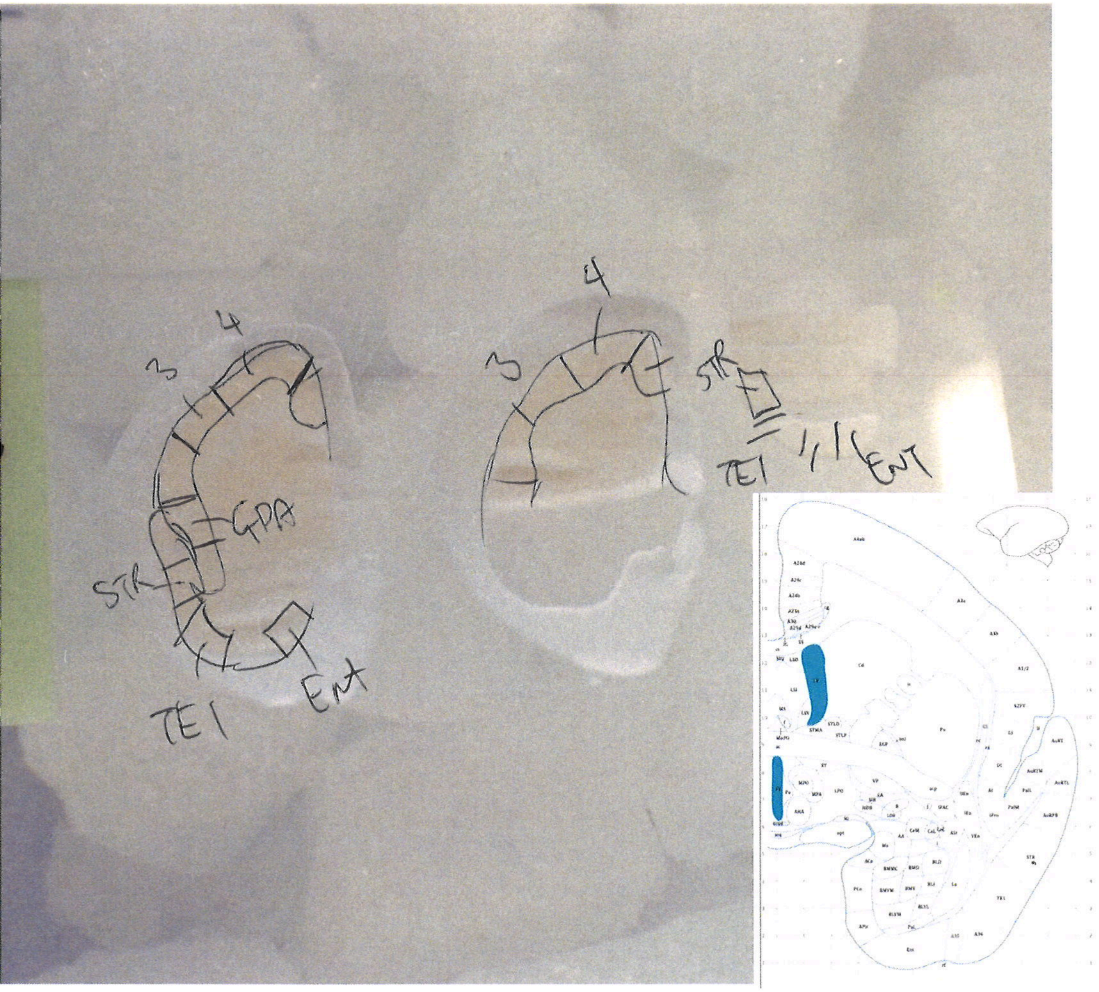

3
4
GPA
STR
TE1
ENT
3
4
STR
TE1
ENT

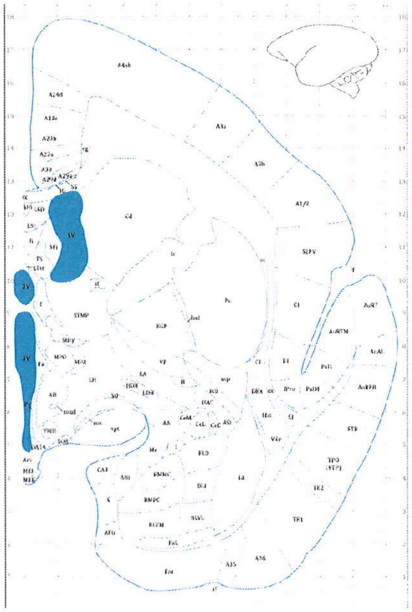

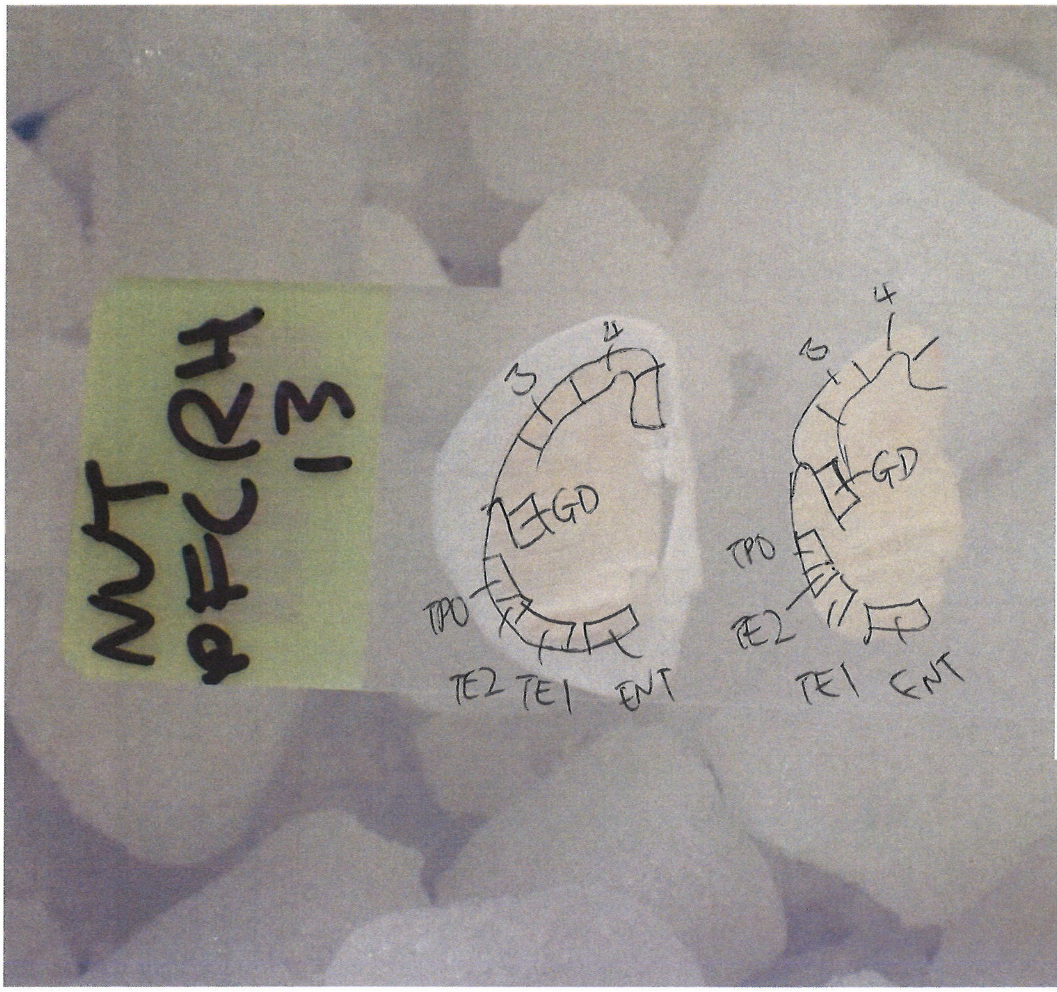
NUT PFC RH 1-3
GD
GD
TPO
TPO
TE2
TE2 TE1 ENT
TE1 ENT
3
4
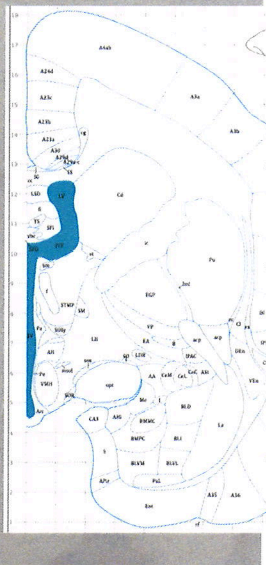
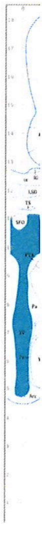

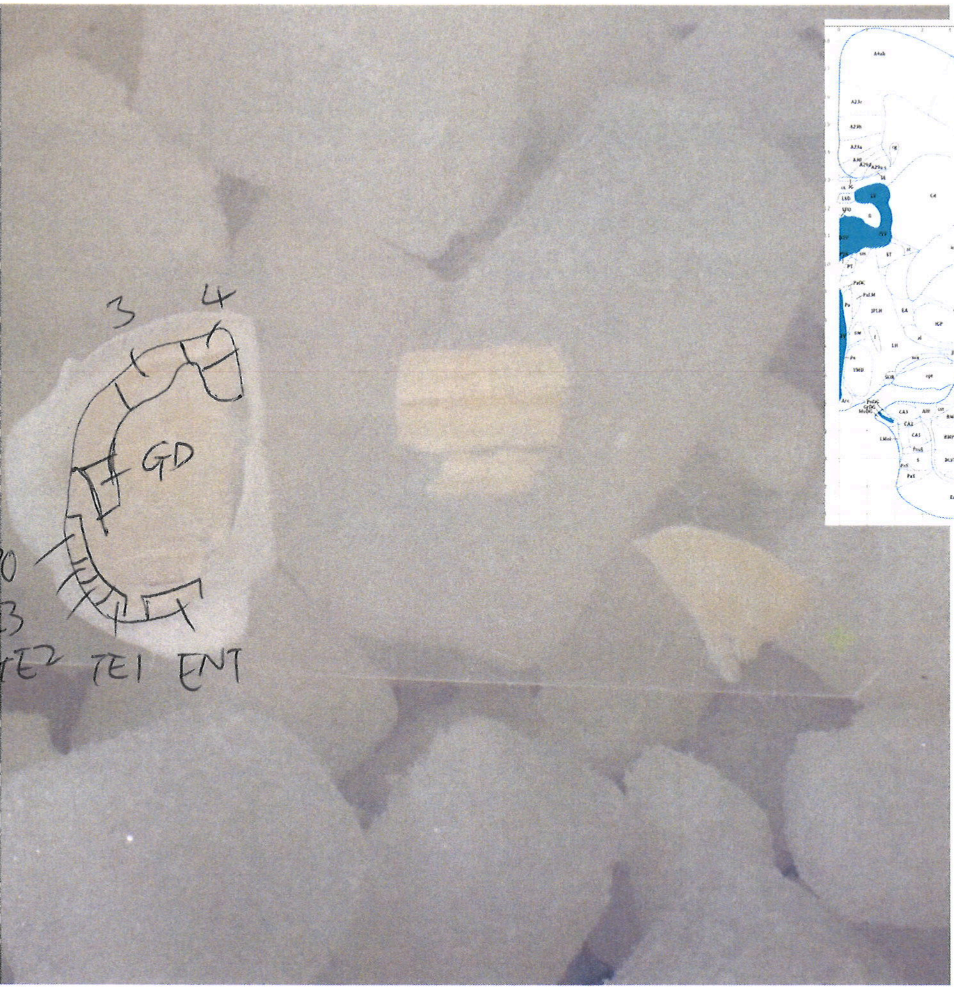
3
4
GD
TE1
ENT

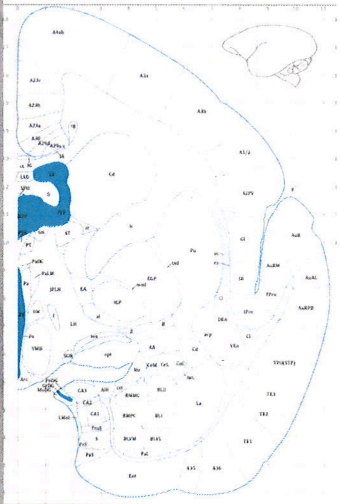

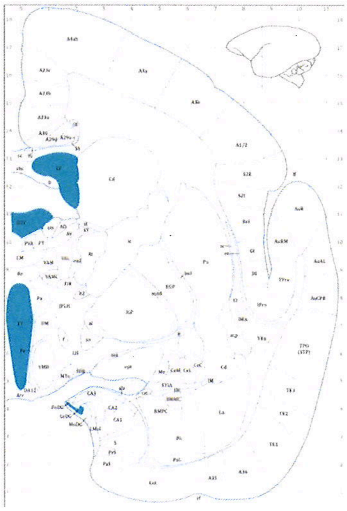

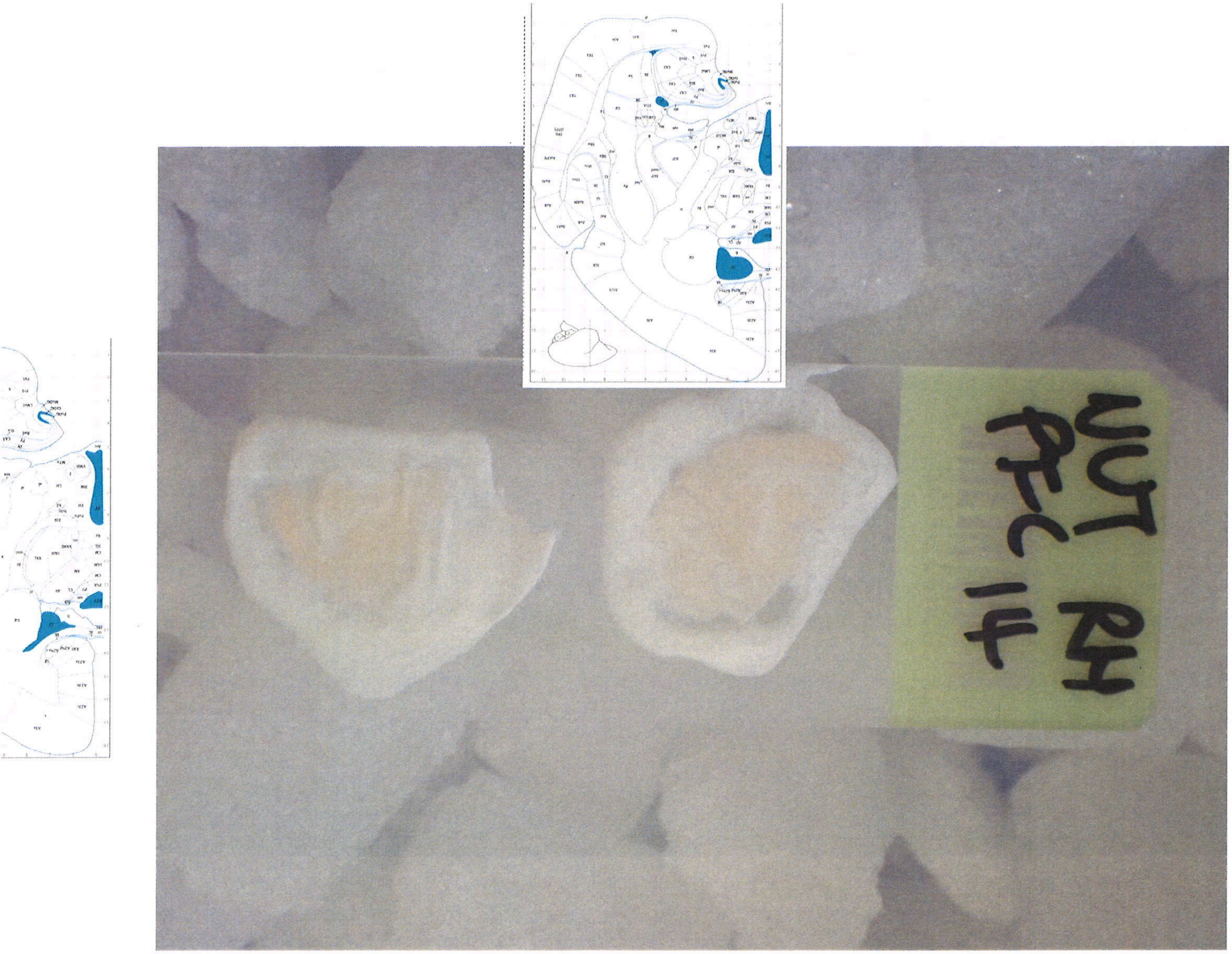
WT RH
PFC 1-4

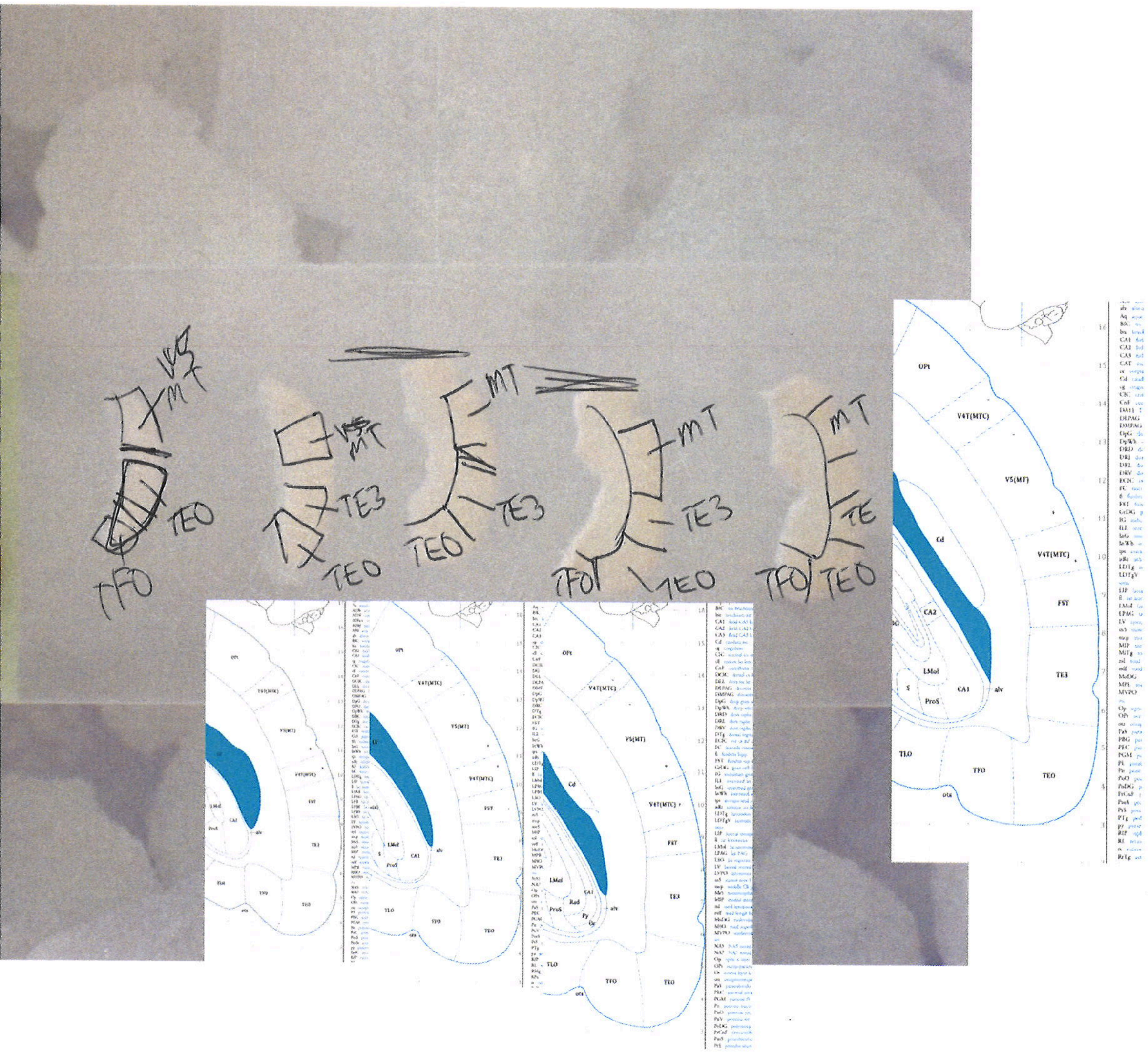

V5 MT
TEO
TFO
TE3
OPt
V4T(MTC)
V5(MT)
FST
TLO
Cd
CA1
CA2
LMol
ProS
S
alv
Rad
Py
Or
ots

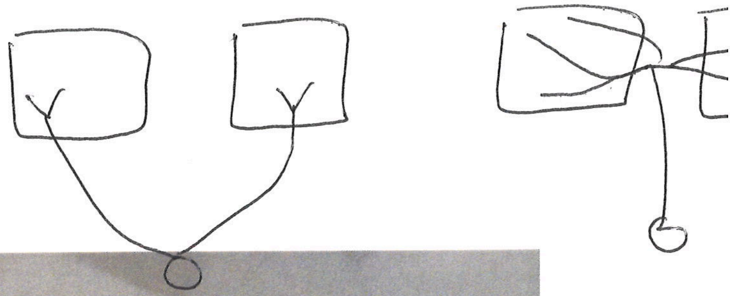

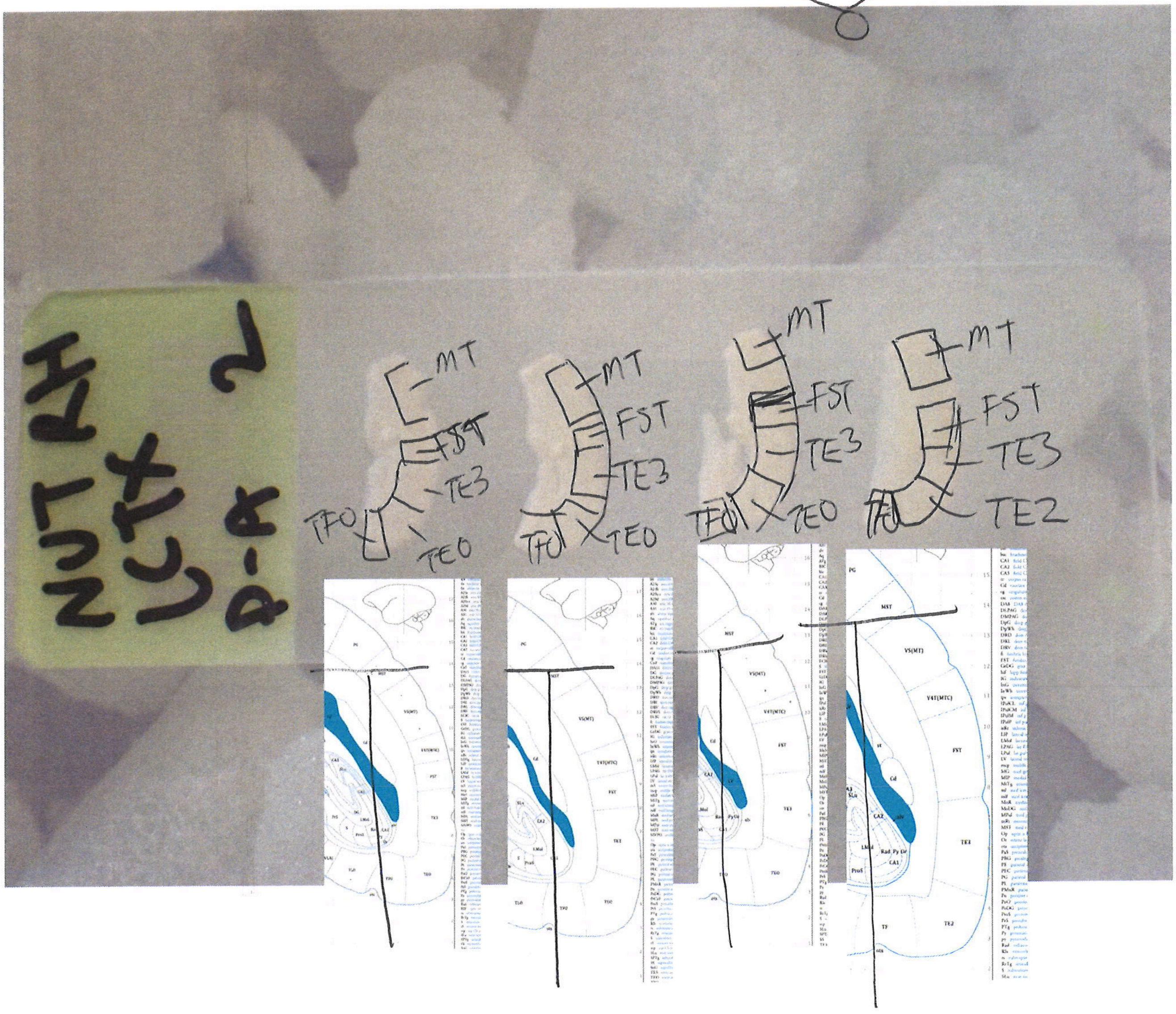
NUT RH
LCTX
P-A 2
MT
FST
TE3
TEO
TFO
TE2

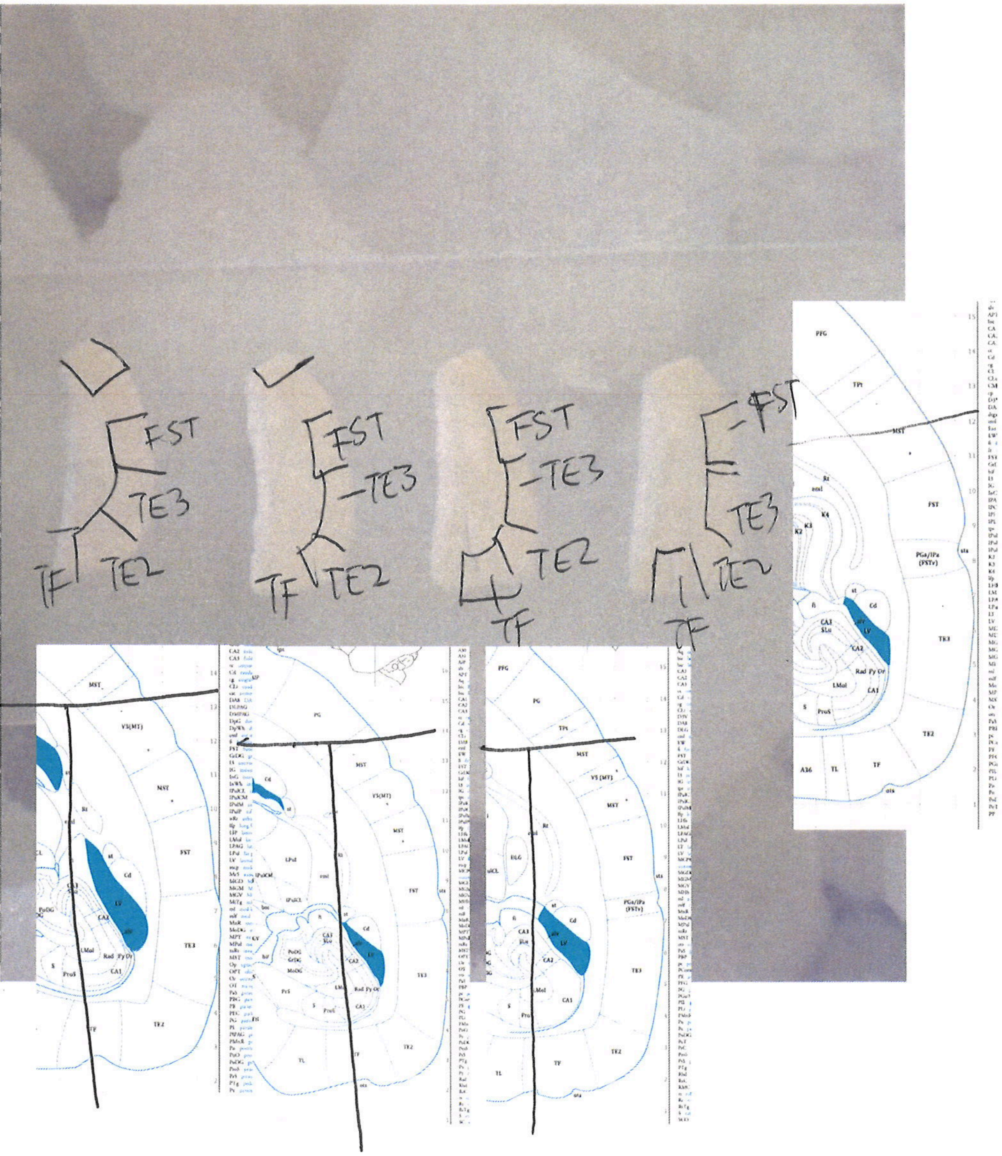
FST
TE3
TE2
TF
FST
TE3
TE2
TF
FST
TE3
TE2
TF
FST
TE3
TE2
TF
MST
V5(MT)
FST
TE3
TE2
PG
TPt
PFG
PGa/IPa (FSTv)
Cd
LV
CA1
CA2
CA3
A36
TL
TF

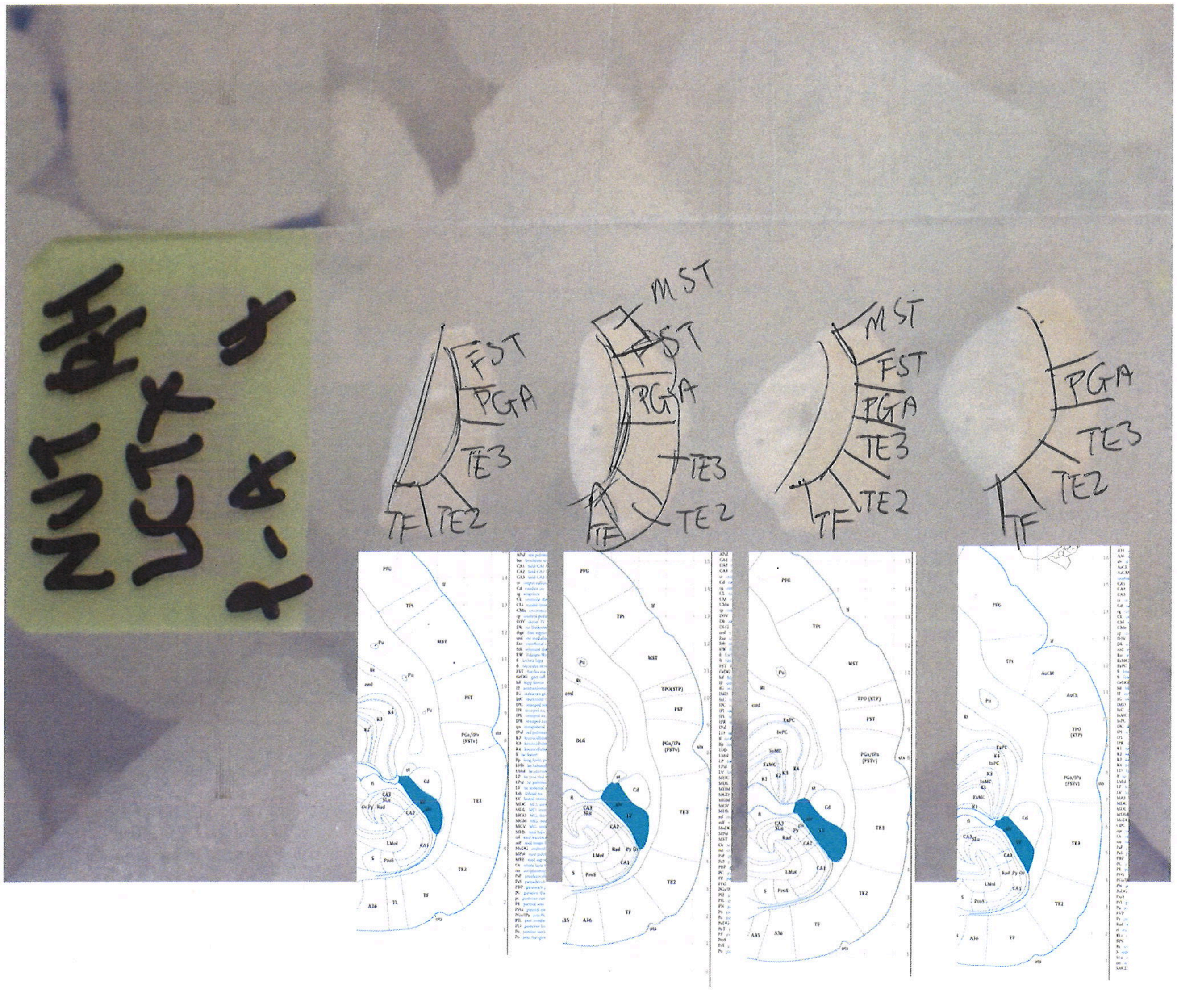
NUT RH
LCTX 4
1-4
FST
PGA
TE3
TE2
TF
MST
FST
PGA
TE3
TE2
TF
MST
FST
PGA
TE3
TE2
TF
PGA
TE3
TE2
TF

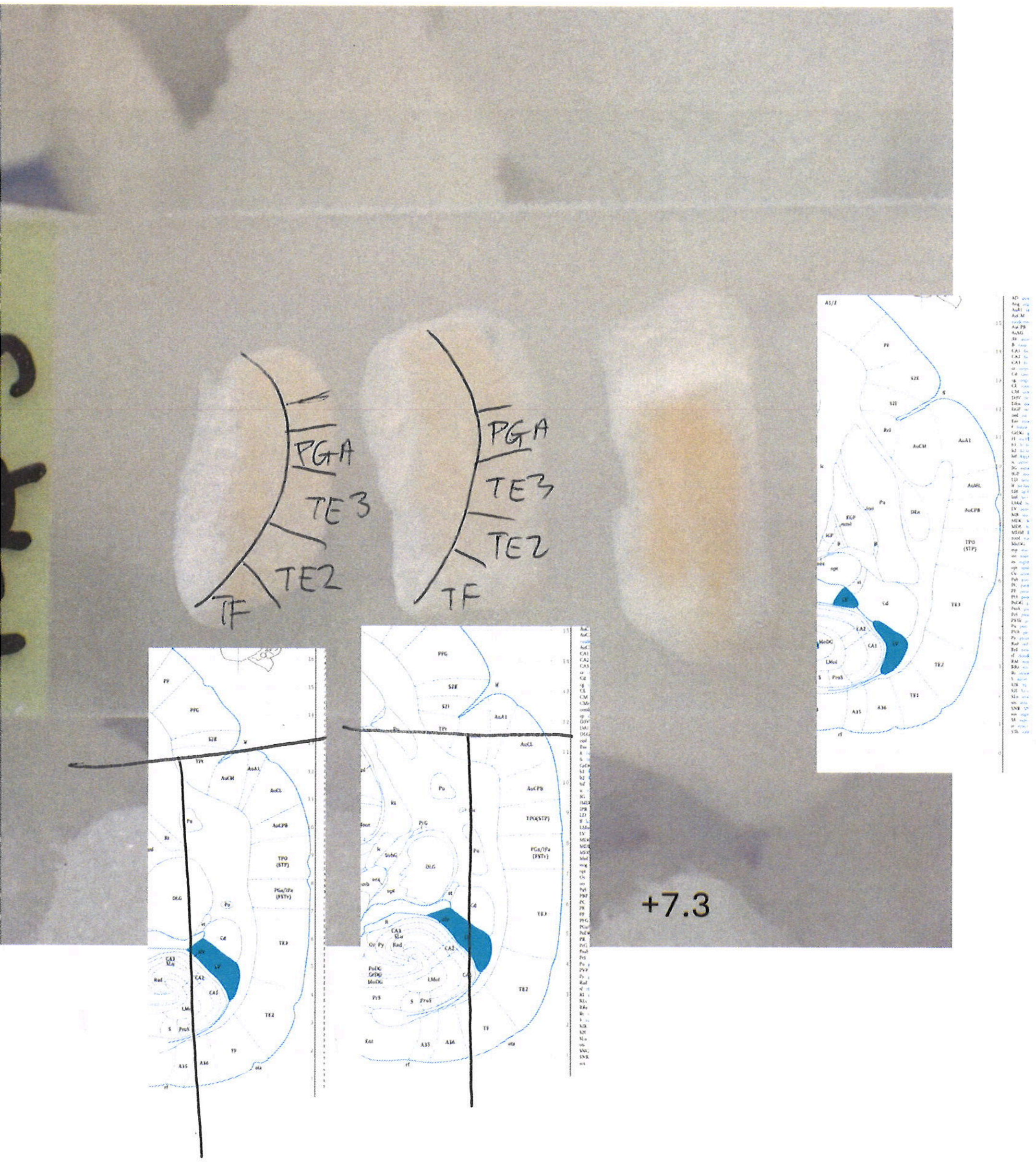
PGA
TE3
TE2
TF
PGA
TE3
TE2
TF
+7.3

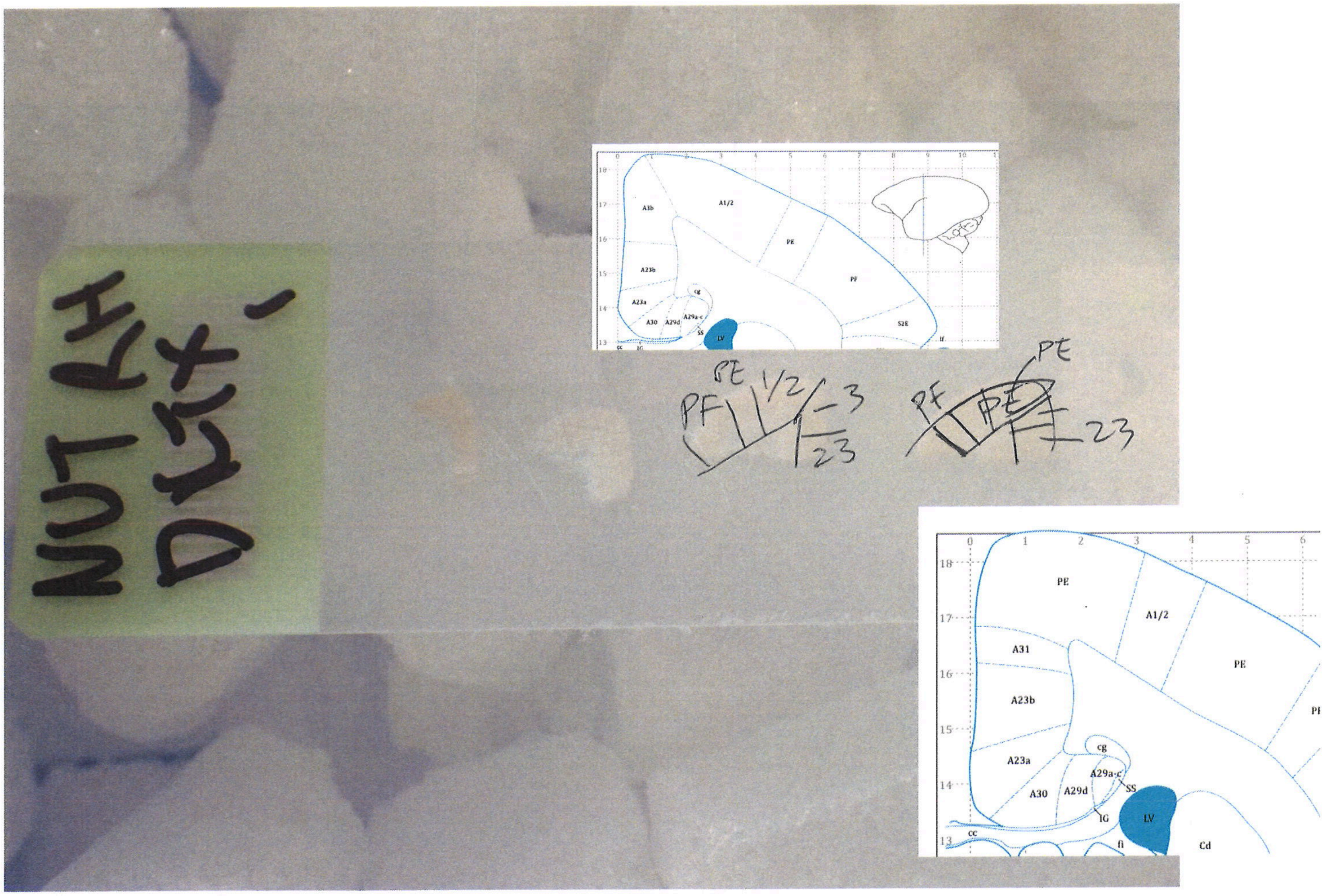
PE
A1/2
PF
A3b
A31
A23b
A23a
A30
A29d
A29a-c
SS
IG
LV
cg
cc
fi
Cd
S2E
1/2
23

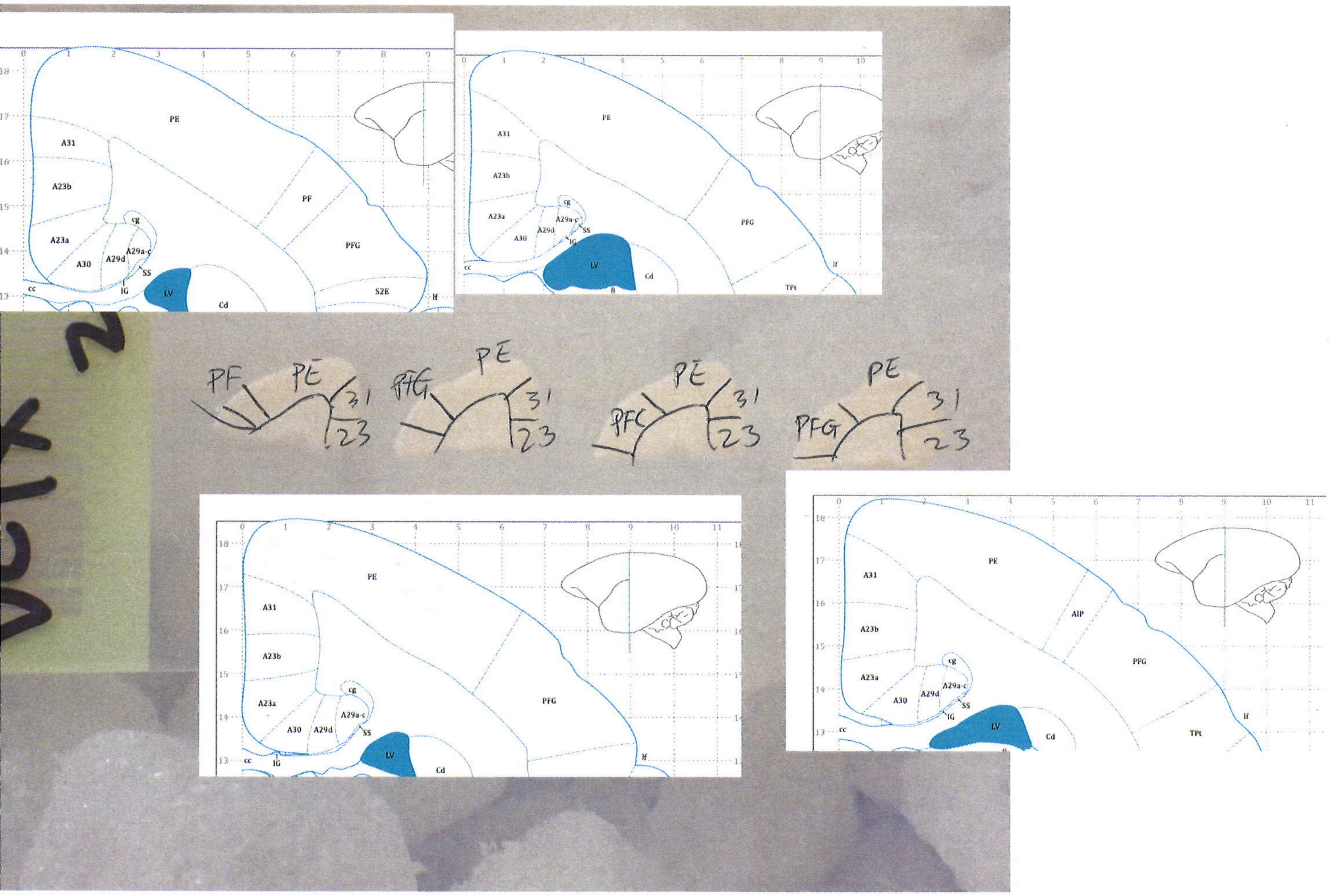
PE
A31
A23b
A23a
A30
A29d
A29a-c
cg
SS
IG
LV
Cd
cc
PF
PFG
S2E
AIP
TPt
lf
PF PE 31 23
PFG PE 31 23
PFG PE 31 23
PFG PE 31 23

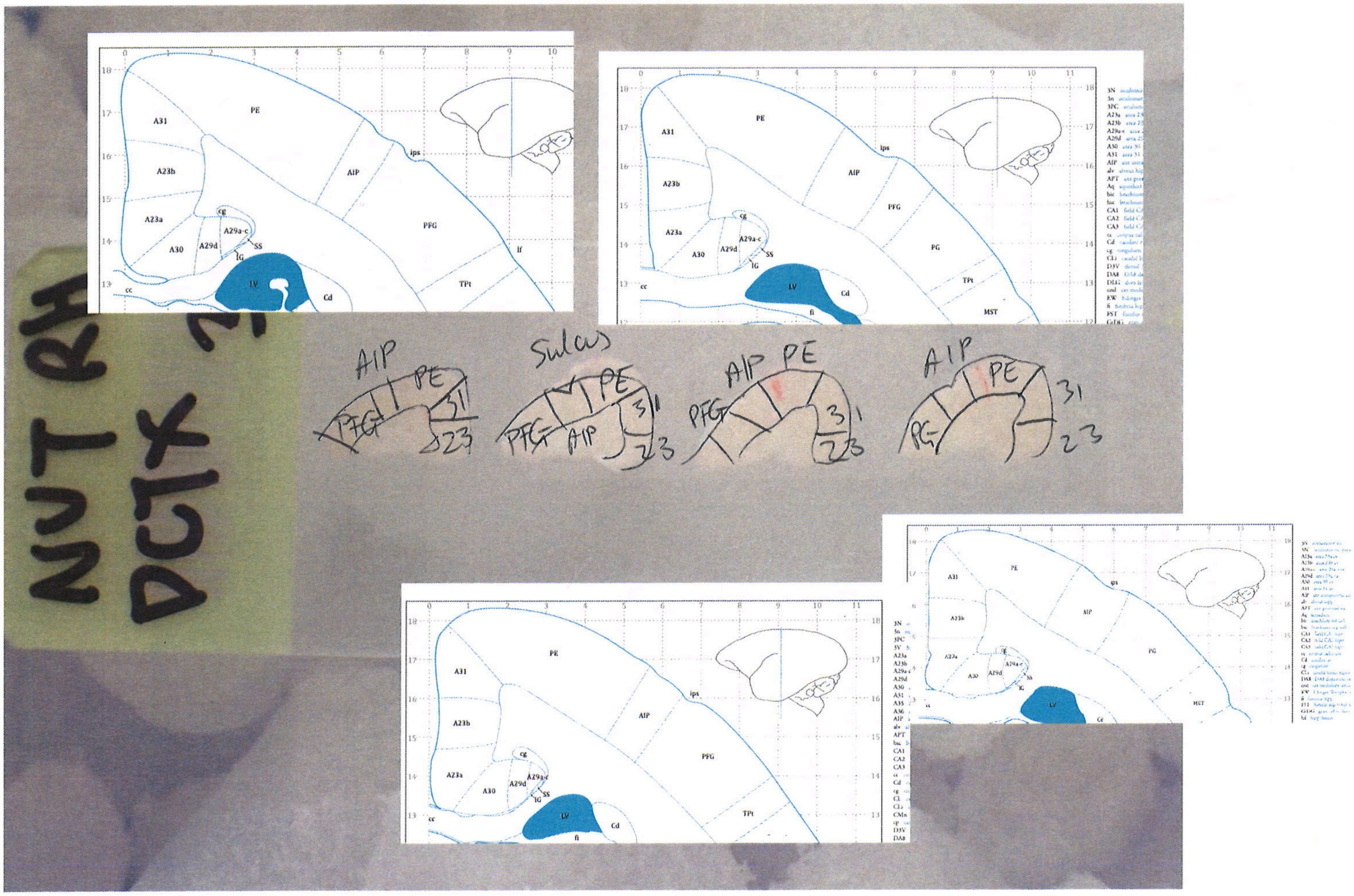
NUT RH
DCTX
AIP
PE
PFG
31
23
Sulcus
PG
PE
A31
A23b
A23a
A30
A29d
A29a-c
SS
IG
cg
LV
Cd
cc
AIP
ips
PFG
PG
TPt
MST
lf
fi

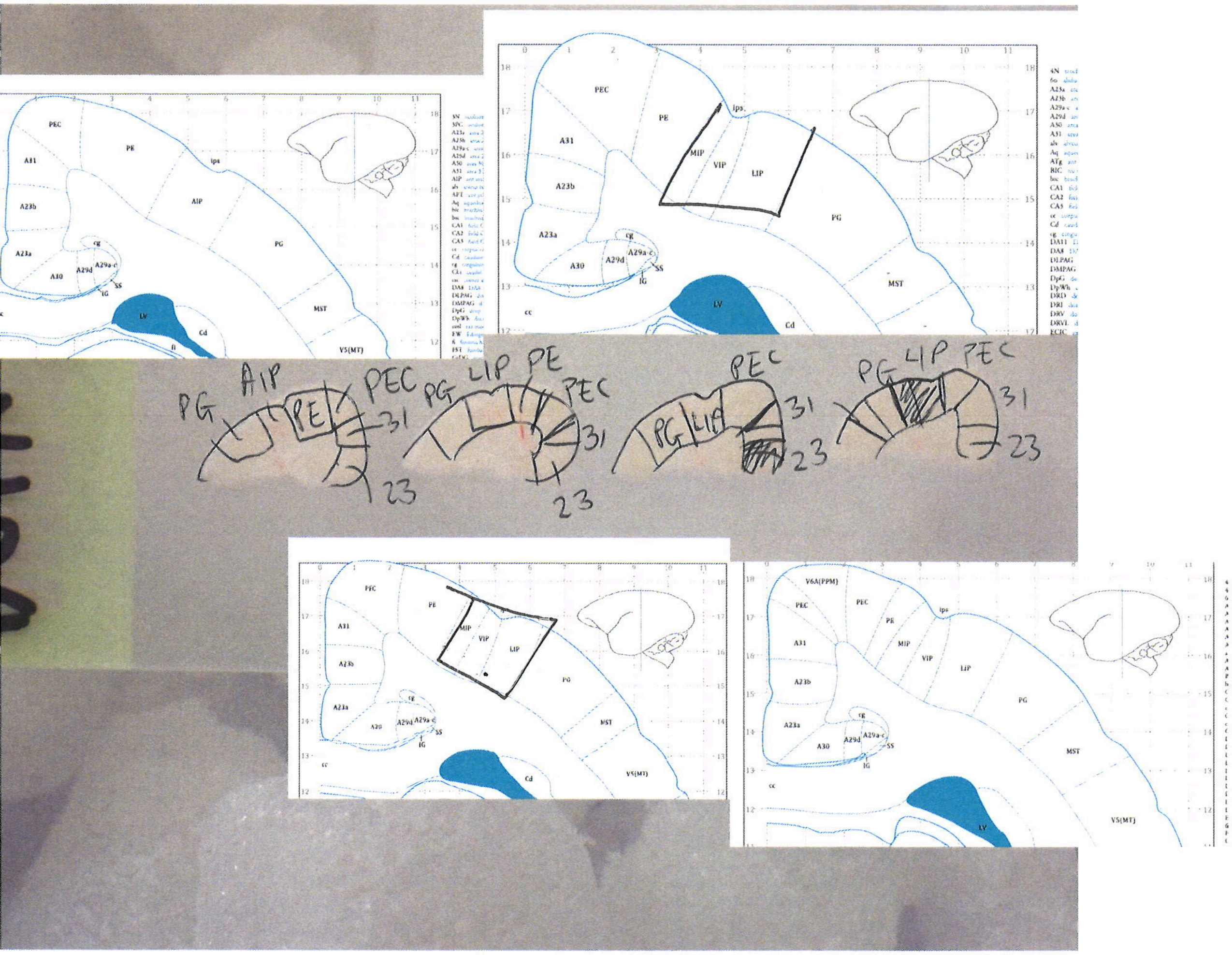

PEC
PE
A31
ips
A23b
AIP
A23a
PG
cg
A30
A29d
A29a-c
SS
IG
LV
MST
Cd
V5(MT)
MIP
VIP
LIP
cc
PG
AIP
PE
PEC
31
23
PG
LIP
PE
PEC
31
23
PEC
PG
LIP
31
23
PG
LIP
PEC
31
23
V6A(PPM)

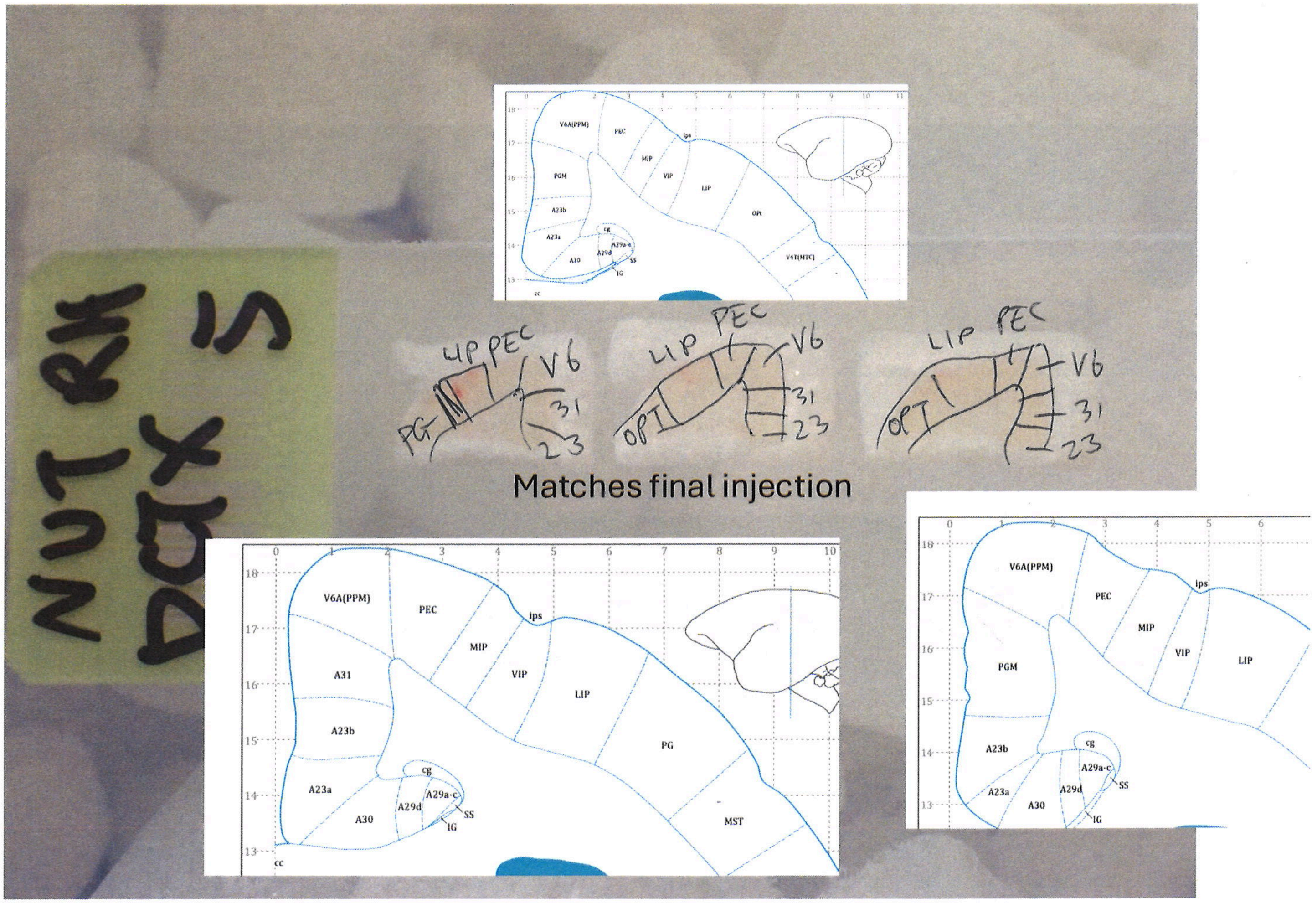
NUT RH
DAX 5
LIP PEC V6 PG 31 23
LIP PEC V6 OPT 31 23
LIP PEC V6 OPT 31 23
Matches final injection
V6A(PPM)
PEC
ips
MIP
VIP
LIP
PG
MST
A31
A23b
A23a
A30
A29d
A29a-c
cg
SS
IG
cc
PGM
OPt
V4T(MTC)

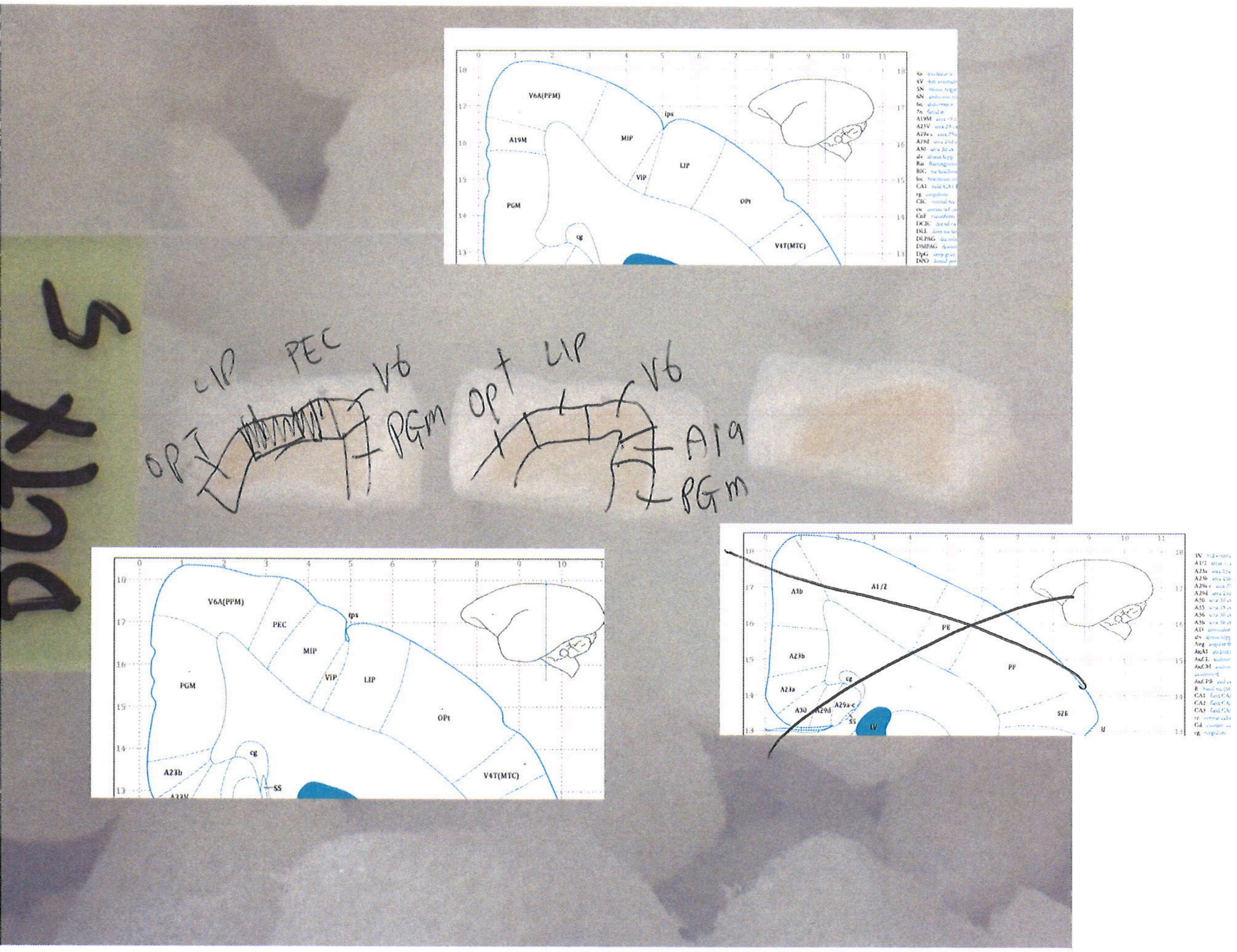
DCTX 5
LIP
PEC
V6
OPT
PGM
OPT
LIP
V6
A19
PGM
V6A(PPM)
ips
A19M
MIP
LIP
VIP
PGM
OPt
cg
V4T(MTC)
V6A(PPM)
PEC
ips
MIP
VIP
LIP
PGM
OPt
cg
A23b
V4T(MTC)
A3b
A1/2
PE
A23b
PF
A23a
cg
A30
A29a-c
LV
S2E

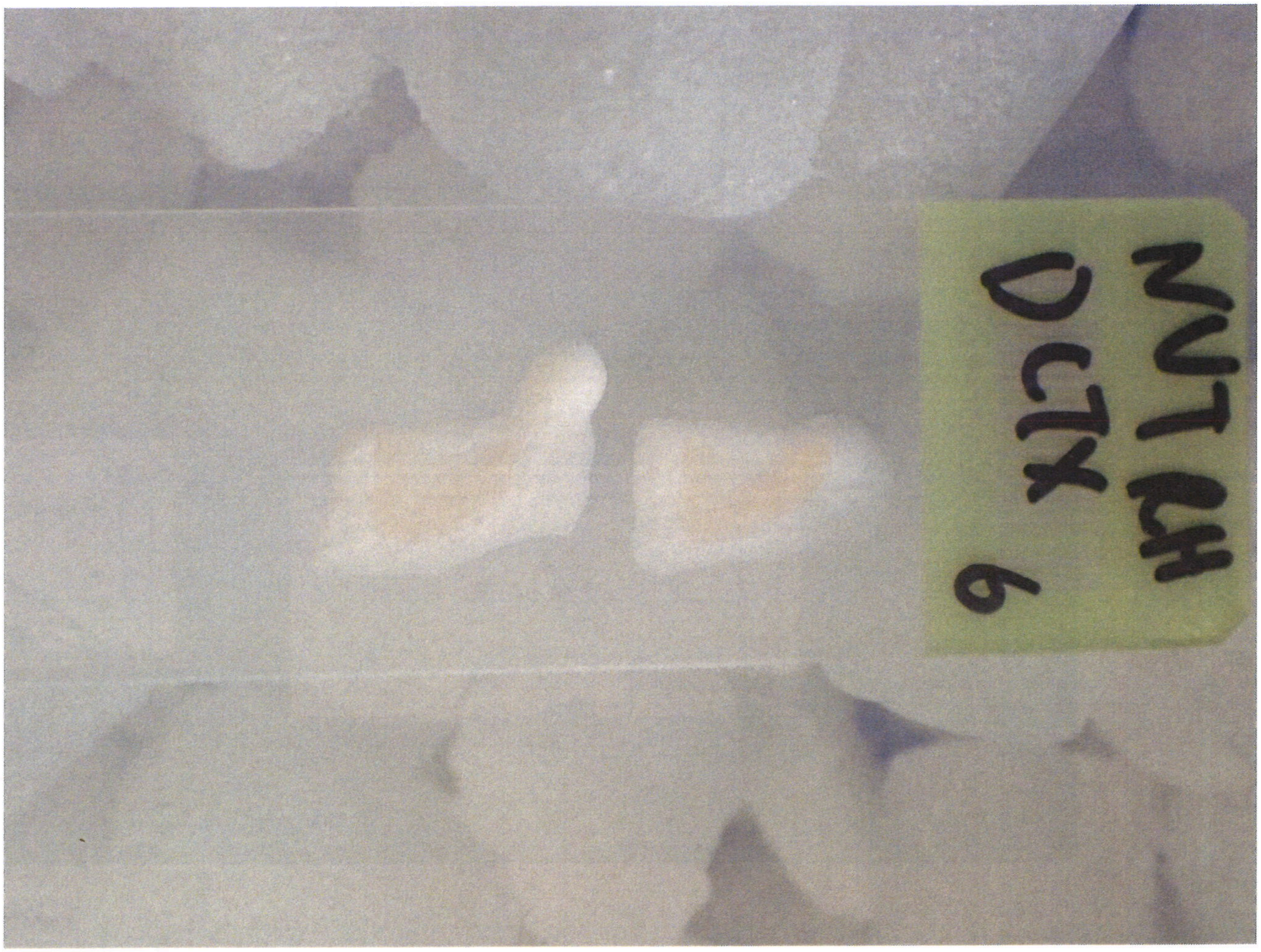
NUTCH
DCTX 6

HEMISPHERE

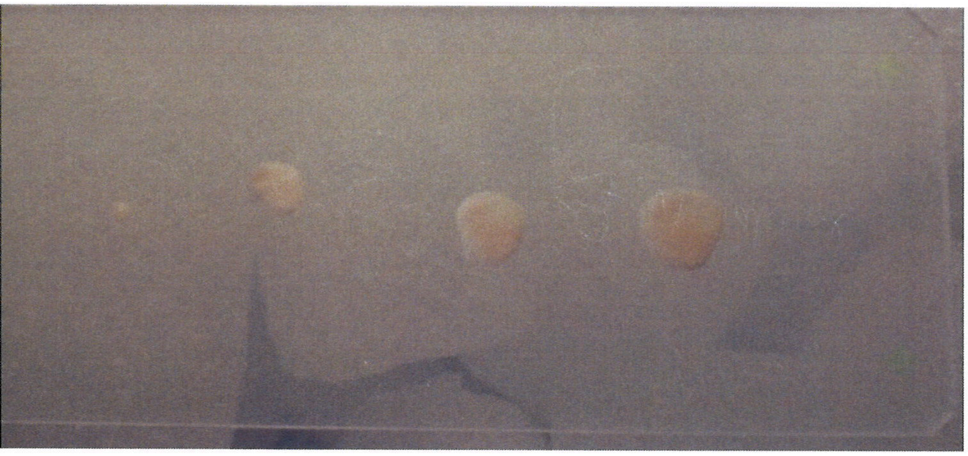

ALL V1

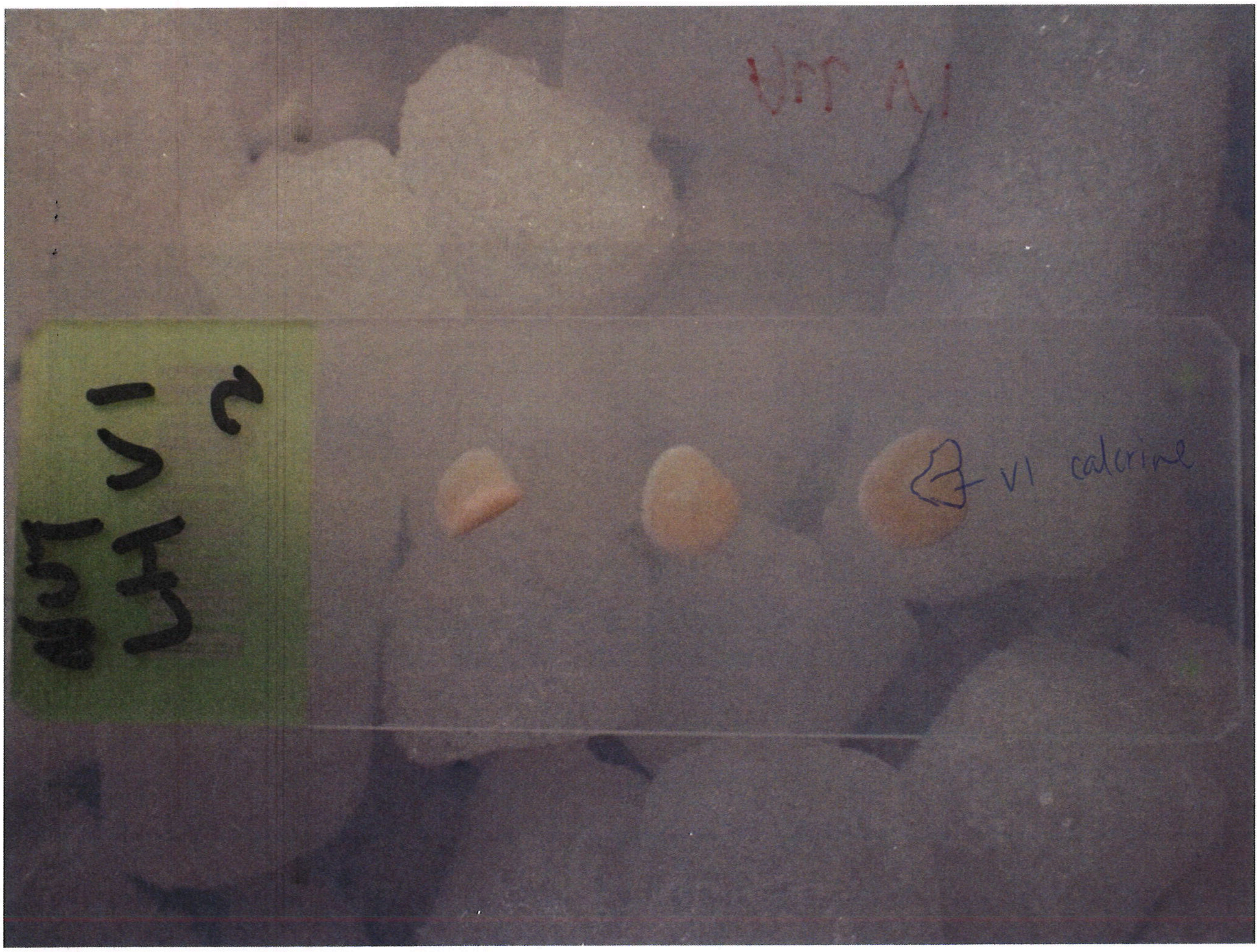


All V1

LEFT HEMISPHERE

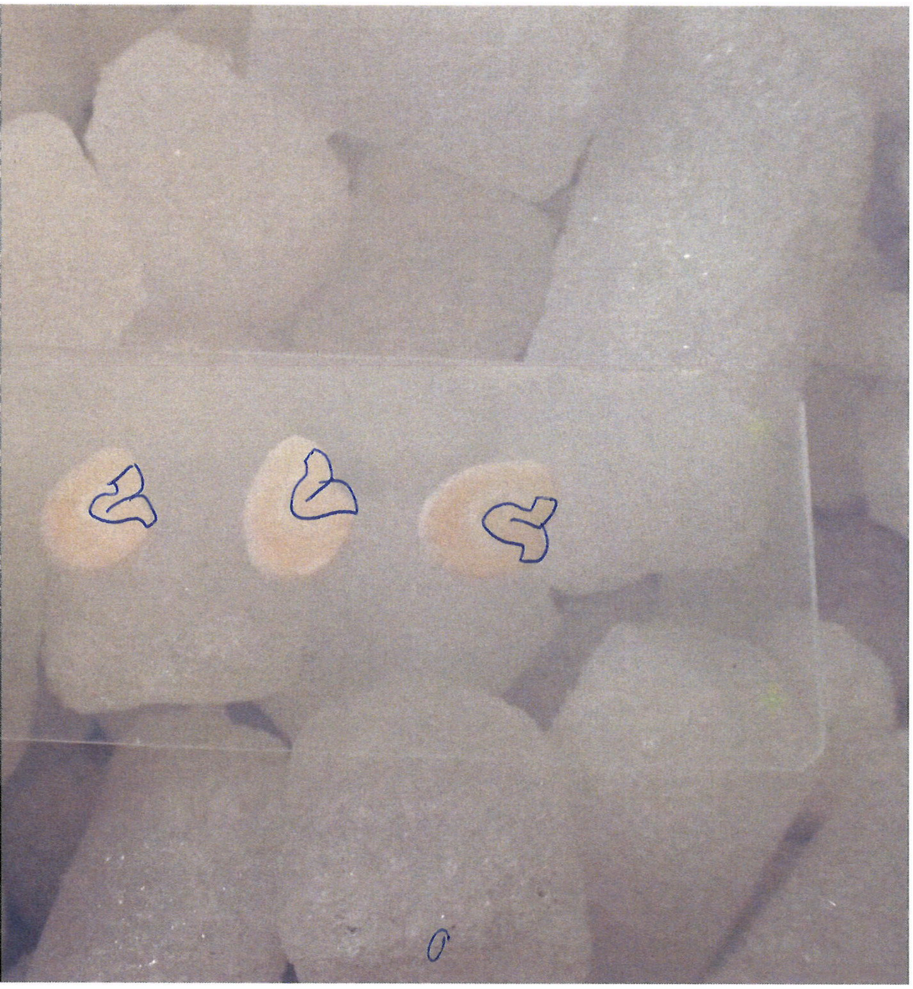

all VI

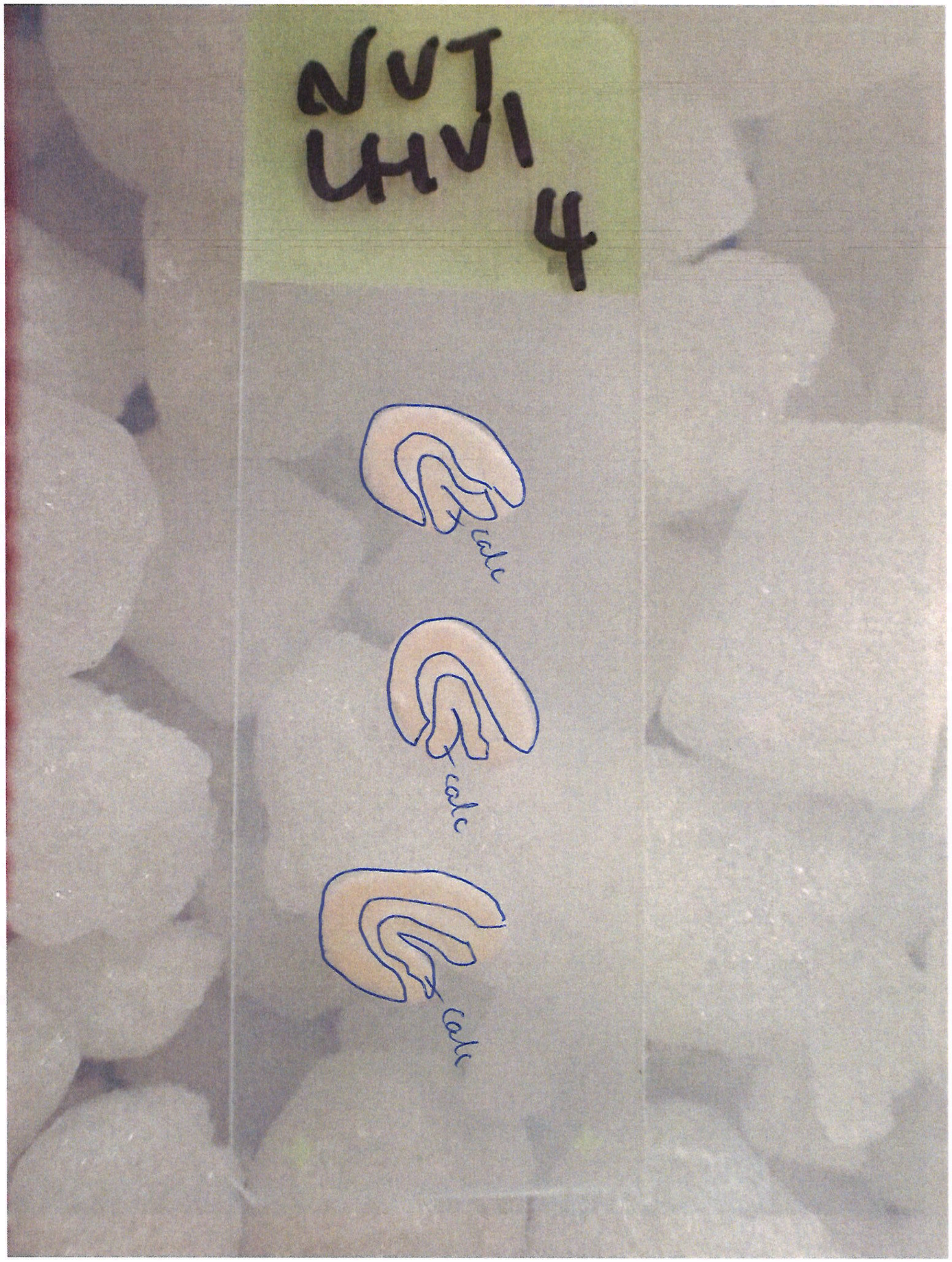
NVT
LHVI
4
calc
calc
calc

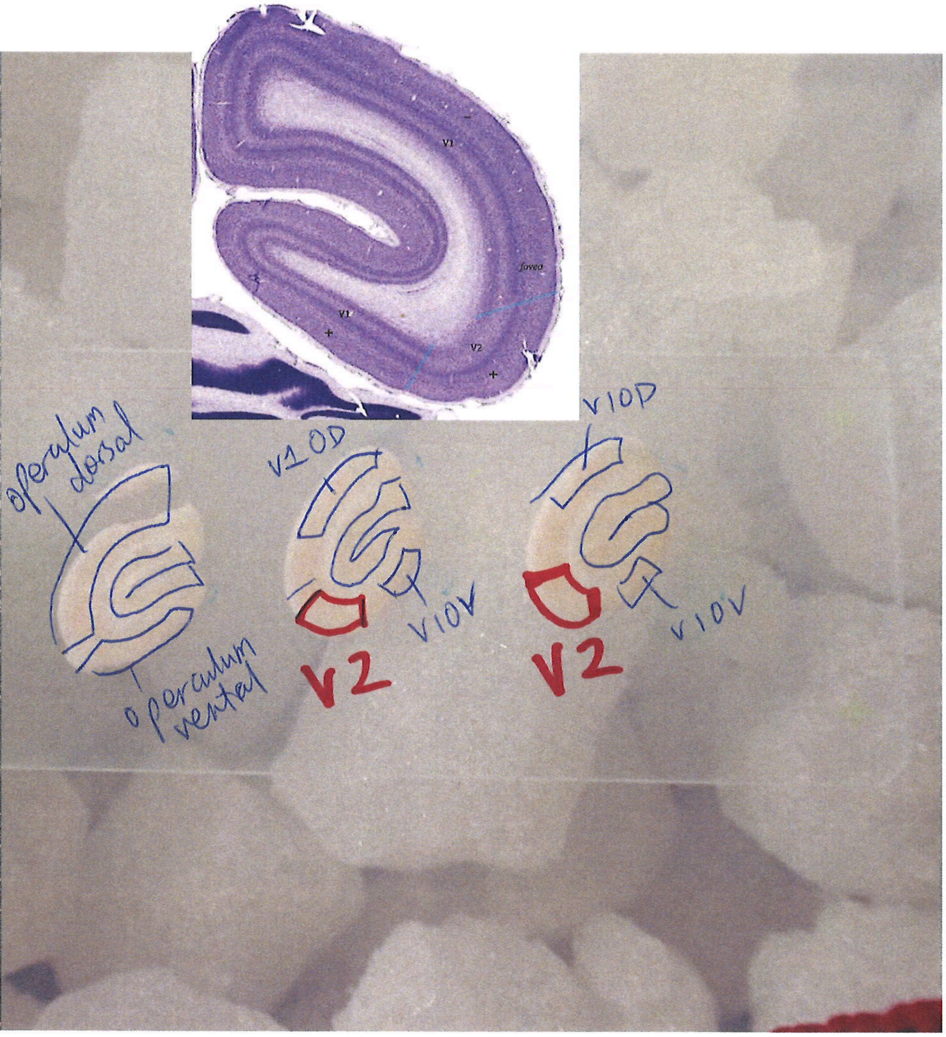
V1
fovea
V1
V2
operculum dorsal
operculum ventral
V1 OD
V1 OV
V2
V1 OD
V1 OV
V2

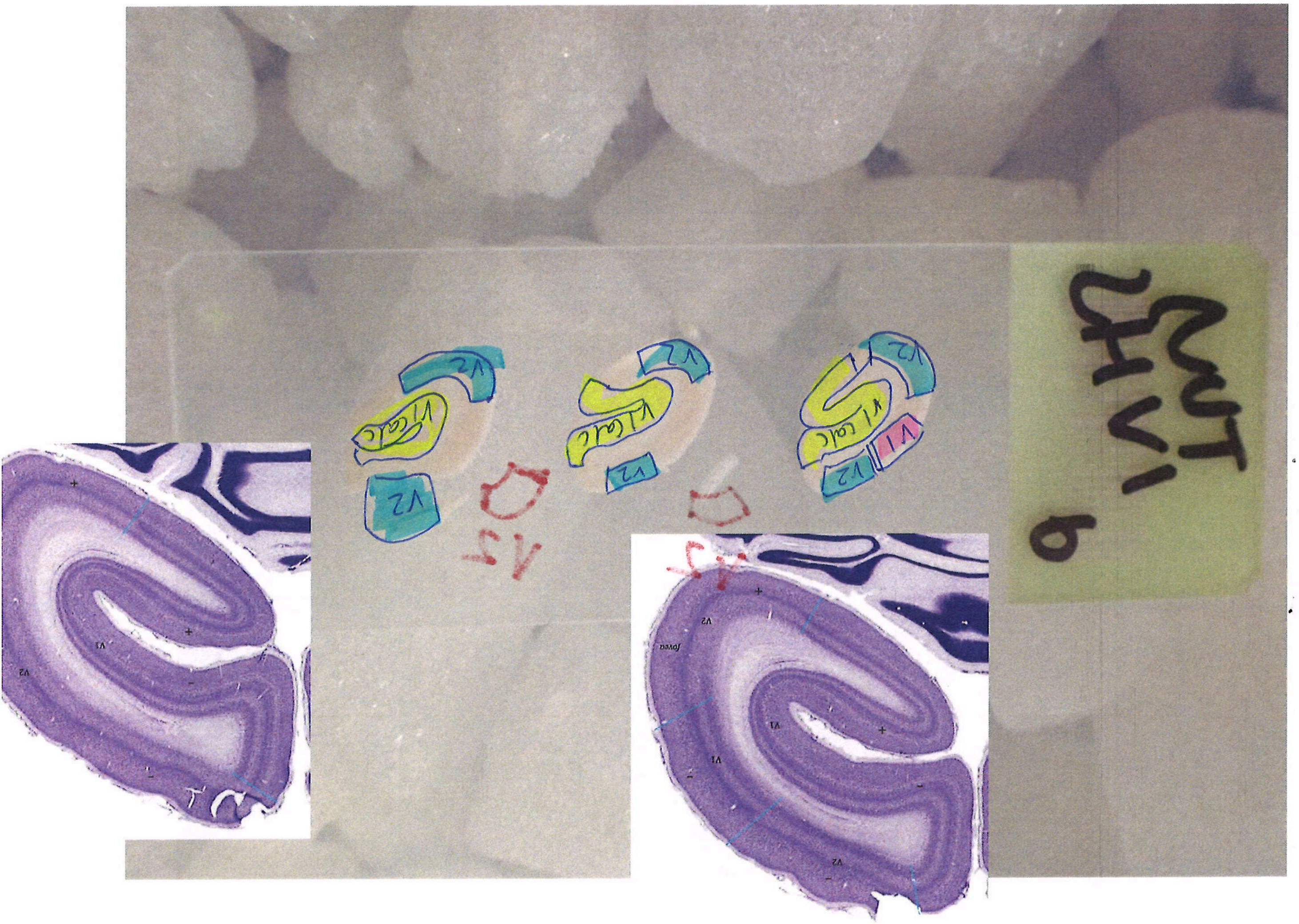
CUT
LHV1
6
V1 calc
V2
V1

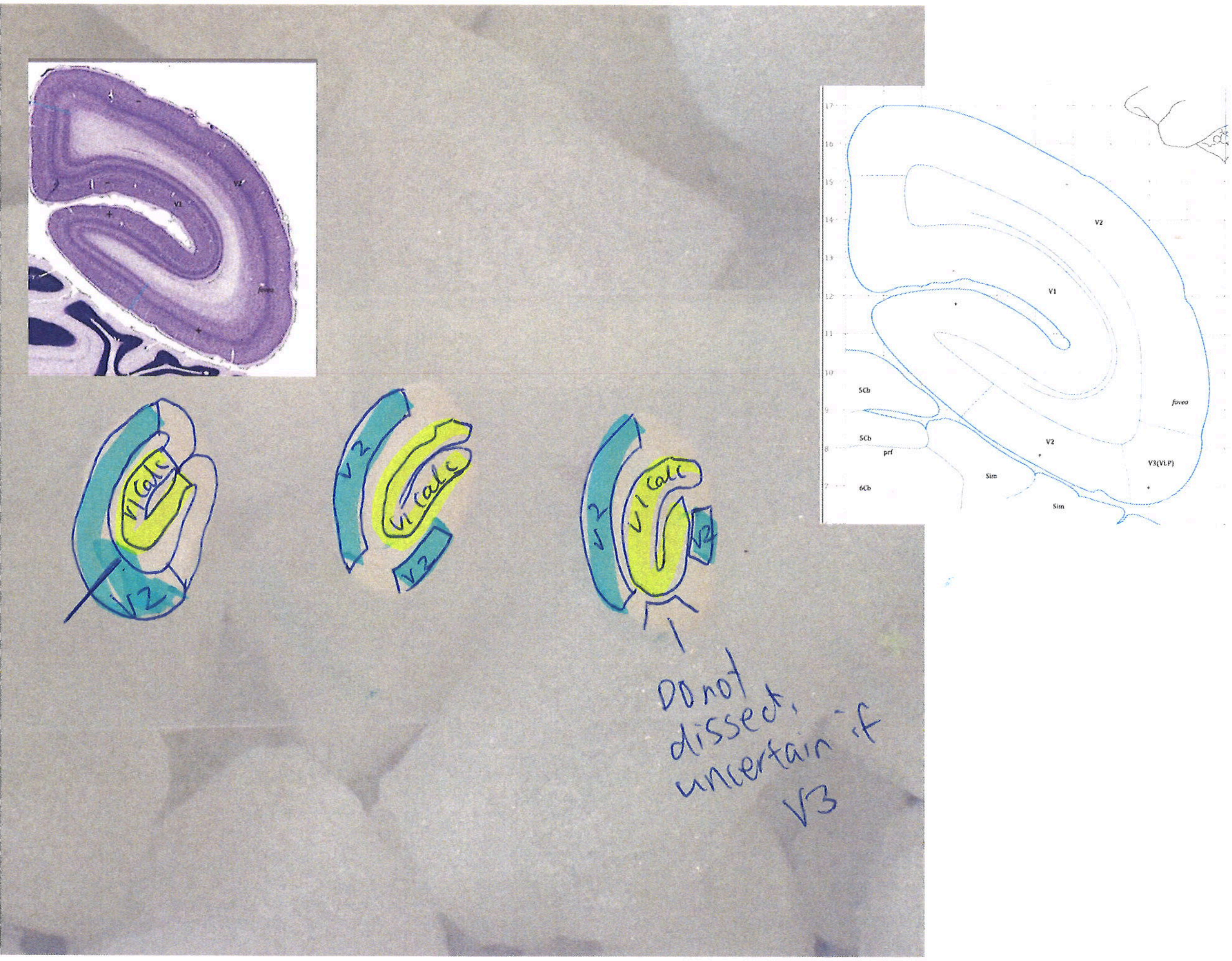
V1 calc
V2
V1 calc
V2
V2
V1 calc
V2
V2
Do not dissect, uncertain if V3
V2
V1
fovea
V2
V3(VLP)
5Cb
5Cb
prf
6Cb
Sim
Sim

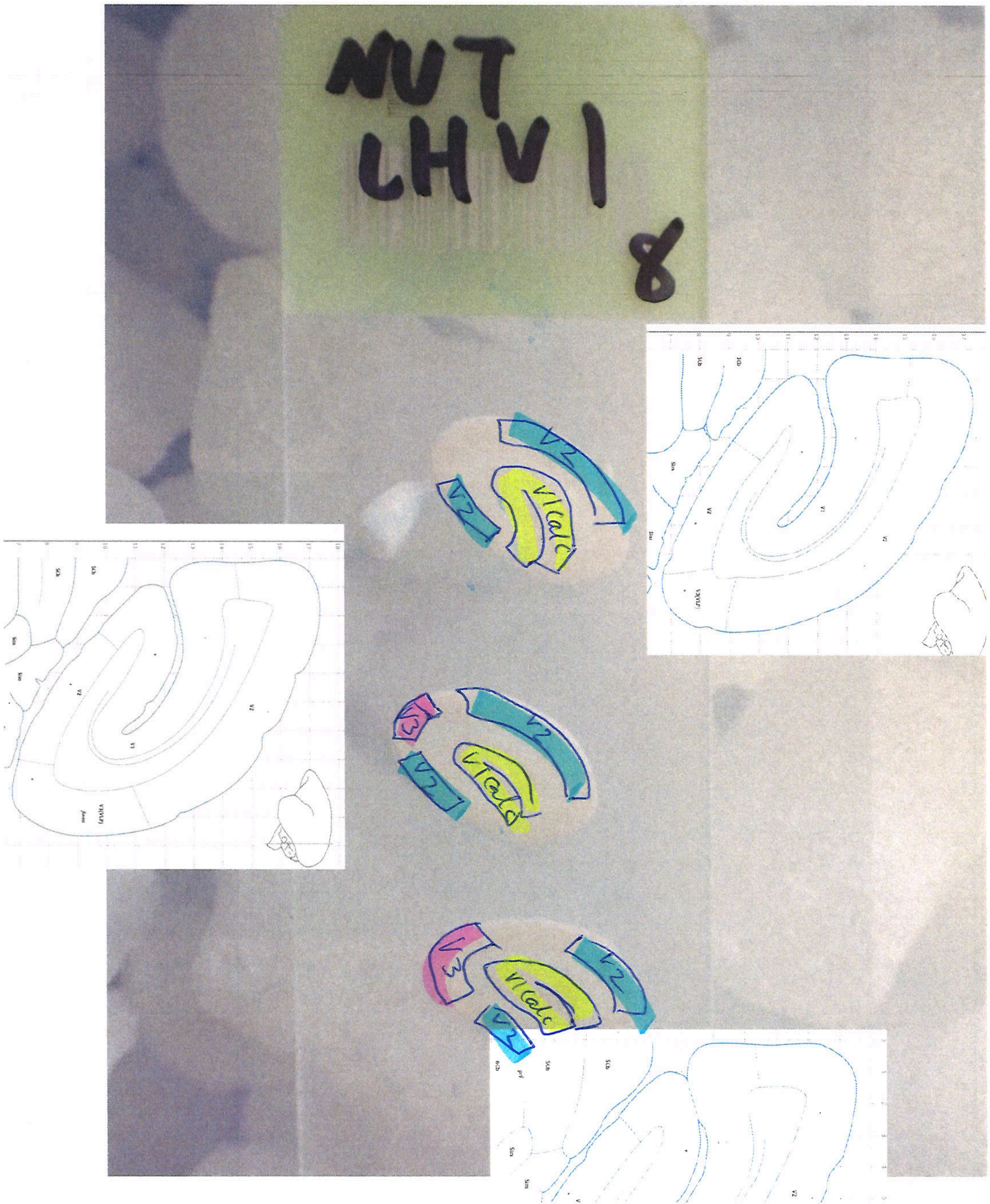
MUT
LHV1
8
V2
V1calc
V3

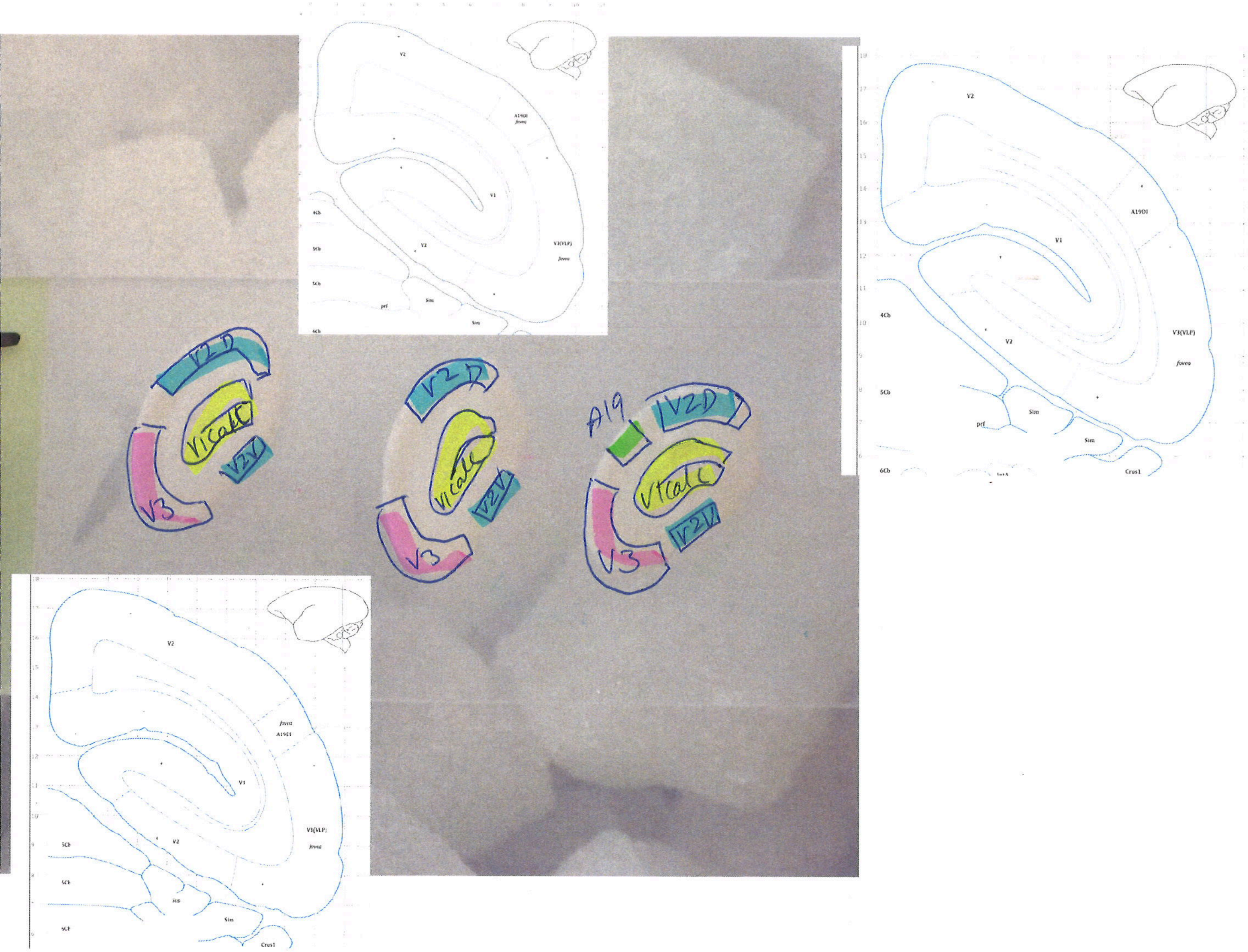

V2D
V1calc
V2V
V3
A19
V2
V1
A19DI
V3(VLP)
fovea
Sim
Crus1

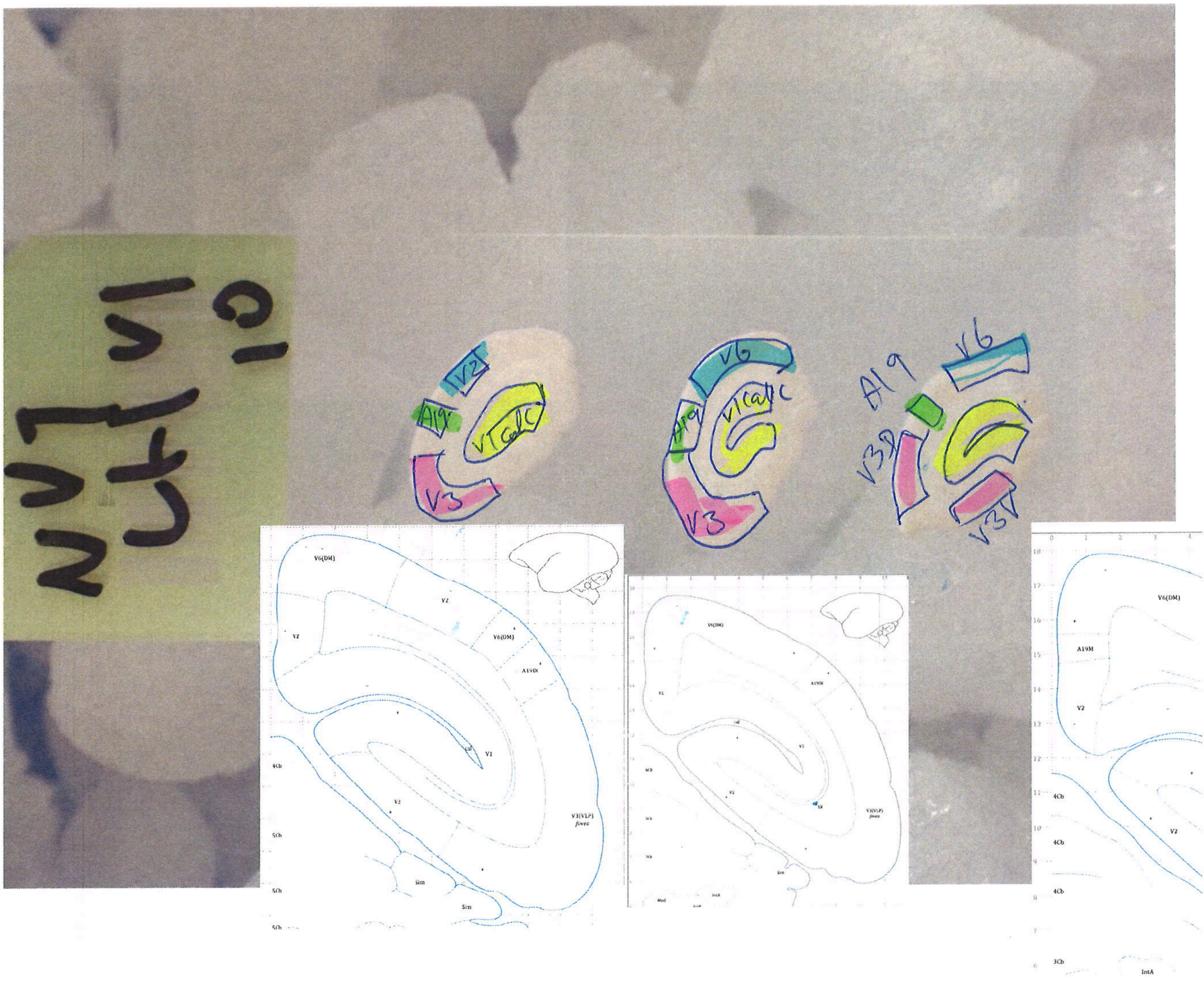
V3
A19
V2
V1 (calc)
V6
V3D
V3V
V6(DM)
A19VDi
A19M
V1
V2
V3(VLP)
fovea
4Cb
5Cb
Sim
IntA

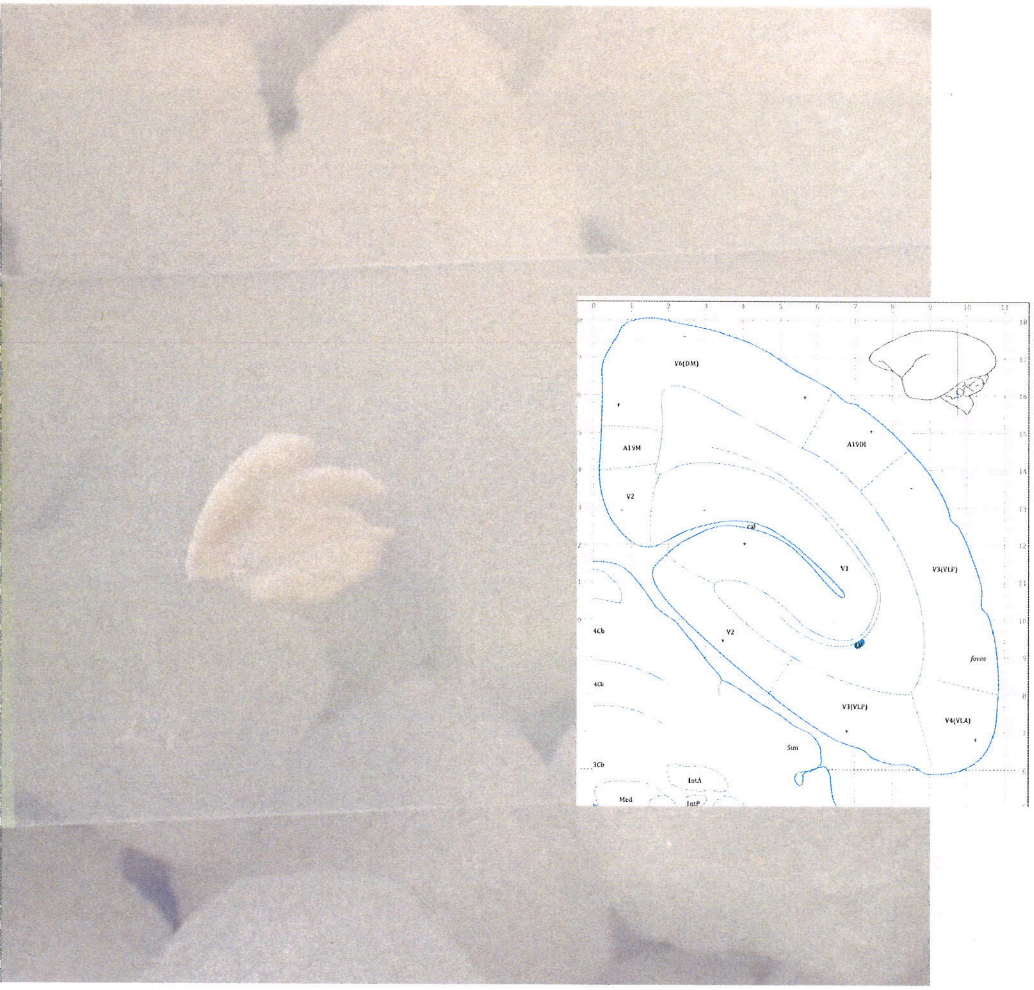
V6(DM)
A19M
A19DI
V2
cal
V1
V3(VLF)
4Cb
V2
fovea
4Cb
V3(VLP)
V4(VLA)
Sim
3Cb
IntA
Med
IntP

All area 10

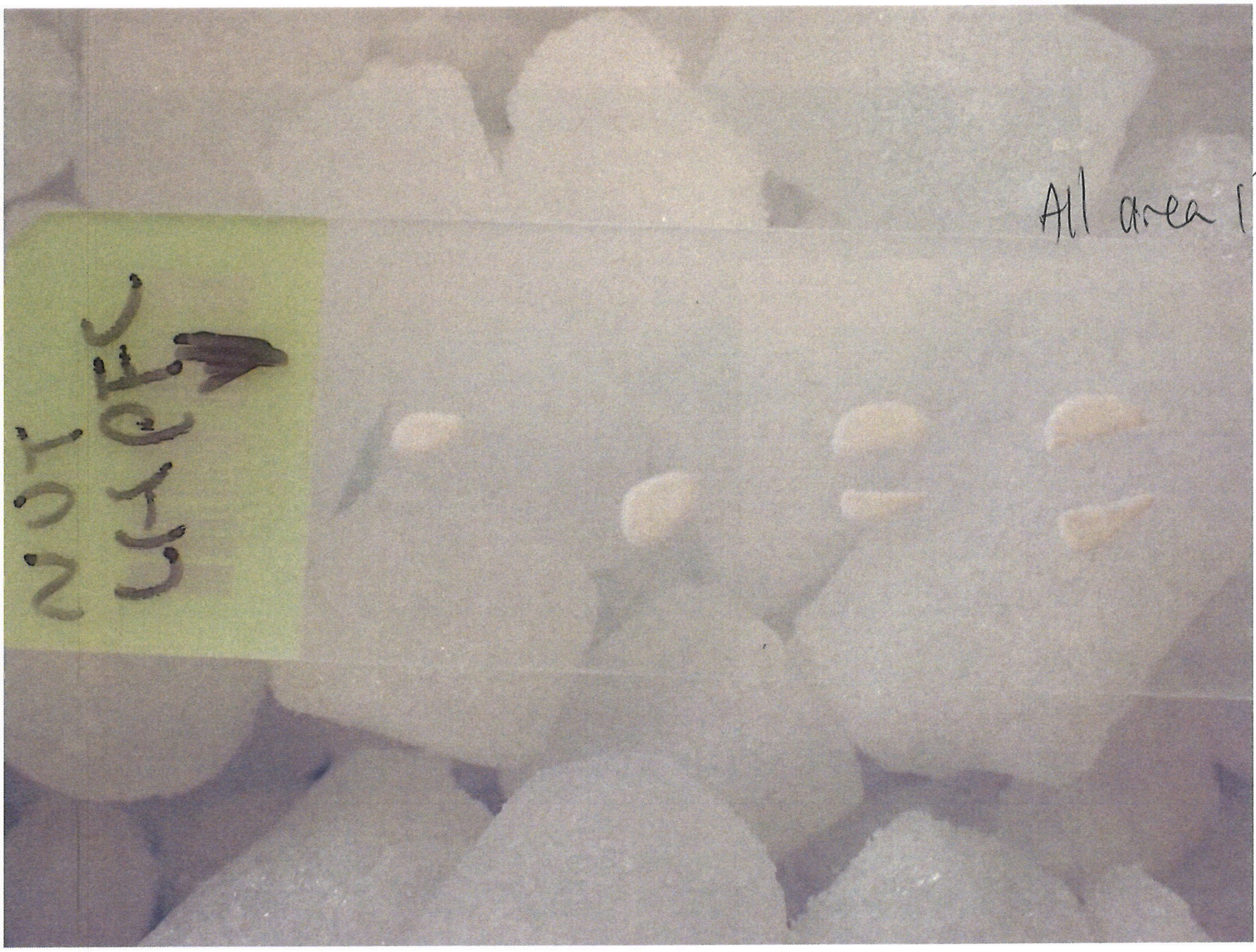

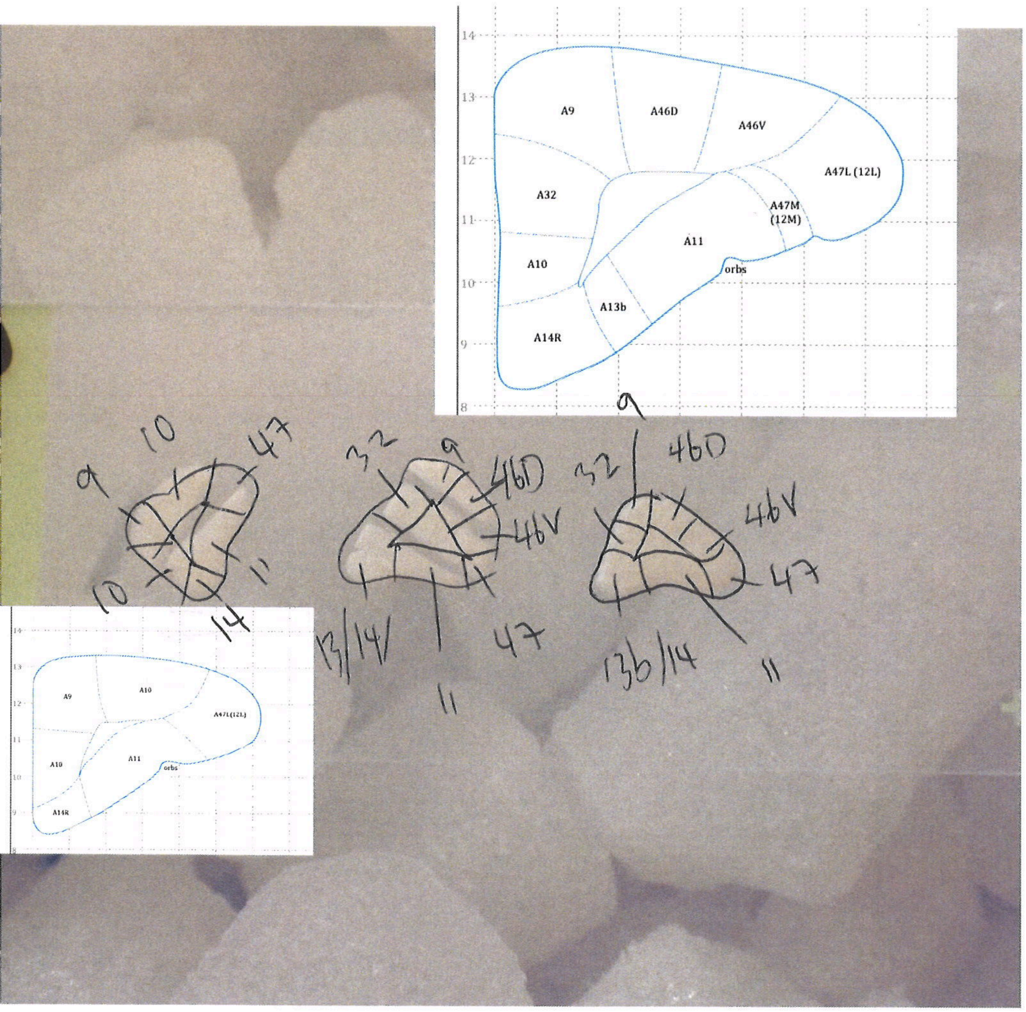

14
13
12
11
10
9
8
A9
A46D
A46V
A32
A47L (12L)
A47M
(12M)
A11
A10
orbs
A13b
A14R
9
10
47
10
11
14
32
9
46D
46V
13/14
11
47
9
32
46D
46V
47
13b/14
11
A9
A10
A47L(12L)
A10
A11
orbs
A14R

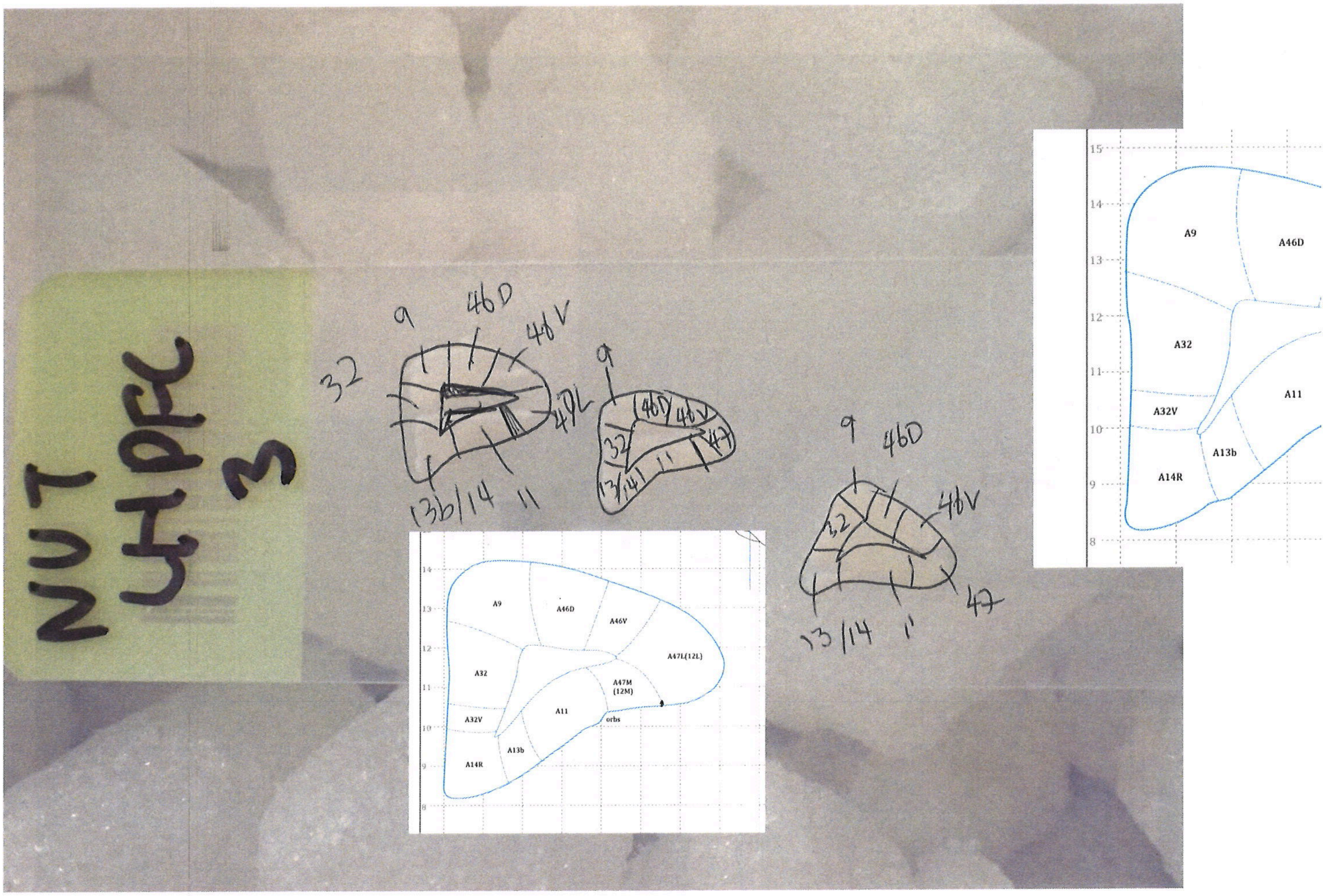
NUT
LH PFC
3
9
46D
46V
32
47L
13b/14
11
47
13/14
A9
A46D
A46V
A47L(12L)
A32
A47M
(12M)
A11
A32V
orbs
A14R
A13b

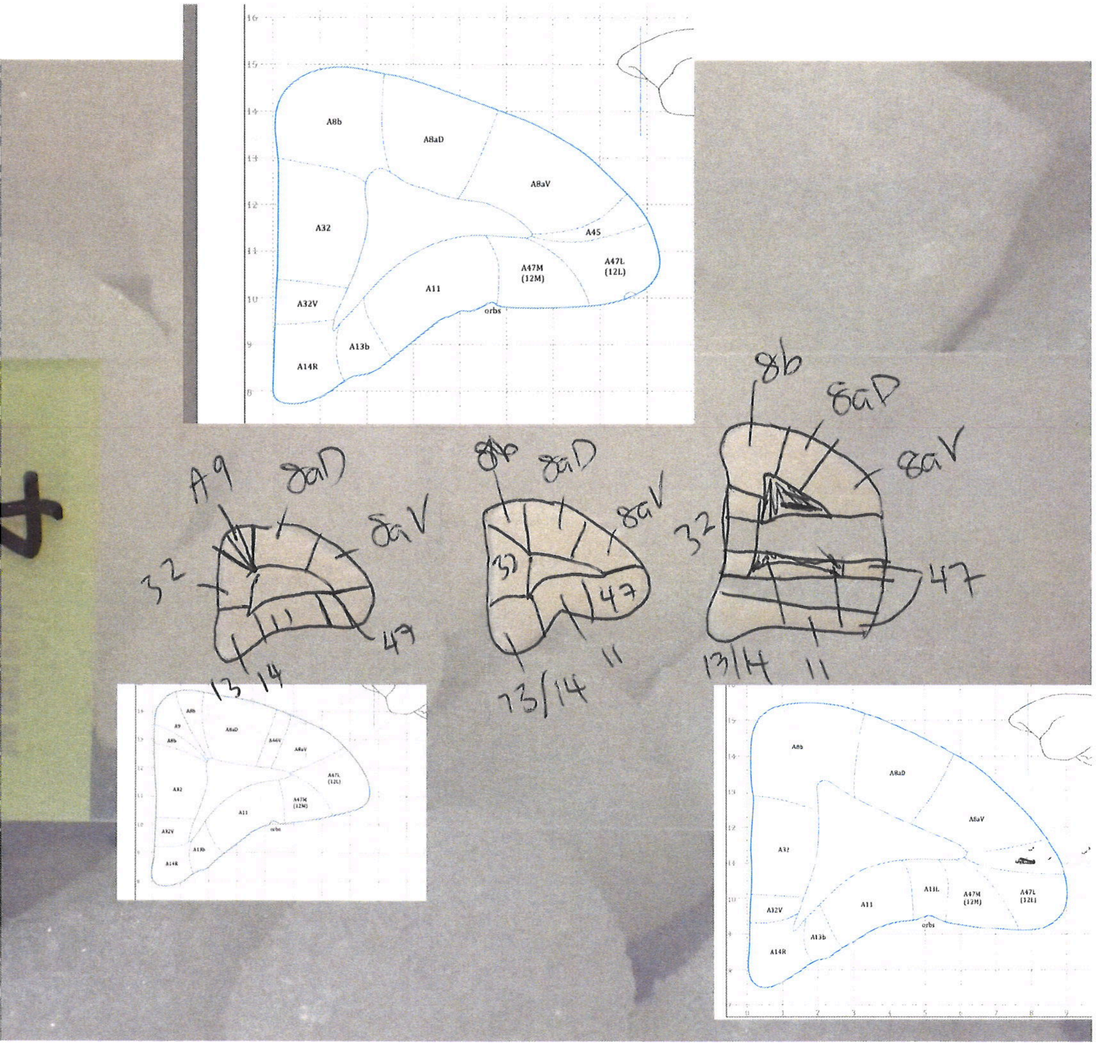
A8b
A8aD
A8aV
A45
A32
A47M (12M)
A47L (12L)
A11
A32V
orbs
A13b
A14R
A9
8aD
8aV
32
47
13
14
11
8b
13/14
A11L

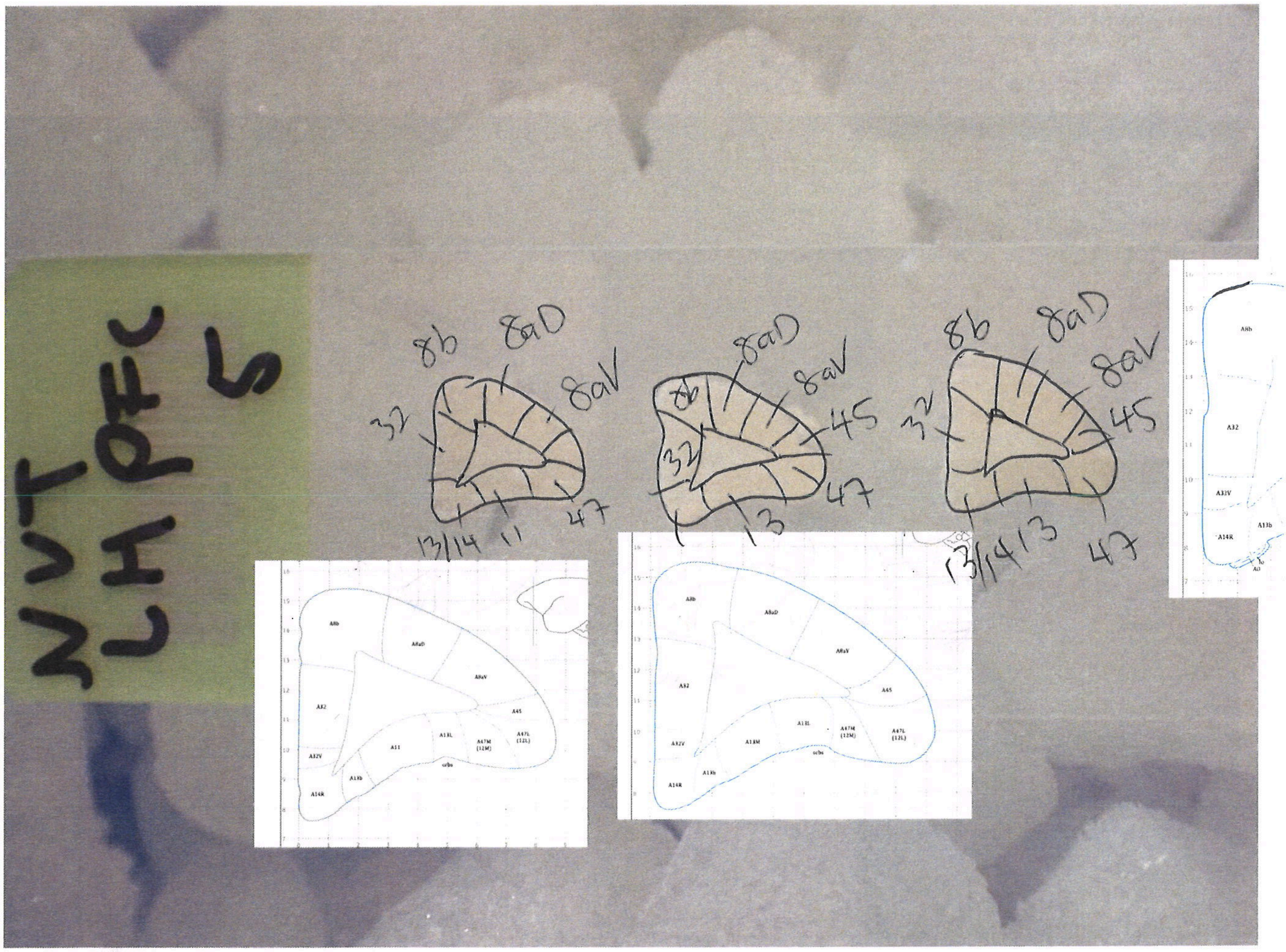
NVT
LH PFC
5
8b
8aD
8aV
32
13/14
11
47
45
13
A8b
A8aD
A8aV
A45
A32
A32V
A14R
A13b
A11
A13L
A13M
A47M
(12M)
A47L
(12L)
orbs

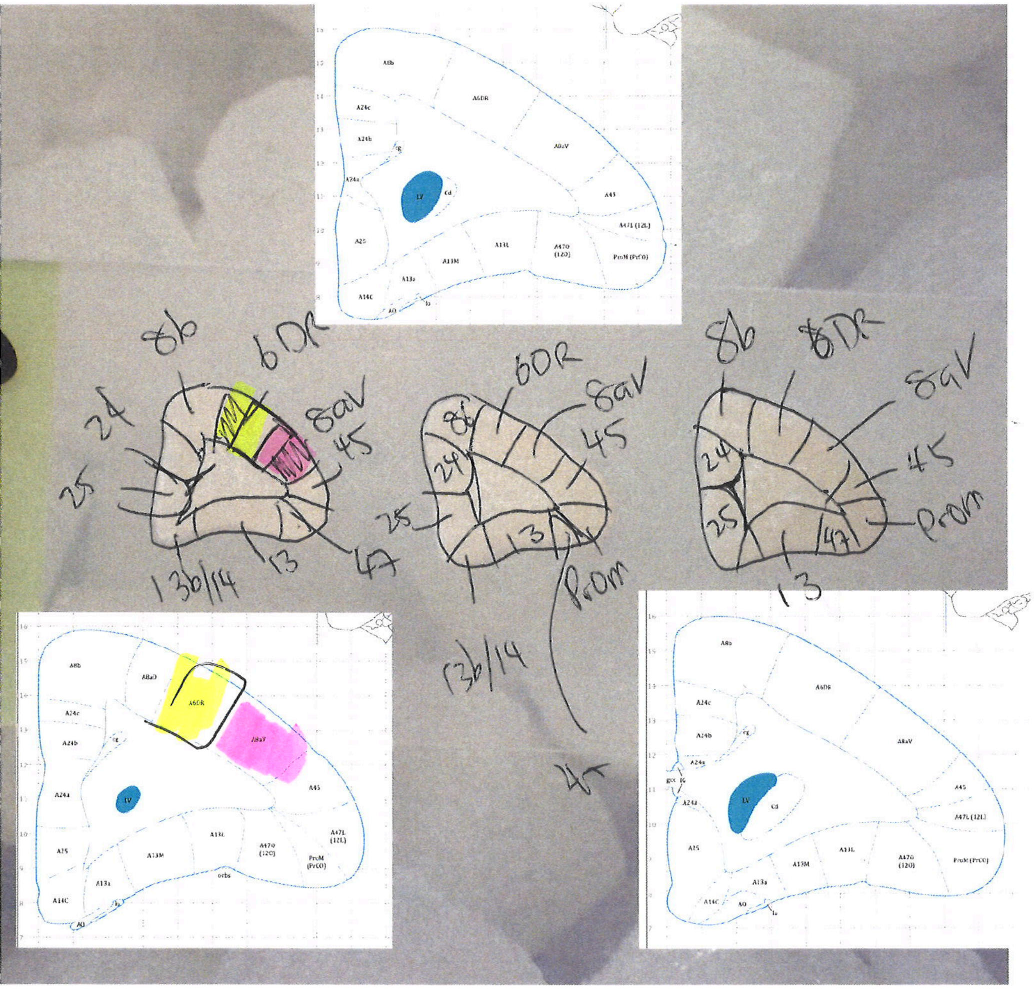
8b
6DR
24
8aV
45
25
13b/14
13
47
Prom
A8b
A6DR
A24c
A24b
A24a
A8aV
A45
A47L (12L)
A25
A13M
A13L
A47O (12O)
ProM (PrCO)
A13a
A14C
LV
Cd

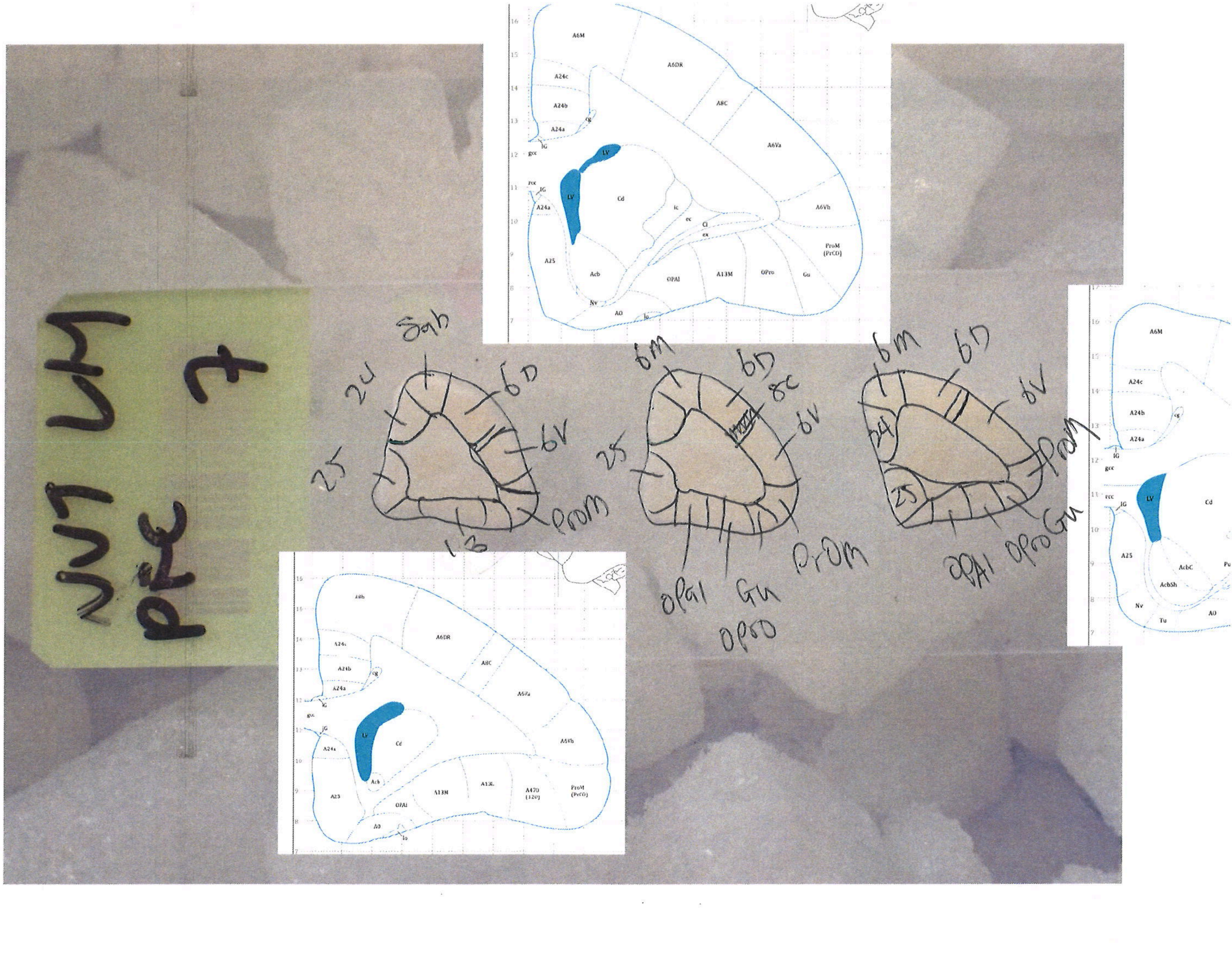

Sgb
24
6D
6V
25
ProM
13
6M
6D
8C
6V
25
ProM
OPAI
Gu
OPro
6M
6D
6V
24
25
ProM
OPAI
OPro
Gu

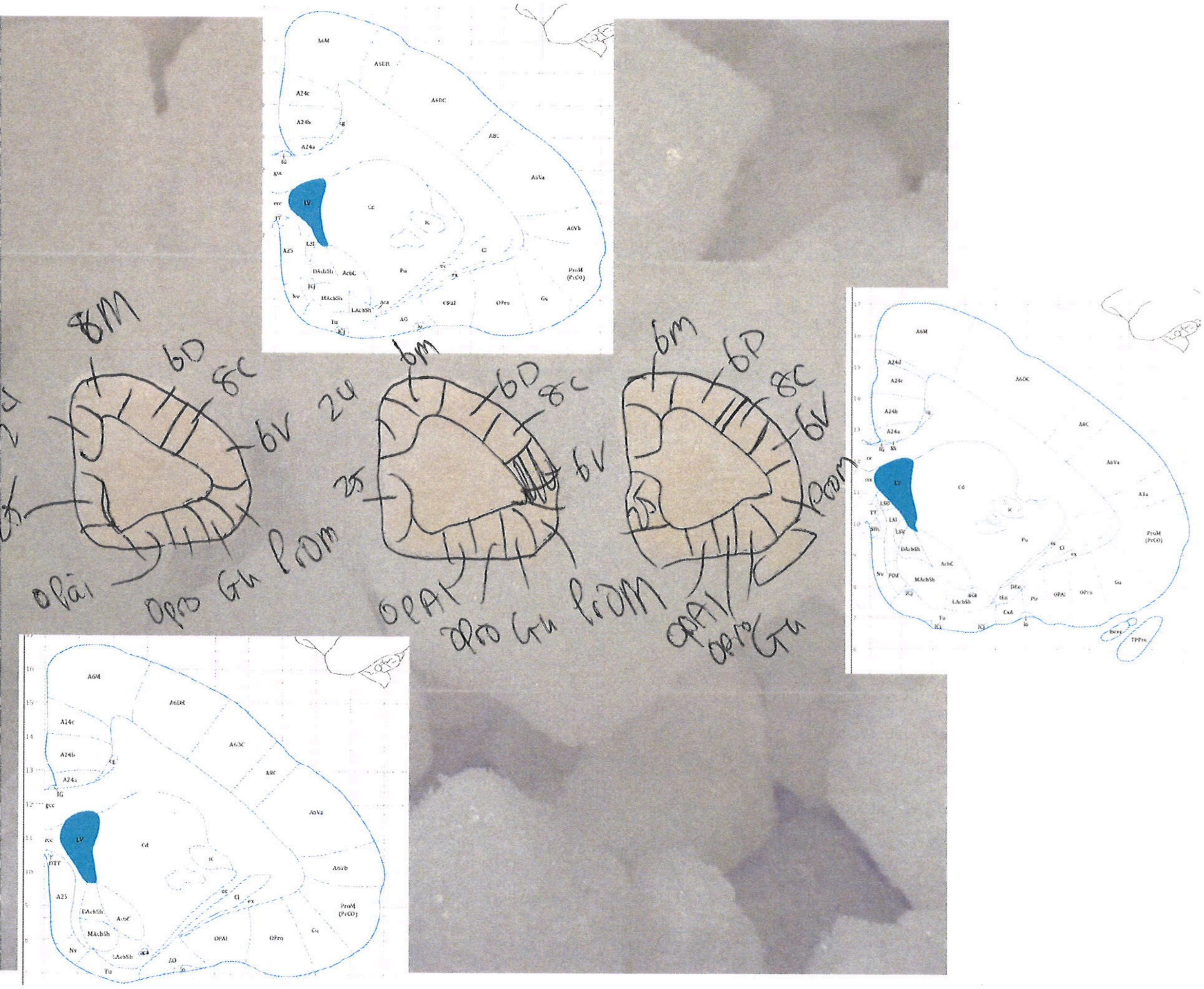

8M
6D
8C
6V
OPAI
OPro
Gu
ProM
6m
25
LV
Cd
Pu
A25
Tu
Nv
OPAl
ProM (PrCO)
A6M
A24c
A24b
A24a
IG

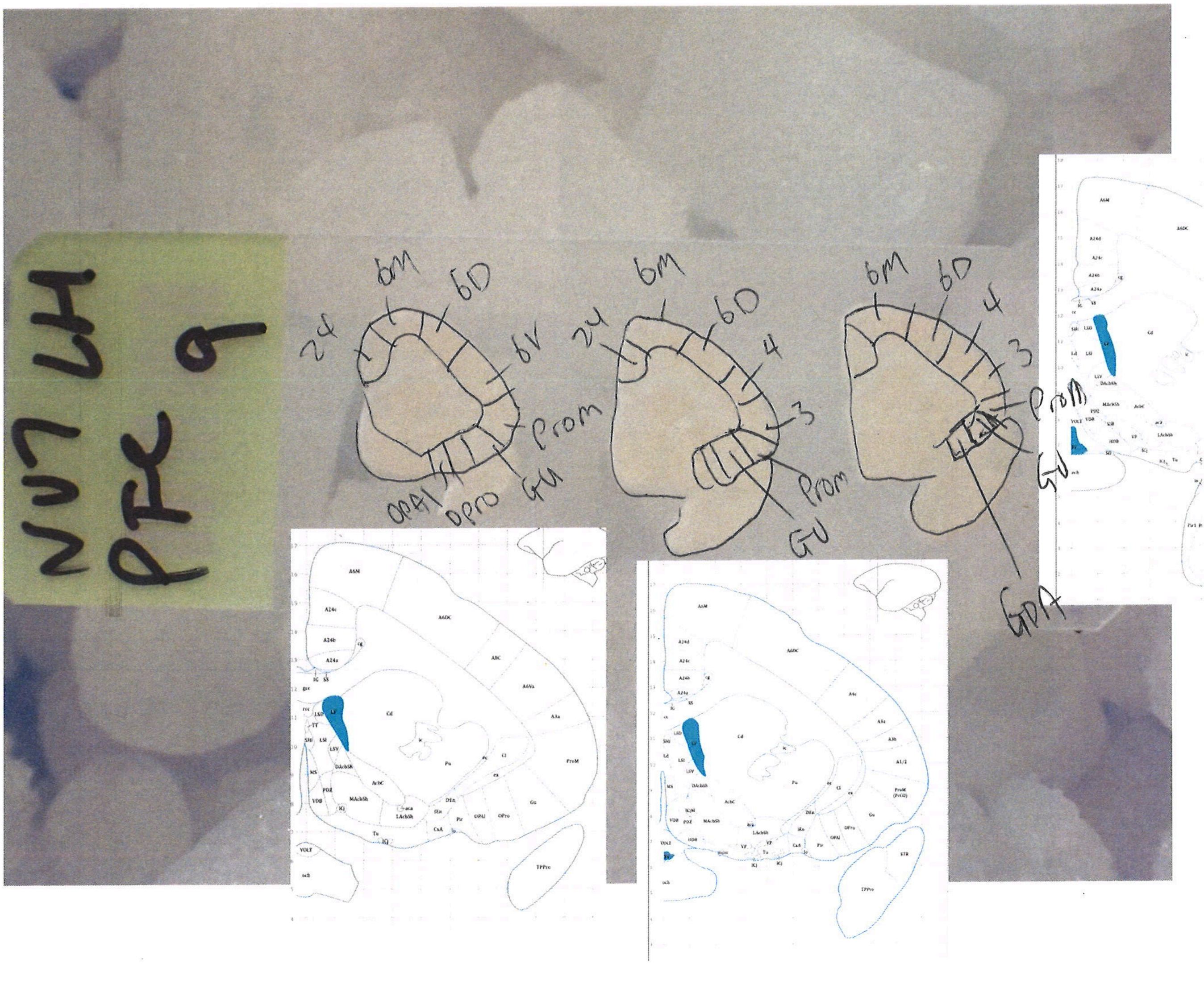
NU7 LH
PFC 9
24
6M
6D
6V
4
3
Prom
GU
OPAl
OPro
GPA

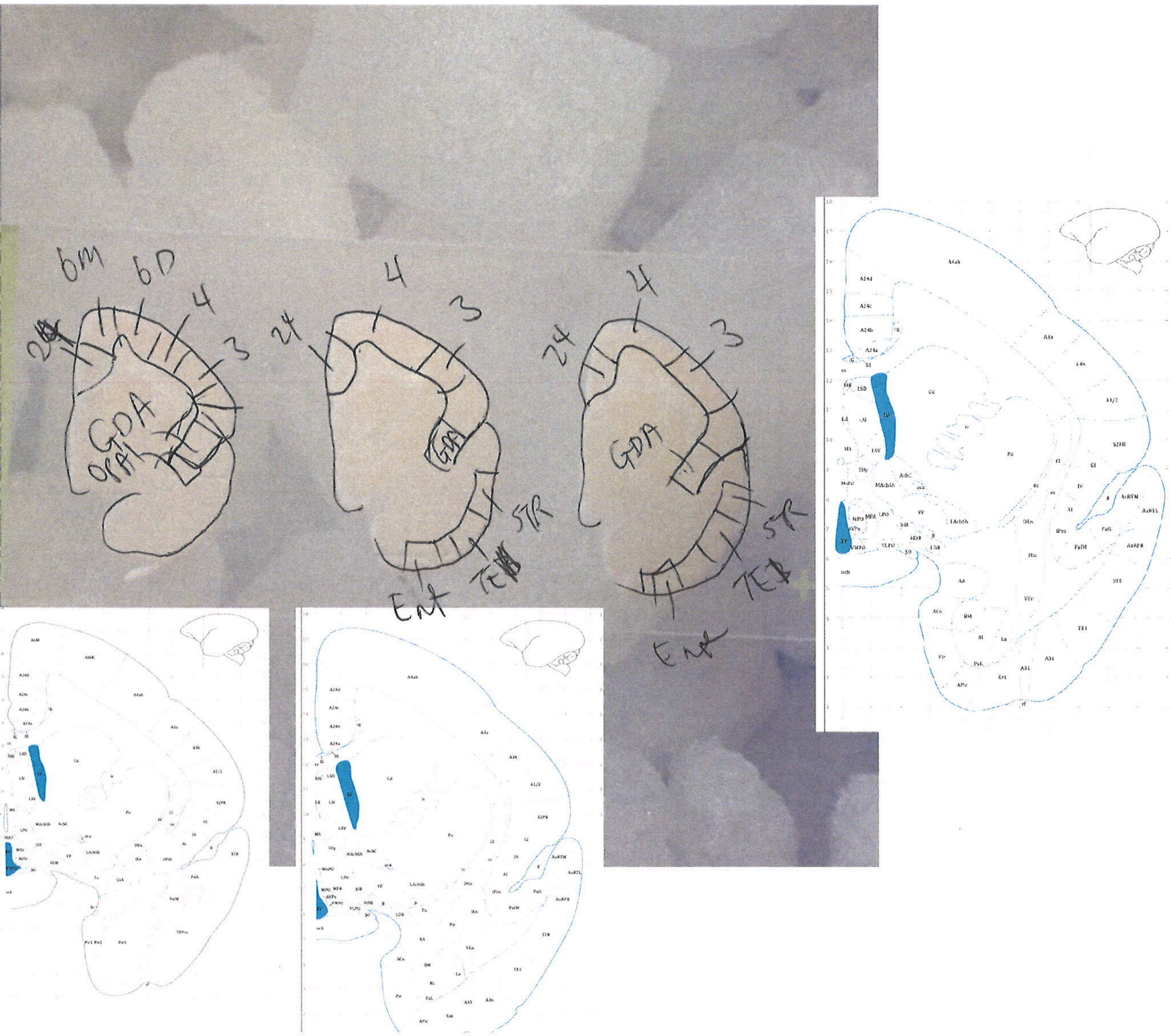
6m
6D
4
3
24
GDA
4
3
24
STR
Ent
4
3
24
GDA
STR
Ent

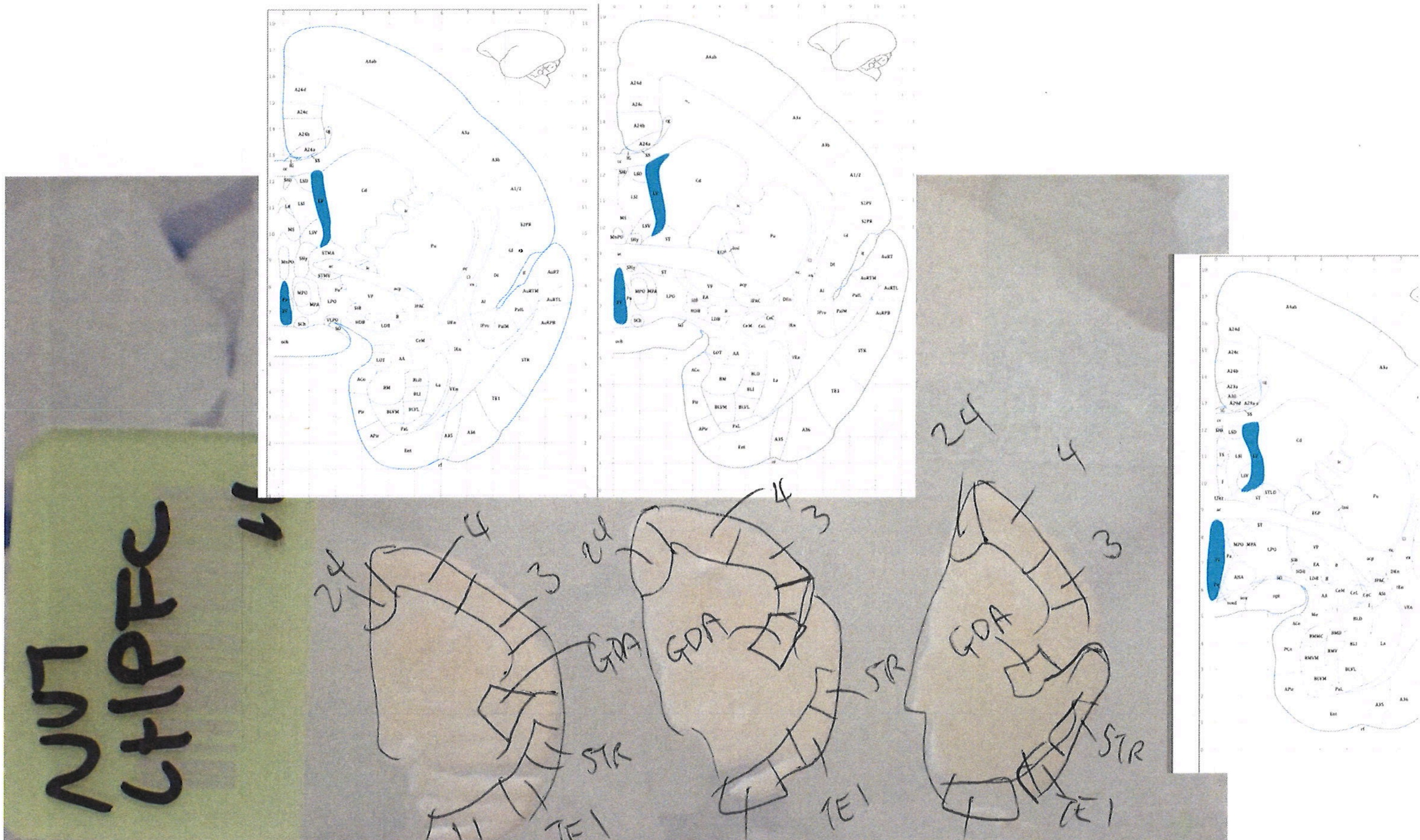
NUT
CHPFC
11
24
4
3
GDA
STR
TE1
ENT
PRh

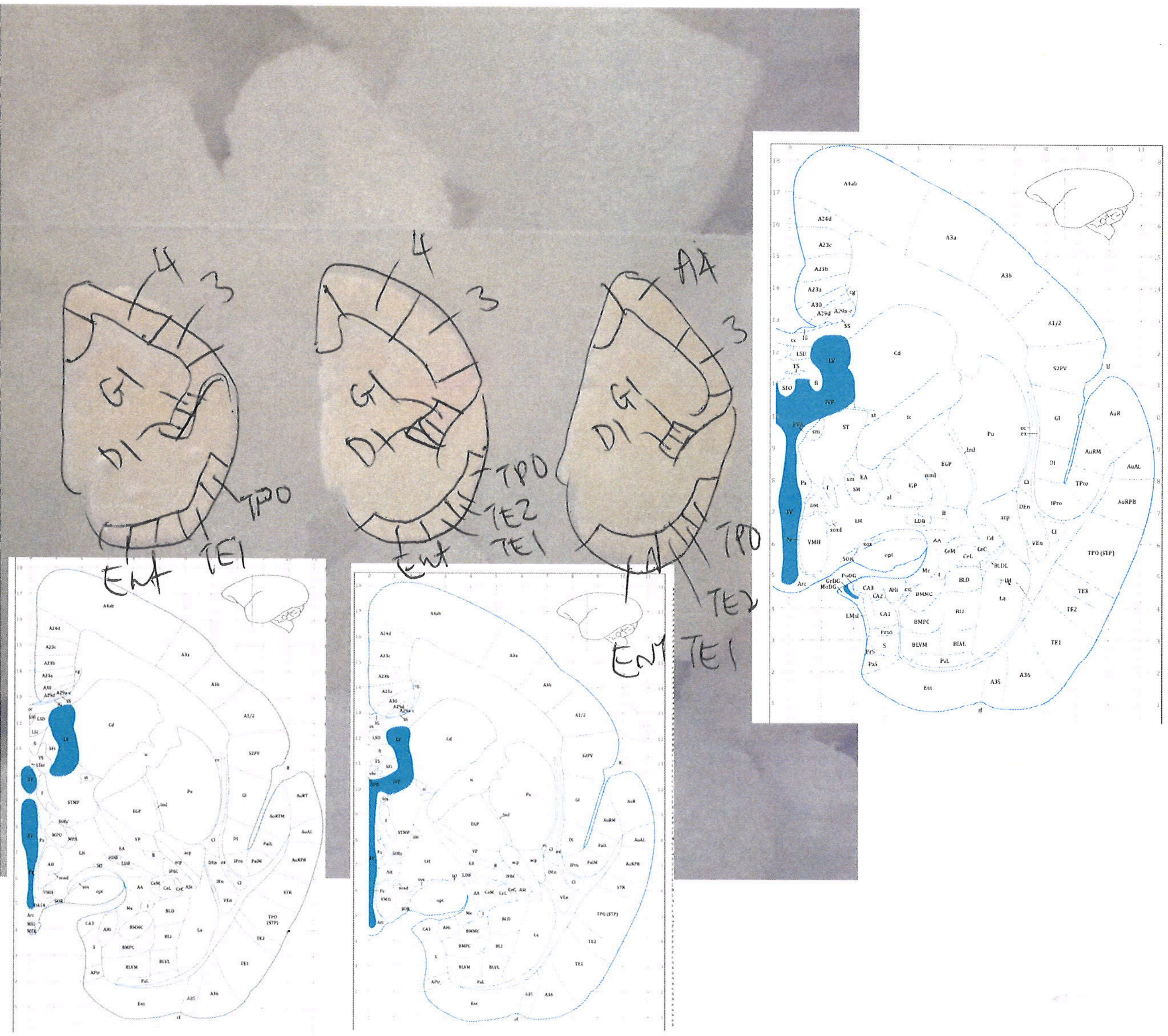
4
3
GI
DI
TPO
TE1
Ent
4
3
GI
DI
TPO
TE2
TE1
Ent
A4
3
GI
DI
TPO
TE2
TE1
Ent

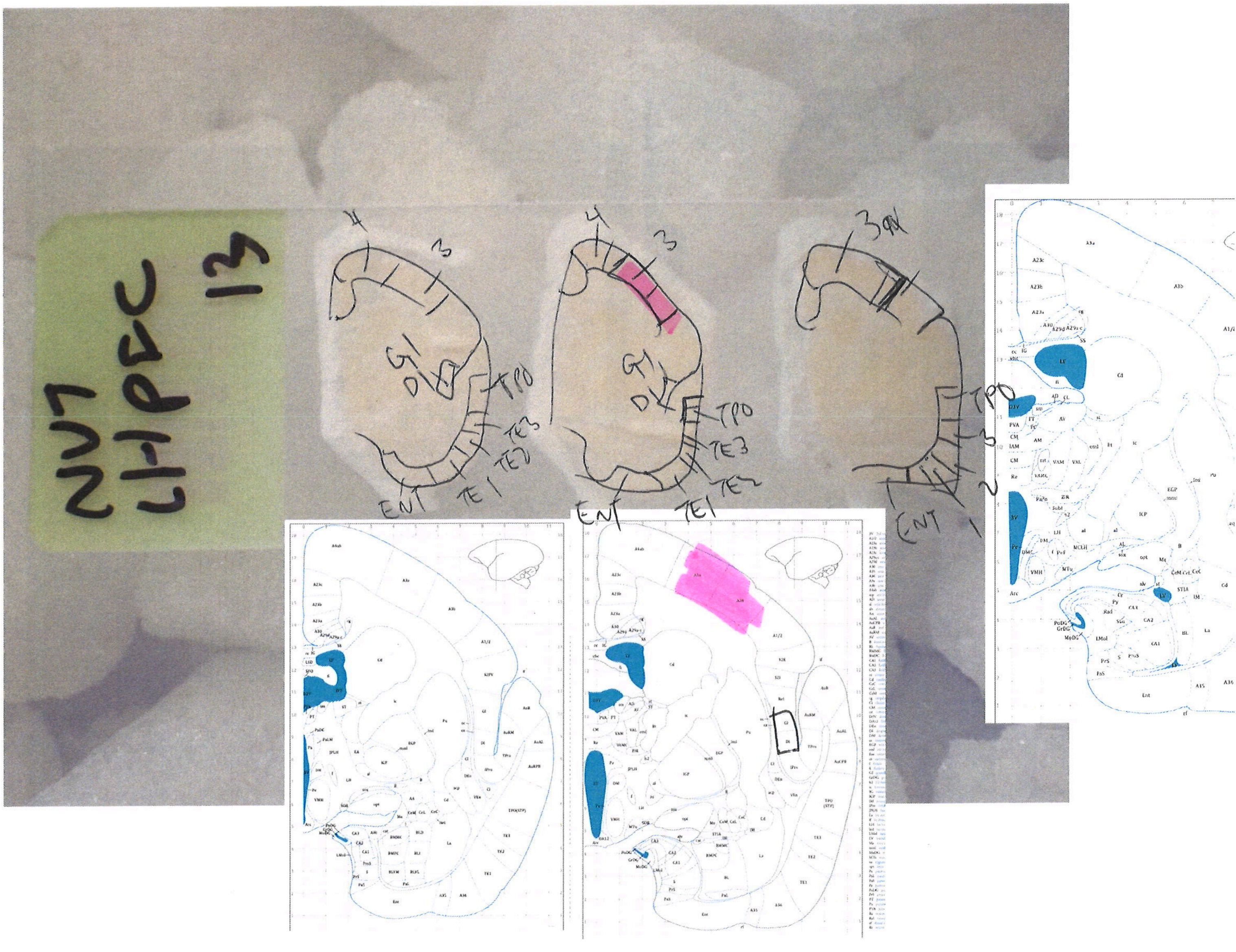
NV7
LHPEC
13
4
3
GI
DI
TPO
TE3
TE2
TE1
ENT
3av
1
2
3

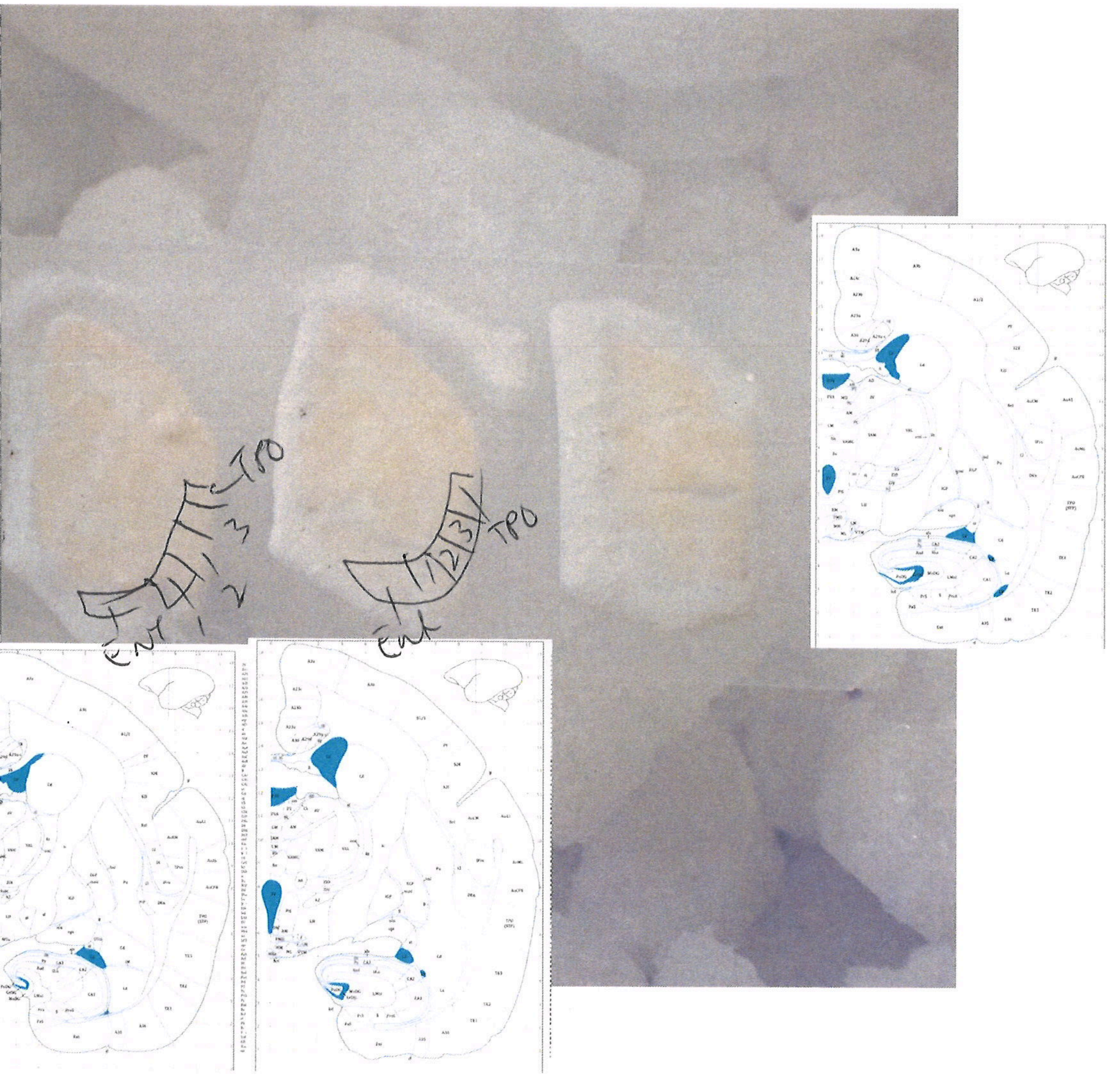

TPO
3
2
1
Ent
TPO
Ent

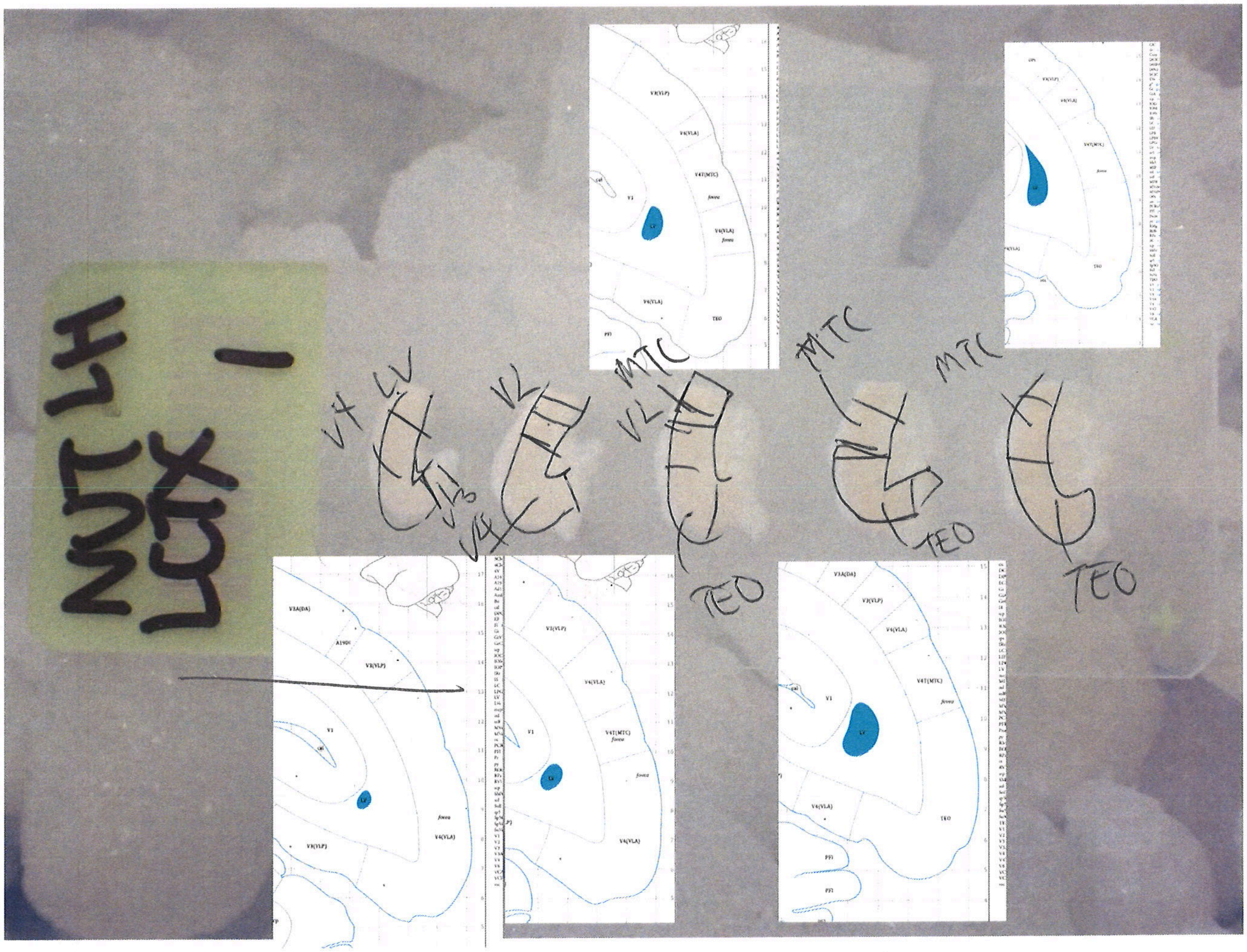
NUT LH
LCTX
1
V4
LV
V3
V2
MTC
TEO

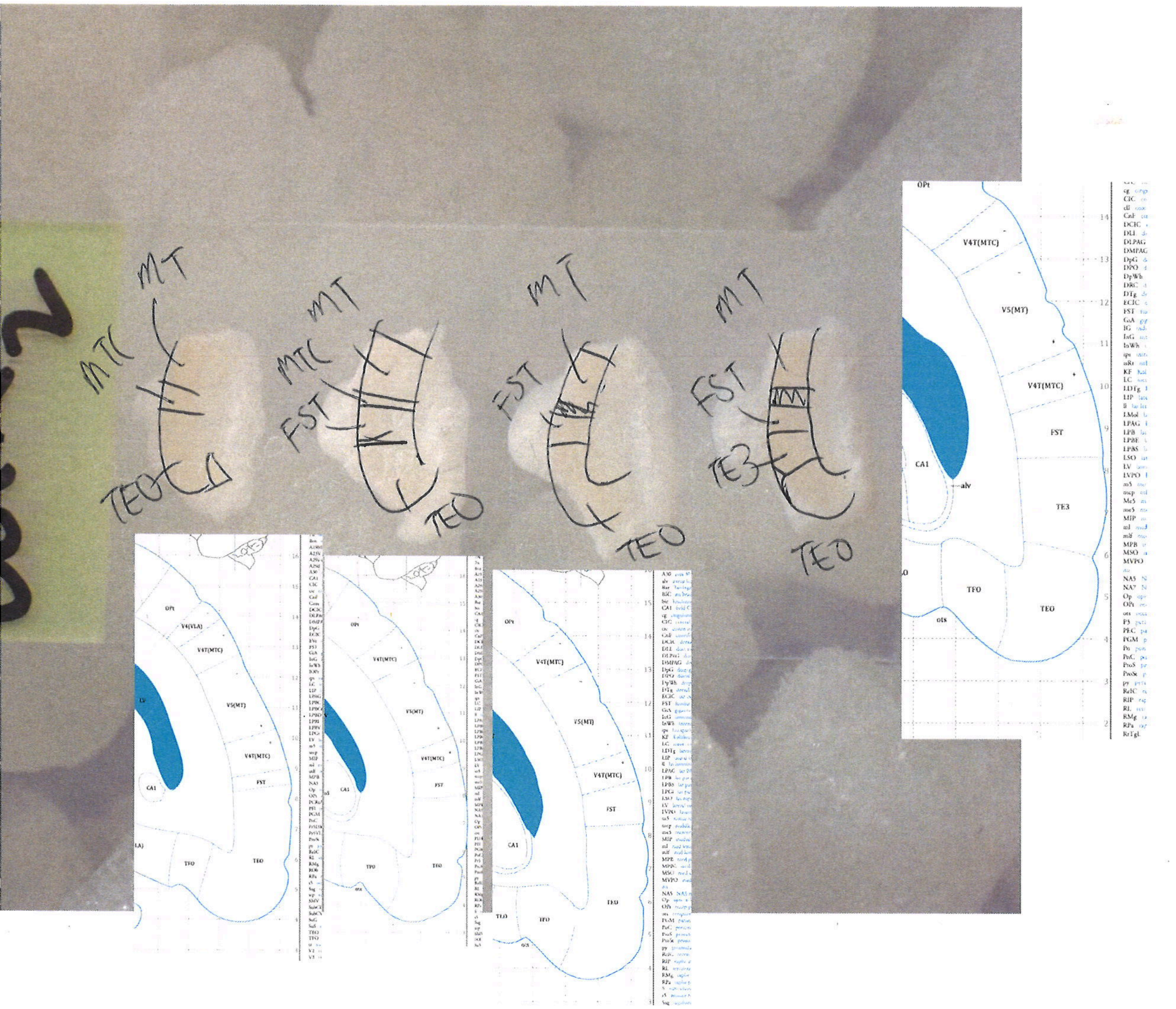

MT
MTC
TEO
MT
MTC
FST
TEO
MT
FST
TEO
MT
FST
TE3
TEO
OPt
V4T(MTC)
V5(MT)
V4T(MTC)
FST
CA1
alv
TE3
TFO
TEO
ots
OPt
V4(VLA)
V4T(MTC)
V5(MT)
V4T(MTC)
FST
CA1
TFO
TEO
OPt
V4T(MTC)
V5(MT)
V4T(MTC)
FST
CA1
TFO
TEO
OPt
V4T(MTC)
V5(MT)
V4T(MTC)
FST
CA1
TEO
TFO
TEO

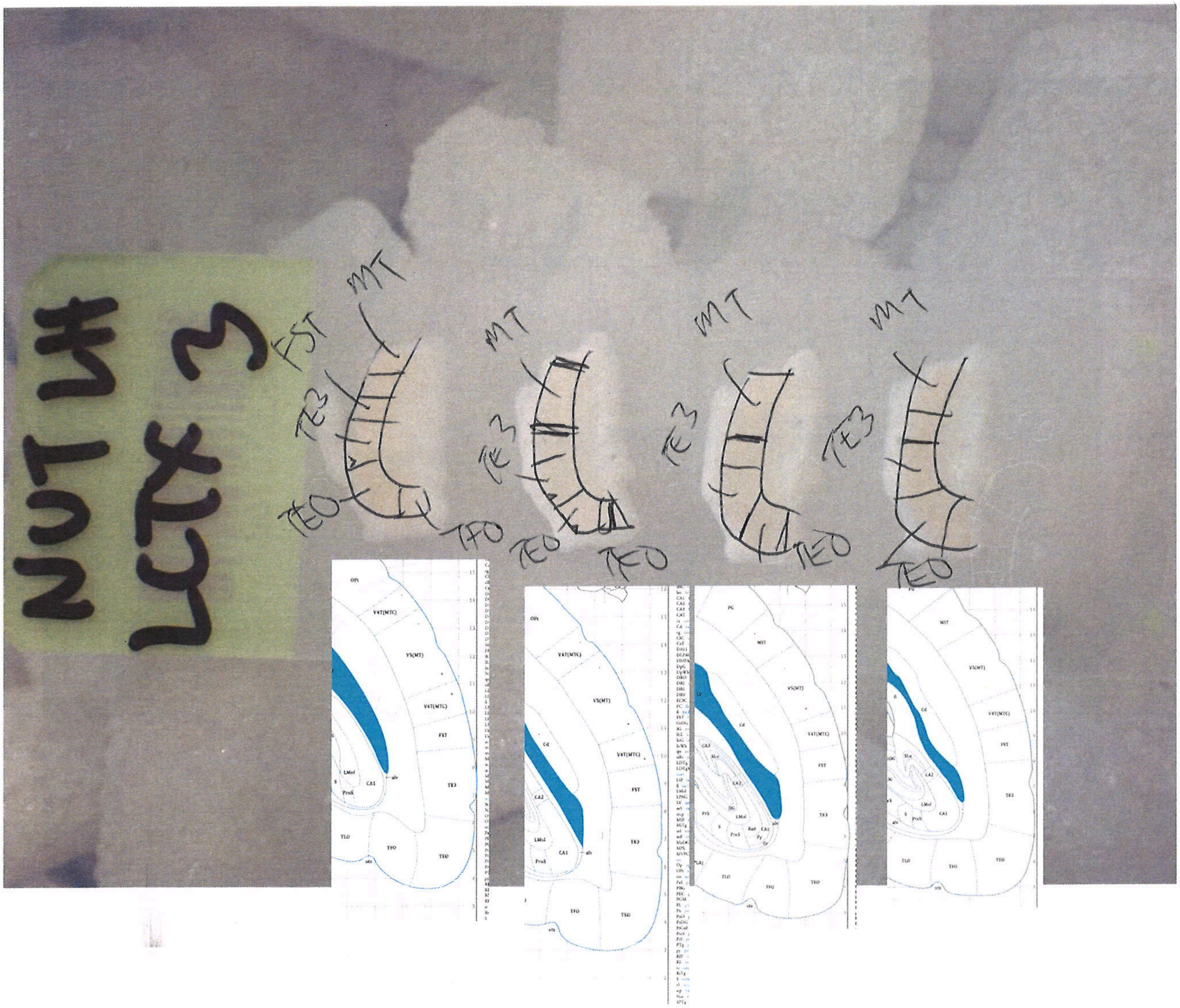
NUT LH
LCTX 3
MT
FST
TE3
TEO
TFO

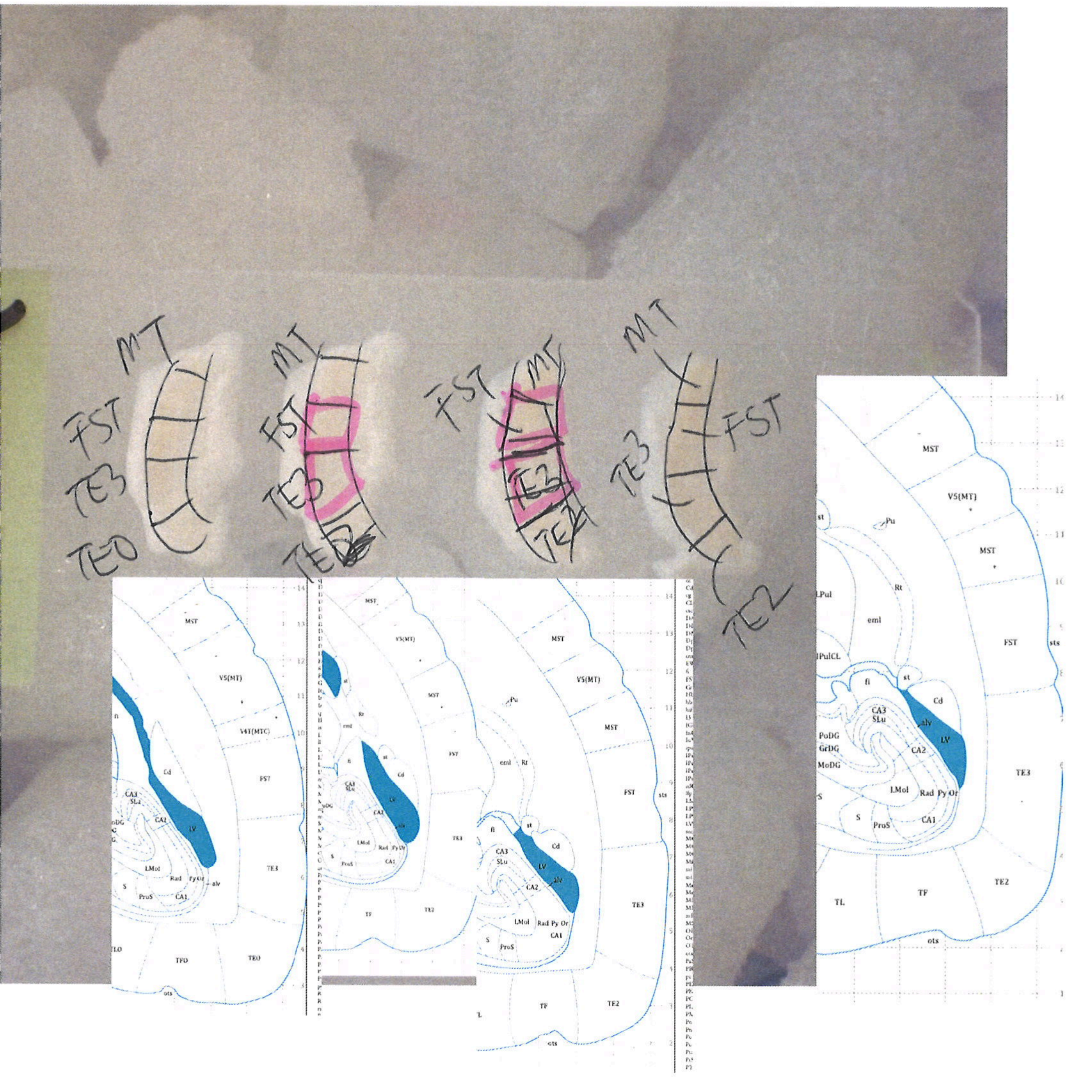
MT
FST
TE3
TEO
TE2
MST
V5(MT)
V4t(MTC)
FST
TE3
TE2
TEO
TFO
TF
TL
sts
ots
Cd
LV
alv
CA1
CA2
CA3
SLu
LMol
Rad
Py
Or
S
ProS
Pu
Rt
eml
st
fi
LPul
LPulCL
PoDG
GrDG
MoDG

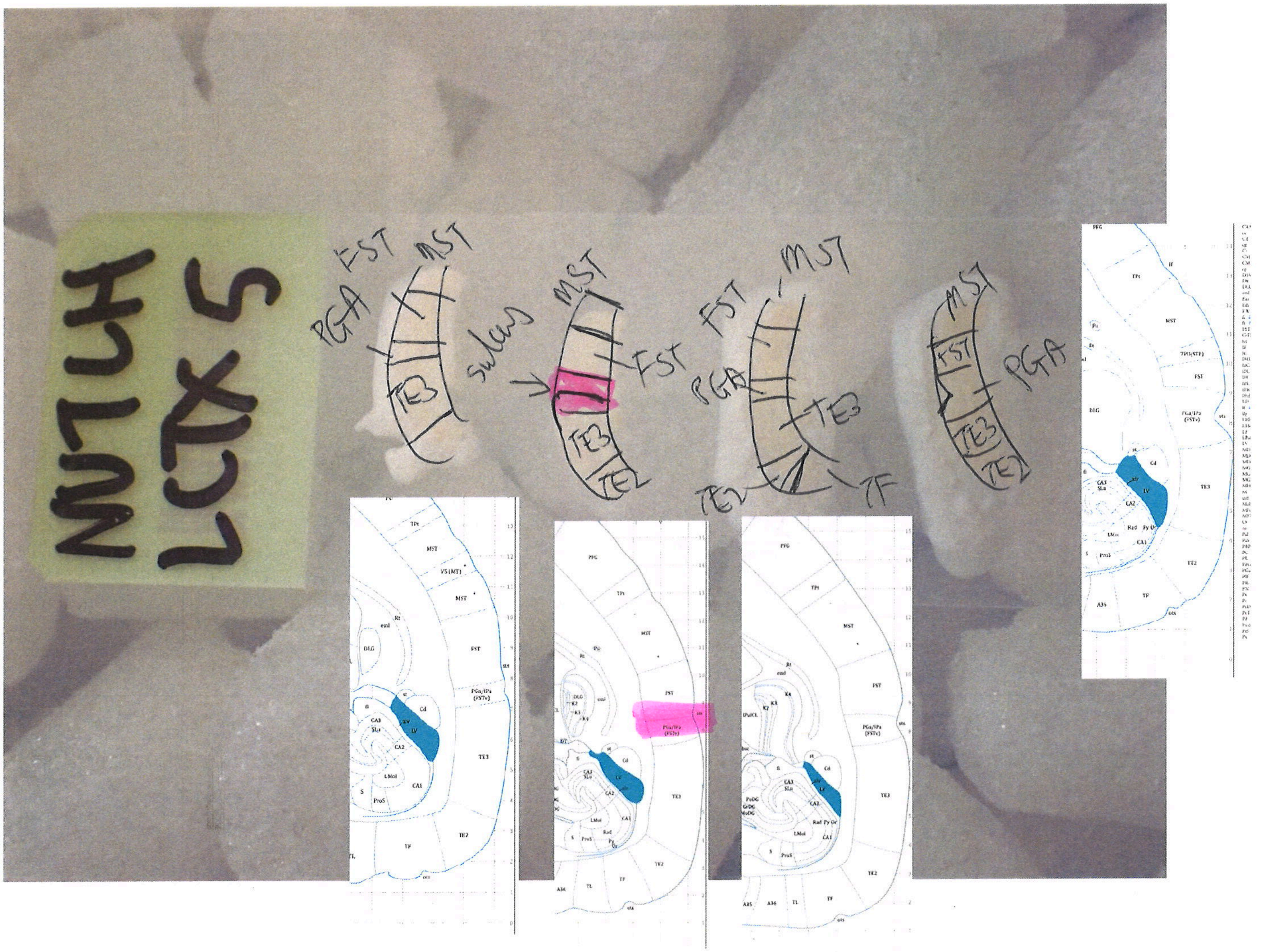
NU7LH
LCTX 5
PGA
FST
MST
TE3
sulcus
MST
FST
TE3
TE2
FST
MST
PGA
TE3
TE2
TF
MST
FST
PGA
TE3
TE2

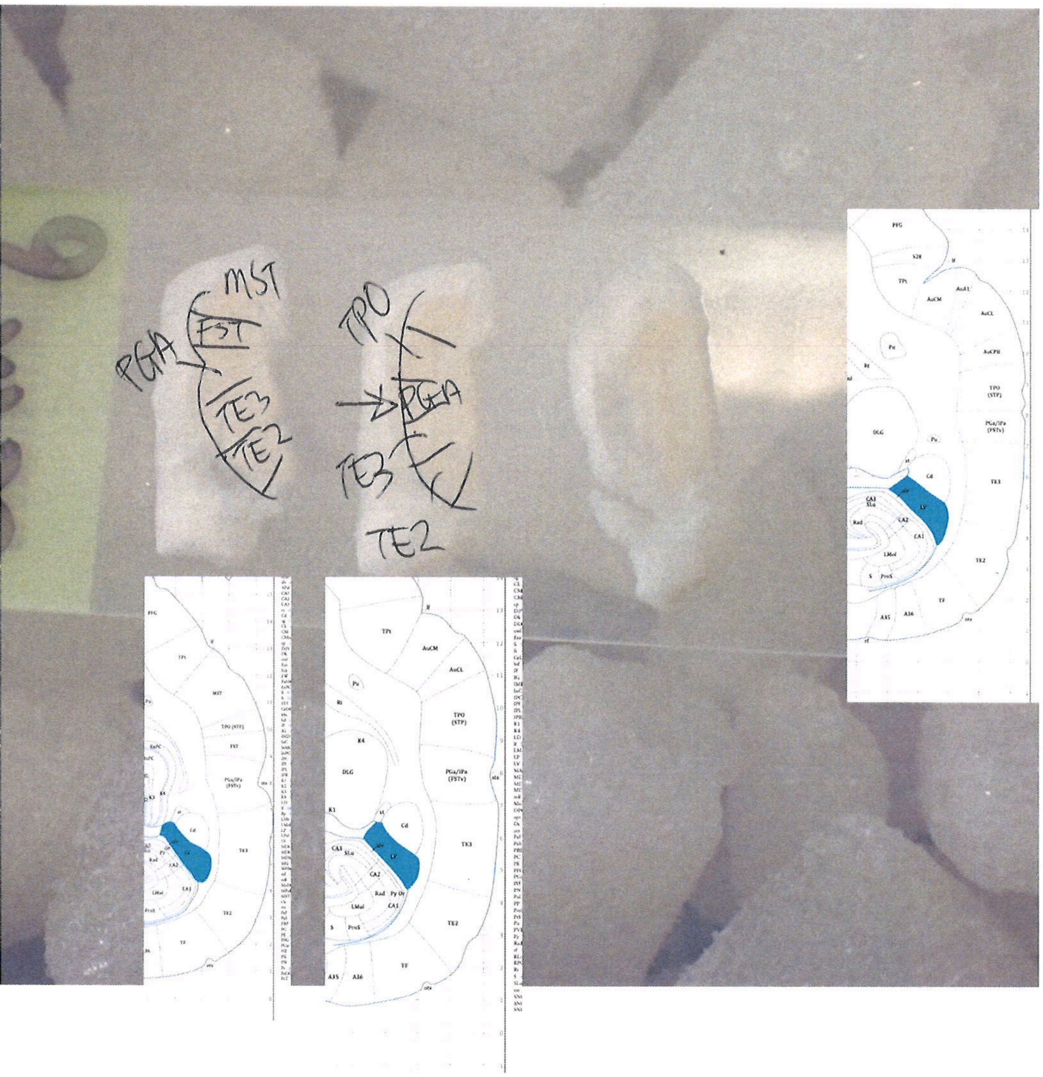

MST
FST
PGA
TE3
TE2
TPO
PGA
TE3
TE2
PFG
TPt
AuCM
AuCL
TPO (STP)
PGa/IPa (FSTv)
TE3
TE2
TF
DLG
Pu
Cd
LV
CA3
CA2
CA1
LMol
ProS
A35
A36

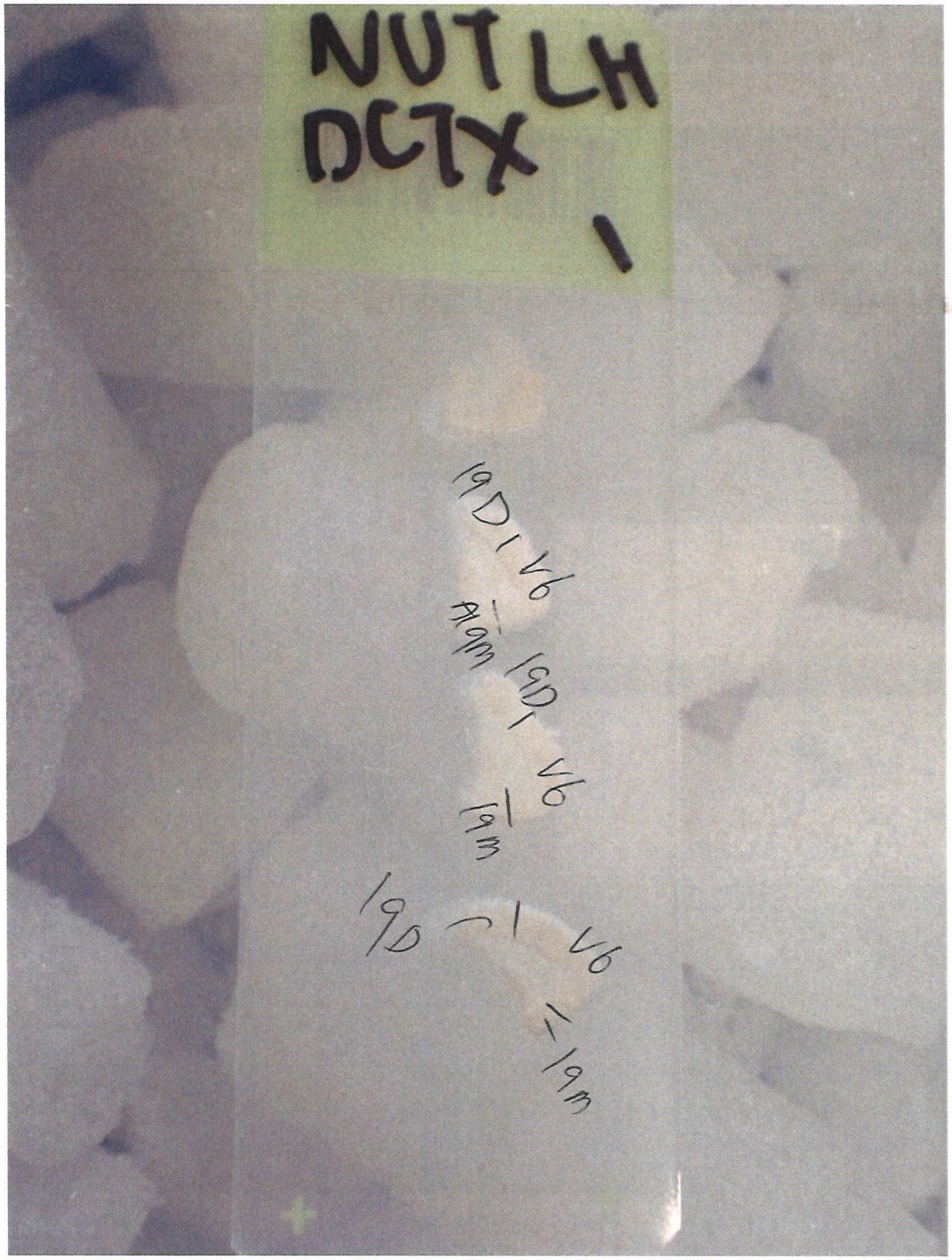
NUT LH
DC7X
1
19D 1 V6
19D 1 V6
19D 1 V6

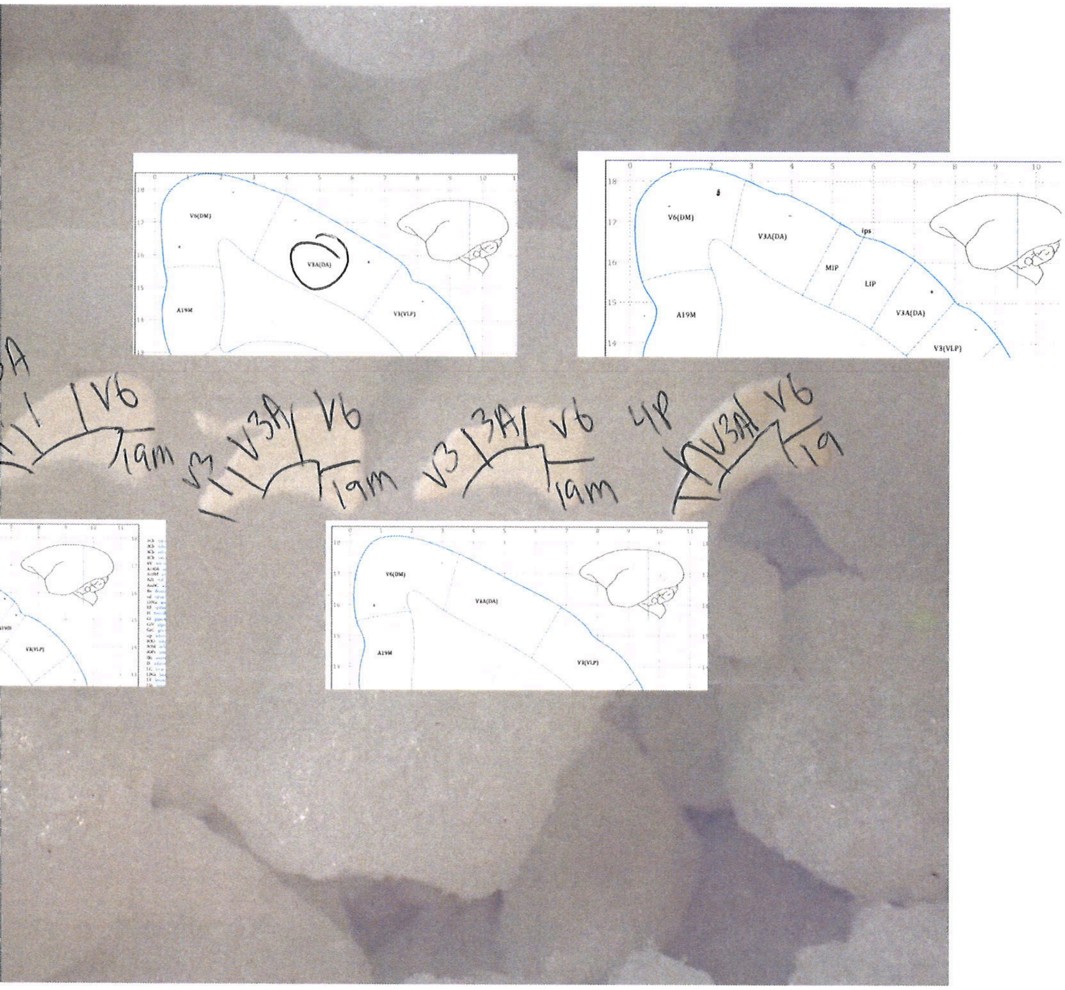
V6(DM)
V3A(DA)
A19M
V3(VLP)
ips
MIP
LIP
V6
19m
V3A
V3
19

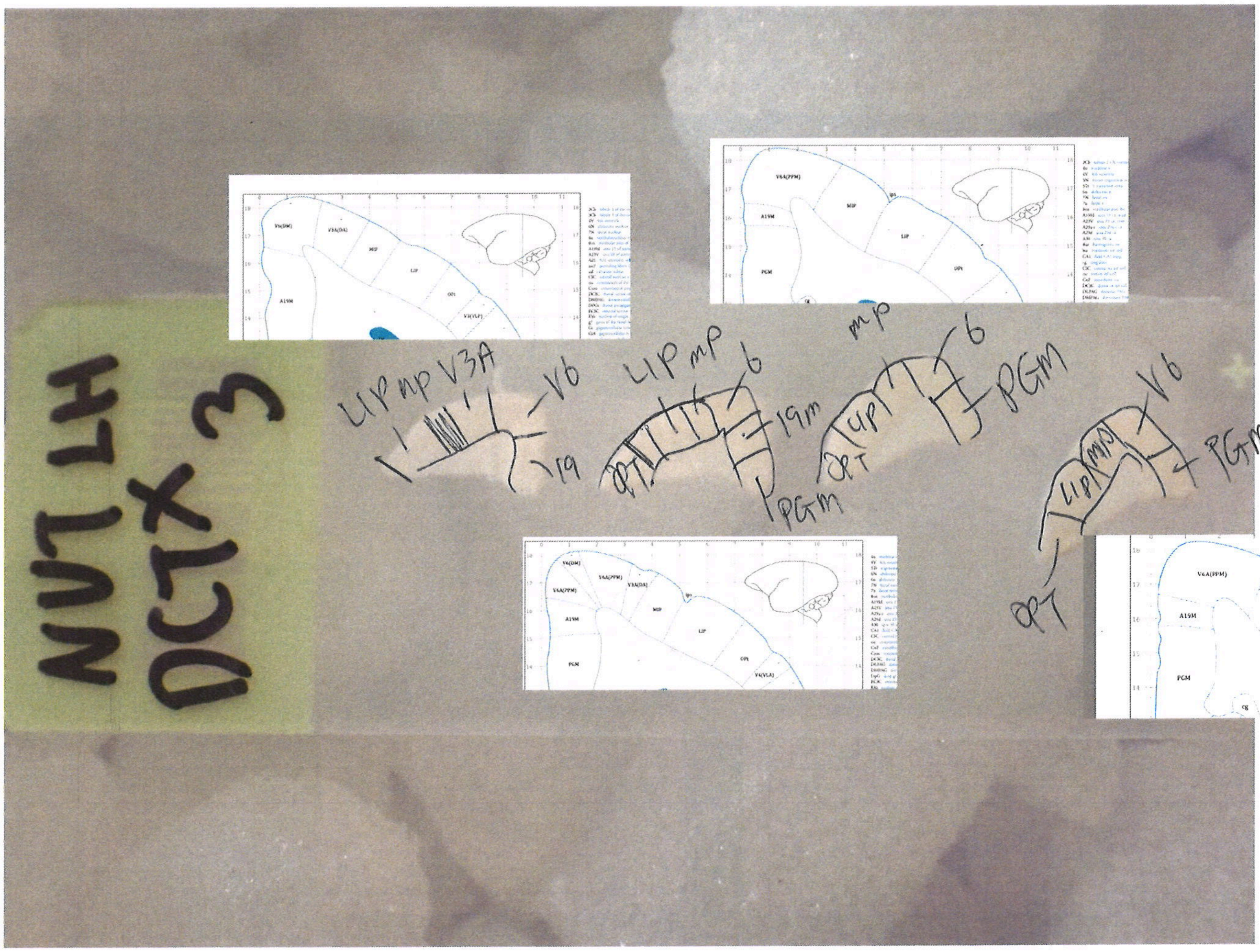
NUT LH
DCTX 3
LIP mp V3A
V6
19
LIP mp
6
19m
OPT
PGM
mp
6
PGM
LIP
OPT
V6
PGM
LIP
mp
OPT
V6A(PPM)
A19M
PGM
MIP
LIP
OPt
V6(DM)
V3A(DA)
V3(VLF)
V4(VLA)

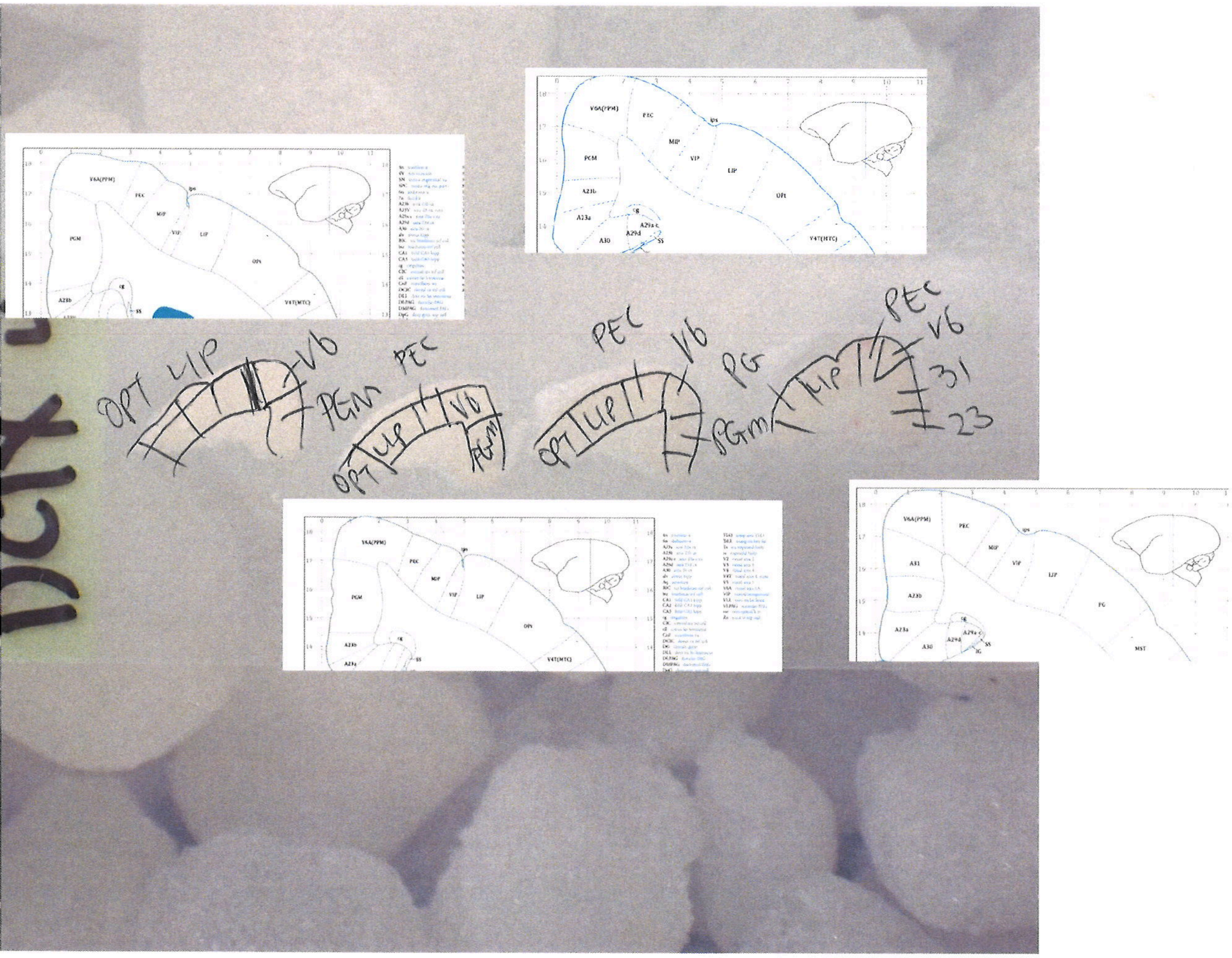
V6A(PPM)
PEC
MIP
VIP
LIP
OPt
PGM
ips
V4T(MTC)
A23a
A23b
A30
A31
PG
MST
OPT LIP V6
PEM
PEC
OPT LIP V6 PGM
PEC
V6
OPT LIP
PG
PGm
PEC
V6
LIP
31
23

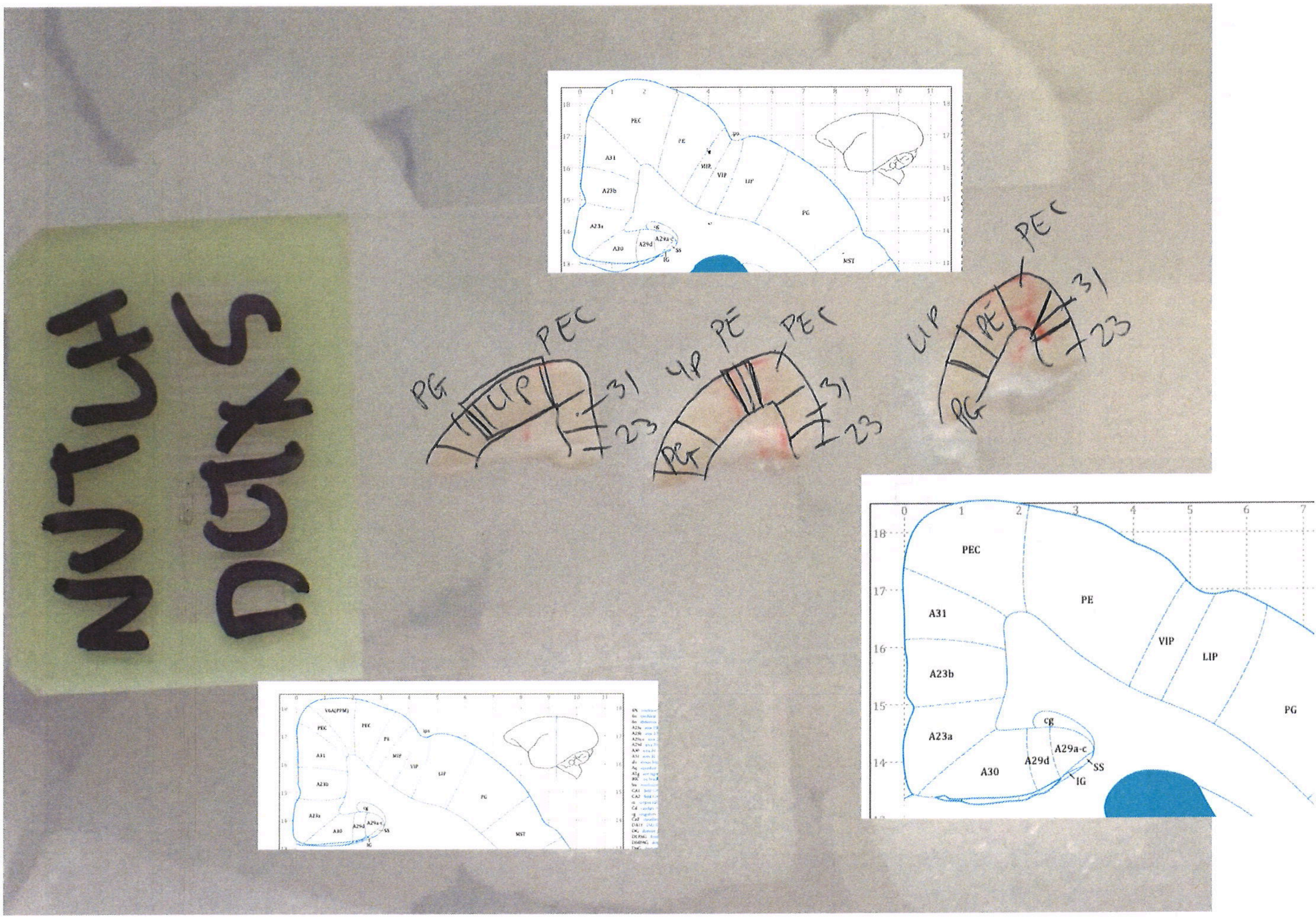
NUTLH
DCTX 5
PG
LIP
PEC
31
23
VP
PE
PEC
PE
VIP
LIP
PG
A31
A23b
A23a
A30
A29d
A29a-c
cg
SS
IG
MIP
MST

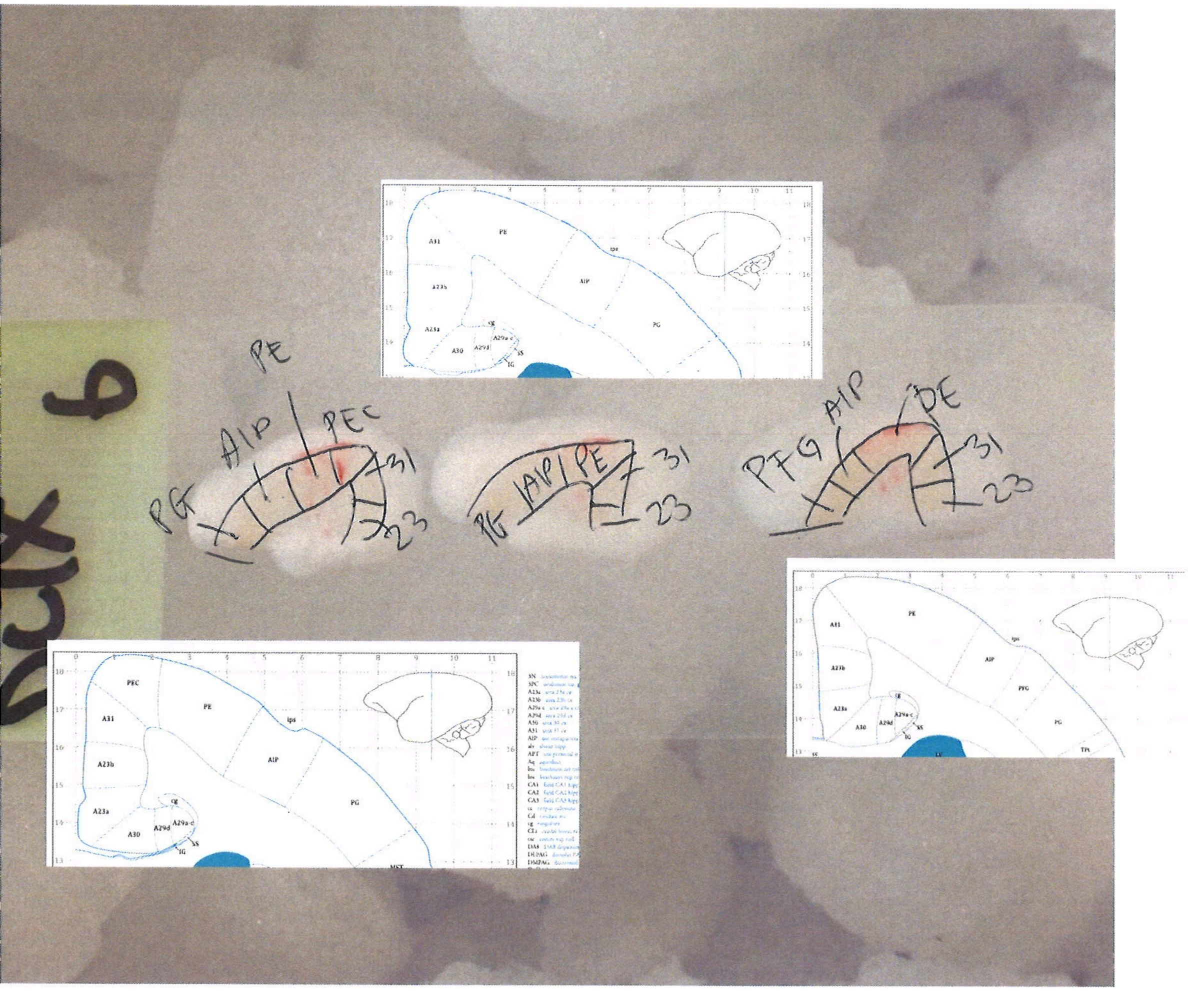
PE
AIP
PEC
PG
31
23
AIP
PE
PG
31
23
PFG
AIP
PE
31
23
A31
A23b
A23a
A30
A29d
A29a-c
AIP
PG
PFG
ips
SS
IG

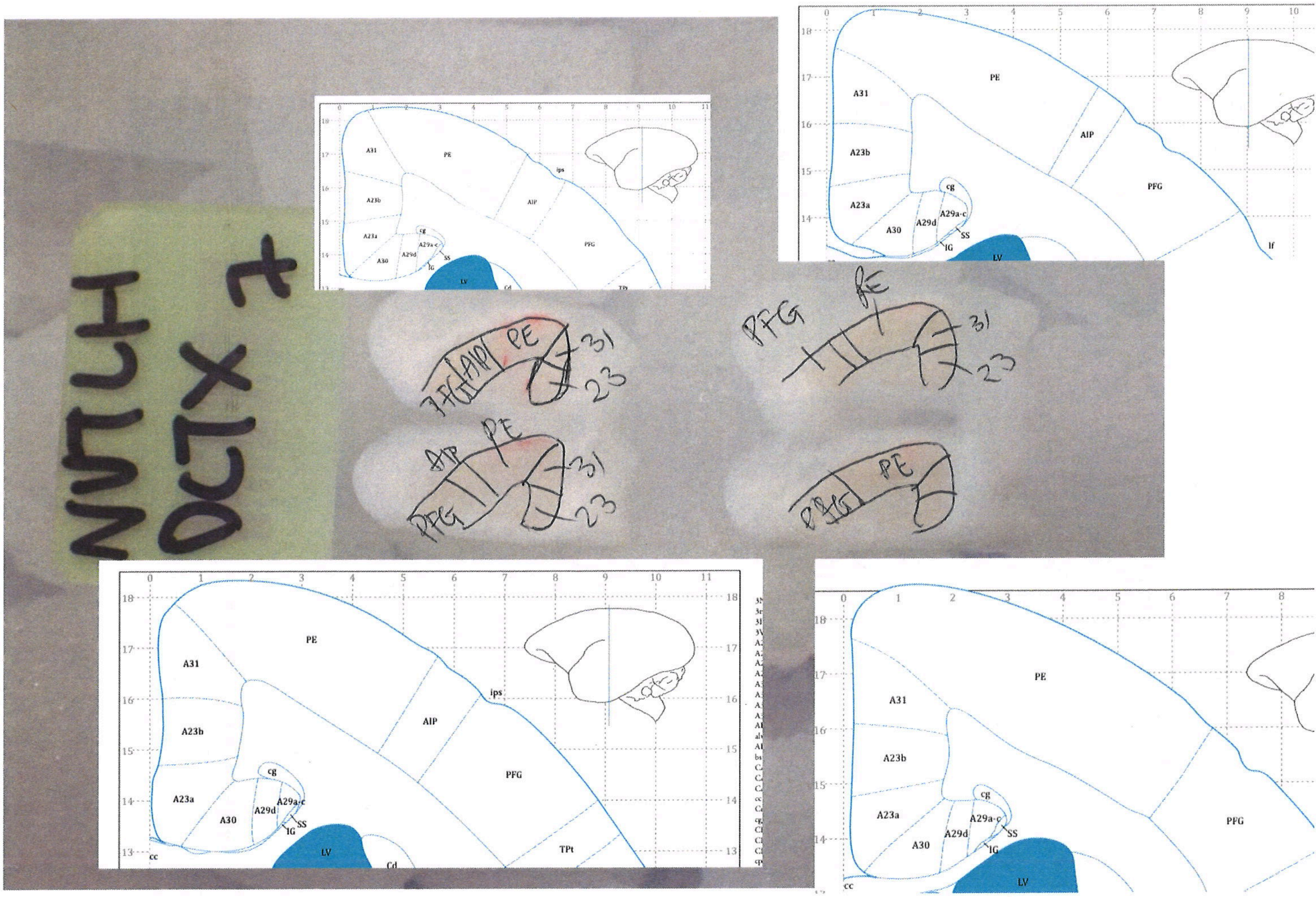

PE
A31
A23b
A23a
A30
A29d
A29a-c
cg
SS
IG
LV
Cd
AIP
PFG
ips
TPt
cc
lf
31
23

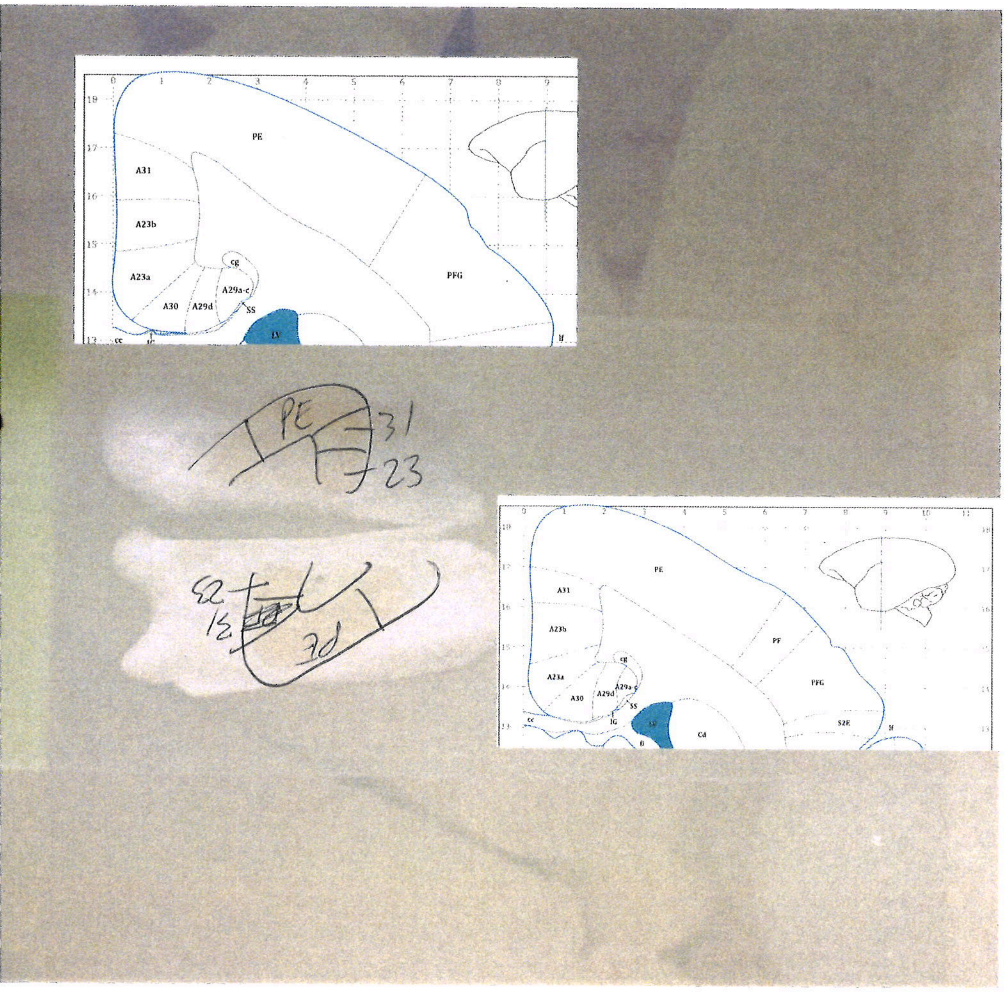
PE
A31
A23b
A23a
A30
A29d
A29a-c
cg
SS
LV
PFG
lf
PE
31
23
PE
3d
PE
A31
A23b
A23a
A30
A29d
cg
SS
IG
Cd
PF
PFG
S2E
lf
