## Supplementary material for "Conserved and specialized features of thalamocortical wiring revealed by single-cell projection mapping in mouse and marmoset": Supp. File 1: scan_xiaoyin.chen_2025-11-23-15-28-00.pdf

frontal CTX : 1mm - 1.5mm
(3-5 section)
other: 1.5mm - 2.5
(5-8 sections)

### LH PFC

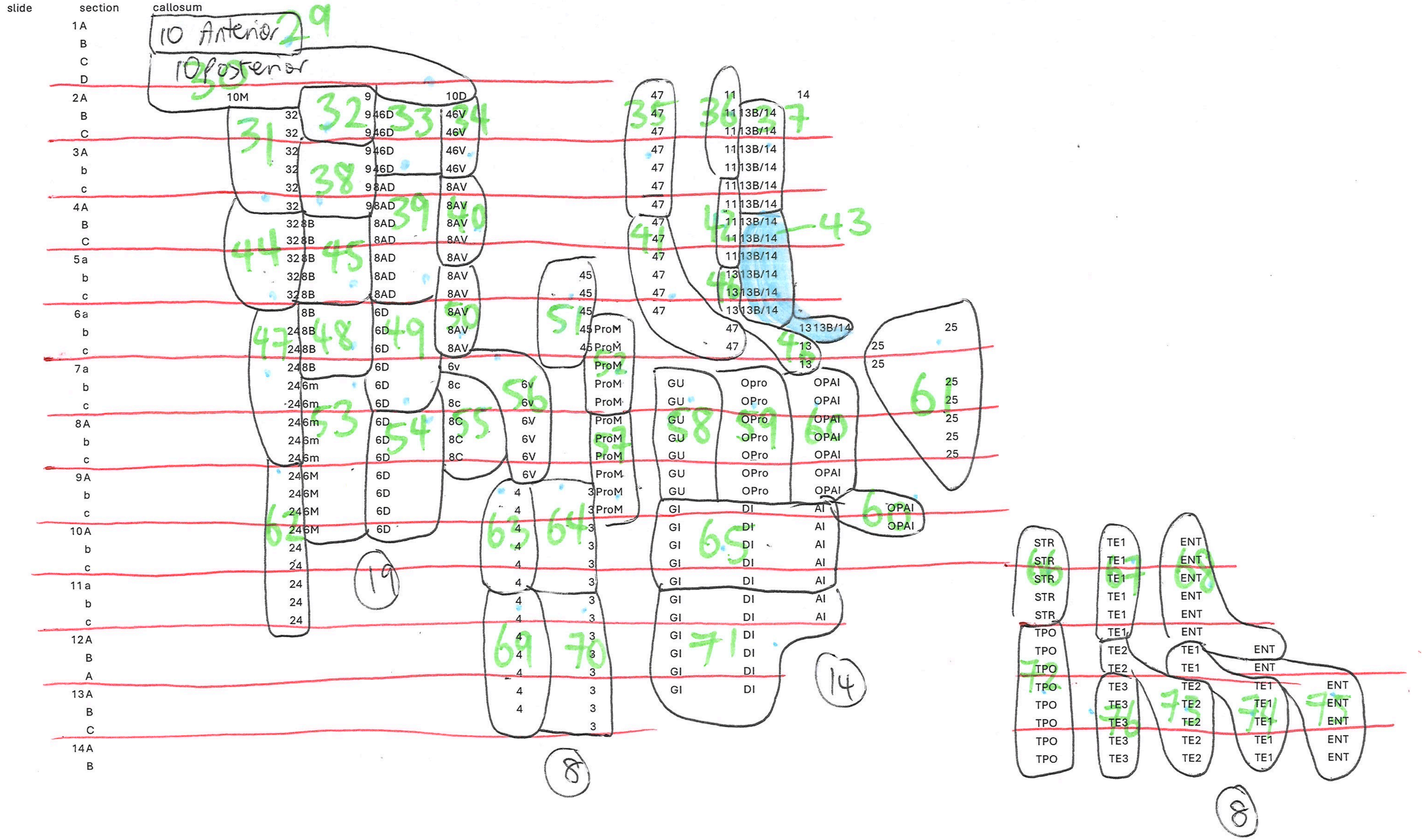


49 total tubes

### LH lateral cortex

| slide | section | | | | | | |
|---|---|---|---|---|---|---|---|
| 1 | a | VL | V4 | | | | |
| | b | VL | V4 | | | | |
| | c | MTC | VL | | TEO | | |
| | d | MTC | | | TEO | | |
| | e | MTC | | | TEO | | |
| 2 | A | MT | MTC | | TEO | | |
| | B | MT | MTC | FST | TEO | | |
| | C | MT | | FST | TEO | | |
| | D | MT | | FST | TE3 | TEO | |
| 3 | A | MT | | FST | TE3 | TEO | TFO |
| | B | MT | | | TE3 | TEO | TFO |
| | C | MT | | | TE3 | TEO | TFO |
| | D | MT | | | TE3 | TEO | |
| 4 | A | MT | | FST | TE3 | TEO | |
| | B | MT | | FST | TE3 | TE2 | |
| | C | MT | | FST | TE3 | TE2 | |
| | D | MT | | FST | TE3 | TE2 | |
| 5 | A | MST | FST | PGA | TE3 | TE2 | |
| | B | MST | FST | PGA | TE3 | TE2 | |
| | C | MST | FST | PGA | TE3 | TE2 | |
| | D | MST | FST | PGA | TE3 | TE2 | |
| 6 | A | MST | FST | PGA | TE3 | TE2 | |
| | B | | TPO | PGA | TE3 | TE2 | |
| | C | | | | | | |

14

LH dorsal cortex

| slide | section | | | | | | | |
|---|---|---|---|---|---|---|---|---|
| 1 | a | | | | | | | |
| | b | 19M | V6 | | | 19D | | |
| | c | 19M | V6 | | | 19D | | |
| | d | 19M | V6 | | | 19D | | |
| 2 | A | 19M | V6 | | 3A | | | |
| | B | 19M | V6 | | 3A | | | |
| | C | 19M | V6 | | 3A | | | |
| | D | 19M | V6 | | 3A | MIP | LIP | |
| 3 | A | 19M | V6 | | 3A | MIP | LIP | |
| | B | 19M | V6 | | | MIP | LIP | OPT |
| | C | PGM | V6 | | | MIP | LIP | OPT |
| | D | PGM | V6 | | | MIP | LIP | OPT |
| 4 | A | PGM | V6 | | | | LIP | OPT |
| | B | PGM | V6 | | PEC | | LIP | OPT |
| | C | PGM | V6 | | PEC | | LIP | OPT |
| | D | | 23 | 31 | PEC | | LIP | PG |
| 5 | A | | 23 | 31 | PEC | PE | LIP | PG |
| | B | | 23 | 31 | PEC | PE | LIP | PG |
| | C | | 23 | 31 | PEC | PE | LIP | PG |
| 6 | A | | 23 | 31 | PEC | PE | AIP | PG |
| | B | | 23 | 31 | | PE | AIP | PG |
| | C | | 23 | 31 | | PE | AIP | PFG |
| 7 | A | | 23 | 31 | | PE | AIP | PFG |
| | B | | 23 | 31 | | PE | AIP | PFG |
| | C | | 23 | 31 | | PE | | PFG |
| | D | | 23 | 31 | | PE | | PFG |
| 8 | A | | 23 | 31 | | PE | | |
| | B | | 23 | 31 | | PE | | |

29 30 31 32 33 34 35 36 37 38 39 40 41 42 43 44

45

V3
V3
V3

16 tubes

### LH Visual cortex

| slide | section | area | are | area | | |
|---|---|---|---|---|---|---|
| | 1 a | V1 fovea | | | | |
| | 1 b | v1 fovea | | | | |
| | 1 c | v1 fovea | | | | |
| | 1 d | v1 fovea | | | | |
| | 2 a | v1 fovea | | | | |
| | 2 b | v1 fovea | | | | |
| | 2 c | v1 fovea | v1 calcrine | | | |
| | 3 a | v1 fovea | v1 calcrine | | | |
| | 3 b | v1 fovea | v1 calcrine | | | |
| | 3 c | v1 fovea | v1 calcrine | | | |
| | 4 a | v1 fovea | v1 calcrine | | | |
| | 4 b | v1 fovea | v1 calcrine | | | |
| | 4 c | v1 fovea | v1 calcrine | | | |
| | 5 a | V1 operculum dorsal | V1 opervulum ventral | V1 calcrine | | |
| | 5 b | V1 operculum dorsal | V1 opervulum ventral | V1 calcrine | V2 ventrolatera | |
| | 5 c | V1 operculum dorsal | V1 opervulum ventral | V1 calcrine | V2 ventrolatera | |
| | 6 a | V1 operculum dorsal | V2 dorsal | v2 ventral | V1 calcrine | |
| | 6 b | V1 calcrine | V2 dorsal | v2 ventral | | |
| | 6 c | V1 calcrine | V2 dorsal | v2 ventral | | |
| | 7 a | V2 dorsal | v2 ventral | V1 calcrine | | |
| | 7 b | V2 dorsal | v2 ventral | V1 calcrine | | |
| | 7 c | V2 dorsal | v2 ventral | V1 calcrine | | |
| | 8 a | V2 dorsal | v2 ventral | V1 calcrine | | |
| | b | V2 dorsal | v2 ventral | V1 calcrine | V3 | |
| | c | V2 dorsal | v2 ventral | V1 calcrine | V3 | |
| | 9 a | V2 dorsal | v2 ventral | V1 calcrine | V3 | |
| | b | V2 dorsal | v2 ventral | V1 calcrine | V3 | |
| | c | V2 dorsal | v2 ventral | V1 calcrine | V3 | A19 |
| | 10 a | V2 dorsal | V3 | V1 calcrine | A19 | |
| | b | V6 | V3 | V1 calcrine | A19 | |
| | c | V6 | V3 dorsal | V1 calcrine | A19 | V3 ventral |

Tube 1, V1 fovea #1

Tube 2, V1 calc #1

Tube 3, V1 fov #2

#4

#5

#6 calcrine #2, include 6B, 6C

V2 ventral

#8

calcrine #3

12

### RH PFC

| slide | section | Callosum | area | area | | | | | | | | | | sulcus | | | | | |
|---|---|---|---|---|---|---|---|---|---|---|---|---|---|---|---|---|---|---|---|
| | 1a | | | 10 | | | | | | | | | | | | | | | |
| | b | | | 10 | | | | | | | | | | | | | | | |
| | c | | | 10 | | | | | | | | | | | | | | | |
| | d | | | 10 | | | | | | | | | | | | | | | |
| | 2a | | 10M | | | 9 | 10D | | | 47 | 11 | 14 | | | | | | | |
| | b | | 10M | | | 9 46D | 46V | | | 47 | 11 13B/14 | | | | | | | | |
| | c | | | 32 | | 9 46D | 46V | | | 47 | 11 13B/14 | | | | | | | | |
| | 3A | | | 32 | | 9 46D | 46V | | | 47 | 11 13B/14 | | | | | | | | |
| | B | | | 32 | | 9 46D | 46V | | | 47 | 11 13B/14 | | | | | | | | |
| | C | | | 32 | | 9 8AD | 8AV | | | 47 | 11 13B/14 | | | | | | | | |
| | 4A | | | 32 | | 8AD | 8AV | | | 47 | 11 13B/14 | | | | | | | | |
| | B | | | 32 | 8B | 8AD | 8AV | | | 47 | 11 13B/14 | | | | | | | | |
| | C | | | 32 | 8B | 8AD | 8AV | | | 47 | 11 13B/14 | | | | | | | | |
| | 5a | | | 32 | 8B | 8AD | 8AV | | | 47 | 11 13B/14 | | | | | | | | |
| | b | | | 32 | 8B | 8AD | 8AV | | 45 | 47 | 13 13B/14 | | | | | | | | |
| | 6a | | | | 8B | 8AD | 6D | 8AV | | | | | | | | | | | |
| | b | | | 24 | 8B | 6D | 8AV | | 45 | 47 | 13 13B/14 | | 25 | | | | | | |
| | c | | | 24 | 8B | 6D | 8AV | | 45 ProM | | 47 | 13 | 25 | | | | | | |
| | 7a | | | 24 | 8B | 6D | 6v | | ProM | | | 13 | 25 | | | | | | |
| | b | | | 24 | 6m | 6D | 6v | | ProM | GU | Opro | | 25 | | | | | | |
| | c | | | 24 | 6m | 6D | 6v | | ProM | GU | OPro | | 25 | | | | | | |
| | 8A | | | 24 | 6m | 6D | 8C | 6V | ProM | GU | OPro | OPAI | | 25 | | | | | |
| | b | | | 24 | 6m | 6D | 8C | 6V | ProM | GU | OPro | OPAI | | 25 | | | | | |
| | c | | | 24 | 6m | 6D | 8C | 6V | ProM | GU | OPro | OPAI | | | | | | | |
| | 9A | | | 24 | 6M | 6D | 8C | 6V | ProM | GU | OPro | OPAI | | | | | | | |
| | b | | | 24 | 6M | 6D | | 4 | ProM | GU | OPro | OPAI | | | | | | | |
| | 10A | | | 24 | 6M | 6D | | 4 | 3 ProM | GI | DI | AI | OPAI | | STR | TE1 | ENT | | |
| | B | | | 24 | | | | 4 | 3 | GI | DI | AI | | | STR | TE1 | ENT | | |
| | C | | | 24 | | | | 4 | 3 | GI | DI | AI | | | STR | TE1 | ENT | | |
| | 11A | | | 24 | | | | 4 | 3 | GI | DI | AI | | | STR | TE1 | ENT | | |
| | B | | | 24 | | | | 4 | 3 | GI | DI | AI | | | STR | TE1 | ENT | | |
| | 12A | | | 24 | | | | 4 | 3 | GI | DI | AI | | | STR | TE1 | ENT | | |
| | B | | | | | | | 4 | 3 | | | | | | STR | TE1 | ENT | | |
| | 13A | | | | | | | 4 | 3 | GI | DI | | | | TPO | TE2 | TE1 | ENT | |
| | B | | | | | | | 4 | 3 | GI | DI | | | | TPO | TE2 | TE1 | ENT | |
| | 14A | | | | | | | 4 | 3 | GI | DI | | | | TPO | TE3 | TE2 | TE1 | ENT |

total: 47

#### RH lateral cortex

| slide | section | | | | | | |
|---|---|---|---|---|---|---|---|
| | 1 a | MT | | TEO | TFO | | |
| | b | MT | TE3 | TEO | | | |
| | c | MT | TE3 | TEO | | | |
| | d | MT | TE3 | TEO | TFO | | |
| | e | MT | TE3 | TEO | TFO | | |
| | 2 A | MT | FST | TE3 | TEO | TFO | |
| | B | MT | FST | TE3 | TEO | TFO | |
| | C | MT | FST | TE3 | TEO | TFO | |
| | D | MT | FST | TE3 | TE2 | TF | |
| | 3 A | MT | FST | TE3 | TE2 | TF | |
| | B | MT | FST | TE3 | TE2 | TF | |
| | C | MT | FST | TE3 | TE2 | TF | |
| | D | MT | FST | TE3 | TE2 | TF | |
| | 4 A | FST | PGA | TE3 | TE2 | TF | |
| | B | MST | FST | PGA | TE3 | TE2 | TF |
| | C | MST | FST | PGA | TE3 | TE2 | TF |
| | D | | | PGA | TE3 | TE2 | TF |
| | 5 A | | | PGA | TE3 | TE2 | TF |
| | B | | | PGA | TE3 | TE2 | TF |

14

RH dorsal cortex

| slide | section | | | | | | | |
|---|---|---|---|---|---|---|---|---|
| | 1A | | | | | | | |
| | B | | | | | | | |
| | C | | 3 | 1 | PE | PF | | |
| | D | | 23 | | PE | PF | | |
| | 2A | | 23 | 31 | PE | PF | | |
| | B | | 23 | 31 | PE | PFG | | |
| | C | | 23 | 31 | PE | PFG | | |
| | D | | 23 | 31 | PE | PFG | | |
| | 3A | | 23 | 31 | PE | AIP | PFG | |
| | B | | 23 | 31 | PE | AIP | PFG | |
| | C | | 23 | 31 | PE | AIP | PFG | |
| | D | | 23 | 31 | PE | AIP | PFG | |
| | 4A | | 23 | 31 | PEC | PE | AIP | PG |
| | B | | 23 | 31 | PEC | PE | LIP | PG |
| | C | | 23 | 31 | PEC | PE | LIP | PG |
| | D | | 23 | 31 | PEC | PE | LIP | PG |
| | 5A | | 23 | 31 | V6 | PEC | LIP | PG |
| | B | | 23 | 31 | V6 | PEC | LIP | OPT |
| | C | | 23 | 31 | V6 | PEC | LIP | OPT |
| | 6A | PGM | | | V6 | PEC | LIP | OPT |
| | B | PGM | A19 | | V6 | | LIP | OPT |

15

MIP 81

slides 5 B, C
6AB.

### RH visual cortex

| slide | section | | | | | | | | | | |
|---|---|---|---|---|---|---|---|---|---|---|---|
| | 1A | V1 fovea | | | | | | | | | |
| | B | v1 fovea | | | | | | | | | |
| | C | V1 fovea | | | | | | | | | |
| | 2A | v1 fovea | | | | | | | | | |
| | B | V1 fovea | | | | | | | | | |
| | C | v1 fovea | | | | | | | | | |
| | 3A | V1 calcrine | V1 fovea | | | | | | | | |
| | B | V1 calcrine | V1 fovea | | | | | | | | |
| | C | V1 calcrine | V1 fovea | | | | | | | | |
| | 4A | V1 calcrine | V1 fovea | | | | | | | | |
| | B | V1 calcrine | V1 fovea | | | | | | | | |
| | C | V1 calcrine | V1 fovea | | | | | | | | |
| | 5a | | | | | | | | | | |
| | b | V1 calcrine | v1OD' | V2 | V1OV | | | | | | |
| | c | V1 calcrine | v2D | v1OD | V2 V | V1OV | | | | | |
| | 6a | V1 calcrine | v2D | | V2 V | V1OV | | | | | |
| | b | V1 calcrine | v2D | | V2 V | V1OV | | | | | |
| | c | V1 calcrine | v2D | | V2 V | V1OV | | | | | |
| | 7a | V1 calcrine | v2D | | V2 V | | | | | | |
| | b | V1 calcrine | v2D | V3? | V2 V | | | | | | |
| | 8a | V1 calcrine | v2D | V3 | V2 V | | | | | | |
| | b | V1 calcrine | v2D | V3 | V2 V | | | | | | |
| | 9a | V1 calcrine | v2D | V3 | V2 V | | | | | | |
| | b | V1 calcrine | v2D | A19D(?) | V3 | V2 V | | | | | |
| | 10a | V1 calcrine | v2D | A19D | V3 | V2 V | | | | | |
| | b | V1 calcrine | v2D | A19D | V3 | V2 V | | | | | |
| | 11a | V1 calcrine | v2M | V6 | V2D | A19D | V3 | V2V | | | |
| | b | V1 calcrine | v2M | V6 | | A19D | V3 | V2V | | | |
| | 12a | V1 calcrine | v2M | V6 | | A19D | V3 | V2V | | | |
| | b | V1 calcrine | v2M | A19M | V6 | A19D | V3D | V4(?) | V3V | V2V | |
| | c | V1 calcrine | v2M | A19M | V6 | A19D | V3D | V4 | V3V | V2V | |
| | 13a | V1 calcrine | v2M | A19M | V6 | A19D | V3D | V4 | V3V' | V2V | |
| | b | V1 calcrine | v2M | A19M | V6 | A19D | V3D | V4 | V3V | V2V | |
| | c | V1 calcrine | v2M | A19M | V6 | A19D | V3D | V4 | V3V | V2V | |
| | 14a | V1 calcrine | v2M | A19M | V6 | V3A | VL | V4 | V3V | V2V | |
| | b | V1 calcrine | v2M | A19M | V6 | V3A | VL | V4 | TEO | V3V | V2V |
