## Supplementary material for "Conserved and specialized features of thalamocortical wiring revealed by single-cell projection mapping in mouse and marmoset": Supp. File 1: tissue printout.pdf

### LH PFC

| slide | section | callosum | | | | | | | | | | | | | | | | |
|---|---|---|---|---|---|---|---|---|---|---|---|---|---|---|---|---|---|---|
| 1 | A | | | | | | | | | | | | | | | | | |
| | B | | | | | | | | | | | | | | | | | |
| | C | | | | | | | | | | | | | | | | | |
| | D | | | | | | | | | | | | | | | | | |
| 2 | A | | 10M | 9 | | 10D | | 47 | 11 | 14 | | | | | | | | |
| | B | | 32 | 9 | 46D | 46V | | 47 | 11 | 13B/14 | | | | | | | | |
| | C | | 32 | 9 | 46D | 46V | | 47 | 11 | 13B/14 | | | | | | | | |
| 3 | A | | 32 | 9 | 46D | 46V | | 47 | 11 | 13B/14 | | | | | | | | |
| | b | | 32 | 9 | 46D | 46V | | 47 | 11 | 13B/14 | | | | | | | | |
| | c | | 32 | 9 | 8AD | 8AV | | 47 | 11 | 13B/14 | | | | | | | | |
| 4 | A | | 32 | 9 | 8AD | 8AV | | 47 | 11 | 13B/14 | | | | | | | | |
| | B | | 32 | 8B | 8AD | 8AV | | 47 | 11 | 13B/14 | | | | | | | | |
| | C | | 32 | 8B | 8AD | 8AV | | 47 | 11 | 13B/14 | | | | | | | | |
| 5 | a | | 32 | 8B | 8AD | 8AV | | 47 | 11 | 13B/14 | | | | | | | | |
| | b | | 32 | 8B | 8AD | 8AV | 45 | 47 | 13 | 13B/14 | | | | | | | | |
| | c | | 32 | 8B | 8AD | 8AV | 45 | 47 | 13 | 13B/14 | | | | | | | | |
| 6 | a | | | 8B | 6D | 8AV | 45 | 47 | 13 | 13B/14 | | | | | | | | |
| | b | | 24 | 8B | 6D | 8AV | 45 | ProM | 47 | 13 | 13B/14 | 25 | | | | | | |
| | c | | 24 | 8B | 6D | 8AV | 45 | ProM | 47 | 13 | 25 | | | | | | | |
| 7 | a | | 24 | 8B | 6D | 6v | | ProM | | 13 | 25 | | | | | | | |
| | b | | 24 | 6m | 6D | 8c | 6v | ProM | GU | Opro | OPAI | 25 | | | | | | |
| | c | | 24 | 6m | 6D | 8c | 6v | ProM | GU | OPro | OPAI | 25 | | | | | | |
| 8 | A | | 24 | 6m | 6D | 8C | 6V | ProM | GU | OPro | OPAI | 25 | | | | | | |
| | b | | 24 | 6m | 6D | 8C | 6V | ProM | GU | OPro | OPAI | 25 | | | | | | |
| | c | | 24 | 6m | 6D | 8C | 6V | ProM | GU | OPro | OPAI | 25 | | | | | | |
| 9 | A | | 24 | 6M | 6D | | 6V | ProM | GU | OPro | OPAI | | | | | | | |
| | b | | 24 | 6M | 6D | 4 | 3 | ProM | GU | OPro | OPAI | | | | | | | |
| | c | | 24 | 6M | 6D | 4 | 3 | ProM | GI | DI | AI | OPAI | | | | | | |
| 10 | A | | 24 | 6M | 6D | 4 | 3 | | GI | DI | AI | OPAI | | | | | | |
| | b | | 24 | | | 4 | 3 | | GI | DI | AI | | | STR | TE1 | ENT | | |
| | c | | 24 | | | 4 | 3 | | GI | DI | AI | | | STR | TE1 | ENT | | |
| 11 | a | | 24 | | | 4 | 3 | | GI | DI | AI | | | STR | TE1 | ENT | | |
| | b | | 24 | | | 4 | 3 | | GI | DI | AI | | | STR | TE1 | ENT | | |
| | c | | 24 | | | 4 | 3 | | GI | DI | AI | | | STR | TE1 | ENT | | |
| 12 | A | | | | | 4 | 3 | | GI | DI | | | | TPO | TE1 | ENT | | |
| | B | | | | | 4 | 3 | | GI | DI | | | | TPO | TE2 | TE1 | ENT | |
| | A | | | | | 4 | 3 | | GI | DI | | | | TPO | TE2 | TE1 | ENT | |
| 13 | A | | | | | 4 | 3 | | GI | DI | | | | TPO | TE3 | TE2 | TE1 | ENT |
| | B | | | | | 4 | 3 | | | | | | | TPO | TE3 | TE2 | TE1 | ENT |
| | C | | | | | | 3 | | | | | | | TPO | TE3 | TE2 | TE1 | ENT |
| 14 | A | | | | | | | | | | | | | TPO | TE3 | TE2 | TE1 | ENT |
| | B | | | | | | | | | | | | | TPO | TE3 | TE2 | TE1 | ENT |

### RH PFC

| slide | section | Callosum | area | area | | | | | | | | | sulcus | | | | | |
|---|---|---|---|---|---|---|---|---|---|---|---|---|---|---|---|---|---|---|
| 1 | a | | 10 | | | | | | | | | | | | | | | |
| | b | | 10 | | | | | | | | | | | | | | | |
| | c | | 10 | | | | | | | | | | | | | | | |
| | d | | 10 | | | | | | | | | | | | | | | |
| 2 | a | | 10M | 9 | | 10D | | 47 | 11 | 14 | | | | | | | | |
| | b | | 10M | 9 | 46D | 46V | | 47 | 11 | 13B/14 | | | | | | | | |
| | c | | 32 | 9 | 46D | 46V | | 47 | 11 | 13B/14 | | | | | | | | |
| 3 | A | | 32 | 9 | 46D | 46V | | 47 | 11 | 13B/14 | | | | | | | | |
| | B | | 32 | 9 | 46D | 46V | | 47 | 11 | 13B/14 | | | | | | | | |
| | C | | 32 | 9 | 8AD | 8AV | | 47 | 11 | 13B/14 | | | | | | | | |
| 4 | A | | 32 | | 8AD | 8AV | | 47 | 11 | 13B/14 | | | | | | | | |
| | B | | 32 | 8B | 8AD | 8AV | | 47 | 11 | 13B/14 | | | | | | | | |
| | C | | 32 | 8B | 8AD | 8AV | | 47 | 11 | 13B/14 | | | | | | | | |
| 5 | a | | 32 | 8B | 8AD | 8AV | | 47 | 11 | 13B/14 | | | | | | | | |
| | b | | 32 | 8B | 8AD | 8AV | 45 | 47 | 13 | 13B/14 | | | | | | | | |
| 6 | a | | | 8B | 8AD | 6D | 8AV | | | | | | | | | | | |
| | b | | 24 | 8B | 6D | 8AV | 45 | 47 | 13 | 13B/14 | 25 | | | | | | | |
| | c | | 24 | 8B | 6D | 8AV | 45 | ProM | 47 | 13 | 25 | | | | | | | |
| 7 | a | | 24 | 8B | 6D | 6v | | ProM | | 13 | 25 | | | | | | | |
| | b | | 24 | 6m | 6D | 6v | | ProM | GU | Opro | 25 | | | | | | | |
| | c | | 24 | 6m | 6D | 6v | | ProM | GU | OPro | 25 | | | | | | | |
| 8 | A | | 24 | 6m | 6D | 8C | 6V | ProM | GU | OPro | OPAI | 25 | | | | | | |
| | b | | 24 | 6m | 6D | 8C | 6V | ProM | GU | OPro | OPAI | 25 | | | | | | |
| | c | | 24 | 6m | 6D | 8C | 6V | ProM | GU | OPro | OPAI | | | | | | | |
| 9 | A | | 24 | 6M | 6D | 8C | 6V | ProM | GU | OPro | OPAI | | | | | | | |
| | b | | 24 | 6M | 6D | 4 | | ProM | GU | OPro | OPAI | | | | | | | |
| 10 | A | | 24 | 6M | 6D | 4 | 3 | ProM | GI | DI | AI | OPAI | | | | | | |
| | B | | 24 | | | 4 | 3 | | GI | DI | AI | | | STR | TE1 | ENT | | |
| | C | | 24 | | | 4 | 3 | | GI | DI | AI | | | STR | TE1 | ENT | | |
| 11 | A | | 24 | | | 4 | 3 | | GI | DI | AI | | | STR | TE1 | ENT | | |
| | B | | 24 | | | 4 | 3 | | GI | DI | AI | | | STR | TE1 | ENT | | |
| 12 | A | | 24 | | | 4 | 3 | | GI | DI | AI | | | STR | TE1 | ENT | | |
| | B | | | | | 4 | 3 | | | | | | | STR | TE1 | ENT | | |
| 13 | A | | | | | 4 | 3 | | GI | DI | | | | TPO | TE2 | TE1 | ENT | |
| | B | | | | | 4 | 3 | | GI | DI | | | | TPO | TE2 | TE1 | ENT | |
| 14 | A | | | | | 4 | 3 | | GI | DI | | | | TPO | TE3 | TE2 | TE1 | ENT |

### RH lateral cortex

| slide | section | | | | | | |
|---|---|---|---|---|---|---|---|
| 1 | a | MT | | TEO | TFO | | |
| | b | MT | TE3 | TEO | | | |
| | c | MT | TE3 | TEO | | | |
| | d | MT | TE3 | TEO | TFO | | |
| | e | MT | TE3 | TEO | TFO | | |
| 2 | A | MT | FST | TE3 | TEO | TFO | |
| | B | MT | FST | TE3 | TEO | TFO | |
| | C | MT | FST | TE3 | TEO | TFO | |
| | D | MT | FST | TE3 | TE2 | TF | |
| 3 | A | MT | FST | TE3 | TE2 | TF | |
| | B | MT | FST | TE3 | TE2 | TF | |
| | C | MT | FST | TE3 | TE2 | TF | |
| | D | MT | FST | TE3 | TE2 | TF | |
| 4 | A | FST | PGA | TE3 | TE2 | TF | |
| | B | MST | FST | PGA | TE3 | TE2 | TF |
| | C | MST | FST | PGA | TE3 | TE2 | TF |
| | D | | | PGA | TE3 | TE2 | TF |
| 5 | A | | | PGA | TE3 | TE2 | TF |
| | B | | | PGA | TE3 | TE2 | TF |

### RH dorsal cortex

| slide | section | | | | | | |
|---|---|---|---|---|---|---|---|
| 1 | A | | | | | | |
| | B | | | | | | |
| | C | 3 | 1 | PE | PF | | |
| | D | 23 | | PE | PF | | |
| 2 | A | 23 | 31 | PE | PF | | |
| | B | 23 | 31 | PE | PFG | | |
| | C | 23 | 31 | PE | PFG | | |
| | D | 23 | 31 | PE | PFG | | |
| 3 | A | 23 | 31 | PE | AIP | PFG | |
| | B | 23 | 31 | PE | AIP | PFG | |
| | C | 23 | 31 | PE | AIP | PFG | |
| | D | 23 | 31 | PE | AIP | PFG | |
| 4 | A | 23 | 31 | PEC | PE | AIP | PG |
| | B | 23 | 31 | PEC | PE | LIP | PG |
| | C | 23 | 31 | PEC | PE | LIP | PG |
| | D | 23 | 31 | PEC | PE | LIP | PG |
| 5 | A | 23 | 31 | V6 | PEC | LIP | PG |
| | B | 23 | 31 | V6 | PEC | LIP | OPT |
| | C | 23 | 31 | V6 | PEC | LIP | OPT |
| 6 | A | PGM | | V6 | PEC | LIP | OPT |
| | B | PGM | A19 | V6 | | LIP | OPT |

### RH visual cortex

| slide | section | | | | | | | | | | |
|---|---|---|---|---|---|---|---|---|---|---|---|
| 1 | A | V1 fovea | | | | | | | | | |
| | B | v1 fovea | | | | | | | | | |
| | C | V1 fovea | | | | | | | | | |
| 2 | A | v1 fovea | | | | | | | | | |
| | B | V1 fovea | | | | | | | | | |
| | C | v1 fovea | | | | | | | | | |
| 3 | A | V1 calcrine | V1 fovea | | | | | | | | |
| | B | V1 calcrine | V1 fovea | | | | | | | | |
| | C | V1 calcrine | V1 fovea | | | | | | | | |
| 4 | A | V1 calcrine | V1 fovea | | | | | | | | |
| | B | V1 calcrine | V1 fovea | | | | | | | | |
| | C | V1 calcrine | V1 fovea | | | | | | | | |
| 5 | a | | | | | | | | | | |
| | b | V1 calcrine | v1OD | V2 | V1OV | | | | | | |
| | c | V1 calcrine | v2D | v1OD | V2 V | V1OV | | | | | |
| 6 | a | V1 calcrine | v2D | | V2 V | V1OV | | | | | |
| | b | V1 calcrine | v2D | | V2 V | V1OV | | | | | |
| | c | V1 calcrine | v2D | | V2 V | V1OV | | | | | |
| 7 | a | V1 calcrine | v2D | | V2 V | | | | | | |
| | b | V1 calcrine | v2D | V3? | V2 V | | | | | | |
| 8 | a | V1 calcrine | v2D | V3 | V2 V | | | | | | |
| | b | V1 calcrine | v2D | V3 | V2 V | | | | | | |
| 9 | a | V1 calcrine | v2D | V3 | V2 V | | | | | | |
| | b | V1 calcrine | v2D | A19D(?) | V3 | V2 V | | | | | |
| 10 | a | V1 calcrine | v2D | A19D | V3 | V2 V | | | | | |
| | b | V1 calcrine | v2D | A19D | V3 | V2 V | | | | | |
| 11 | a | V1 calcrine | v2M | V6 | V2D | A19D | V3 | V2V | | | |
| | b | V1 calcrine | v2M | V6 | | A19D | V3 | V2V | | | |
| 12 | a | V1 calcrine | v2M | V6 | | A19D | V3 | V2V | | | |
| | b | V1 calcrine | v2M | A19M | V6 | A19D | V3D | V4(?) | V3V | V2V | |
| | c | V1 calcrine | v2M | A19M | V6 | A19D | V3D | V4 | V3V | V2V | |
| 13 | a | V1 calcrine | v2M | A19M | V6 | A19D | V3D | V4 | V3V | V2V | |
| | b | V1 calcrine | v2M | A19M | V6 | A19D | V3D | V4 | V3V | V2V | |
| | c | V1 calcrine | v2M | A19M | V6 | A19D | V3D | V4 | V3V | V2V | |
| 14 | a | V1 calcrine | v2M | A19M | V6 | V3A | VL | V4 | V3V | V2V | |
| | b | V1 calcrine | v2M | A19M | V6 | V3A | VL | V4 | TEO | V3V | V2V |
