## Supplementary material for "Conserved and specialized features of thalamocortical wiring revealed by single-cell projection mapping in mouse and marmoset": Supp. File 1: 818353_dissection_annotation.pdf

Sectioned from anterior : left side is right side

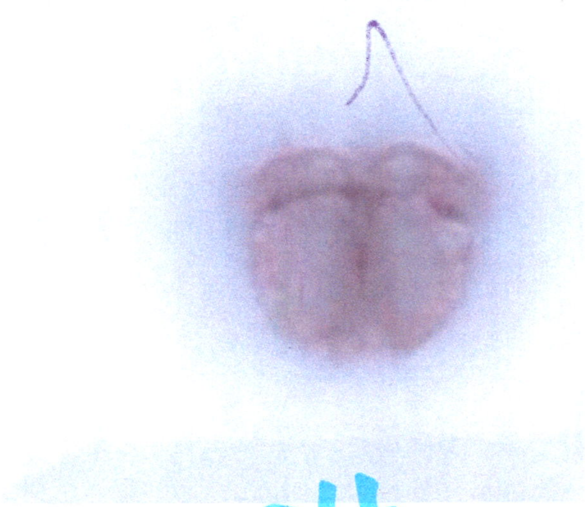

24

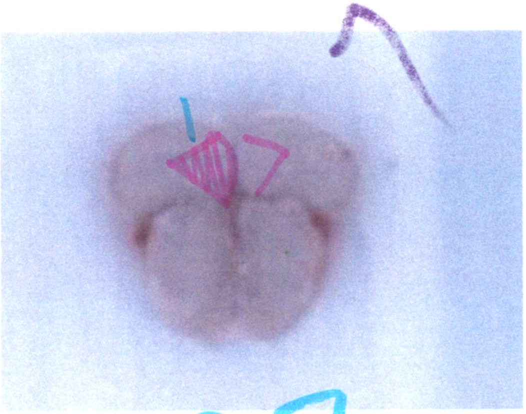

27

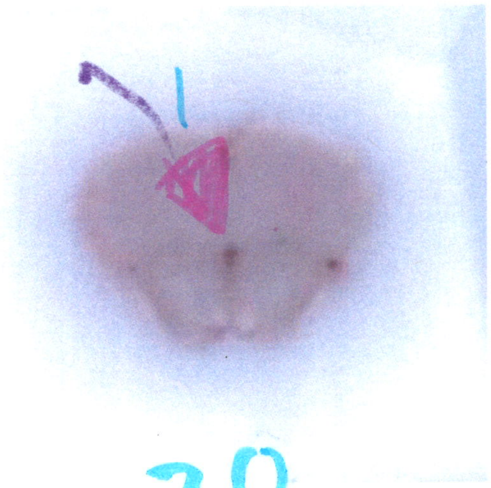

30

Sectioned from anterior : left side is right side

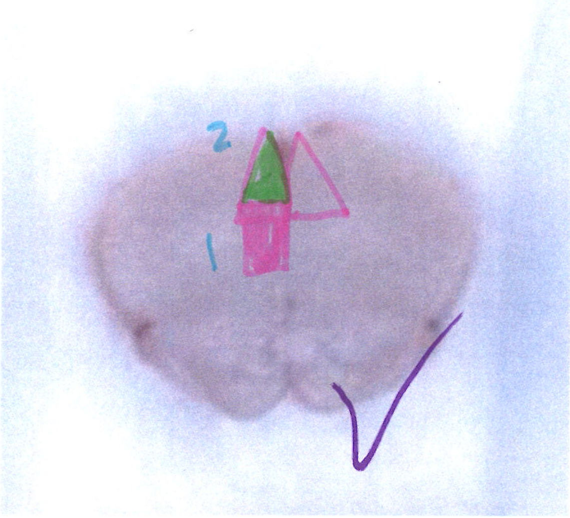


33

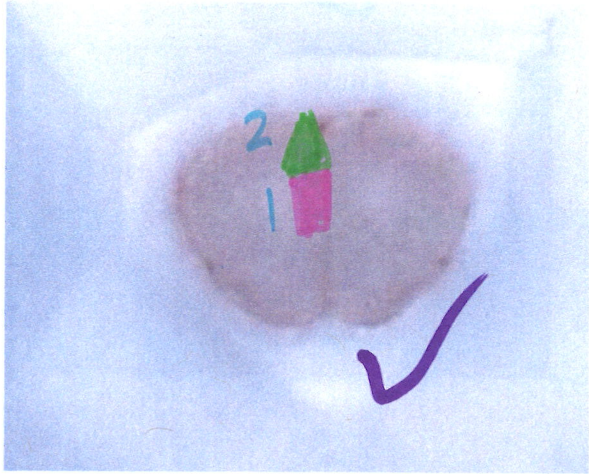


36

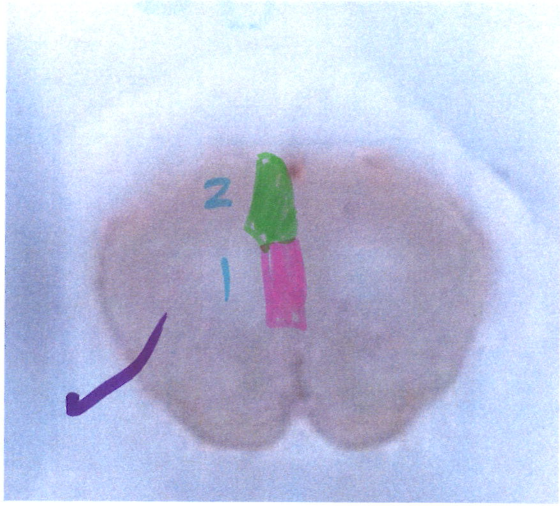


39

Sectioned from anterior : left side is right side

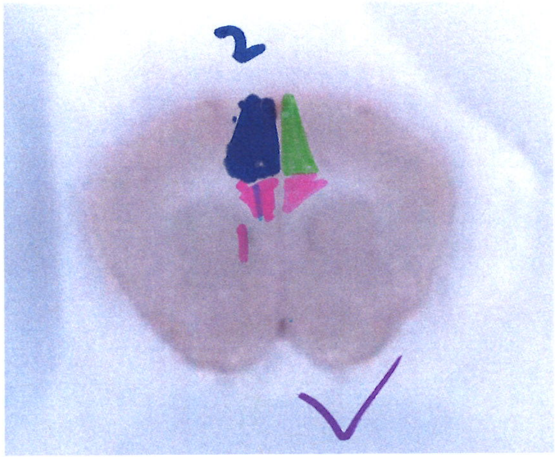


42

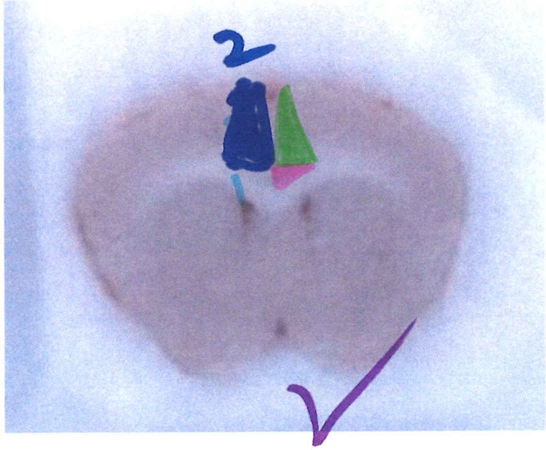


45
callosum connects

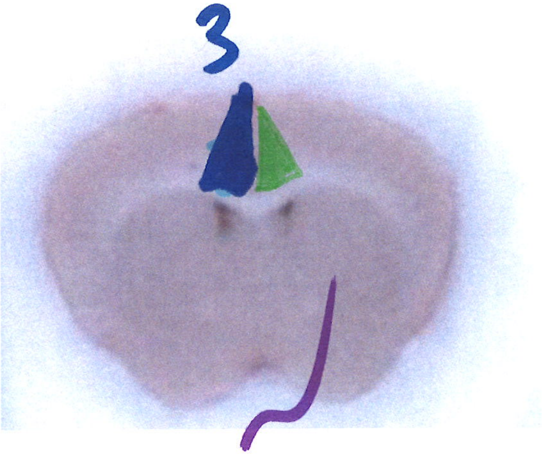


48

Sectioned from anterior : left side is right side

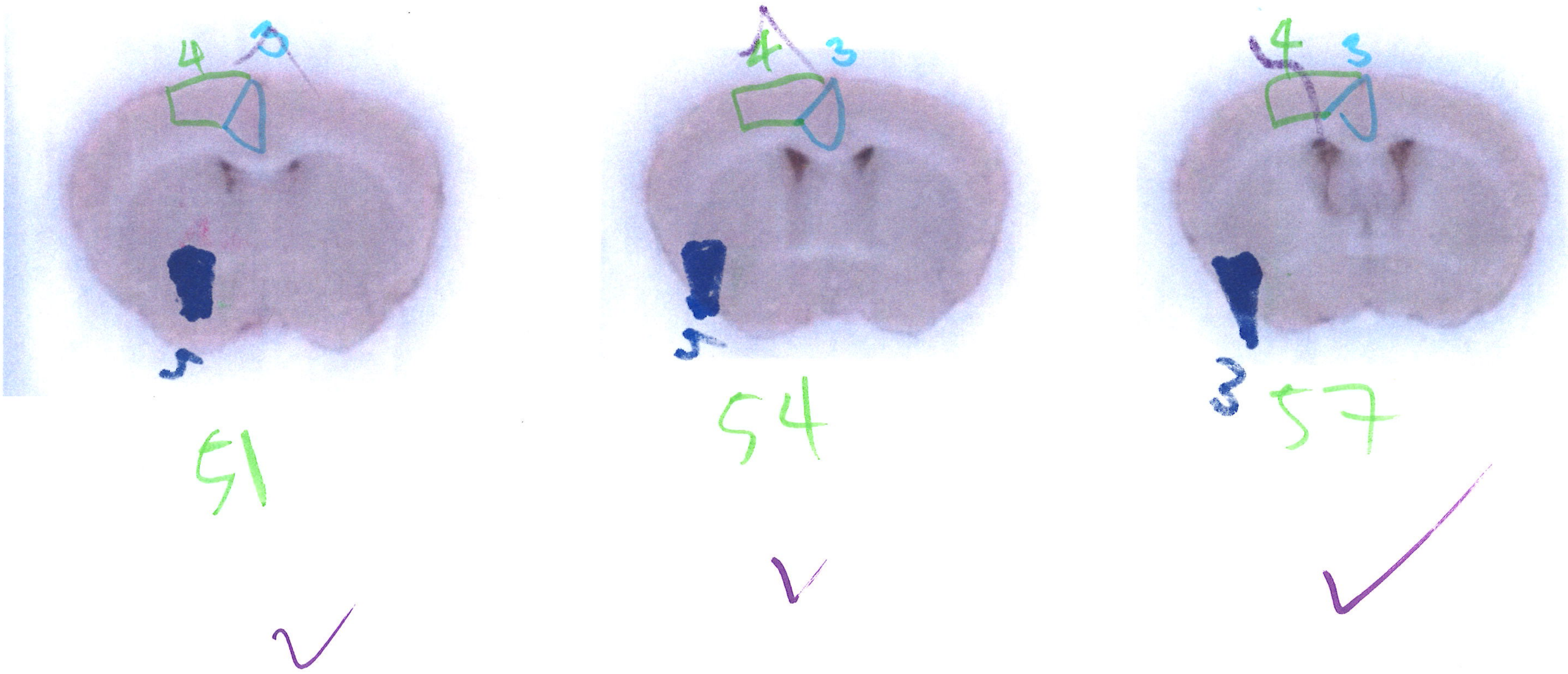

[illegible]

Sectioned from anterior : left side is right side

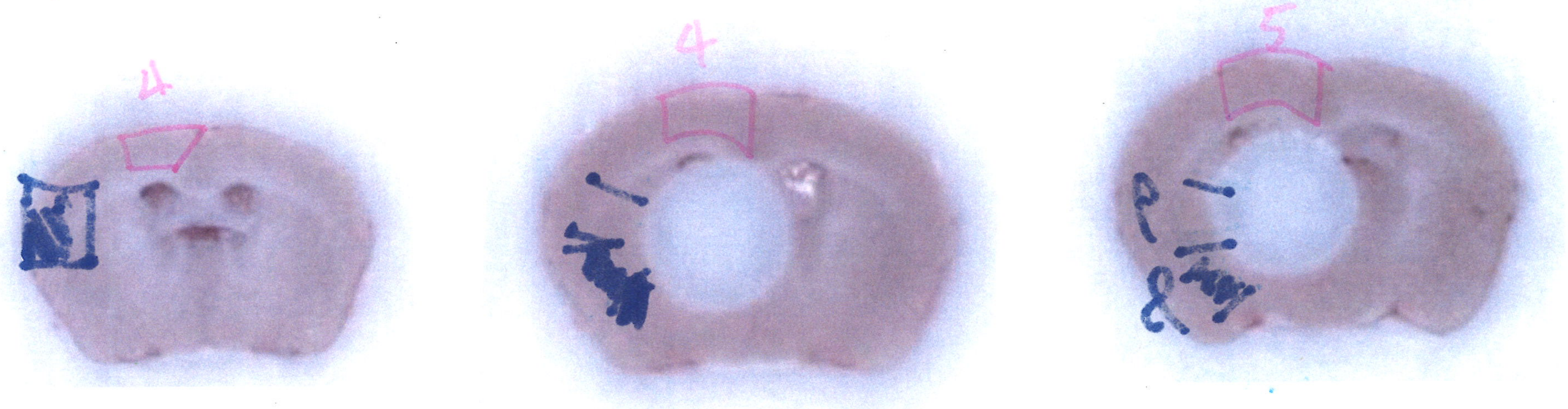


60 ✓ 63 ✓ 67

Sectioned from anterior : left side is right side

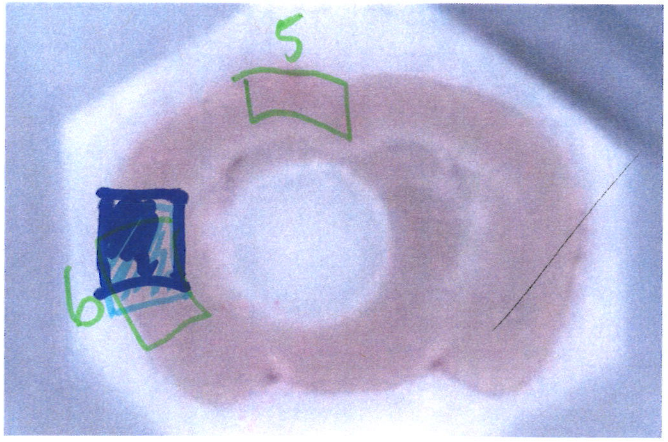


69

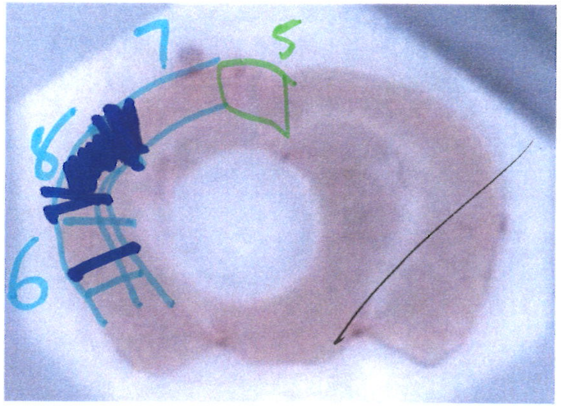


72

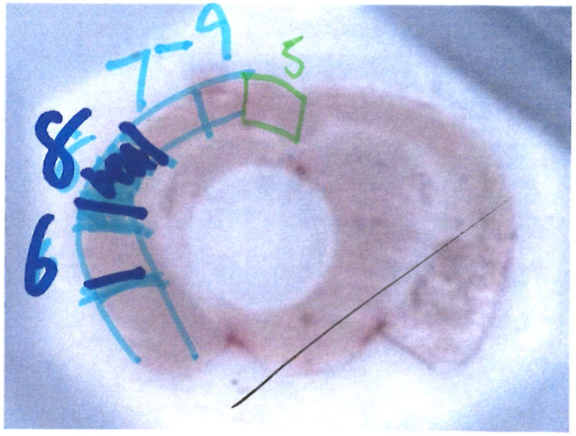


75

Notch appears

Sectioned from anterior : left side is right side

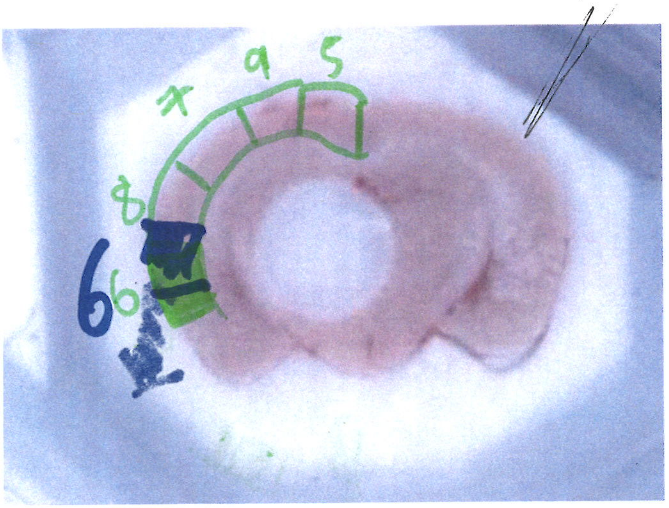


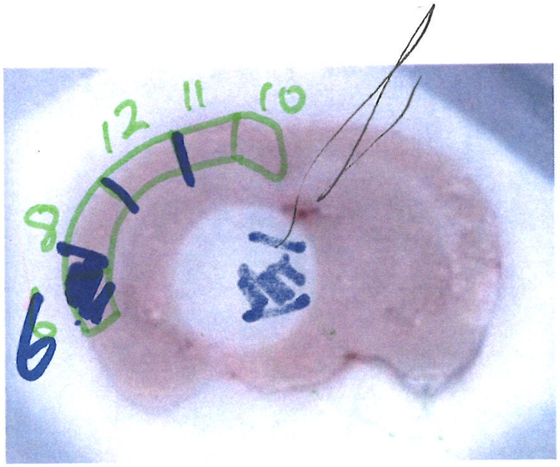


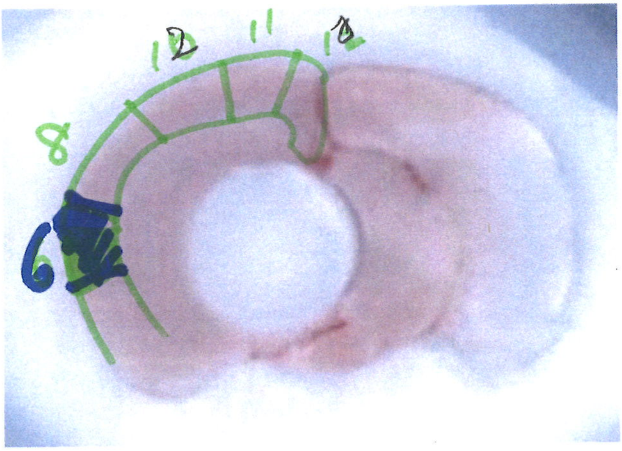


callosum connects:

78

81

84

Sectioned from anterior : left side is right side

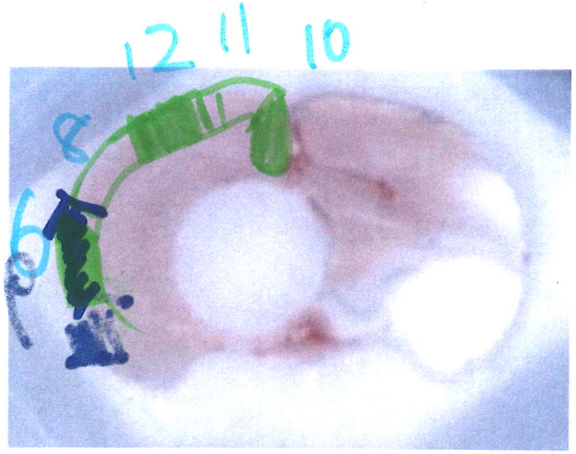


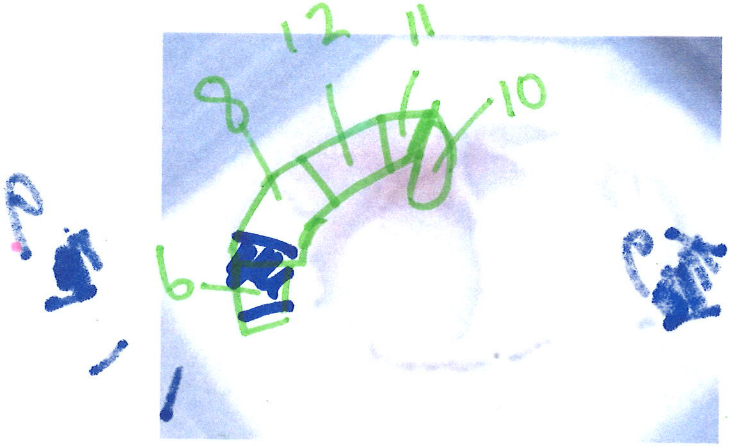


87

90

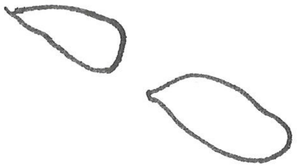

Sectioned from posterior : left side is left side

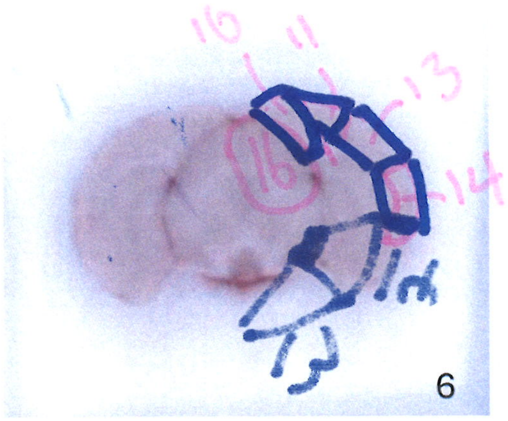


93

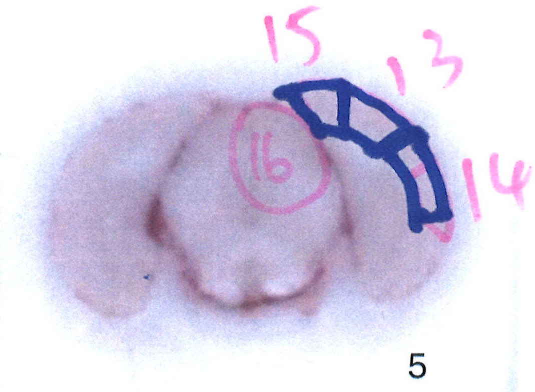


96

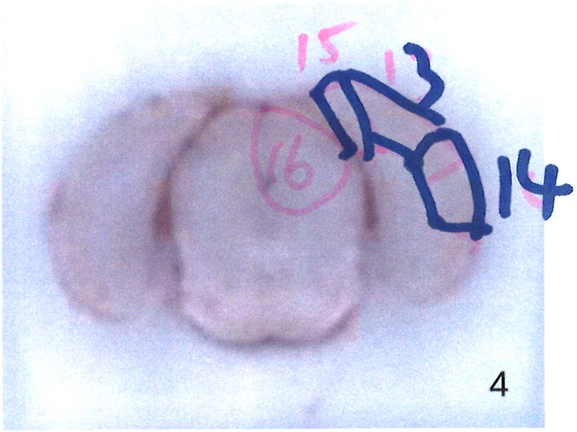


99

Sectioned from posterior : left side is left side

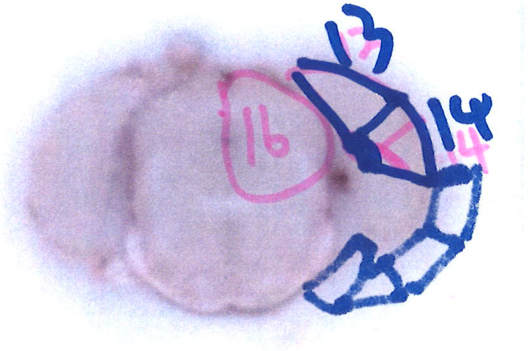


3

102

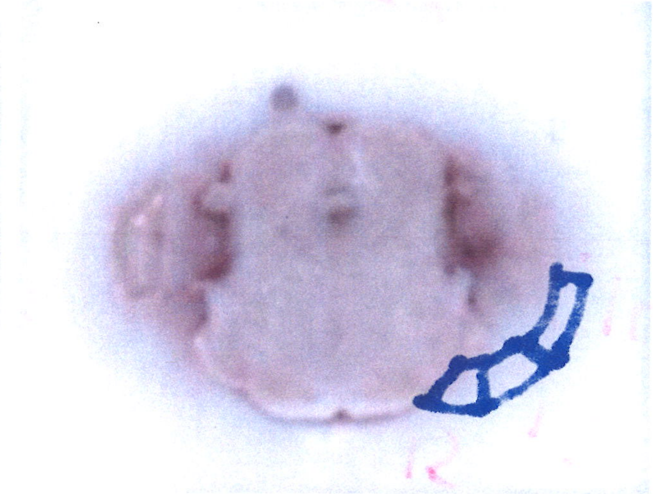

2

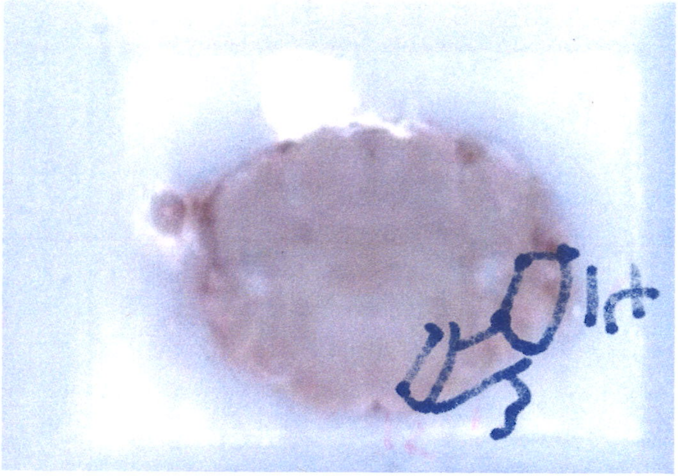


1
