## Supplementary material for "Conserved and specialized features of thalamocortical wiring revealed by single-cell projection mapping in mouse and marmoset": Supp. File 1: 839168_dissection_annotation.pdf

839168_11

CL40

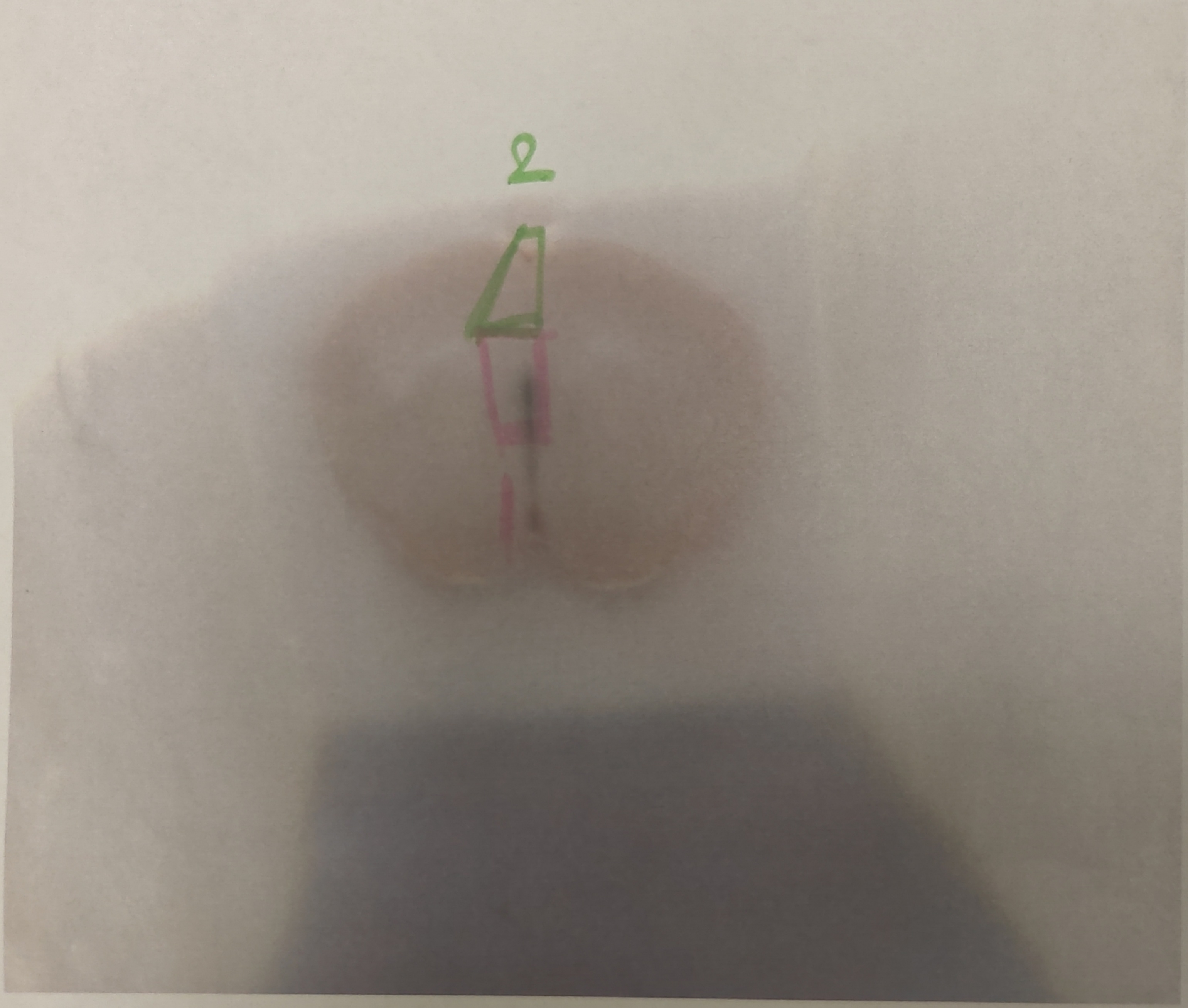


Preceeding 2 sections

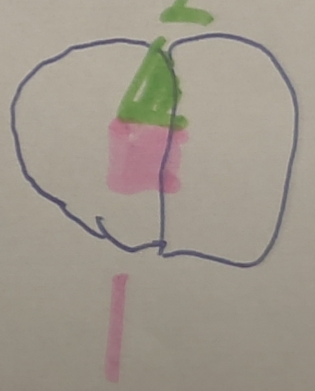


Preceeding ~~[illegible]~~ 3-4 sections

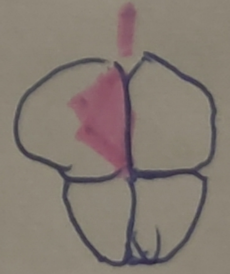

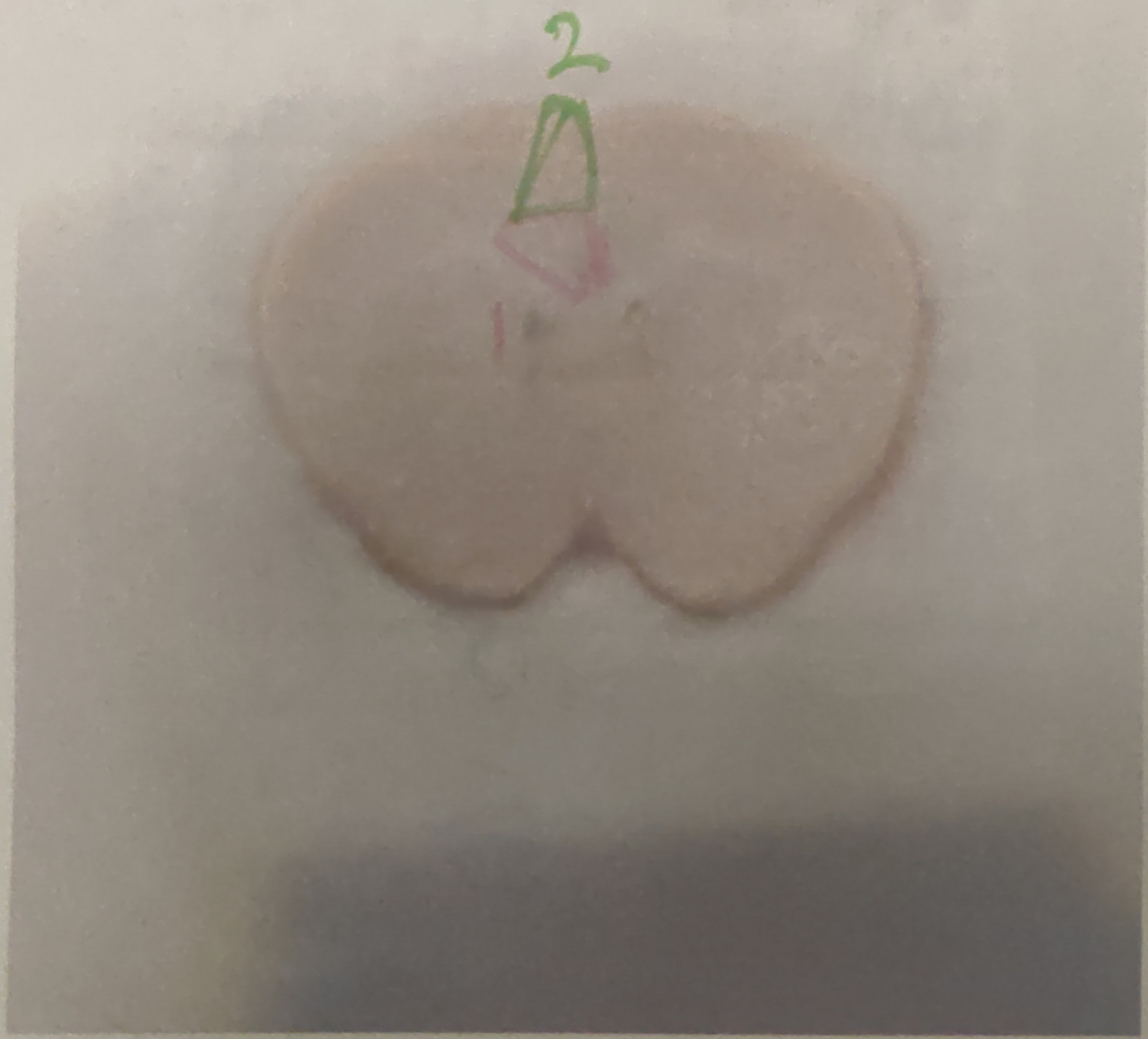


839168_21

*CL43

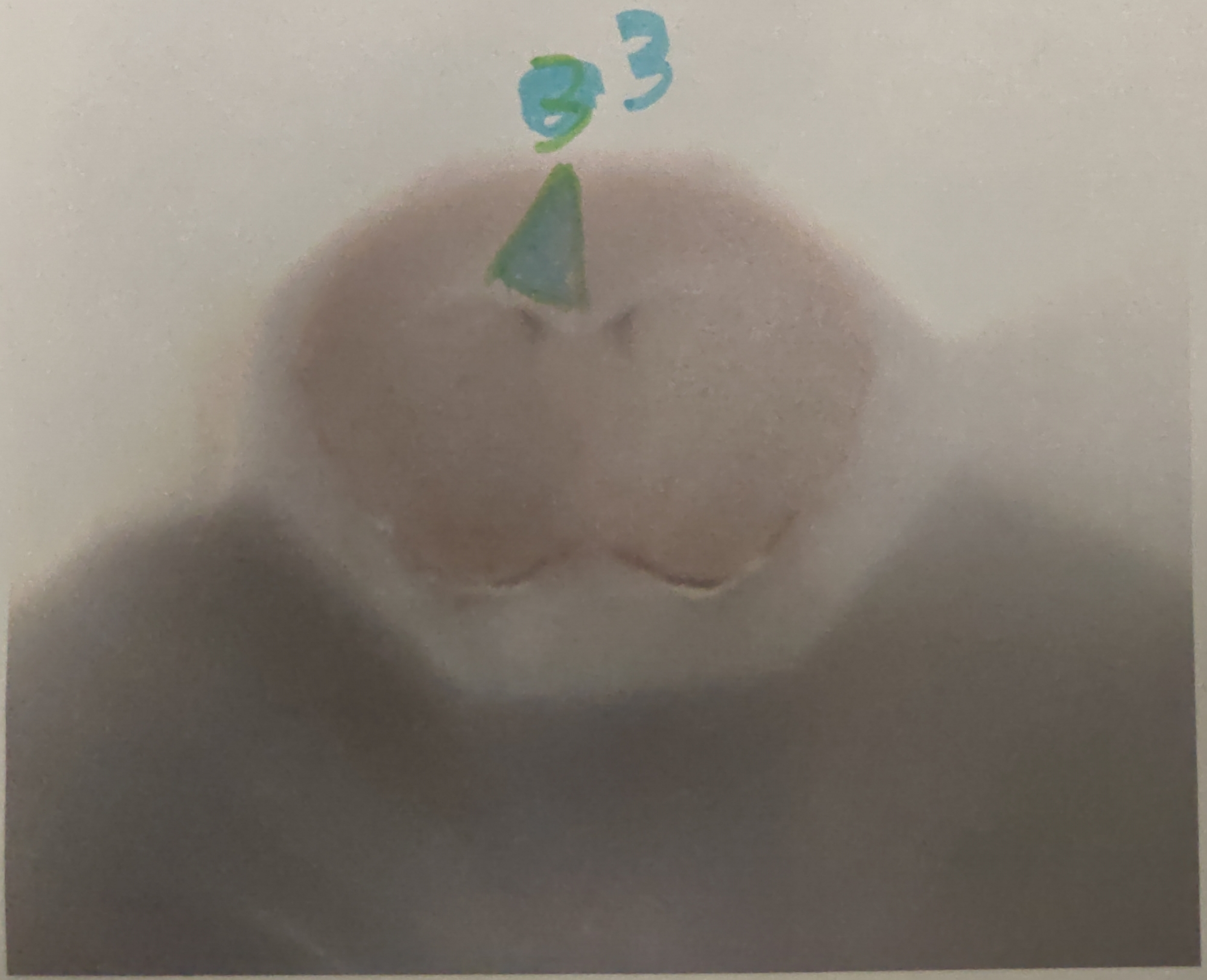


839168_3 1

CL47

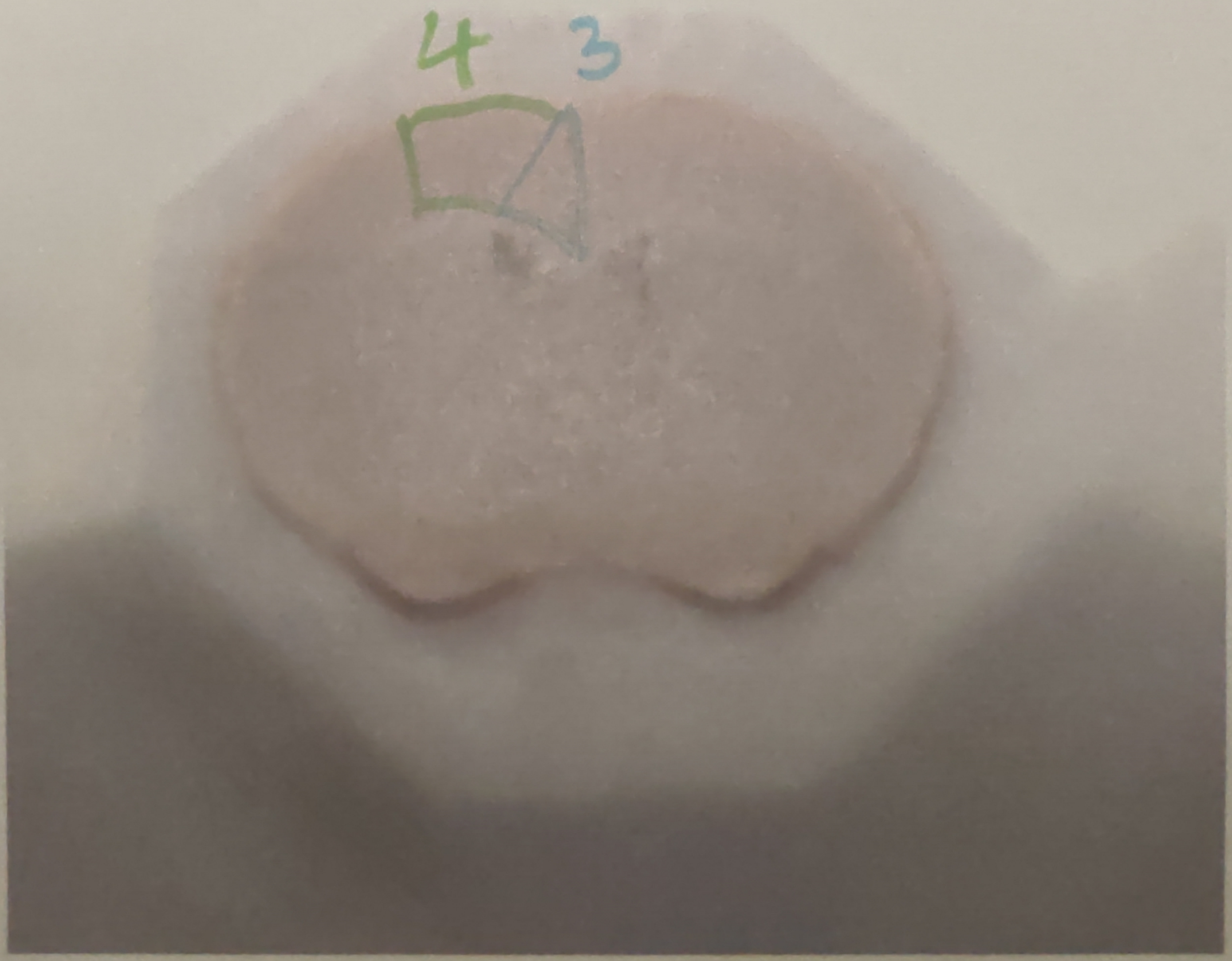


839168_4

CL50

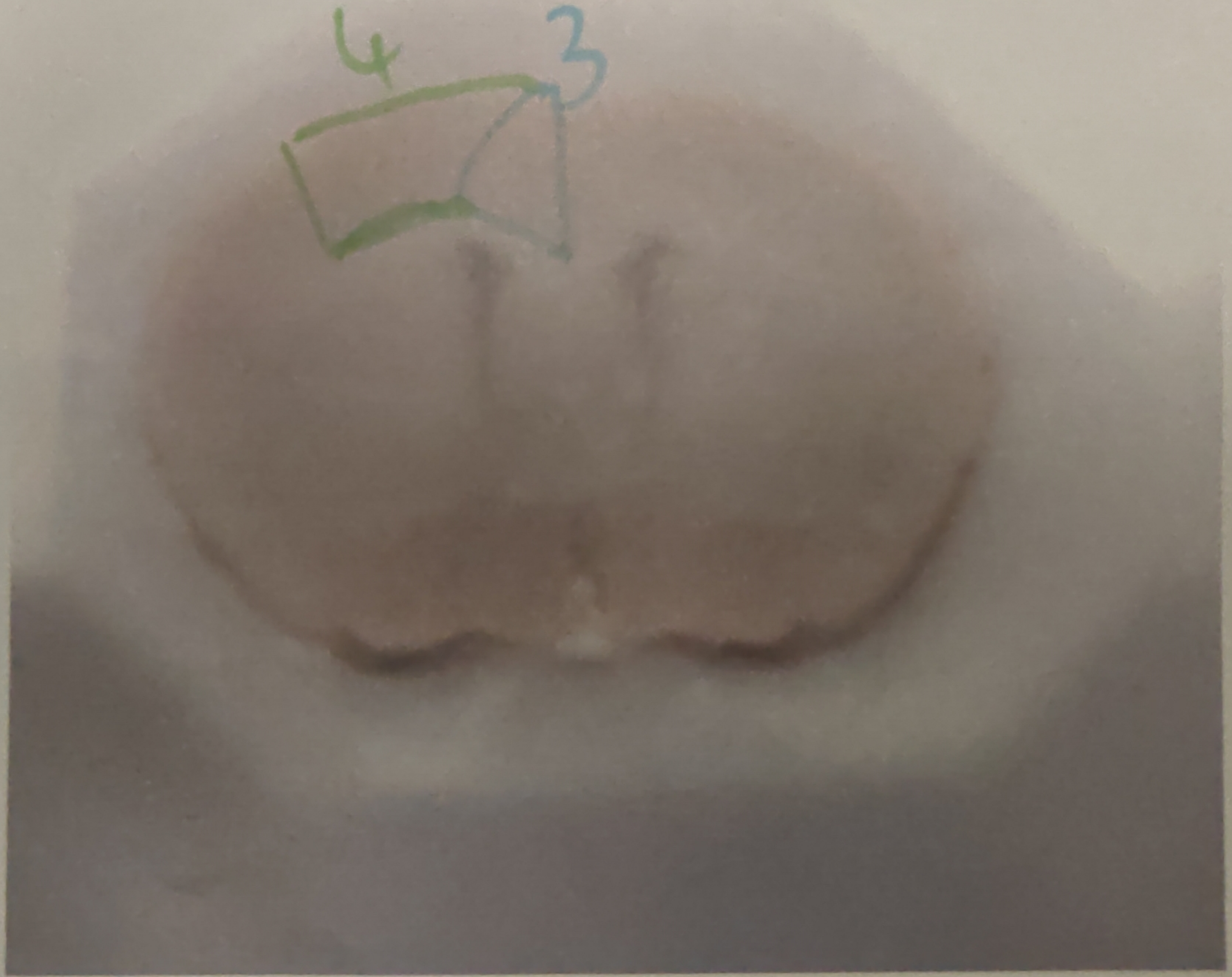


839168_5

CL53

Last section
from anterior

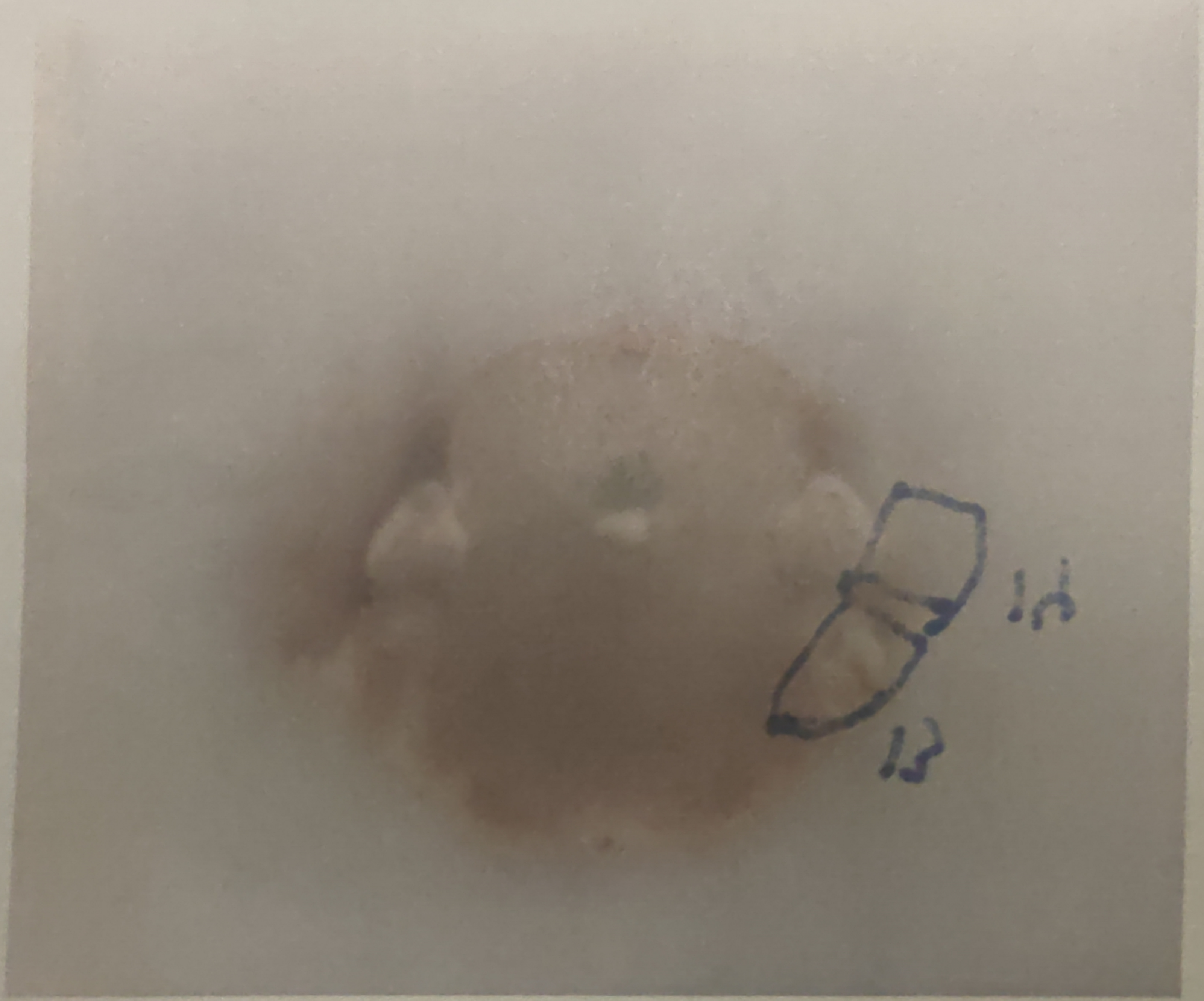


839168_2.1

CL 107

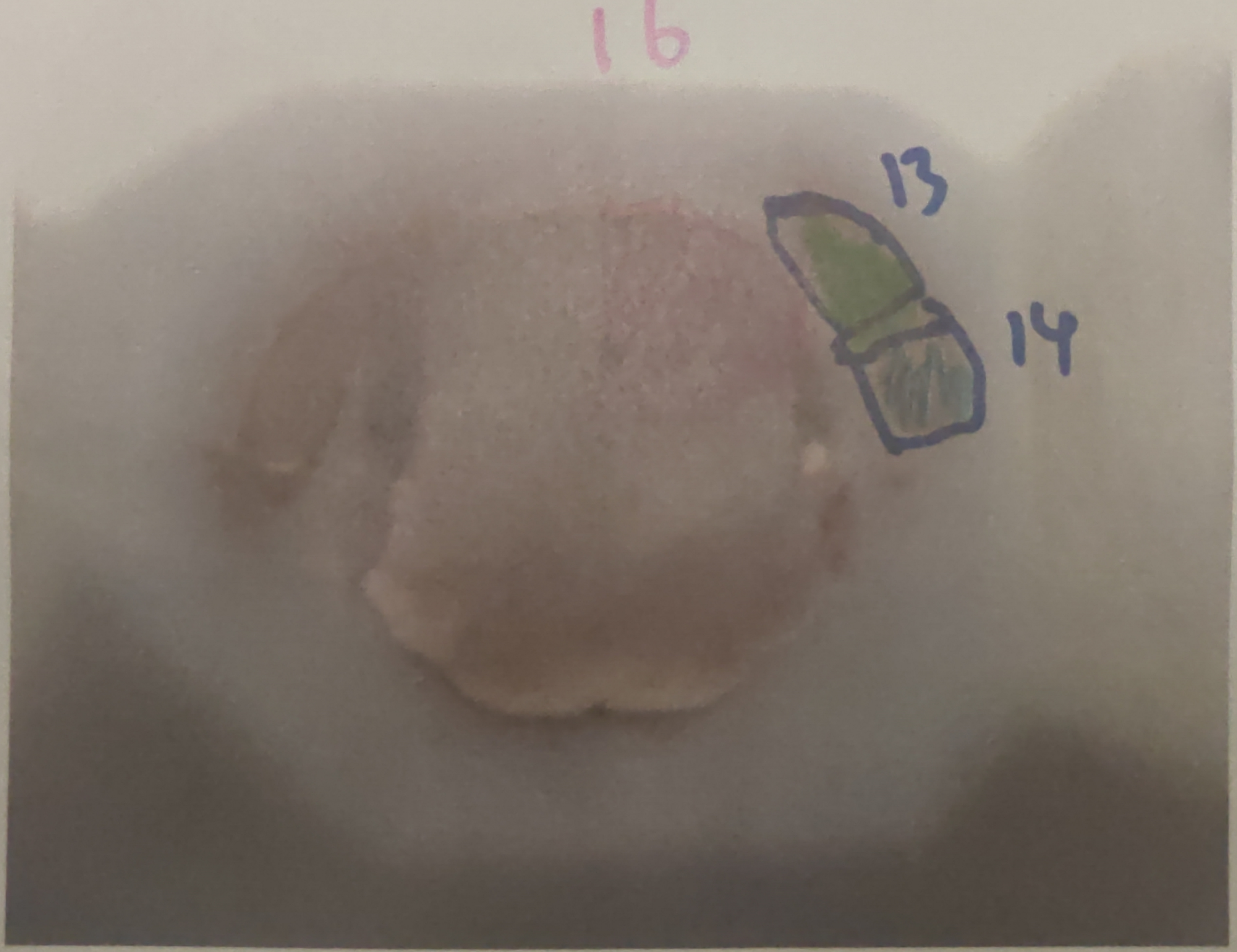


839168_2.2
1.tif

☆ CL[illegible] 103

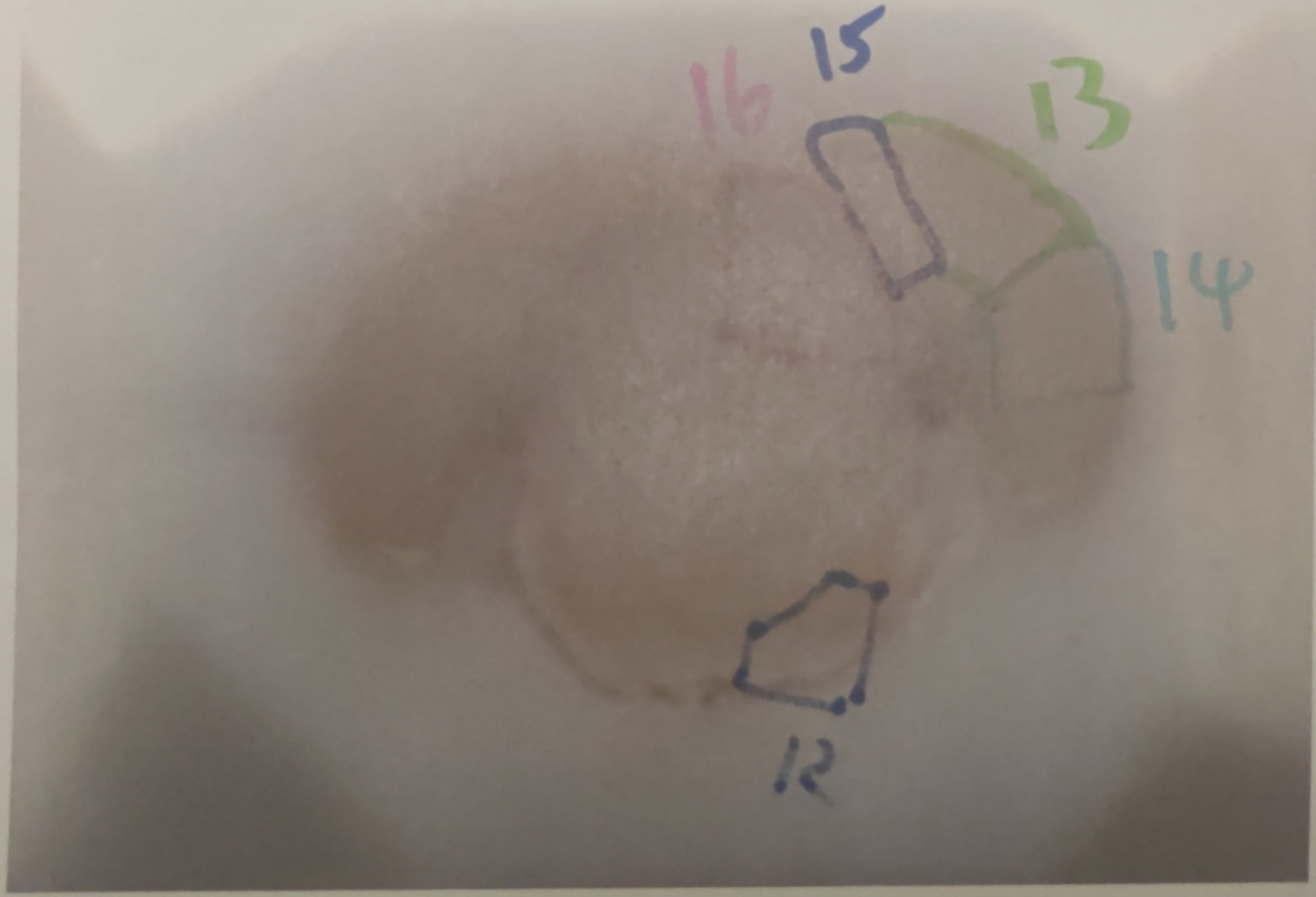


2.3

~~CL107 100?~~

CL95?

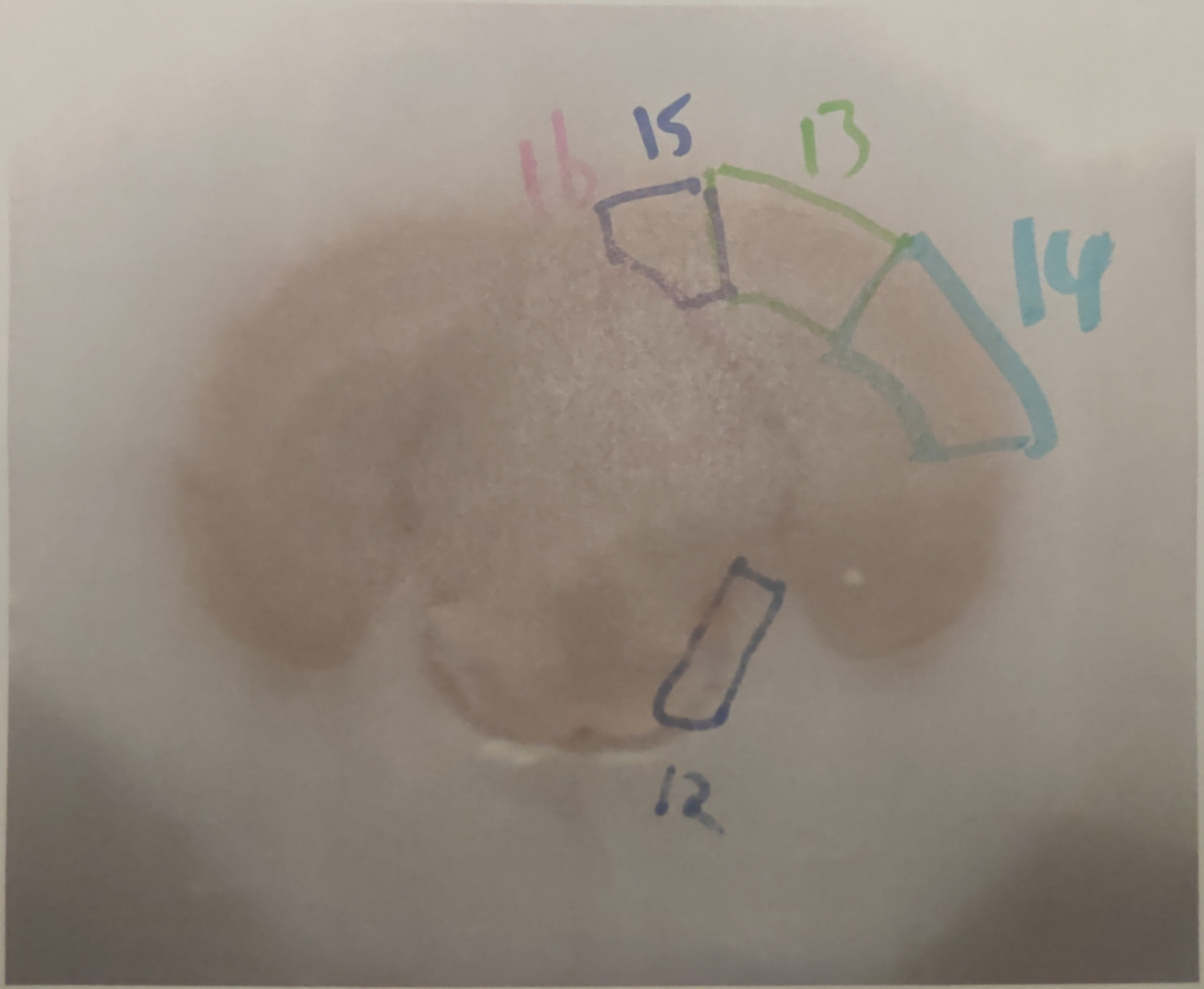


839168_2.4

* CL95

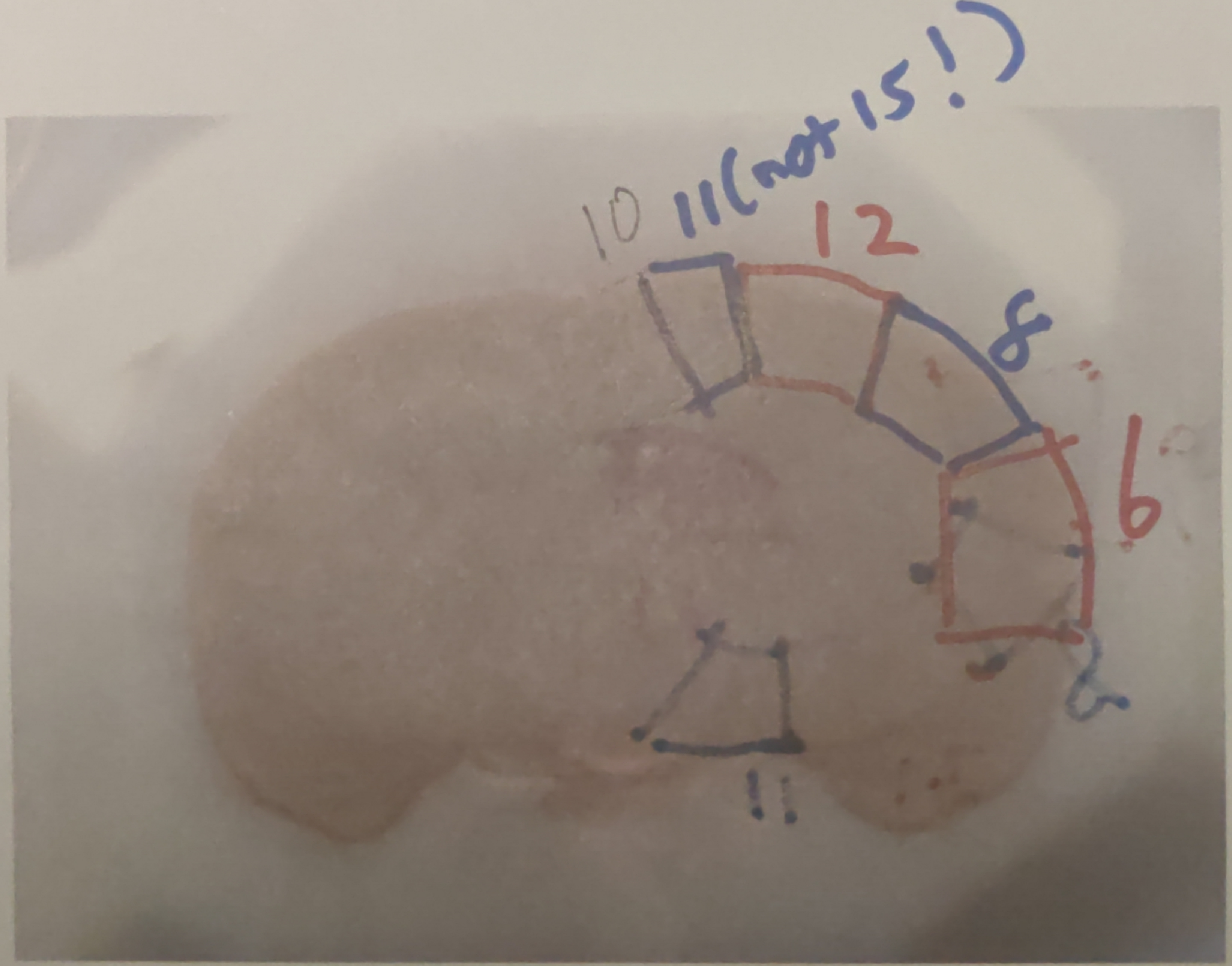


839168_2.5

* CL 89?

Note the punched slices are collected
P→A not A→P (as in 1st brain)
This causes 300μm shift in corresponding
slice images

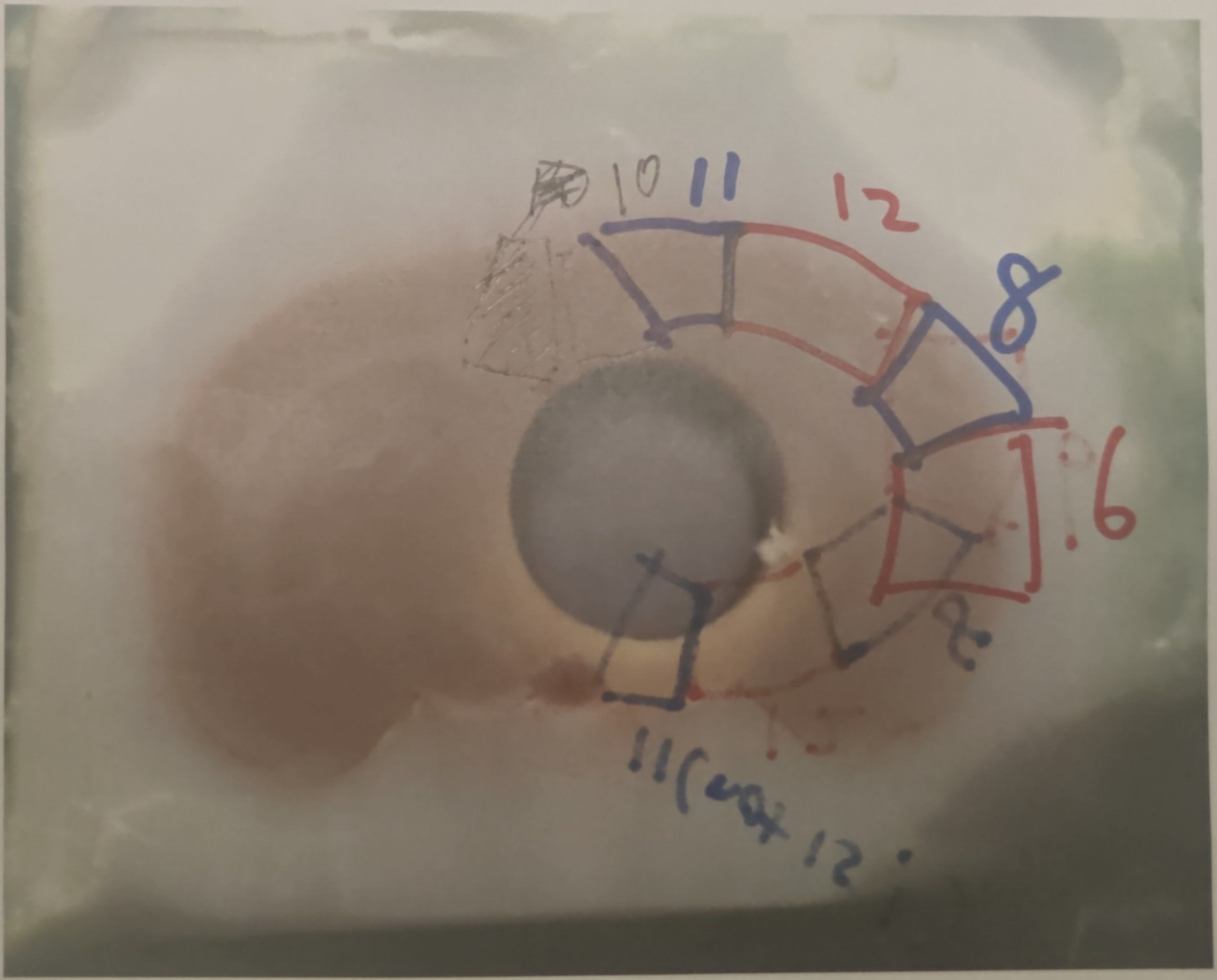


839168_3.1

*CL86?

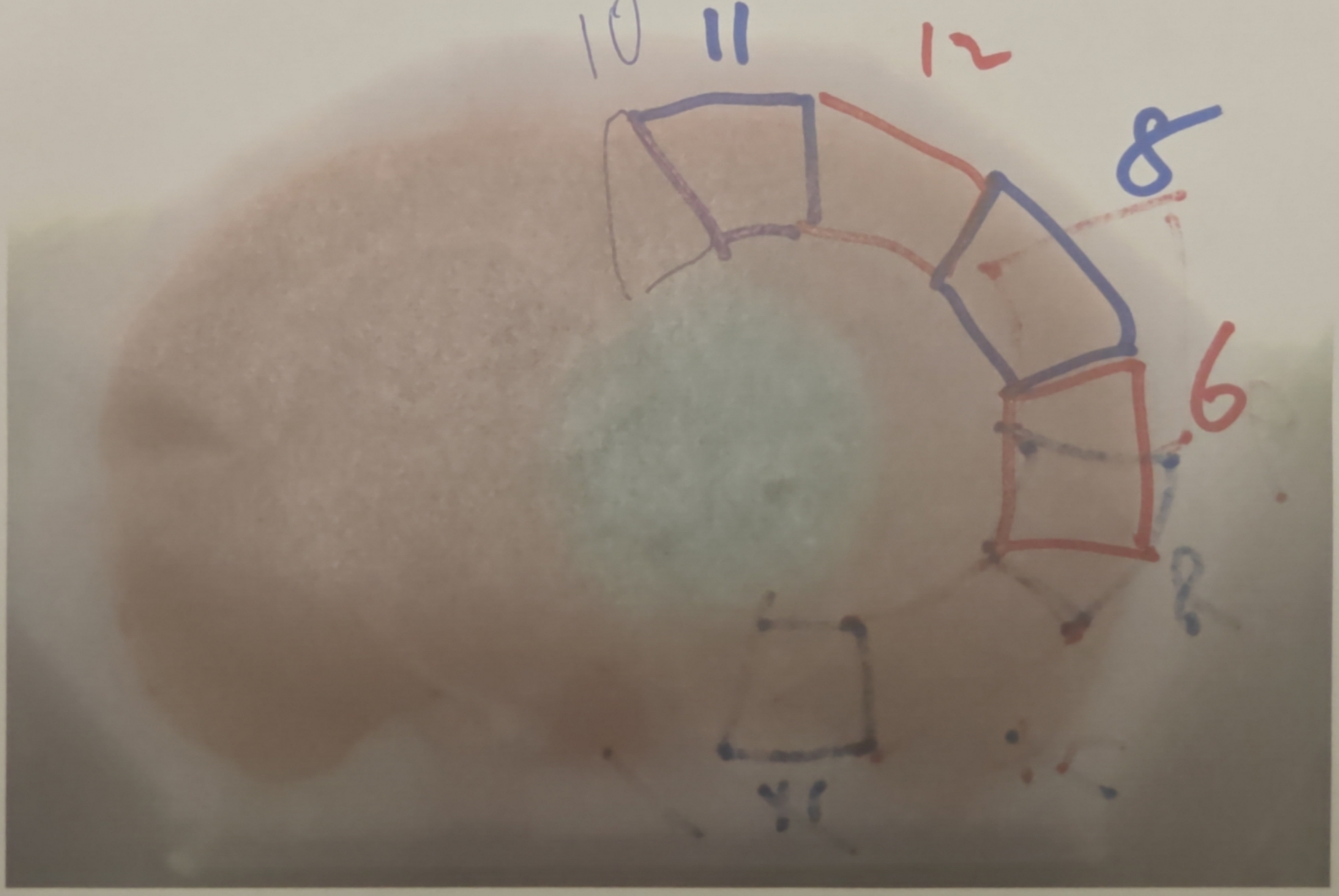


3.2

CL87

→ cpce

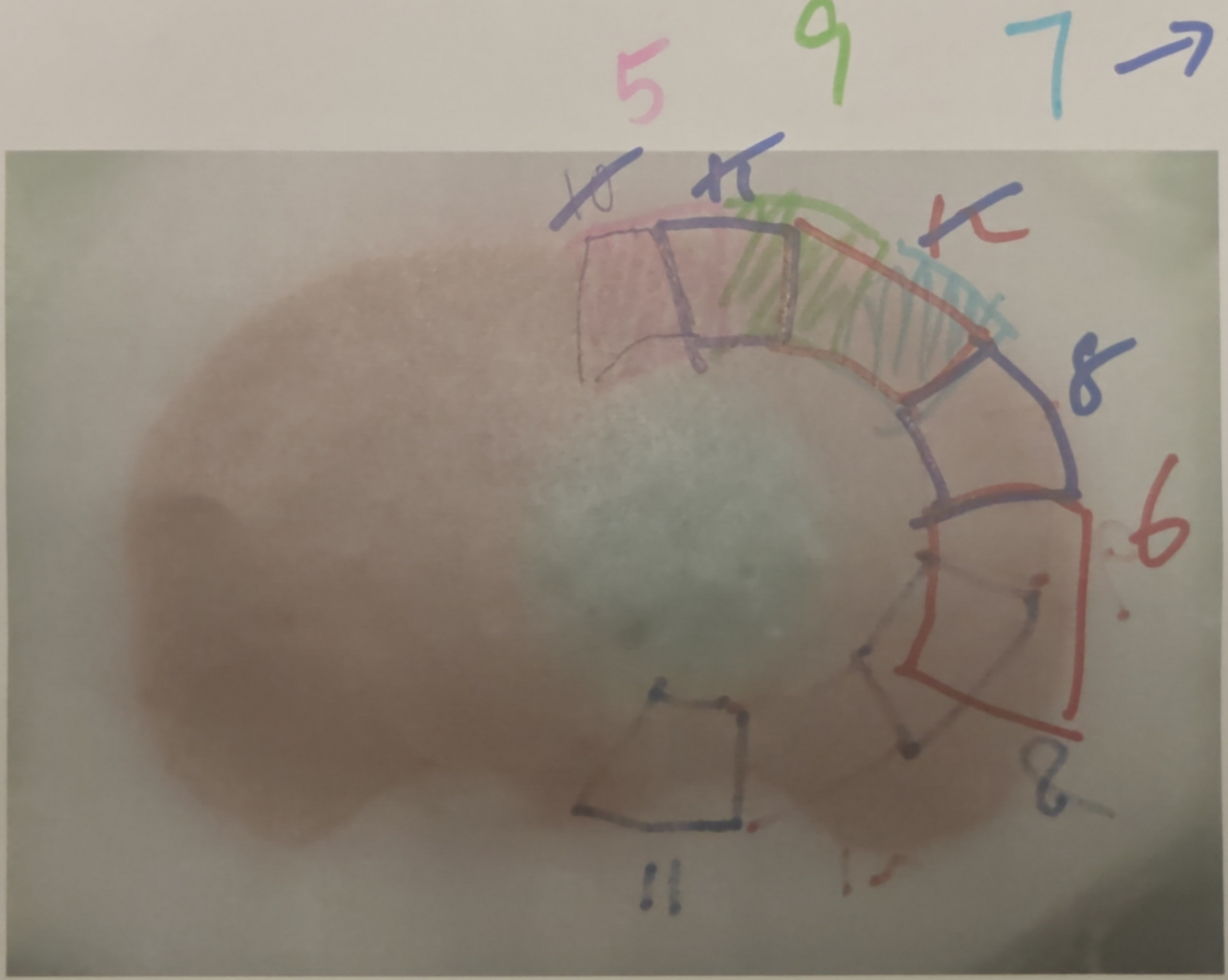


core → because P→A, corresponds to CL 77 in first brain

3.3

CL 80

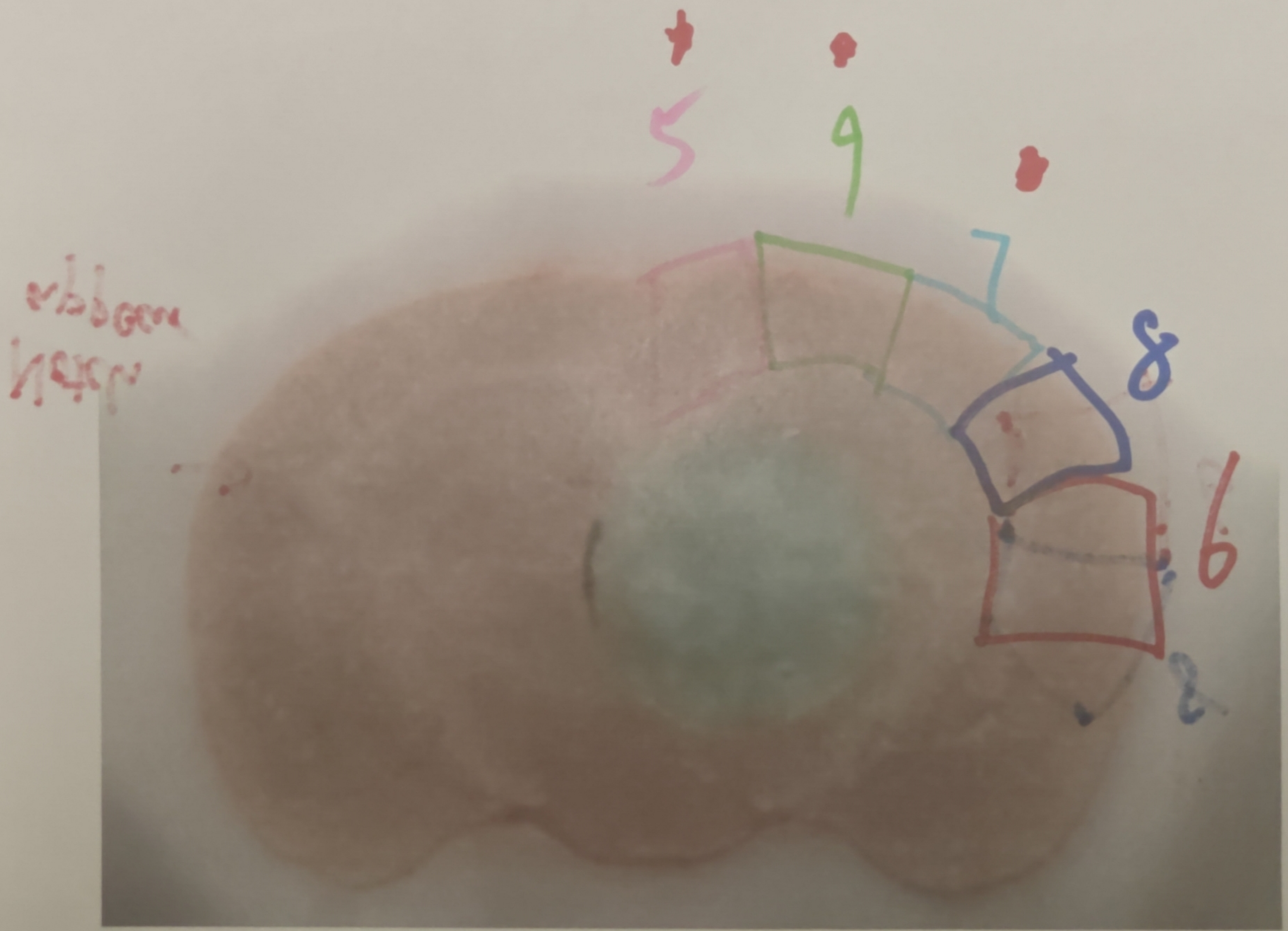


839168_3.4

→ because this is P→A, so correspond to CL71 in previous brain

3.5

*CL74

Notch appear →

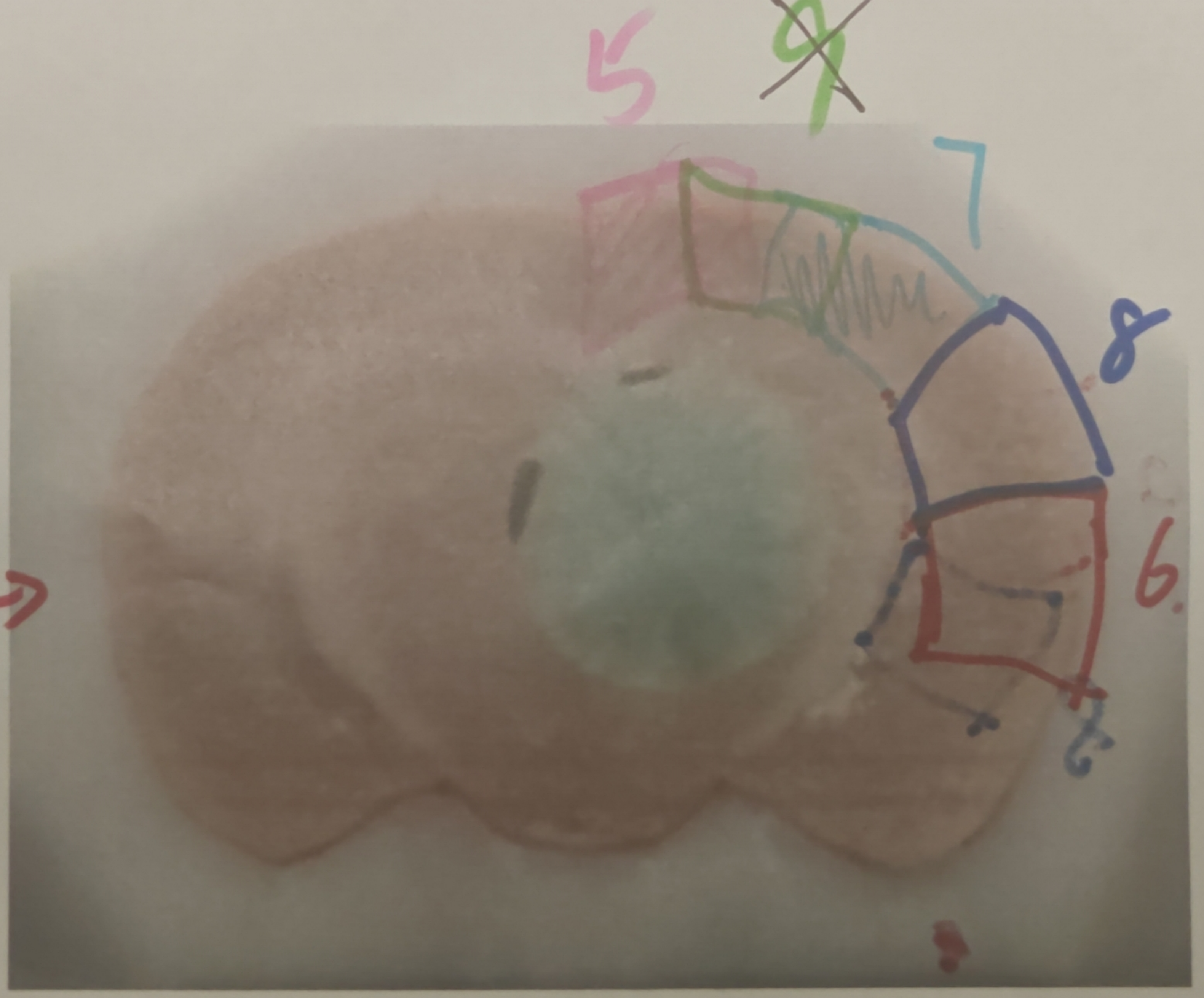

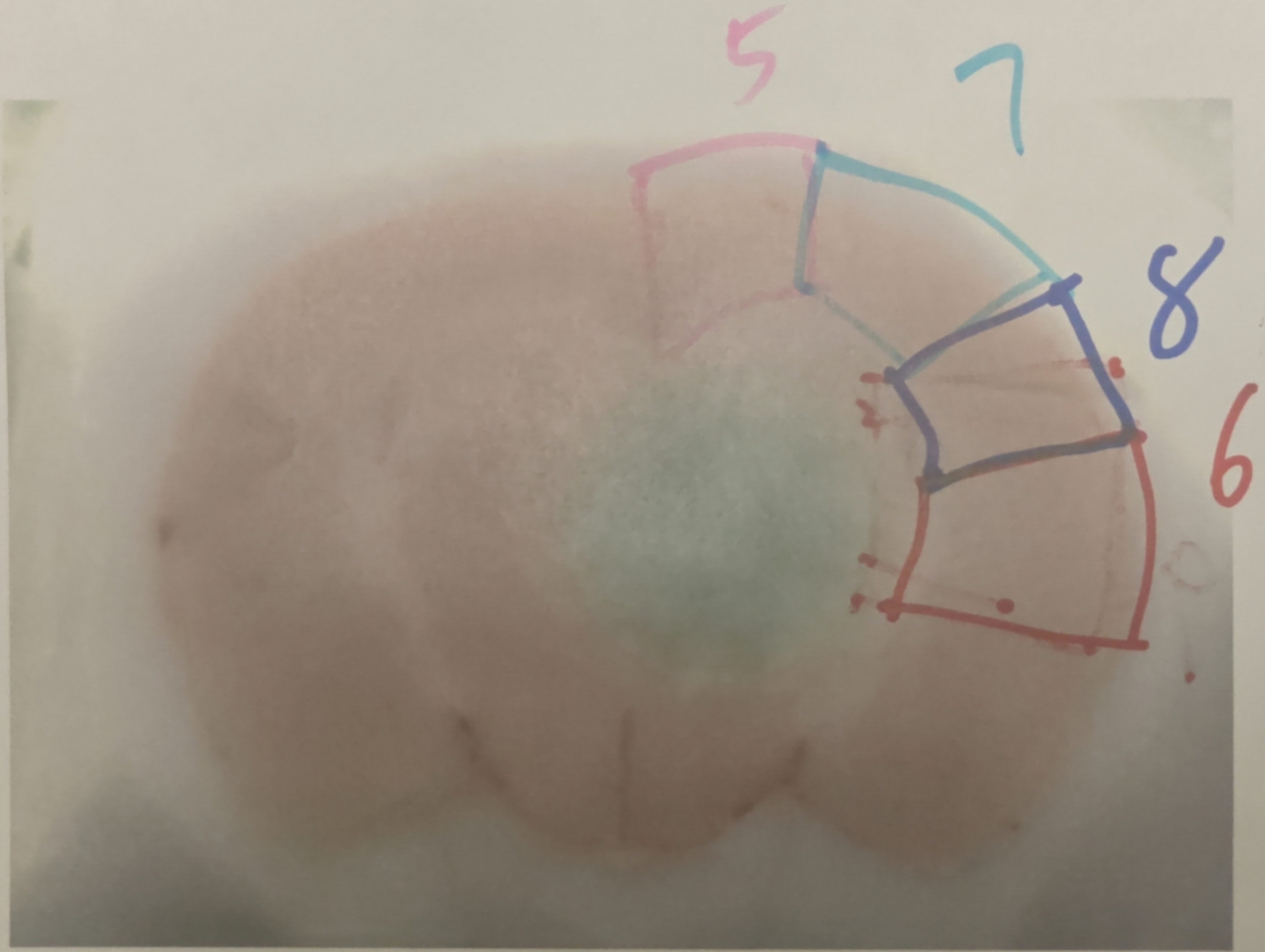


3.6

Cut 1

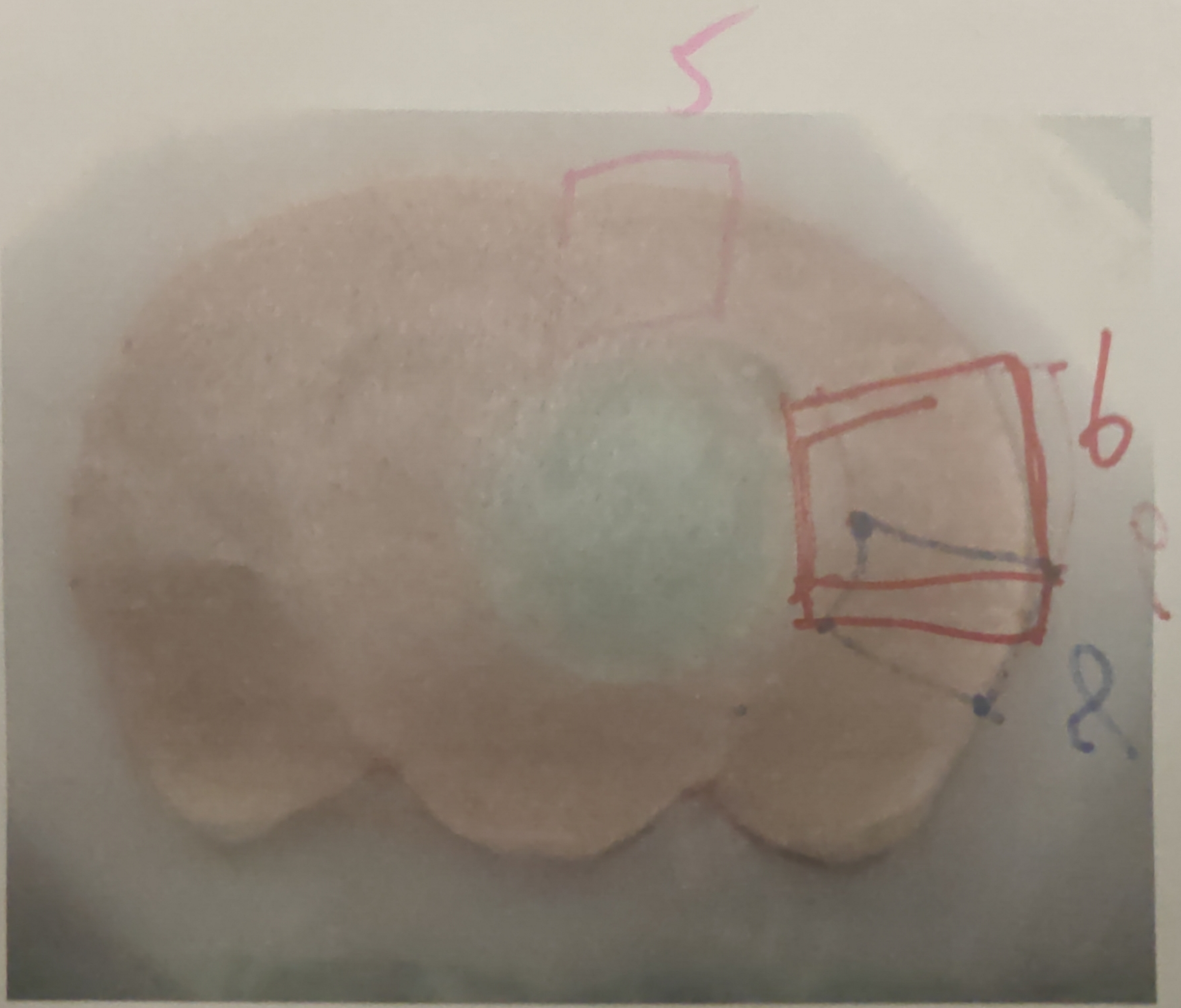


3.7

CL68

★★ see notes

Note this brain the punched sections are cut P→A, whereas previous A→P, 300μm diff across slices

3.8

CL 65

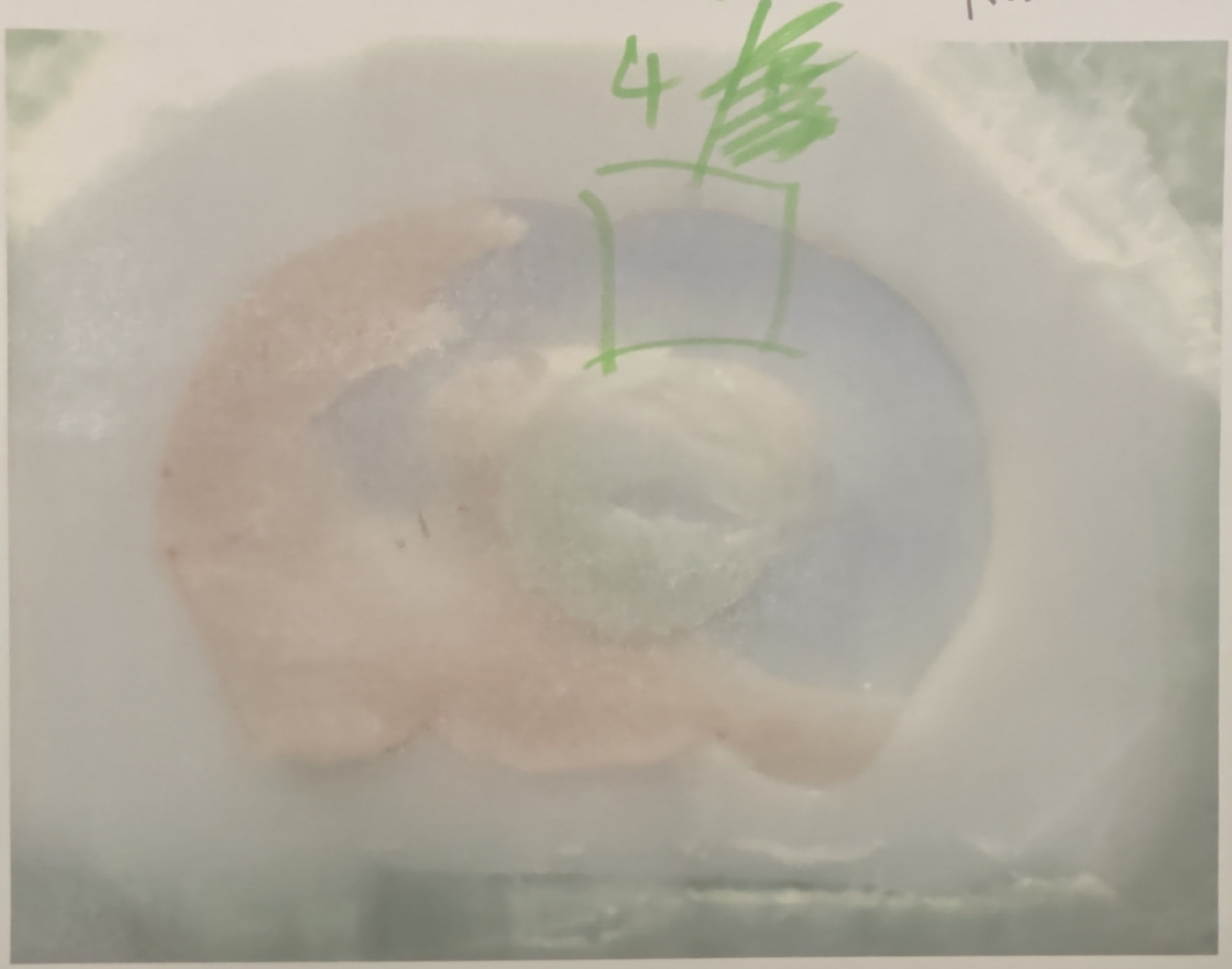


Missing 53–65 (1.2mm)

If last punched section is thick collect 4 not 5

(4 is till 63

5 starts on 67 in first brain)
