## Supplementary figures and images for "Conserved and specialized features of thalamocortical wiring revealed by single-cell projection mapping in mouse and marmoset"

### case04L_11L_19R_21R.jpg

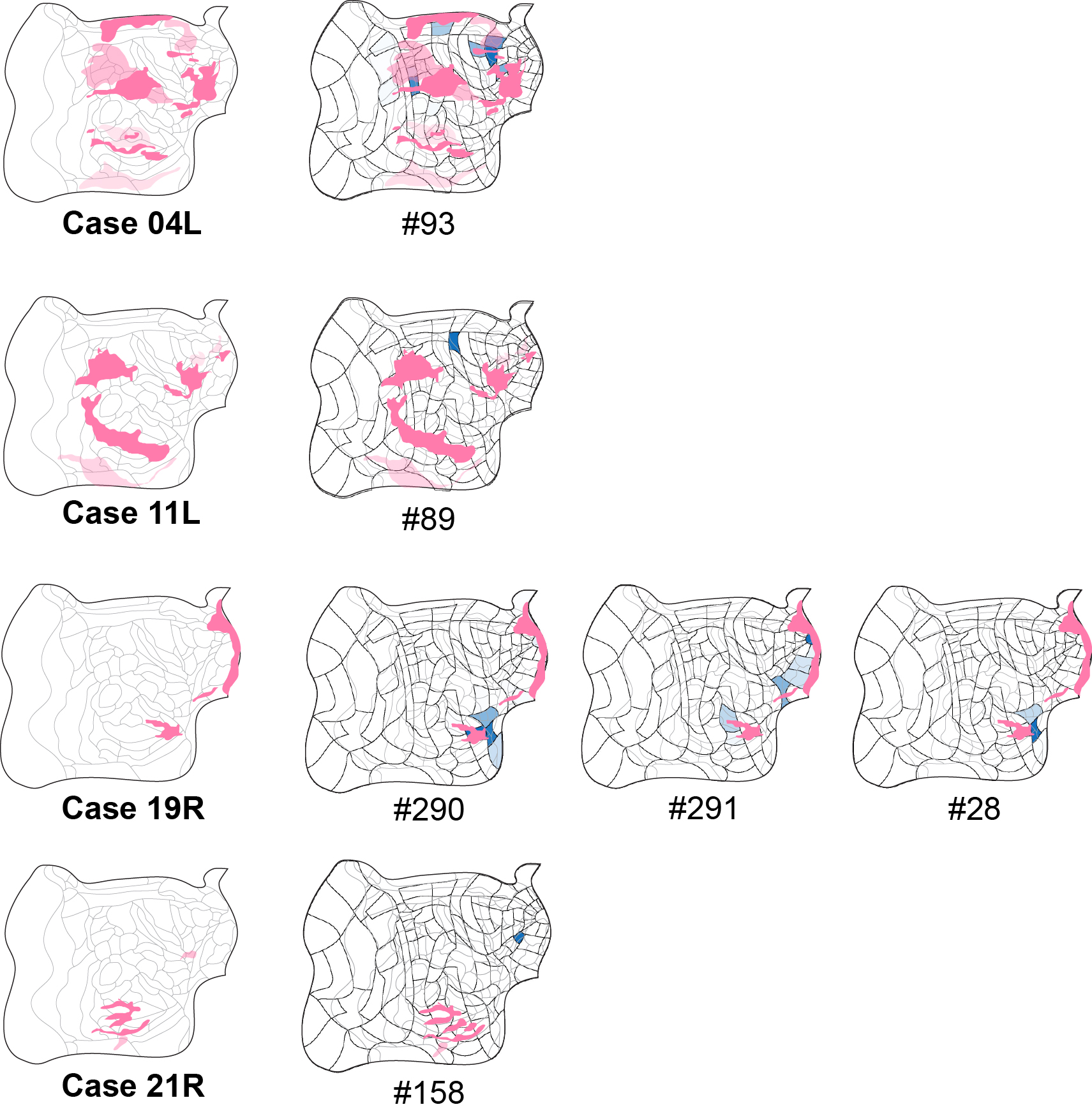

### case04L_11L_19R_21Rrandom.jpg

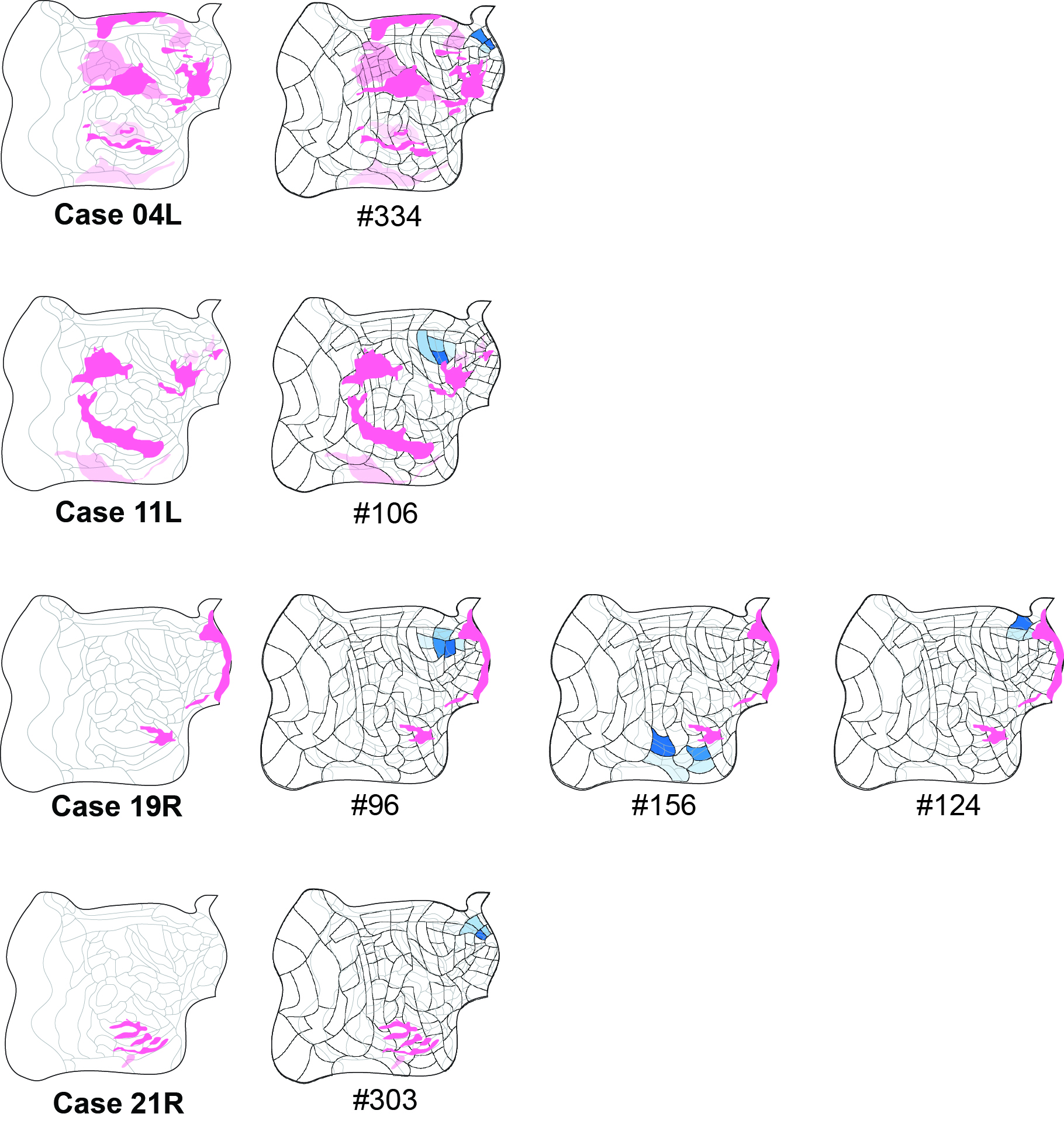

### case13L.jpg

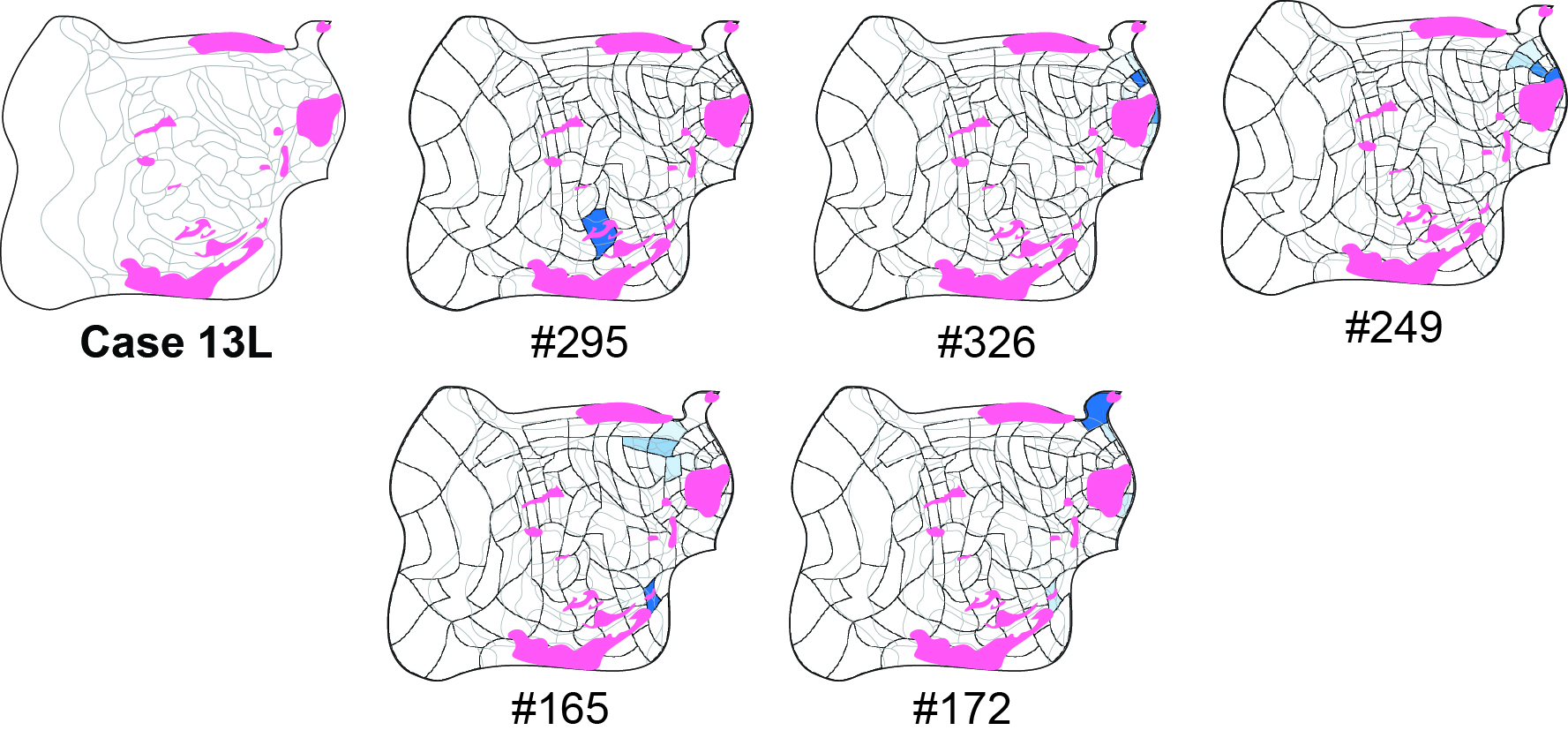

### case22L.jpg

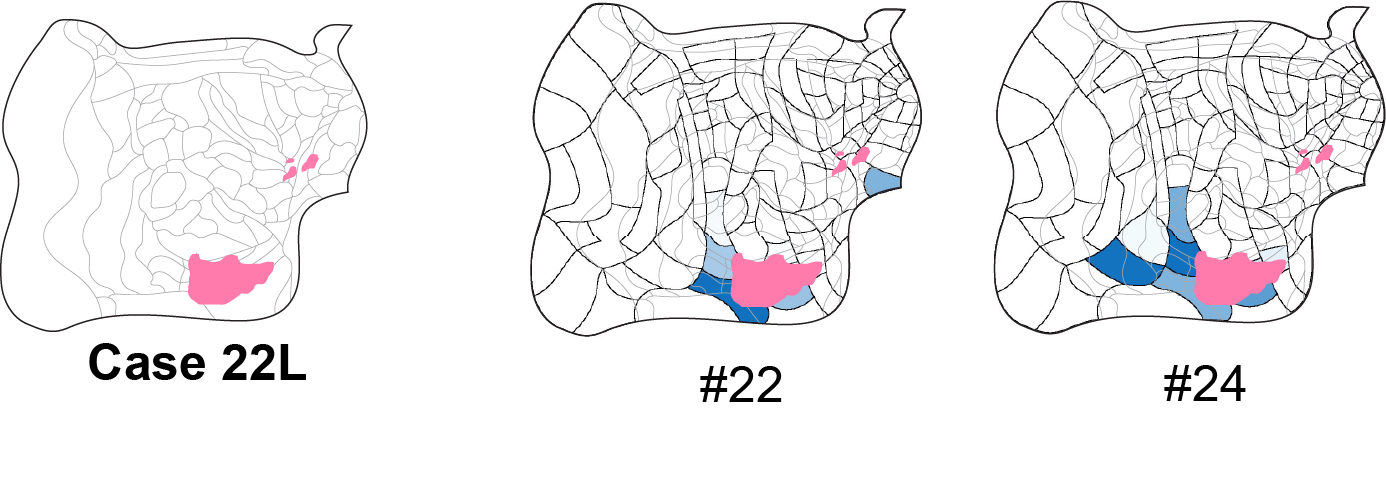

### case22Lrandom.jpg

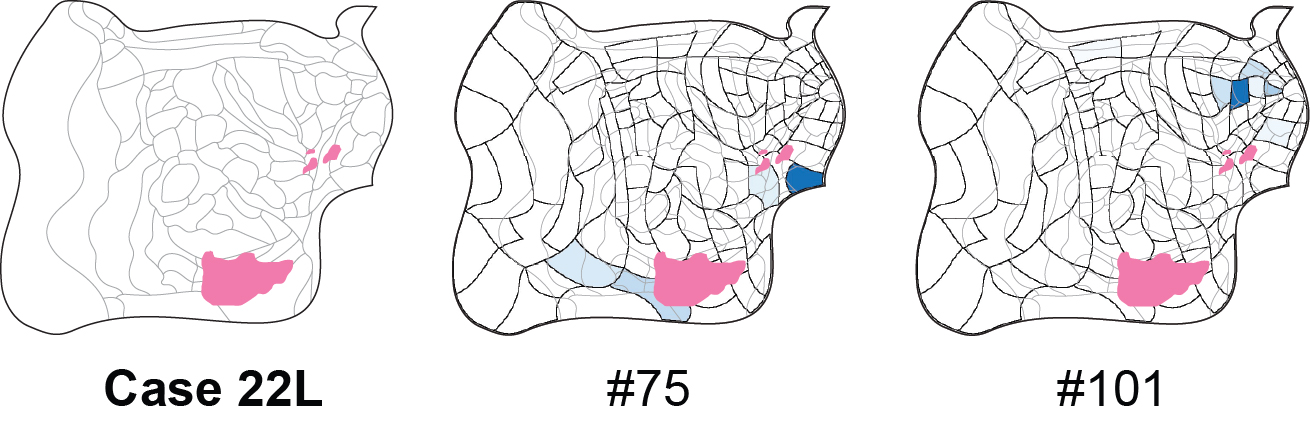

### case22R.jpg

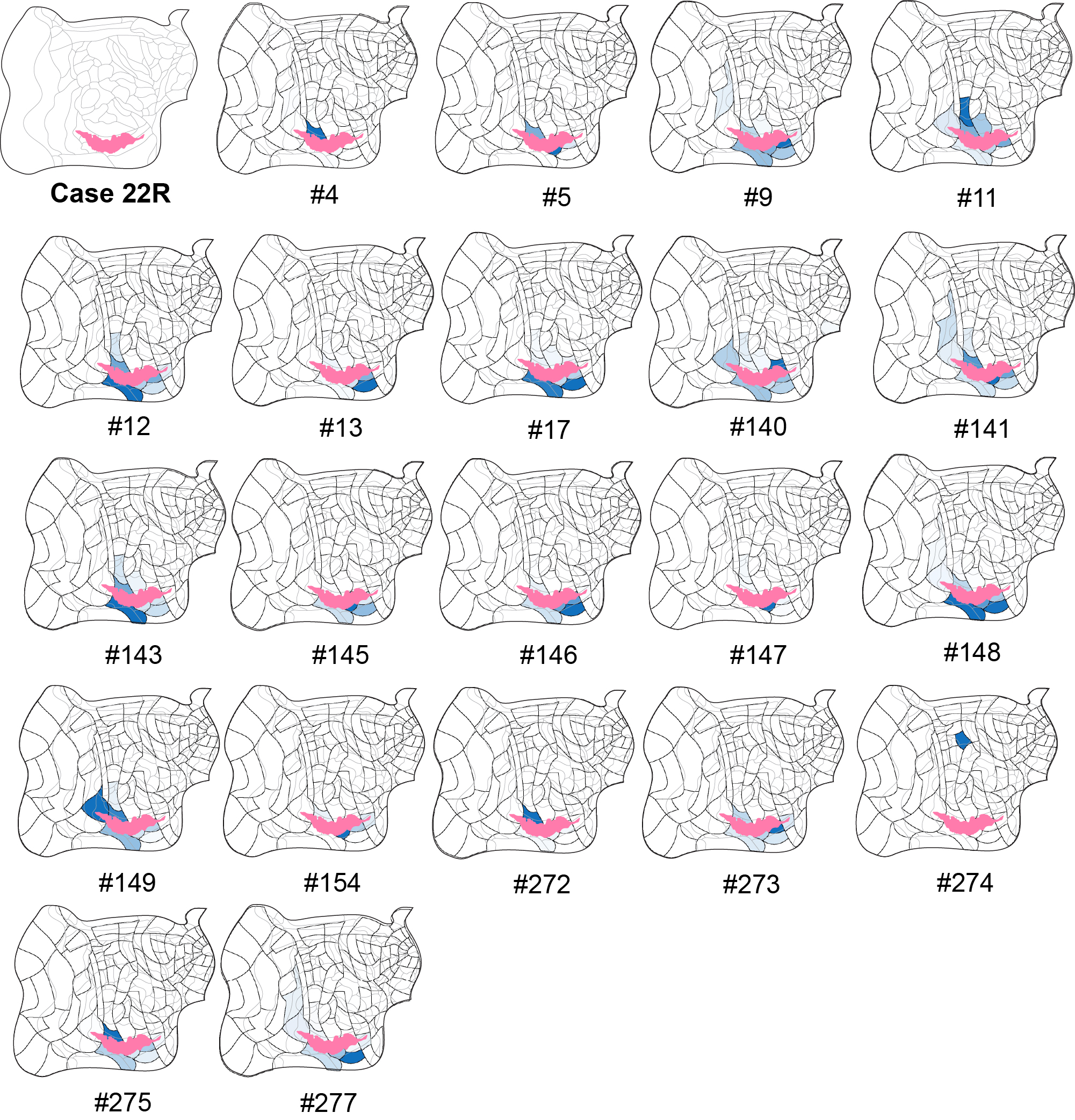

### case22Rrandom.jpg

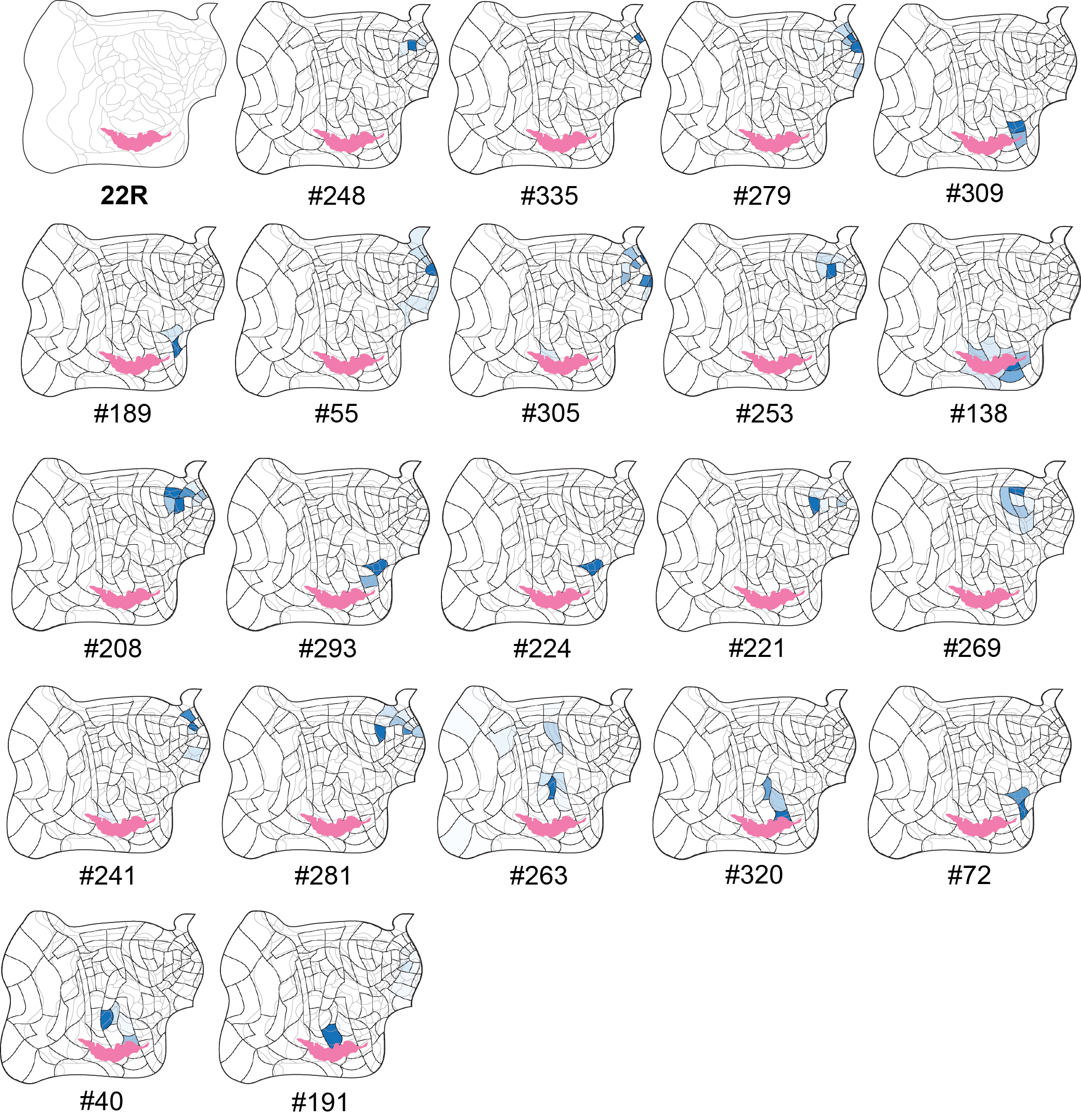

### case28L.jpg

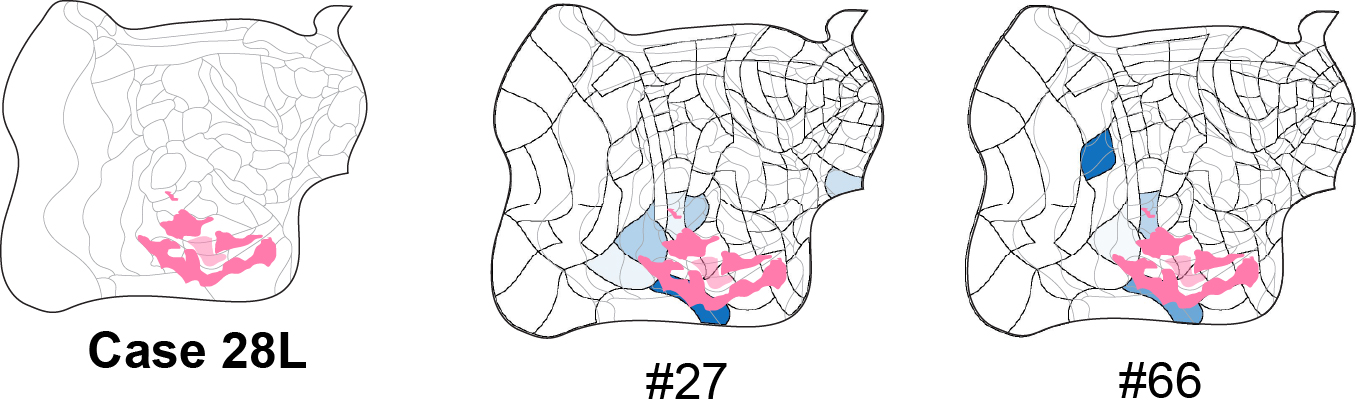

### case28L_random.jpg

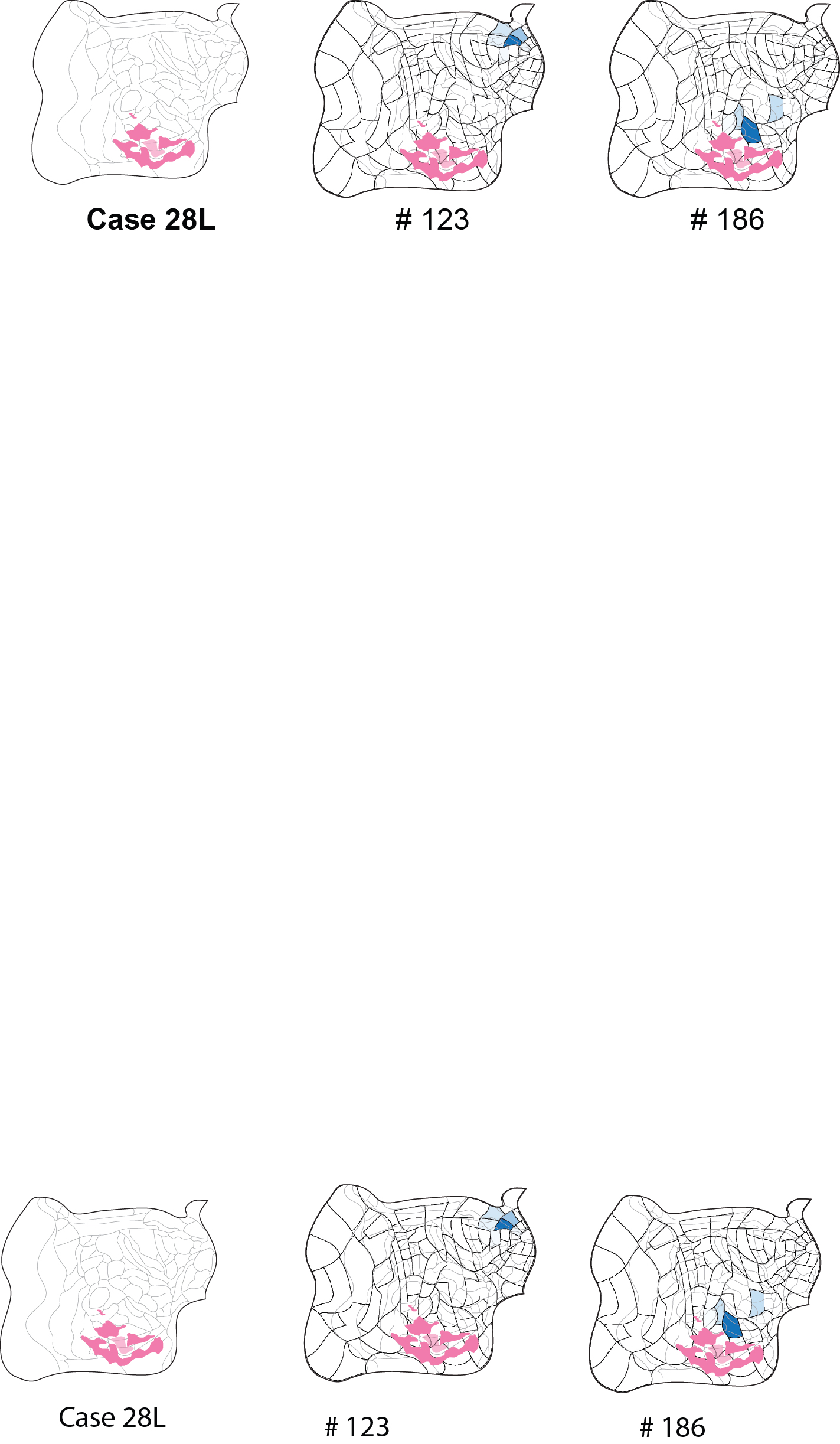
